# Experimental evolution of phosphomannomutase-deficient yeast reveals compensatory mutations in a phosphoglucomutase

**DOI:** 10.1101/2022.04.06.487342

**Authors:** Ryan C. Vignogna, Mariateresa Allocca, Maria Monticelli, Joy W. Norris, Richard Steet, Giuseppina Andreotti, Ethan O. Perlstein, Gregory I. Lang

## Abstract

The most common cause of human congenital disorders of glycosylation (CDG) are mutations in the phosphomannomutase gene *PMM2*, which affect protein *N*-linked glycosylation. The yeast gene *SEC53* encodes a nearly-identical homolog of human *PMM2*. We evolved 384 populations of yeast harboring one of two human-disease-associated alleles, *sec53*-V238M and *sec53*-F126L, or wild-type *SEC53*. We find that after 1,000 generations, most populations compensate for the slow-growth phenotype associated with the *sec53* human-disease-associated alleles. Through whole-genome sequencing we identify compensatory mutations, including known *SEC53* genetic interactors. We observe an enrichment of compensatory mutations in other genes whose human homologs are associated with Type 1 CDG, including *PGM1*, which encodes the minor isoform of phosphoglucomutase in yeast. By genetic reconstruction, we show that evolved *pgm1* mutations are dominant and allele-specific genetic interactors that restore both protein glycosylation and growth of yeast harboring the *sec53*-V238M allele. Finally, we characterize the enzymatic activity of purified Pgm1 mutant proteins. We find that reduction, but not elimination, of Pgm1 activity best compensates for the deleterious phenotypes associated with the *sec53*-V238M allele. Broadly, our results demonstrate the power of experimental evolution as a tool for identifying genes and pathways that compensate for human-disease associated alleles.

## INTRODUCTION

Protein glycosylation is an important cotranslational and posttranslational modification involving the attachment of glycans to polypeptides. These glycans play vital roles in protein folding, stability, activity, and transport (Varki, 2017). Glycosylation is one of the most abundant protein modifications, with evidence indicating that over 50% of human proteins are glycosylated (Roth et al., 2012; Wong, 2005). Despite this, we lack a global understanding of the complex pathologies involving protein glycosylation.

Congenital disorders of glycosylation (CDG) are a group of inherited metabolic disorders arising from defects in the protein glycosylation pathway, including *N*-linked glycosylation, *O*-linked glycosylation, and lipid/glycosylphosphatidylinositol anchor biosynthesis (Chang et al., 2018). *N*-linked CDG are further categorized into two groups based on the affected process: synthesis and transfer of glycans (Type 1) or processing of protein-bound glycans (Type 2) (Aebi et al., 1999). Mutations in the human phosphomannomutase 2 gene (*PMM2*) cause Type 1 CDG and are the most common cause of CDG (Ferreira et al., 2018). PMM2 forms a homodimer and catalyzes the interconversion of mannose-6-phosphate and mannose-1-phosphate (EC 5.4.2.8). Mannose-1-phosphate is then converted to GDP-mannose, a required substrate for *N*-linked glycosylation glycosyltransferases.

Two of the most common pathogenic *PMM2* variants found in humans are p.Val231Met p.Phe119Leu. The V231M protein is less stable and exhibits defects in protein folding (Citro et al., 2018; Silvaggi et al., 2006), whereas the F119L protein is stable but exhibits dimerization defects (Andreotti et al., 2015; Kjaergaard et al., 1999; Pirard et al., 1999). The budding yeast *Saccharomyces cerevisiae* contains a nearly identical phosphomannomutase enzyme encoded by the gene *SEC53. SEC53* is essential in yeast and expression of human *PMM2* rescues lethality (Hansen et al., 1997; Lao et al., 2019). Previously, one of us (E.O. Perlstein) constructed strains harboring the yeast-equivalent mutations of the most common human disease-associated *PMM2* alleles including V231M (*sec53*-V238M) and F119L (*sec53*-F126L) (Lao et al., 2019).

Using experimental evolution, we sought to identify compensatory mutations able to overcome glycosylation-deficiency. Previous studies have used experimental evolution of compromised organisms, to study evolutionary outcomes and identify compensatory mutations (Harcombe et al., 2009; Helsen et al., 2020; Laan et al., 2015; Michel et al., 2017; Moser et al., 2017; Szamecz et al., 2014). Experimental evolution allows for a more robust approach than traditional suppressor screens, enabling the identification of weak suppressors and compensatory interactions between multiple mutations (Cooper, 2018; LaBar et al., 2020).

Here we present a 1,000-generation evolution experiment of yeast models of PMM2-CDG. We evolved 96 populations of yeast with wild-type *SEC53*, 192 populations with the *sec53*-V238M allele, and 96 populations with the *sec53*-F126L allele for 1,000 generations. We sequenced 188 evolved clones to identify mutations that arose specifically in the populations harboring the *sec53* human-disease-associated alleles. We find an overrepresentation of mutations in genes whose human homologs are other Type 1 CDG genes, including *PGM1* (the minor isoform of phosphoglucomutase), the most commonly mutated gene in our experiment. We show that evolved mutations in *PGM1* restore protein *N*-linked glycosylation and alleviate the slow-growth phenotype caused by the *sec53*-V238M disease-associated allele. We performed genetic and biochemical characterization of the *pgm1* mutations and show that these mutations are dominant and allele-specific *SEC53* genetic-interactors.

## RESULTS

### Experimental evolution improves growth of yeast models of PMM2-CDG

Mutations in the phosphomannomutase 2 gene, *PMM2*, are the most common cause of congenital disorders of glycosylation (CDG). Previously Lao et al., 2019 constructed yeast models of PMM2-CDG by making the yeast-equivalent mutations for five human-disease-associated alleles in *SEC53*, the yeast ortholog of *PMM2*. These strains have growth rate defects that correlate with enzymatic activity and promoter strength. Increasing expression levels of the mutant *sec53* alleles two-fold, by replacing the endogenous promoter (p*SEC53*) with the *ACT1* promoter (*pACT1*), modestly improves the growth rate of *sec53* disease-associated alleles (Figure 1A).

**Figure 1.**
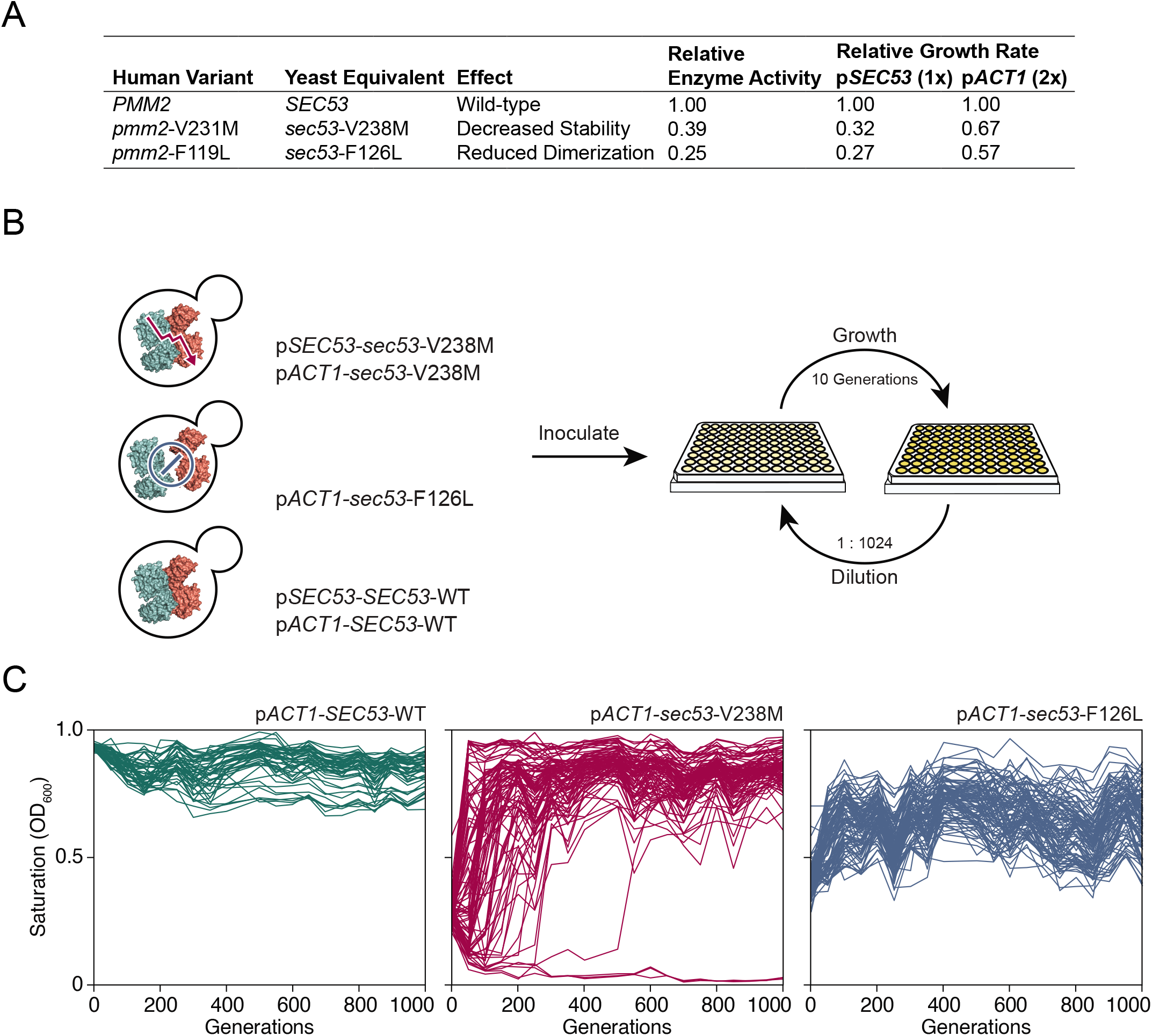
Experimental evolution of yeast models of congenital disorders of glycosylation. (A) Table of the various *sec53* alleles used in this study. In vitro enzymatic activities of Pmm2 are relative to wild-type Pmm2 and were previously reported (Pirard et al., 1999). Relative growth rates of yeast carrying various *SEC53* alleles were previously reported (Lao et al., 2019). (B) Diagram of the evolution experiment. Yeast carrying *sec53* mutations implicated in human disease were used to initiate replicate populations in 96-well plates: 96 populations of each mutant genotype and 48 populations of each wild type genotype. Populations were propagated in rich glucose media, unshaken, for 1,000 generations. (C) Single time-point OD_600_ readings of populations were taken every 50 generations during the evolution experiment as a measure of growth rate. Each line represents one population.

We chose to focus on *sec53*-V238M and *sec53*-F126L because they are two of the most common disease alleles in humans and because the mutations have distinct and well-characterized effects on protein structure and function (Figure 1A) (Andreotti et al., 2015; Briso-Montiano et al., 2022; Citro et al., 2018; Kjaergaard et al., 1999; Pirard et al., 1999; Silvaggi et al., 2006). We evolved 96 haploid populations of each mutant *sec53* genotype (p*SEC53-sec53-V238M*, *pACT1-sec53-V238M, pACT1-sec53-F126L*) and 48 haploid populations of each wild-type *SEC53* genotype (p*SEC53*-*SEC53*-WT, *pACT1-SEC53-WT*) for 1,000 generations in rich glucose medium (Figure 1B). Strains with the p*SEC53*-*sec53*-F126L allele grew too slowly to keep up with a 1:2^10^ dilution every 24 hours. Every 50 generations we measured culture density (single time-point OD_600_) for each population as a proxy for growth rate (Figure 1C; Figure 1 — figure supplement 1).

Populations with the *pACT1-sec53-V238M* allele show a range of dynamics, with some populations reaching maximum saturation (OD_600_ ≈ 1.0) early in the experiment, while some do not even by Generation 1,000 (Figure 1C). Three *pACT1-sec53-V238M* populations went extinct before Generation 200. While saturation levels of *pACT1-sec53-F126L* populations also increased over the course of the evolution experiment, none reached an OD_600_ close to 1.0, indicating that these populations are likely less-fit than the evolved *pACT1-sec53-V238M* populations. Each p*SEC53*-*sec53*-V238M population started the evolution experiment with growth rates comparable to *SEC53-WT* populations, indicating compensatory mutation(s) likely arose in the starting inocula (Figure 1 — figure supplement 1). We therefore exclude these populations from subsequent analyses. Together, our data show that the PMM2-CDG yeast models acquired compensatory mutations throughout the evolution experiment and that the extent of compensation depends on the specific disease-associated allele.

### Putative compensatory mutations are enriched for other Type 1 CDG-associated homologs

To identify the compensatory mutations in our evolved populations, we sequenced single clones from 188 populations isolated at Generation 1,000 to an average sequencing depth of ~50x. These included 91 *pACT1-sec53-V238M* clones, 36 p*ACT1*-*sec53*-F126L clones, 32 p*SEC53*-*SEC53*-WT clones, 11 *pACT1-SEC53-WT* clones, and 18 of the initially-suppressed p*SEC53-sec53-*V238M clones (Supplemental Dataset 1). Autodiploids are a common occurrence in our experimental system (Fisher et al., 2018; Johnson et al., 2021). We attempted to avoid sequencing autodiploids by screening our evolved populations for sensitivity to benomyl, an antifungal agent that inhibits growth of *S. cerevisiae* diploids more severely than haploids (Venkataram et al., 2016).

We find that the average number of de novo mutations (SNPs and small indels) varied between *SEC53* genotypes, with the p*ACT1*-*sec53*-F126L clones accruing more mutations per clone (8.14 ± 1.00, 95% CI) compared to *pACT1-sec53-V238M* (5.76 ± 0.54) and *SEC53*-WT clones (combined p *SEC53*-*SEC53*-WT and p*ACT1*-*SEC53*-WT) (6.49 ± 0.90) (Figure 2 — figure supplement 1). Despite screening for benomyl sensitivity, most of the sequenced clones (131/188) appear to be autodiploids, with evidence of multiple heterozygous loci. In addition, we detect several copy number variants and aneuploidies shared between independent populations (Figure 2 — figure supplement 2).

We identified putative adaptive targets of selection as genes with more mutations than expected by chance across replicate clones. Some of these common targets of selection are not specific to *sec53*. For example, mutations in negative regulators of Ras (*IRA1* and *IRA2*) are found in clones from each experimental group and are also observed in other laboratory evolution experiments across a wide range of conditions (Figure 2A) (Fisher et al., 2018; Gresham and Hong, 2014; Johnson et al., 2021; Kvitek and Sherlock, 2013; Lang et al., 2013; Venkataram et al., 2016).

**Figure 2.**
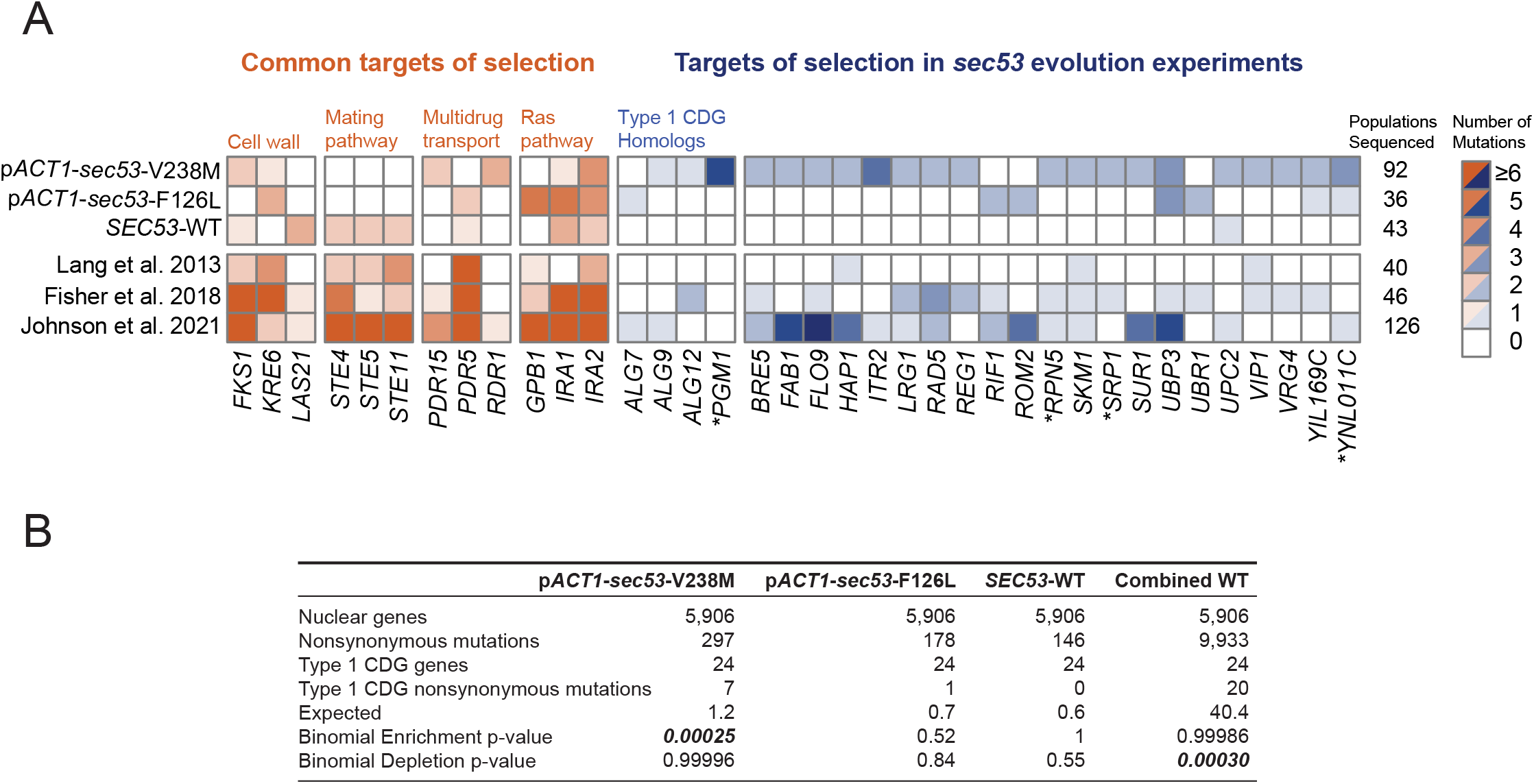
Mutations in Type 1 CDG homologs are enriched in *sec53*-V238M populations. (A) Heatmap showing number of nonsynonymous mutations per gene that arose in the evolution experiment. Genes with 2 or more unique nonsynonymous mutations (and each Type 1 CDG homolog with at least 1 mutation) are shown. p*SEC53-SEC53-WT* and *pACT1-SEC53-WT* are grouped as *“SEC53-WT”.*For comparison we show data from previously reported evolution experiments where the experimental conditions were identical to the conditions used here, aside from strain background and experiment duration (bottom 3 rows). Commonly mutated pathways are grouped for clarity. Asterisks (*) indicate previously-known *SEC53* genetic interactors (Costanzo et al., 2016; Kuzmin et al., 2018). (B) Quantification of enrichment and depletion of mutations in Type 1 CDG homologs in the listed evolution experiments. Lang et al., Fisher et al., Johnson et al., and *SEC53*-WT populations comprise “Combined WT”.

We identified putative *sec53*-compensatory targets as genes mutated exclusively in *pACT1-sec53-V238M* and/or *pACT1-sec53-F126L* populations (Figure 2A). The most frequently mutated genes among p*ACT1*-*sec53*-F126L clones are common targets in other evolution experiments such as *IRA1, IRA2*, and *KRE6*, suggesting that mutations capable of compensating for Sec53-F126L dimerization defects are rare or not easily accessible. In contrast, while recurrently mutated genes in *pACT1-sec53-V238M* populations include common targets, we also identified a number of unique targets of selection, most notably in homologs of other Type 1 CDG-associated genes. We find an enrichment of mutations in CDG homologs in *pACT1-sec53-V238M* clones (Figure 2B; Figure 2 — figure supplement 3).

Across all sequenced populations we identified ~620 nonsynonymous mutations. As there are nearly 6,000 yeast genes, the probability of any given gene receiving even a single nonsynonymous mutation is low. We are therefore well-powered to detect enrichment of mutations, but underpowered to detect depletion of mutations. To gain statistical power we aggregated our *SEC53*-WT data with three other large data sets using similar strains and propagation regimes (Fisher et al., 2018; Johnson et al., 2021; Lang et al., 2013). In this aggregate dataset, we observe a statistically-significant depletion of mutations in Type 1 CDG homologs, suggesting that they are typically under purifying selection in experimental evolution (Figure 2B).

One of the CDG homologs, *PGM1*, is the most frequently mutated gene among *pACT1-sec53*-V238M clones. *PGM1* encodes the minor isoform of phosphoglucomutase in yeast (Bevan and Douglas, 1969). We only find *PGM1* mutations in *pACT1-sec53-V238M* populations which suggests their compensatory effects are unique to the *sec53*-V238M mutation and not glycosylation-deficiency in general. Each of the five mutations in *PGM1* are missense mutations, rather than frameshift or nonsense. This is consistent with selection acting on alteration-of-function rather than loss-of-function (posterior probability of non-loss-of-function = 0.96, see Methods).

To identify the compensatory mutation(s) that arose prior to the start of the evolution experiment in the suppressed p*SEC53-sec53-*V238M populations, we analyzed the 18 sequenced clones. For each clone, we find both wild-type *SEC53* and *sec53*-V238M alleles among sequencing reads. This could result from integration or maintenance of the covering plasmid. No sequencing reads align to the *URA3* locus of our reference genome, suggesting that the plasmid marker was lost, as expected following counterselection.

### Compensatory mutations restore growth

In order to validate putative compensatory mutations we reconstructed evolved mutations in *PGM1* and *ALG9*, two genes whose human homologs are implicated in Type 1 CDG (Frank et al., 2004; Timal et al., 2012). These heterozygous evolved mutations were constructed in a homozygous *pACT1-sec53-V238M* diploid background (note that although each *pgm1* and *alg9* mutation assayed here arose in haploid-founded populations, they arose as heterozygous mutations in autodiploids). We quantified the fitness effect of each mutation using a flow cytometry-based fitness assay, competing the reconstructed strains against a fluorescently-labeled, diploid version of the p*ACT1*-*SEC53*-WT ancestor.

The p*ACT1*-*sec53*-V238M ancestor has a fitness deficit of −25.86 ± 2.3% (95% CI) relative to wild-type. We find that each of the five evolved *pgm1* mutations is compensatory in the *pACT1-sec53-V238M* background. The fitness effects of the *sec53/pgm1* double-mutants range between −19.10 ± 1.07% and −9.84 ± 0.77% (i.e., the *pgm1* mutations confer a fitness benefit between 6.76% and 16.02% in the *pACT1-sec53-V238M* background) (Figure 3A). We also find that the evolved *alg9*-S230R mutation is compensatory, as the *sec53*/*alg9* double-mutant has a fitness effect of −17.89 ± 0.59% (Figure 3 — figure supplement 1). We compared the fitness of these reconstructed double-mutants to the fitness of the evolved clones which carry 3-9 additional SNPs. In each case the evolved clones were more fit than the reconstructed strain, except for the evolved clone containing the *pgm1*-D295N mutation (Figure 3 — figure supplement 1). Thus, while the *alg9* and *pgm1* mutations compensate for the *sec53*-V238M defect, other mutations contribute to the fitness of these evolved clones.

**Figure 3.**
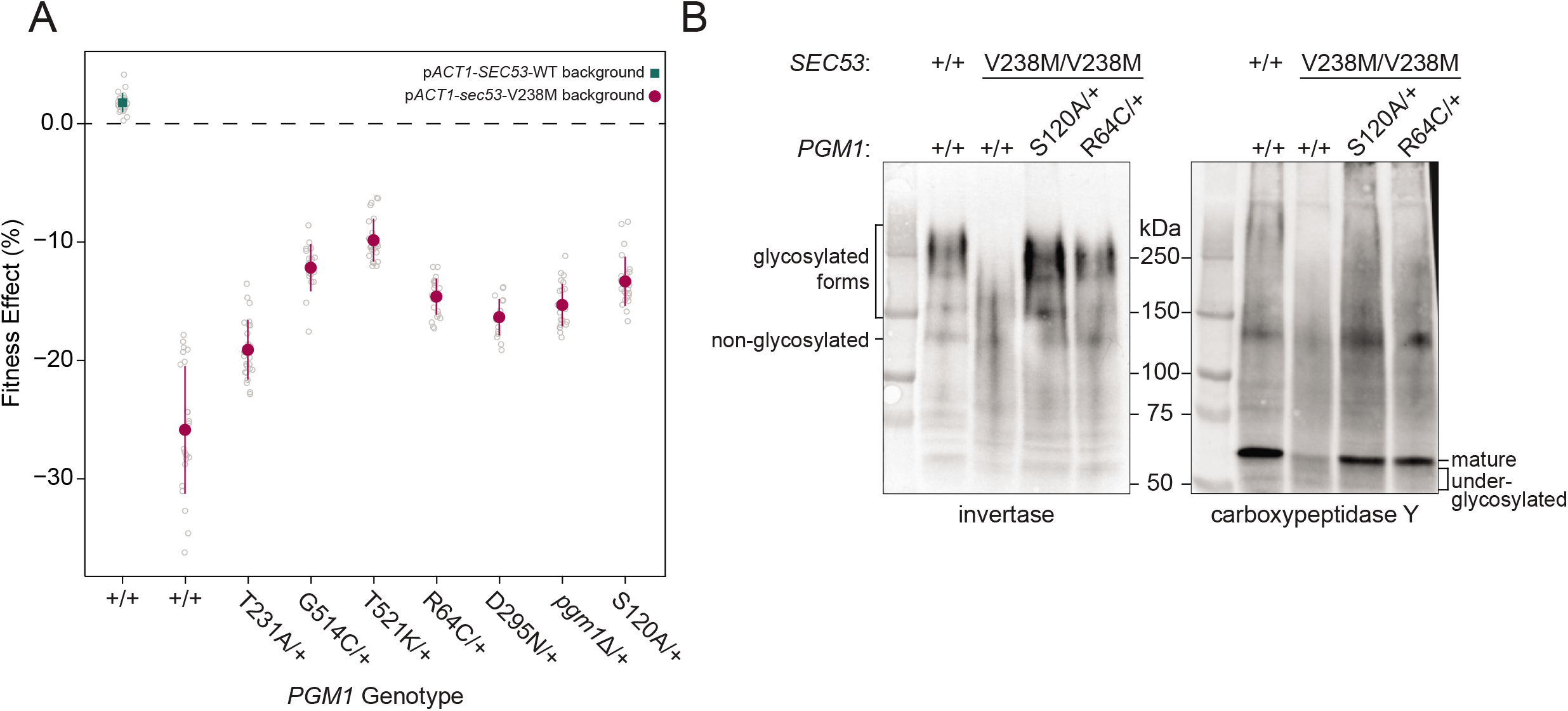
Evolved mutations rescue fitness and protein glycosylation defects of *pACT1-sec53-V238M*. (A) Average fitness effects and standard deviations of reconstructed heterozygous *pgm1* mutations. Fitness effects were determined by competitive fitness assays against a fluorescently-labeled version of the diploid *pACT1-SEC53-WT* ancestor. Replicate measurements are plotted as gray circles. Pairs of *pgm1* fitness effects with non-statistically-significant differences: R64C-D295N, R64C-*pgm1*Δ, R64C-S120A, D295N-*pgm1*Δ, G514C-T521K, G514C-S120A, and *pgm1*Δ-S120A (*df*=194, *F*=208.1, each p>0.05, one-way ANOVA with Tukey post hoc test). (B) Western blots of invertase (left) and carboxypeptidase Y (right) from ancestral and reconstructed strains. In panels A and B, plus signs (+) indicate wild-type alleles. Genotypes are either homozygous wild-type (+/+), homozygous mutant (mutation/mutation), or heterozygous (mutation/+).

We constructed and assayed the fitness effects of the *pgm1* mutations in diploid *pACT1-sec53-F126L* and p*ACT1-SEC53*-WT backgrounds. Each p*ACT1*-*sec53*-F126L strain was too quickly outcompeted by the reference strain to measure fitness, suggesting that the compensatory effects of the evolved *pgm1* mutations are specific for the *sec53*-V238M mutation or any fitness improvements in the p*ACT1*-*sec53*-F126L background are too minimal to detect. Each *pgm1* mutation is nearly neutral in the *pACT1-SEC53-WT* background (Figure 3 — figure supplement 2).

To determine if loss-of-function (LOF) of *PGM1* would phenocopy evolved mutations, we deleted *PGM1* (*pgm1*Δ) in the diploid *pACT1-sec53-V238M* background and again measured fitness. We find that the heterozygous *pgm1*Δ mutation improves fitness (−15.28 ± 0.76%), indicating that LOF of *PGM1* is compensatory in the conditions of the evolution experiment (Figure 3A). It is not immediately clear, then, why we only identify missense mutations in *PGM1*. Each of the mutated Pgm1 residues are located around the active site of the enzyme, based on predicted protein structure (Jumper et al., 2021; Stiers and Beamer, 2018). It could be that the evolved *pgm1* mutations are loss-of-function given this clustering but maintaining protein expression provides an ancillary benefit. To account for this possibility, we constructed a catalytically-dead allele of *PGM1* by mutating the enzyme’s catalytic serine (*pgm1*-S120A) (Stiers et al., 2017a). We find that *pgm1*-S120A improves *pACT1-sec53-V238M* fitness (−13.32 ± 0.88%) and this does not significantly differ from *pgm1Δ* (Figure 3A), indicating that *PGM1*LOF is compensatory regardless of if that arises from loss of coding sequence or enzyme activity. We also measured fitness of homozygous *pgm1* mutations (Figure 3 — figure supplement 3). In all cases the fitness effect of heterozygous and homozygous *pgm1* mutants are not substantially different from each other, indicating that, by this assay, the evolved *pgm1*mutations are dominant in the *pACT1-sec53-V238M* background, as are the *pgm1*Δ and *pgm1*-S120A mutations.

### Compensatory mutations restore protein glycosylation

We have established that *pgm1* mutations compensate for the growth rate defect of the *sec53*-V238M allele. To determine whether the evolved pgm1 mutants also compensate for the molecular defects in *N*-linked glycosylation we examined two representative yeast glycoproteins, invertase and carboxypeptidase Y (CPY), in our reconstructed *pgm1* strains. Invertase forms both a non-glycosylated homodimer (120 kDa) and a secreted homodimer that is heavily glycosylated (approximate range of 140-270 kDa) (Gascón et al., 1968; Zeng and Biemann, 1999). Mature CPY only exists in its glycosylated form, with a molecular weight of 61 kDa (Hasilik and Tanner, 1978).

The ancestral *pACT1-sec53-V238M* strain shows underglycosylation of invertase and CPY. We find a reduced abundance of the higher-molecular-weight glycosylated form of invertase and mature CPY, accompanied by the appearance of underglycosylated forms of these proteins (Figure 3B). We find that evolved *pgm1* mutations, *pgm1*Δ, and *pgm1*-S120A each restore glycosylation of these proteins to near-wild-type levels (Figure 3B, Figure 3 — figure supplements 4,5). We do not observe rescue of protein glycosylation in *pACT1-sec53*-F126L strains carrying *pgm1* mutations (Figure 3 — figure supplement 6). Together, these suggest that *pgm1*-mediated rescue of *pACT1-sec53-V238M* fitness is due to restoration of protein glycosylation and the effect is specific for the *pACT1-sec53-V238M* background.

### Compensatory *PGM1* mutations are dominant suppressors of *sec53*-V238M

To further demonstrate that *pgm1*-mediated compensation is specific for *sec53*-V238M, we performed a genetic analysis by dissecting tetrads from strains that are heterozygous for both *sec53* and *pgm1*. In the absence of a compensatory mutation, we expect 50% large colonies and all tetrads showing a 2:2 segregation of colony size due to the single segregating locus (*SEC53/sec53*). However, if an evolved mutation is compensatory then we expect 75% large colonies with three types of segregation patterns: 2:2, 3:1, and 4:0, large to small colonies, respectively. These three segregation patterns are expected to follow a 1:4:1 ratio assuming no genetic linkage (Figure 4A). For each genetic test, we dissected ten tetrads and performed a log likelihood test to determine whether the segregation pattern is more consistent with genetic suppression than non-suppression (Figure 4 — source data 1).

**Figure 4.**
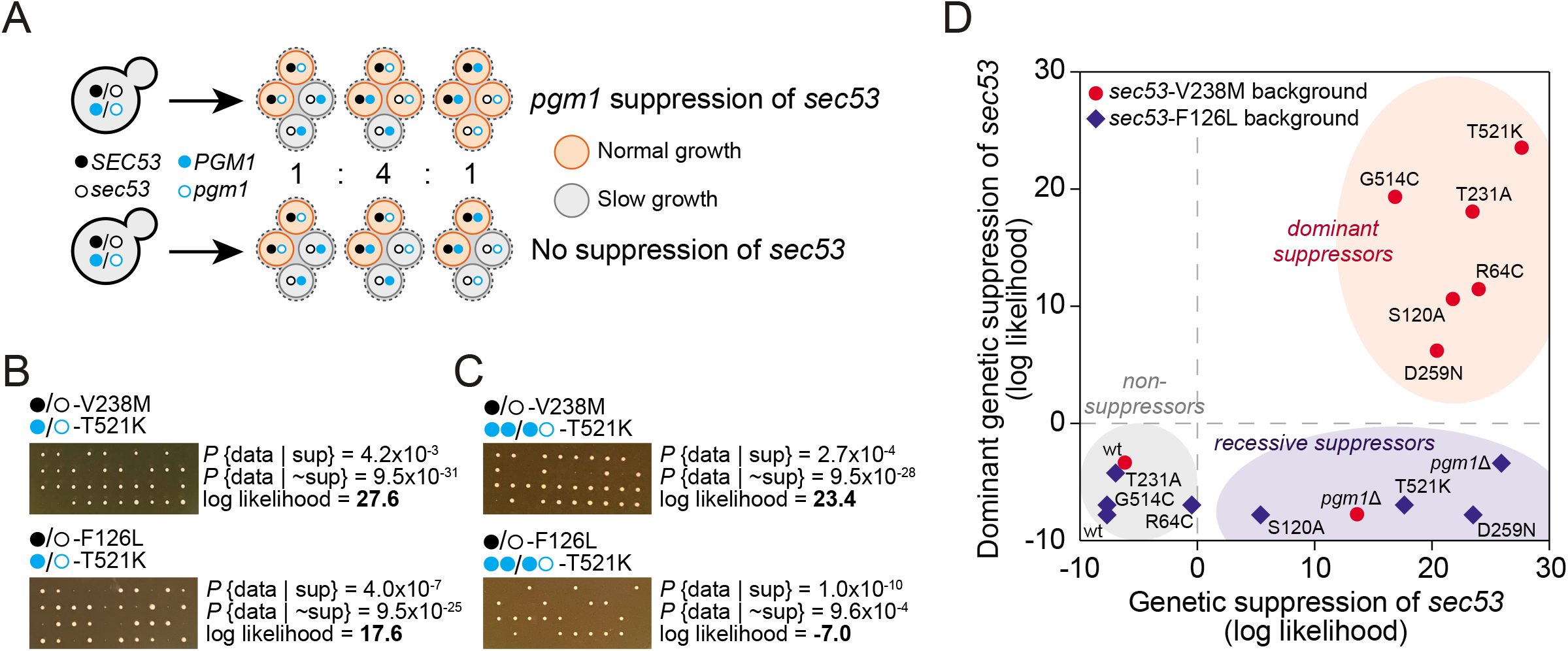
*pgm1* mutations are dominant suppressors of *sec53*-V238M. (A) Diagram of possible spore genotypes in the tetrad dissections. Heterozygous (e.g. *SEC53/sec53 PGM1/pgm1*) diploid strains could produce one of three tetrad genotypes based on allele segregation. (B) Example of *pgm1*-T521K dissections in a *sec53*-V238M background (top) and a *sec53*-F126L background (bottom). Listed are the probability and log likelihood of suppression, based on the ratio of normal growth (large colonies) to slow growth (no colonies or small colonies). (C) Example of *PGM1*::*pgm1*-T521K dissections in a *sec53*-V238M background (top) and *sec53*-F126L background (bottom). (D) Plot of recessive suppression versus dominant suppression of the *pgm1* mutations in the *sec53*-V238M background (red circles) and *sec53*-F126L background (blue diamonds), based on the tetrad dissections.

We find each of the evolved *pgm1* mutations compensates for the V238M allele of *SEC53* (Figure 4 — figure supplement 1). For example, dissections of a strain heterozygous for *pACT1-sec53-V238M* and *pgm1*-T521K show 78% large colonies (one 4:0 and nine 3:1, with a log likelihood ratio of 27.6, Figure 4B). These genetic tests corroborate the results from the fitness assays by showing that all five evolved *pgm1* mutations, the *pgm1Δ* and *pgm1*-S120A mutations, as well as the evolved *alg9* mutation all suppress *sec53*-V238M (Figure 4 — figure supplement 1). We next assayed each of the *pgm1* mutations in a *sec53*-F126L background. In contrast to the fitness assays, we find several *pgm1* alleles (T521K, D295N, S120A, and *pgm1*Δ) that suppress *sec53*-F126L (Figure 4 — figure supplement 1). It is worth noting, however, that the degree of compensation is less than in the *sec53*-V238M background based on colony sizes.

To test if *pgm1* mutations are dominant, we integrated a second wild-type copy of *PGM1*to each of the strains such that each haploid spore will contain either two wild-type copies of *PGM1* (*PGM1::PGM1*) or both wild-type *PGM1* and mutant *pgm1* (*PGM1::pgm1*). We find that the five evolved *pgm1* mutations and the *pgm1*-S120A mutations, but not *pgm1*Δ, are dominant suppressors of *sec53*-V238M background in the genetic assay (Figure 4C, 4D; Figure 4 — figure supplement 2). For the *sec53*-F126L background, we find no dominant suppressors.

### Reduction of Pgm1 activity alleviates deleterious effect of PMM2-CDG in yeast

To determine how the evolved mutations alter Pgm1 enzymatic activity, we cloned mutant and wild-type *PGM1* alleles into bacterial expression vectors, purified recombinant enzyme, and assayed phosphoglucomutase activity using a standard coupled enzymatic assay (Figure 5A; Figure 5 — figure supplement 1). We find that mutant Pgm1 have variable activities, ranging from near-wild type (Pgm1-T231A) to non-detectable (Pgm1-R64C and Pgm1-D295N) (Figure 5B, 5C). The near complete loss of activity of Pgm1-D295N is unsurprising since D295 coordinates a Mg^2+^ ion that plays an essential role during catalysis (Stiers et al., 2016) and we verify that all the active forms show very low activity in the presence of EDTA (Figure 5 — figure supplement 2). Thermostability did not differ between enzymes with detectable activity (Figure 5 — figure supplement 3).

**Figure 5.**
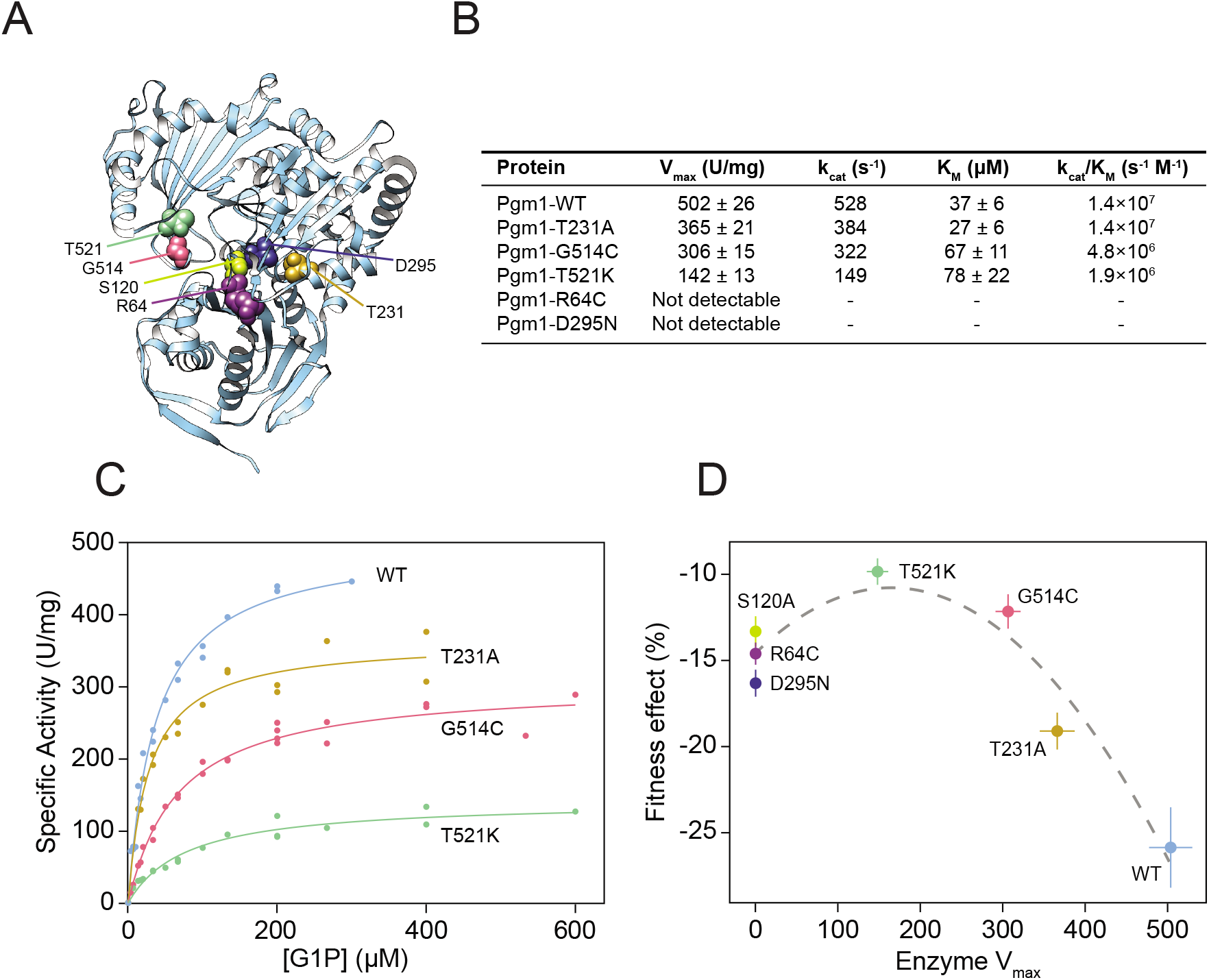
Complete loss of Pgm1 activity overshoots a fitness optimum. (A) AlphaFold structure of *S. cerevisiae* Pgm1 (Jumper et al., 2021). Mutated residues shown as spheres. (B) Kinetic parameters of recombinant Pgm1 enzymes as determined by a coupled enzymatic assay. Values ± 95% confidence intervals. (C) Michaelis-Menten curves of wild-type and mutant Pgm1. Replicate measurements are plotted as circles. (D) Pgm1 enzyme V_max_ versus the corresponding allele’s fitness effect in the diploid *pACT1-sec53-V238M* background (as shown in Figure 3A). Horizontal and vertical error bars represent 95% confidence intervals. Best fit regression shown as a dashed curve (y=14.583+0.04613x-0.00014x^2^, R^2^=0.907, *df*=4, *F*=37.34, p=0.0258, ANOVA). Note that we did not measure Pgm1-S120A activity and assume null activity.

Comparison of the kinetic parameters of active mutant enzymes shows a reduction of maximal velocity (V_max_) between 27%-72% relative to wild-type Pgm1. Based on the apparent Michaelis constant (K_m_), substrate affinities of Pgm1-G514C and Pgm1-T521K are lower than wild-type Pgm1 (*t*=4.30 and 3.14, *df*=35.78 and 20.47, p=0.0001 and 0.0051, respectively, Welch’s modified t-test). Pgm1-T231A shows an increased affinity compared to wild-type Pgm1, but this difference is not statistically significant (*t*=1.94, *df*=34.68, p=0.0601, Welch’s). Together these indicate that the five evolved *pgm1* mutations have varying effects on enzymatic activity. The relationship between *pgm1* fitness and Pgm1 V_max_ is best fit by a quadratic, rather than a linear, function (Figure 5D). Therefore, optimal fitness is attained by tuning down, but not eliminating, Pgm1 activity.

### Mutant Pgm1 does not have increased phosphomannomutase or bisphosphatase activity

Phosphoglucomutases can exhibit low levels of phosphomannomutase activity, and we reasoned that active-site mutations in Pgm1 could alter epimer specificity from glucose to mannose, directly compensating for the loss of Sec53 activity (Lowry and Passonneau, 1969). We tested this in two ways. First, we directly measured phosphomannomutase activity of wild-type Pgm1 and two mutant Pgm1, using a ^31^P-NMR spectroscopy-based assay (Figure 5 — figure supplements 1, 4A). We find that wild-type Pgm1 and Pgm1-G514C show comparably low phosphomannomutase activity, accounting for 3.5% and 3.0% of their phosphoglucomutase activity, respectively. We could not detect any phosphomannomutase activity for Pgm1-D295N. We also tested for phosphomannomutase activity indirectly by determining whether mannose-1-phosphate (M1P) acted as a competitive inhibitor in the phosphoglucomutase assay. If mutant Pgm1 had altered epimer specificity, we would expect M1P to compete with glucose-1-phosphate for residency within the active site, leading to an apparent decrease in phosphoglucomutase activity. However, we find no effect of the addition of M1P on phosphoglucomutase activity for any of the mutant Pgm1 enzymes (Figure 5 — figure supplement 4B). Together, these results indicate that mutant Pgm1 does not have enhanced phosphomannomutase activity.

Glucose-1,6-bisphosphate (G16P) is required as an activator of Pgm1 and each active form exhibits barely-detectable activity in the absence of G16P (Figure 5 — figure supplement 1). We determined an apparent half maximal effective concentration (EC_50_) of G16P for several Pgm1 enzymes (Figure 5 — figure supplement 2). Given that Pgm1 is the minor isoform of phosphoglucomutase in yeast, we also reasoned that it could act as a glucose-1,6-bisphosphatase, similarly to the human PMM1 (Veiga-da-Cunha et al., 2008). We performed a specific glucose-1,6-bisphosphatase assay on wild-type and mutant Pgm1 enzyme (Figure 5 — figure supplement 1). However, none of the mutant Pgm1 enzymes have detectable glucose-1,6-bisphosphatase activity.

Finally, we also reasoned that *pgm1* mutations could differently affect the forward and reverse directions of the phosphoglucomutase reaction, with possible effects on the flux of metabolites within the cell. We analyzed the forward and reverse reactions of Pgm1, Pgm1-T231A and Pgm1-G514C using glucose-1-phosphate or glucose-6-phopshate (G1P or G6P) as the substrate (Figure 5 — figure supplement 1). We again used ^31^P-NMR spectroscopy, as G1P and G6P have clearly distinguishable signals (Figure 5 — figure supplement 5). The ratio of forward to reverse reactions are 3.0 ± 1.2 and 4.3 ± 1.1 (± standard deviation) for Pgm1-T231A and Pgm1-G514C, respectively. Although a slightly higher ratio was obtained for wild-type Pgm1 (7.4 ± 3.1), the differences between wild-type and Pgm1-T231A or Pgm1-G514C are not statistically significant (*W*=1 and 2, p=0.057 and 0.53, respectively, Mann–Whitney U test).

## DISCUSSION

The budding yeast *Saccharomyces cerevisiae* is a powerful system for studying conserved aspects of eukaryotic cell biology due to its fast doubling time, small genome size, genetic tractability, and the wealth of genomics tools available (Botstein and Fink, 2011). Many human genes are functionally identical to—and can complement—their yeast homologs (Kachroo et al., 2015). “Humanizing” yeast by replacing a yeast ortholog with the human gene or making an analogous human mutation in a yeast ortholog enables functional studies of human disease (Franssens et al., 2013; Laurent et al., 2016). For example, Mayfield et al. characterized the severity of 84 alleles of human *CBS* (Cystathionine-β-synthase) that are associated with homocystinuria, identifying those alleles where co-factor supplementation can restore high levels of enzymatic function (Mayfield et al., 2012). Gammie et al. characterized 54 *MSH2* variants associated with hereditary nonpolyposis colorectal cancer (HNPCC) showing that about half of the variants have enzymatic function but are targeted for degradation (Gammie et al., 2007). Subsequent work showed that treatment with proteasome inhibitors restores mismatch repair and chemosensitivity in yeast harboring these disease-associated variants (Arlow et al., 2013). In additional to functional characterization, yeast models of human disease enable screens for genetic interactions, thereby uncovering new avenues for therapeutic intervention. Wiskott-Aldrich syndrome, for example, is due to mutations in *WAS*, the human homolog of *LAS17*. Using a temperature-sensitive lethal allele of *las17*, Filteau et al. isolated compensatory mutations in two yeast strains. The authors identified both common and background-specific mechanisms of compensation, including loss-of-function of *CNB1* (Calcineurin B), which is phenocopied by inhibition with cyclosporin A (Filteau et al., 2015).

Here we use experimental evolution to identify mechanisms of genetic compensation in humanized yeast models of PMM2-CDG. We evolved 96 replicate populations for 1,000 generation in rich glucose medium using strains carrying wild-type *SEC53* (the yeast homolog of *PMM2*) and two human disease-associated alleles (*sec53*-V238M and *sec53*-F126L). We show that compensatory mutations arose throughout the evolution experiment. Fitness gains were greater for populations with the *pACT1-sec53-V238M* allele compared to the *pACT1-sec53-F126L* allele, which showed modest gains in fitness (Figure 1C). This indicates that the number of compensatory mutations and/or the ameliorative effects of compensatory mutations are greater for the *pACT1-sec53-V238M* allele compared to the p*ACT1*-*sec53*-F126L allele. Our sequencing and reconstruction experiments show that both explanations are true. While the *pACT1-sec53-V238M* populations showed several unique targets of selection, the *pACT1-sec53-F126L* populations mostly acquired mutations in targets of selection observed in other evolution experiments (Figure 2A). In addition, the genetic reconstruction experiments show that some of the *pgm1* mutations are recessive suppressors of *sec53*-F126L, whereas all five *pgm1*mutations are dominant suppressors of *sec53*-V238M (Figure 4). The latter is unsurprising given that each *pgm1* mutation arose as a heterozygous mutation in a *sec53*-V238M autodiploid background.

Twenty-four genes have been associated with Type 1 CDG in humans (Aebi et al., 1999). All but two Type 1 CDG genes have at least one functional homolog in yeast, and most map one-to-one (Figure 2 — figure supplement 3). We find an overrepresentation of mutations in Type 1 CDG homologs in the evolved *pACT1-sec53-V238M* populations (Figure 2B). This enrichment is striking considering that we observe a significant depletion of mutations in these same genes aggregated across evolution experiments with wild-type *SEC53*, which suggests that they are normally under purifying selection. Specifically, we find five mutations in *PGM1,*two in *ALG9*, and one in *ALG12*. In addition, one *ALG7* mutation was identified in a single p*ACT1*-*sec53*-F126L population. Of these four, only *PGM1* is a known *SEC53* genetic interactor. *ALG7* encodes an acetylglucosaminephosphotransferase responsible for the first step of lipid-linked oligosaccharide synthesis, downstream of *SEC53* in the *N*-linked glycosylation pathway (Barnes et al., 1984; Kukuruzinska and Robbins, 1987). *ALG9* and *ALG12* act downstream of *SEC53* and encode mannosyltransferases, responsible for transferring mannose residues from dolichol-phosphate-mannose to lipid-linked oligosaccharides (Burda et al., 1999, 1996; Cipollo and Trimble, 2002; Frank and Aebi, 2005). This oligosaccharide is eventually transferred to a polypeptide. Each of the evolved mutations in these genes are predicted to have deleterious effects on protein function based on PROVEAN scores (Supplemental Dataset 1) (Choi and Chan, 2015). This suggests that prodding steps in *N*-linked glycosylation alleviates the deleterious effect of *sec53* disease-associated alleles, perhaps by altering pathway flux to match lowered mannose-1-phosphate availability. Evidence from human studies supports the idea of downstream genetic modifiers affecting PMM2-CDG phenotypes. For example, there is not a direct correlation between clinical phenotypes and biochemical properties of PMM2 (Citro et al., 2018; Freeze and Westphal, 2001). One such modifier is a variant of the glucosyltransferase *ALG6*, which has been implicated in worsening clinical outcomes of PMM2-CDG individuals (Bortot et al., 2013; Westphal et al., 2002).

In addition to Type 1 CDG homologs, we identify several other classes of putative genetic suppressors. Both the *sec53*-V238M and *sec53*-F126L mutations retain partial functionality and Lao et al., 2019 showed that increasing expression of these alleles improves growth. We, therefore, expected to find copy number variants (CNVs) and aneuploidies as compensatory mutations. Though we do identify several strains that show evidence of CNVs at the *HO* locus on Chromosome IV, where the *sec53* alleles reside in our strains, karyotype changes are not a major route of adaptation (Figure 2 — source data 1). Reduced function of mutant Sec53 could be due to destabilization and degradation of the protein. We identify recurrent mutations in the 26S proteasome lid subunit *RPN5* in the *pACT1-sec53-V238M* background, in the E3 ubiquitin ligase *UBR1* in the p*ACT1*-*sec53*-F126L background, and in the ubiquitin-specific protease *UBP3* in both *pACT1-sec53* backgrounds (Figure 2). In addition, we identify a mutation in the ubiquitin-specific protease *UBP15* in a single *pACT1-sec53-V238M* clone. And finally, we observe multiple independent mutations in the known *SEC53*-interactors *RPN5*, *SRP1*, and *YNL011C* in *sec53* populations, which could have positive genetic interactions with *sec53* (Figure 2).

We find a high probability that the mutational spectrum of *PGM1* (5 missense mutations, 0 frameshift/nonsense) resembles selection for non-loss-of-function (posterior probability = 0.96). However, *PGM1* loss-of-function (*pgm1*Δ) provides a fitness benefit indistinguishable from two evolved mutations (*pgm1*-R64C and *pgm1*-D295N) and higher than one evolved mutation (*pgm1*-T231A). Likewise, the *pgm1*Δ allele restores invertase and carboxypeptidase Y glycosylation and growth in the tetrad analyses, albeit recessively. It is not immediately clear, therefore, why we do not observe frameshift/nonsense mutations in *PGM1* in the evolution experiment. We have previously shown that genetic interactions between evolved mutations are not always recapitulated by gene deletion (Vignogna et al., 2021). Given that most of the evolved clones containing *pgm1* mutations are more fit than the reconstructed strains, it is possible that other evolved mutations interact epistatically only with non-loss-of-function *pgm1* mutations. However, there are no shared mutational targets (genes or pathways) among the evolved clones with *pgm1* mutations.

Using tetrad dissections, we show that each evolved *pgm1* mutation and the catalytically-dead *pgm1*-S120A mutation are dominant suppressors of *sec53*-V238M. *pgm1*Δ is not a dominant suppressor based on the tetrad analyses, contrary to the fitness assays where the heterozygous *pgm1*Δ allele confers a fitness benefit comparable to several evolved mutations. This discrepancy likely results from differences in ploidy or growth medium of the two experiments — diploids in liquid culture versus haploids on agar plates. We also find that several *pgm1* mutations (T521K, D295N, *pgm1*Δ, S120A) are recessive suppressors of the *sec53*-F126L allele in the tetrad dissections, whereas we were unable to detect any improvement using fitness assays. Three of these four mutations have no detectable enzymatic activity. However, the R64C allele, which also does not have detectable enzymatic activity does not suppress.

To test specific hypotheses for the mechanism of suppression of *sec53* disease alleles by mutation of *pgm1*, we characterized the biochemical properties of purified recombinant Pgm1 mutant proteins. We find that phosphoglucomutase activity ranges from no-detectable activity (R64C and D295N) to near wild-type (T231). Based on well-characterized human and rabbit PGM1 structures, R64C and D295N mutations likely affect active-site integrity (Liu et al., 1997; Stiers et al., 2016). R64 helps hold the catalytic serine (S120) in place and D295 coordinates a catalytically-essential Mg^2+^ ion. The T231A mutation, which occurs in a residue near the metal binding loop, shows the highest activity levels among our mutant Pgm1 enzymes. The other two mutations with low but detectable activity, G514C and T521K, both occur in the phosphate-binding domain of Pgm1, a region important for substrate binding (Beamer, 2015; Stiers et al., 2017a, 2017b; Stiers and Beamer, 2018). Comparing *pgm1* fitness effects versus their corresponding enzyme V_max_ we find an inverted U-shaped curve, where reducing phosphoglucomutase activity improves fitness, but complete loss of activity overshoots the optimum (Figure 5D). Plotting fitness versus k_cat_/K_m_ results in a qualitatively similar trend, although comparing k_cat_/K_m_ between mutant enzymes may not be appropriate (Eisenthal et al., 2007).

We tested several hypotheses involving Pgm1 alteration-of-function. First, the evolved Pgm1 proteins could directly complement the loss of Sec53 phosphomannomutase activity by switching epimer specificity from glucose to mannose. However, we do not observe increased phosphomannomutase activity in the Pgm1 mutant proteins, nor is this hypothesis consistent with our genetic data: direct complementation would be expected to suppress both the *sec53*-V238M and *sec53*-F126L alleles. A second possible mechanism of suppression is increasing the production of glucose-1,6-bisphosphate (G16P). G16P is an endogenous activator and stabilizer of Pmm2/Sec53. Compounds that raise G16P levels by increasing its production and/or slowing its degradation show promise as candidates for treatment of PMM2-CDG (Iyer et al., 2019; Monticelli et al., 2019). Whereas higher eukaryotes and some bacteria have a dedicated glucose-1,6-bisphosphate synthase, in yeast G16P is produced as a dissociated intermediate of the phosphoglucomutase reaction (Maliekal et al., 2007; Neumann et al., 2021). We hypothesized that evolved mutations in *PGM1* may increase the rate of G16P dissociation. Recent work has shown that a mutation affecting Mg^2+^ coordination in *L. lactis* β-phosphoglucomutase can increase dissociation of β-G16P *in vitro* (Wood et al., 2021). While we did not directly measure G16P synthase activity, we do not find evidence of increased glucose-1,6-bisphosphatase activity in our mutant Pgm1 enzymes. A third possible mechanism of compensation is the indirect increase of G16P by increasing the relative flux of the reverse (G6P to G1P) reaction relative to the forward (G1P to G6P) reaction. This would create a futile cycle of G1P/G6P interconversion, which is expected to result in higher intracellular G16P. This hypothesis is attractive in that it accounts for three observations regarding the evolved *pgm1* mutations: (1) the observation of only missense (not nonsense or frameshift) mutations in the evolution experiment, (2) the dominance of suppressors in our genetic assay, and (3) greater extent of suppression for the stability mutant (V238M) of *SEC53*.

There is truth to the saying that “a selection is worth a thousand screens.” Experimental evolution takes what would otherwise be a screen—looking for larger colonies on a field of smaller ones—and turns it into a selection. Here we demonstrate the power of yeast experimental evolution to identify genetic mechanisms that compensate for the molecular defects of PMM2-CDG disease-associated alleles. This general approach is applicable to other human disease alleles that affect core and conserved biological processes including protein glycosylation, ribosome maturation, mitochondrial function, and RNA modification. Mutations that impinge upon these processes are often highly pleiotropic. In humans, salient phenotypes may be neurological or developmental, but in yeast, these disease alleles invariably lead to slow growth. With a population size of 10^5^, evolution can act on differences as small as 0.001%, far below the ability to resolve on plates. Even our best fluorescence-based assays can only resolve differences of ~0.1% (Lang and Botstein, 2011). With a genome size of 10^7^ bp, a mutation rate of 10^-10^ per bp per generation, and with continuous selective pressure exerted over thousands of generations, each population will sample hundreds of thousands of coding-sequence mutations. Furthermore, evolution can identify complex compensatory interactions involving multiple mutations or mutations in essential genes, both of which are largely absent from current genetic interaction networks.

## METHODS

### Experimental Evolution

The haploid strains used in the evolution experiment are S288c derivatives and were described previously (Lao et al., 2019). Each strain maintains a *URA3*-marked plasmid encoding wild-type *SEC53* (pTC416-SEC53 described in Lao et al. 2019). Strains with the p*SEC53*-sec*53-* F126L allele grew too slowly to keep up with a 1:2^10^ dilution every 24 hours.

To initiate the evolution experiment, strains were struck to synthetic defined (SD) media (CSM minus arginine, yeast nitrogen base, glucose) containing 5-Fluoroorotic Acid (5FOA) (Zymo Research, Irvine, CA, USA). to isolate cells that spontaneously lost pTC416-SEC53. A single colony was chosen for each strain and used to inoculate 10 mL of YPD (yeast extract, peptone, dextrose) plus ampicillin and tetracycline. These cultures grew to saturation then were distributed into 96-well plates. Every 24 hours populations were diluted 1:2^10^ using a BiomekFX liquid handler into 125 μL of YPD plus 100 μg/mL ampicillin and 25 μg/mL tetracycline to prevent bacterial contamination. Plates were then left at 30°C without shaking. The dilution scheme equates to 10 generations of growth per day at an effective population size of ~10^5^. Every 100 generations, 50 μL of 60% glycerol was added to each population and archived at −80°C.

OD_600_ was measured every 50 generations over the course of the evolution experiment. After daily dilutions were completed, populations were resuspended by orbital vortexing at 1050 rpm for 15 seconds then transferred to a Tecan Infinite M200 Pro plate reader for sampling.

### Whole-Genome Sequencing

Benomyl assays were performed by spotting 5 μL of populations onto YPD plus 20 μg/mL benomyl (dissolved in DMSO) and incubating at 25°C. Ploidy was then inferred by visually comparing growth of evolved populations to control haploid and diploid strains.

Single clones were isolated for sequencing by streaking 5 μL of cryo-archived populations to YPD agar. One clone from each population was randomly chosen and grown to saturation in 5 mL YPD, pelleted, and frozen at −20°C. Genomic DNA was harvested from frozen cell pellets using phenol-chloroform extraction and precipitated in ethanol. Total genomic DNA was used in a Nextera library preparation as described previously (Buskirk et al., 2017). Pooled clones were sequenced using an Illumina NovaSeq 6000 sequencer by the Sequencing Core Facility at the Lewis-Sigler Institute for Integrative Genomics at Princeton University.

### Sequencing Analysis

Raw sequencing data were concatenated and then demultiplexed using a dual-index barcode splitter (bitbucket.org/princeton_genomics/barcode_splitter/src). Adapter sequences were trimmed using Trimmomatic (Bolger et al., 2014). Modified S288c reference genomes were constructed using *ref*orm (github.com/gencorefacility/reform) to correct for strain-construction-related differences between our strains and canonical S288c. Trimmed reads were aligned to these modified S288c reference genomes using BWA and mutations were called using FreeBayes (Garrison and Marth, 2012; Li and Durbin, 2009). VCF files were annotated with SnpEff (Cingolani et al., 2012). All calls were confirmed manually by viewing BAM files in IGV (Thorvaldsdóttir et al., 2013). Clones were predicted to be diploids if two or more mutations were called at an allele frequency of 0.5. Copy number variants (CNVs) and aneuploidies were called via changes in sequencing coverage using Control-FREEC (Boeva et al., 2012). CNVs and aneuploidies were confirmed by visually inspecting coverage plots.

### Yeast Strain Construction

Evolved mutations were reconstructed via CRISPR/Cas9 allele swaps. For mutations that fall within a Cas9 gRNA site (*alg9-S230R*, *pgm1-T231A*, *pgm1*-G514C), repair templates were produced by PCR-amplifying 500-2000bp fragments centered around the mutation of interest from evolved clones. Commercially-synthesized dsDNA fragments (Integrated DNA Technologies, Coralville, IA, USA) were used as repair templates for all other mutations (*pgm1*-R64C, *pgm1*-S120A, *pgm1*-D295N, *pgm1*-T521K, *pgm1*Δ). These 500bp fragments contained our mutation of interest as well as 1-2 synonymous mutations in the Cas9 gRNA site to halt cutting. The *pgm1*Δ repair template was a dsDNA fragment consisting of 250bp upstream then 250bp downstream of the ORF. Repair templates were co-transformed into haploid ancestral strains with a plasmid encoding Cas9 and gRNAs targeting near the mutation site (Addgene #83476) (Laughery et al., 2015).

Heterozygous mutant strains were constructed by mating haploid mutants to *MATα* versions of their respective ancestral strains. Homozygous mutant strains were constructed by reconstructing mutations in *MAT*α versions of the ancestral strains and mating them to the respective *MAT***a** mutant strain. Successful genetic reconstructions were confirmed via Sanger sequencing (Psomagen, Rockville, MD, USA).

Linked *PGM1* strains used for tetrad dissections were constructed by transforming strains with a linearized integrating plasmid (Addgene #35121) containing wild-type *PGM1*(Goldstein and McCusker, 1999). Successful integration of the plasmid next to the endogenous *PGM1* locus was confirmed via PCR.

### Competitive Fitness Assays

We measured the fitness effect of evolved mutations using competitive fitness assays described previously with some modifications (Buskirk et al., 2017). Query strains were mixed 1:1 with a fluorescently labeled version of the p*ACT1*-*SEC53*-WT ancestral diploid strain. Each p*ACT1-sec53-* F126L strain was outcompeted immediately by the reference at this ratio, so we also tried mixing at a ratio of 50:1. Cocultures were propagated in 96-well plates in the same conditions in which they evolved for up to 50 generations. Saturated cultures were sampled for flow-cytometry every 10 generations. Each genotype assayed was done so with at least 24 technical replicates (competitions) of 4 biological replicates (2 clones isolated each from 2 isogenic but separately-constructed strains). Analyses were performed in Flowjo and R.

### LOF/non-LOF Bayesian Analysis

From reconstruction data from previously published evolution experiments (Fisher et al., 2018; Lang et al., 2013; Marad et al., 2018) we identified ten genes where selection is acting on loss-of-function (LOF: *ACE2*, *CTS1*, *ROT2*, *YUR1*, *STE11*, *STE12*, *STE4*_haploids_, *STE5*, *IRA1*, and *IRA2*) and six genes where selection is for non-loss-of-function (non-LOF: *CNE1*, *GAS1*, *KEG1*, *KRE5*, *KRE6*, and *STE4*_autodiploids_). These data established prior probabilities that selection is acting on LOF (0.625) or non-LOF (0.375). We then determined the conditional probabilities of missense and frameshift/nonsense mutations given selection for LOF and non-LOF using 240 mutations across these 16 genes as well as 414 5FOA-resistant mutations at *URA3* and 454 canavanine-resistant mutations at *CAN1* (Lang and Murray, 2008). From these data we can estimate the log likelihood and posterior probabilities that selection is acting on LOF or non-LOF given the observed mutational spectrum for any given gene (Supplemental Dataset 2).

### CPY and Invertase Blots

Strains were grown in 5mL YPD plus 100 μg/mL ampicillin and 25 μg/mL tetracycline until saturated, then pelleted, and frozen at −20°C. Cell pellets were lysed in Ripa lysis buffer (150mM NaCl, 1% Nonidet P-40, 0.5% sodium deoxycholate, 0.1% sodium dodecyl sulfate, 25mM Tris (pH 7.4)) with protease inhibitors (Thermo Fisher Scientific, Waltham, MA, USA) and 10mM DTT, incubated for 30 min on ice, sonicated 5 times briefly, incubated for a further 10 min on ice, and centrifuged at 20,000xg for 10 min at 4°C. Supernatant was quantified by Micro BCA Protein Assay (Thermo Fisher). 30 μg of each lysate were prepared in Lamelli buffer and were incubated at 95°C for 5 min and chilled at 4°C. Lysates were separated on a 6% SDS PAGE gel. Protein was transferred to 0.2 μm pore nitrocellulose at 100 volts for 100 min at 4°C in Towbin buffer. The membrane was rinsed and stained with Ponceau S for normalization. The membranes were blocked with 5% milk/TBST for 1 hour at room temperature.

Carboxypeptidase Y blots were incubated with an anti-carboxypeptidase antibody at 1:1000 overnight at 4°C and washed 3 times with TBST. The blots were then incubated with anti-rabbit IgG HRP diluted 1:2000 in milk/TBST for 1 hour, washed 3 times and developed with ECL reagent (Bio-Rad Laboratories, Hercules, CA, USA). Invertase blots were prepared similarly with an anti-invertase antibody and anti-goat IgG HRP. All images were captured on the Bio-Rad ChemiDoc MP Imaging System (Bio-Rad). Analysis was done with Image Lab Software (Bio-Rad). Raw images of blots and Ponceau stain controls are available in Figure 3 — source data 1.

### Tetrad Dissection and Inference of Genetic Interactions

Heterozygous yeast strains (e.g. *SEC53/sec53 PGM1/pgm1*) were sporulated by resuspending 1 ml of overnight YPD culture in 2 ml of SPO++ (1.5% potassium acetate, 0.25% yeast extract, 0.25% dextrose, 0.002% histidine, 0.002% tryptophan) and incubated on a roller-drum for ~7 days at room temperature. 10 tetrads per strain were dissected on YPD agar and then incubated for 48 hours at 30°C. We performed a log-likelihood test to determine if patterns of segregation are indicative of suppression (Figure 4 — source data 1). Note that in order to calculate conditional probabilities we had to include an error term (0.01) to account for biological noise in the real data. Our inference of suppression, however, is robust to our choice of error value.

### Bacterial Growth and Lysis

Wild-type and mutant *PGM1* alleles were cloned into the expression vector pGEX-2T. Cloned vectors were transformed into *E. coli* BL21(DE3). Bacteria were grown in lysogeny broth (LB) medium plus 0.1 mg/ml ampicillin. Each *PGM1* allele was expressed by induction at 0.1 OD with 0.1 mg/mL IPTG for 16 hours at 15°C. Bacteria were harvested by centrifugation, washed with PBS, and stored at −80°C. Bacterial lysis was accomplished in 25 mM Tris buffer (pH 8) containing 1 mM EDTA, 2 mM DTT, 5% glycerol and 0.1 mM PMSF, by adding 1 mg/mL lysozyme and 2.5 μg/mL DNase. Clear extract was then obtained by centrifugation.

### Protein Purification

Protein purification was accomplished using Glutathione Sepharose High Performance resin (GSTrap by Cytiva, Marlborough, MA, USA) and Benzamidine Sepharose 4 Fast Flow resin (HiTrap Benzamidine FF by Cytiva). Cleavage of the GST-tag was conducted on-column by adding thrombin. All the procedures were performed according to manufacturer’s instructions. Briefly, clear extract obtained from 50 mL of bacterial culture was loaded onto the GSTrap column (5mL), previously equilibrated with 25 mM Tris (pH 8) containing 2 mM DTT and 5% glycerol. Unbound protein was eluted, then the column was washed with 20 mM Tris (pH 8) containing 1 mM DTT, 150 mM NaCl, 1 mM MgCl2, and 5% glycerol (buffer A). Thrombin (30 units in 5 mL of buffer A) was loaded and the column was incubated for 14-16 hours at 10°C. GSTrap and HiTrap Benzamidine columns were connected in series, and elution of the untagged protein was accomplished with buffer A. Fractions were analyzed by SDS-PAGE and activity assays. Active fractions were finally collected and concentrated by ultrafiltration. A yield spanning 1.0-4.6 mg of protein per liter of bacterial culture was obtained.

Each protein was obtained in a pure form, as judged by SDS-PAGE and gel filtration analyses, with a single band observed at the expected molecular weight (63 kDa). Gel filtration analysis was performed on BioSep-SEC-S 3000 column (Phenomenex, Torrance, CA, USA) at 1 mL/min. The eluent was 20 mM Tris (pH 8), 1 mM MgCl_2_, 150 mM NaCl.

### Activity Assays

Phosphoglucomutase activity was assayed spectrophotometrically at 340 nm and 29°C by following the reduction of NADP^+^ to NADPH in a 0.3 mL reaction mixture containing 25 mM Tris (pH 8), 5 mM MgCl_2_, 1 mM DTT, 0.25 mM NADP^+^, and 2.7 U/mL glucose 6-phosphate dehydrogenase in the presence of 0.3 mM glucose-1-phosphate (G1P) and 0.02 mM glucose-1,6-bisphosphate (G16P). Additional experiments were performed with the addition of mannose-1-phosphate (M1P; 10, 50, 200 μM). K_m_ were evaluated by measuring activity in the presence of G1P ranging from 0 mM to 0.6 mM. Analyses were performed in R using the *drc* package (Ritz et al., 2015). Bisphosphatase activity was measured spectrophotometrically at 340 nm and 29°C in the presence of 0.1 mM G16P, 0.25 mM NADP+, and 2.7 U/mL glucose 6 phosphate dehydrogenase. Activity expressed as the number of micromoles of substrate transformed per minute per mg of protein under the standard conditions.

Thermostabilities of Pgm1 were determined by incubating purified protein for 5 or 10 minutes at different temperatures (30, 35, 40, 45 °C) in 20 mM Tris (pH 8), 1 mM MgCl2, 150 mM NaCl, 5% glycerol, and 0.1 mg/mL BSA. After cooling on ice, residual activity was measured under standard conditions.

### ^31^P-NMR Spectroscopy

Phosphoglucomutase and phosphomannomutase activity were measured by ^31^P-NMR spectroscopy (Citro et al., 2017). This assay allows us to exclude the effects on the change in absorbance due to impurities that could be substrates for the ancillary enzymes required for the assay. This is of utmost importance when the activity is particularly low and measuring the rate of change in absorbance requires long incubation times, as it is the case of phosphomannomutase activity for Pgm1. A discontinuous assay was performed. Appropriate amounts of proteins were incubated at 30°C with 1 mM G1P or M1P in 20 mM Tris (pH 8), 1 mM MgCl2, 0.1 mg/ml BSA, in the presence of 20 μM G16P for up to 60 min. The reaction was stopped with 50 mM EDTA on ice and heat inactivation. The content of the residual substrate (M1P or G1P) and/or the product (M6P or G6P) were measured recording ^31^P-NMR spectra. Creatine phosphate was added as an internal standard for the quantitative analysis. The ^1^H-decoupled, one-dimensional ^31^P spectra were recorded at 161.976 MHz on a Bruker Avance III HD spectrometer 400MHz, equipped with a BBO BB-H&F-D CryoProbe Prodigy fitted with a gradient along the Z-axis, at a probe temperature of 27°C. Spectral width 120 ppm, delay time 1.2s, pulse width of 12.0μs were applied. All the samples contained 10% ^2^H_2_O for internal lock.

Similarly, forward (G1P to G6P) and reverse (G6P to G1P) phosphoglucomutase activities were measured following the same procedure described above, with 1mM G1P or G6P as the substrate.

## DATA AVAILABILITY

The short-read sequencing data reported in this study have been deposited to the NCBI BioProject database, accession number PRJNA784975.

The following previously published datasets were used:

Lang et al., 2013: NCBI BioProject PRJNA205542 Fisher et al., 2018: NCBI BioProject PRJNA422100 Johnson et al., 2021: NCBI BioProject PRJNA668346

## ACKNOWLEDGEMENTS AND FUNDING

We thank members of the Andreotti Lab and members of the Lang Lab for comments on the manuscript. We thank M. Kathryn Iovine for sharing the pGEX-2T plasmid. We thank Lesa Beamer for useful discussions. This study was supported by the National Institutes of Health: R01GM127420 to GIL and P20GM139769 to RS (Trudy Mackay PI/PD). This study was also supported by DiSTABiF, University of Campania “Luigi Vanvitelli” (fellowship POR Campania FSE 2014/2020 “Dottorati di Ricerca Con Caratterizzazione Industriale” to MA). Portions of this research were conducted on Lehigh University’s Research Computing infrastructure partially supported by the National Science Foundation (Award 2019035). Perlara PBC provided funding support for reagents and personnel time.

## AUTHOR CONTRIBUTIONS

Conceptualization: RCV, GA, EOP, GIL

Formal Analysis: RCV, MA, GA, GIL

Investigation: RCV, MA, MM, JWN

Writing – Original Draft Preparation: RCV, GIL

Writing – Review & Editing: RCV, MA, MM, RS, GA, EOP, GIL

Visualization: RCV, GIL

Supervision: RS, GA, GIL

Funding Acquisition: EOP, GIL

