## Supplemental Data 1 for "Experimental evolution of phosphomannomutase-deficient yeast reveals compensatory mutations in a phosphoglucomutase"

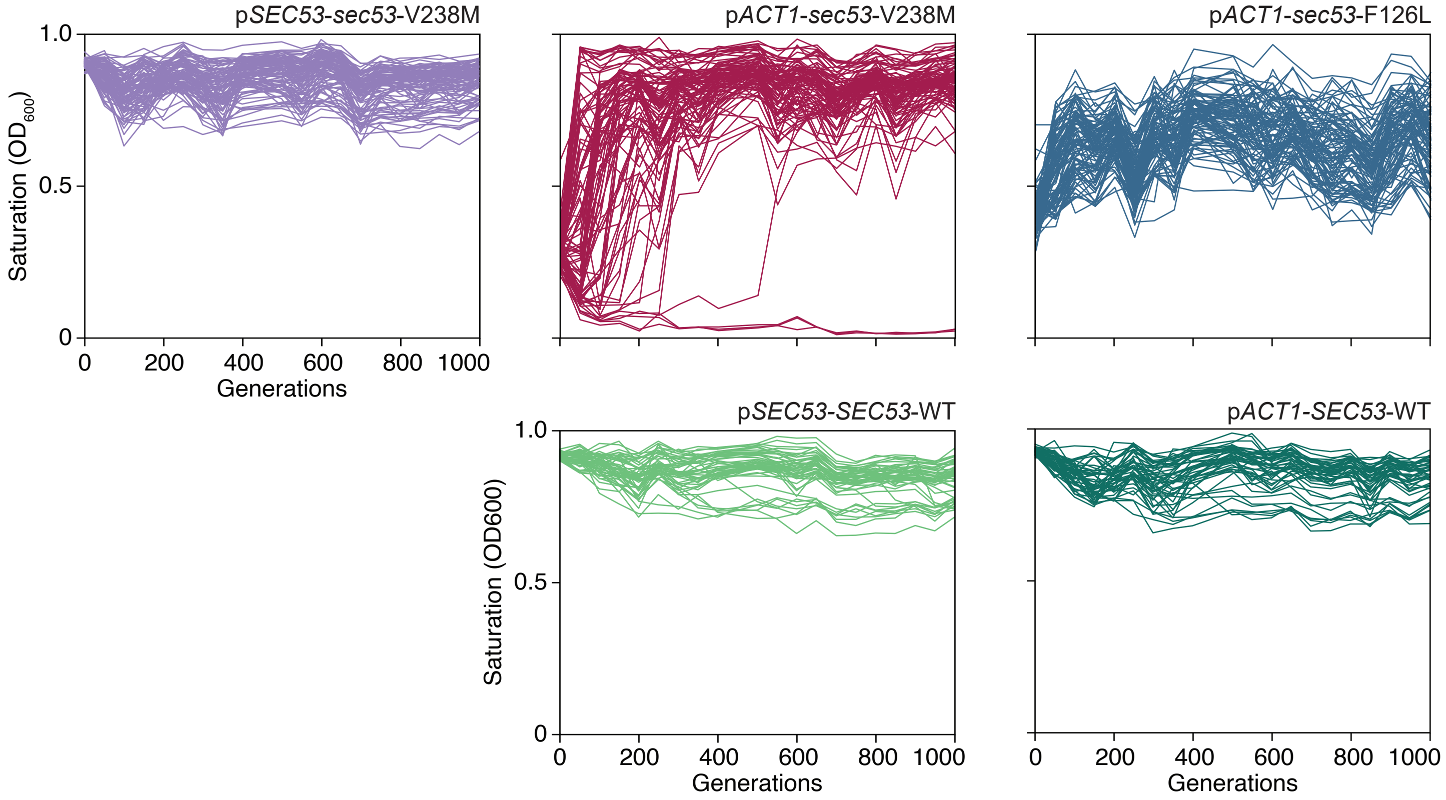


**Figure 1 — figure supplement 1. $OD_{600}$ readings of each population.** Single time-point $OD_{600}$ readings of populations over the course of the evolution experiment. Every 50 generations, following daily transfer, populations were resuspended and OD was measured. Each line represents one population.
