## Supplemental Data 2 for "Experimental evolution of phosphomannomutase-deficient yeast reveals compensatory mutations in a phosphoglucomutase"

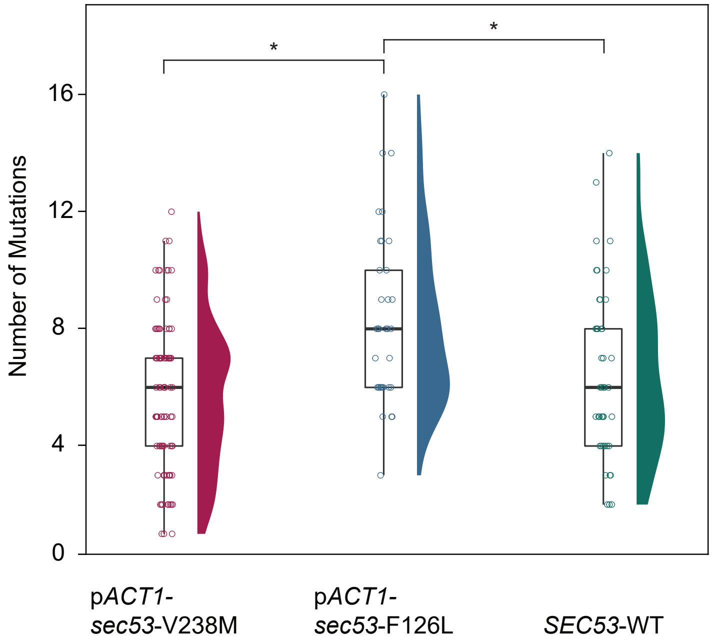


**Figure 2 — figure supplement 1. Number of de novo mutations per *SEC53* genotype.** Each circle represents the number of SNPs and small indels present in a sequenced clone (Supplemental Dataset 1). p*SEC53*-*SEC53*-WT and p*ACT1*-*SEC53*-WT are grouped as "*SEC53*-WT". Distribution of points are shown as violin plots. Only statistically significant differences shown as bars. p*ACT1*-*sec53*-V238M vs. *SEC53*-WT (p=0.326), p*ACT1*-*sec53*-V238M vs. p*ACT1*-*sec53*-F126L (p>0.0001), p*ACT1*-*sec53*-F126L vs. *SEC53*-WT (p=0.0234) (*df*=167, *F*=9.67, one-way ANOVA with Tukey post-hoc test).
