## Supplemental Data 3 for "Experimental evolution of phosphomannomutase-deficient yeast reveals compensatory mutations in a phosphoglucomutase"

| Chr. | Start (kb) | End (kb) | Length (kb) | Description | Copy Number | Type | ***sec53*-V238M** | ***sec53*-F126L** | ***SEC53*-WT** |
|---|---|---|---|---|---|---|---|---|---|
| IV | 1100 | 1220 | 120 | Gain | 3N | CNV | 7/91 | 0/36 | 0/43 |
| V | 450 | 500 | 50 | Loss | 1N | CNV | 30/91 | 7/36 | 8/43 |
| VI | 200 | 270 | 70 | Loss | 0N and 1N | CNV | 4/91 | 0/36 | 0/43 |
| I | 0 | 230 | 230 | Loss | 1N | Aneuploidy | 8/91 | 0/36 | 0/43 |
| II | 0 | 813 | 813 | Gain | 3N | Aneuploidy | 12/91 | 0/36 | 0/43 |
| VI | 0 | 270 | 270 | Loss | 1N | Aneuploidy | 6/91 | 0/36 | 0/43 |
| XIV | 0 | 784 | 784 | Gain | 3N | Aneuploidy | 24/91 | 0/36 | 0/43 |

**Figure 2 — figure supplement 2. Recurrent CNVs and aneuploidies in the evolution experiment.** Listed are the copy number variants (CNVs) and aneuploidies present in three or more sequenced clones among p*ACT1*-*sec53*-V238M, p*ACT1*-*sec53*-F126L, and p*SEC53*-*SEC53*-WT and p*ACT1*-*SEC53*-WT (combined as *SEC53*-WT) populations. We exclude from this table putative CNVs that occur near telomeric and subtelomeric regions. This includes several apparent amplifications of the *HO* locus (where *sec53* resides in our strains) in p*ACT1*-*SEC53*-V238M clones.
