## Supplemental Data 4 for "Experimental evolution of phosphomannomutase-deficient yeast reveals compensatory mutations in a phosphoglucomutase"

| Human gene(s) | Yeast gene(s) | V238M | F126L |
|---|---|---|---|
| *ALG1* | *ALG1* | 0 | 0 |
| *ALG3* | *ALG3* | 0 | 0 |
| *ALG6* | *ALG6* | 0 | 0 |
| *DPAGT1* | *ALG7* | 0 | 1 |
| *ALG8* | *ALG8* | 0 | 0 |
| *ALG9* | *ALG9* | 2 | 0 |
| *ALG11* | *ALG11* | 0 | 0 |
| *ALG12* | *ALG12* | 1 | 0 |
| *ALG13* | *ALG13* | 0 | 0 |
| *DPM1, DPM2, DPM3* | *DPM1, YIL102C-A* | 0, 0 | 0, 0 |
| *SRD5A3* | *DFG10* | 0 | 0 |
| *NUS1* | *NUS1* | 0 | 0 |
| *MAGT1* | *OST3, OST6* | 0, 0 | 0, 0 |
| *PGM1* | *PGM1, PGM2* | 5, 0 | 0, 0 |
| *MPI* | *PMI40* | 0 | 0 |
| *DHDD5* | *RER2* | 0 | 0 |
| *RFT1* | *RFT1* | 0 | 0 |
| *PMM2* | *SEC53* | 0 | 0 |
| *DOLK* | *SEC59* | 0 | 0 |
| *STT3A, STT3B* | *STT3* | 0 | 0 |
| *DDOST* | *WBP1* | 0 | 0 |
| *MPDU1* | n/a | n/a | n/a |
| *SSR4* | n/a | n/a | n/a |

**Figure 2 — figure supplement 3. Number of mutations in Type 1 CDG homologs.** Listed are the number of nonsynonymous mutations in Type 1 CDG homologs in the evolution experiment among p*ACT1*-*sec53*-V238M and p*ACT1*-*sec53*-F126L clones. The list of known Type 1 CDG genes was retrieved from Online Mendelian Inheritance in Man (omim.org) and Alliance Of Genome Resources (alliancegenome.org). No mutations arose in known homologs of Type 2 CDG genes. *MPDU1* and *SSR4* have no known yeast homolog.
