## Supplemental Data 5 for "Experimental evolution of phosphomannomutase-deficient yeast reveals compensatory mutations in a phosphoglucomutase"

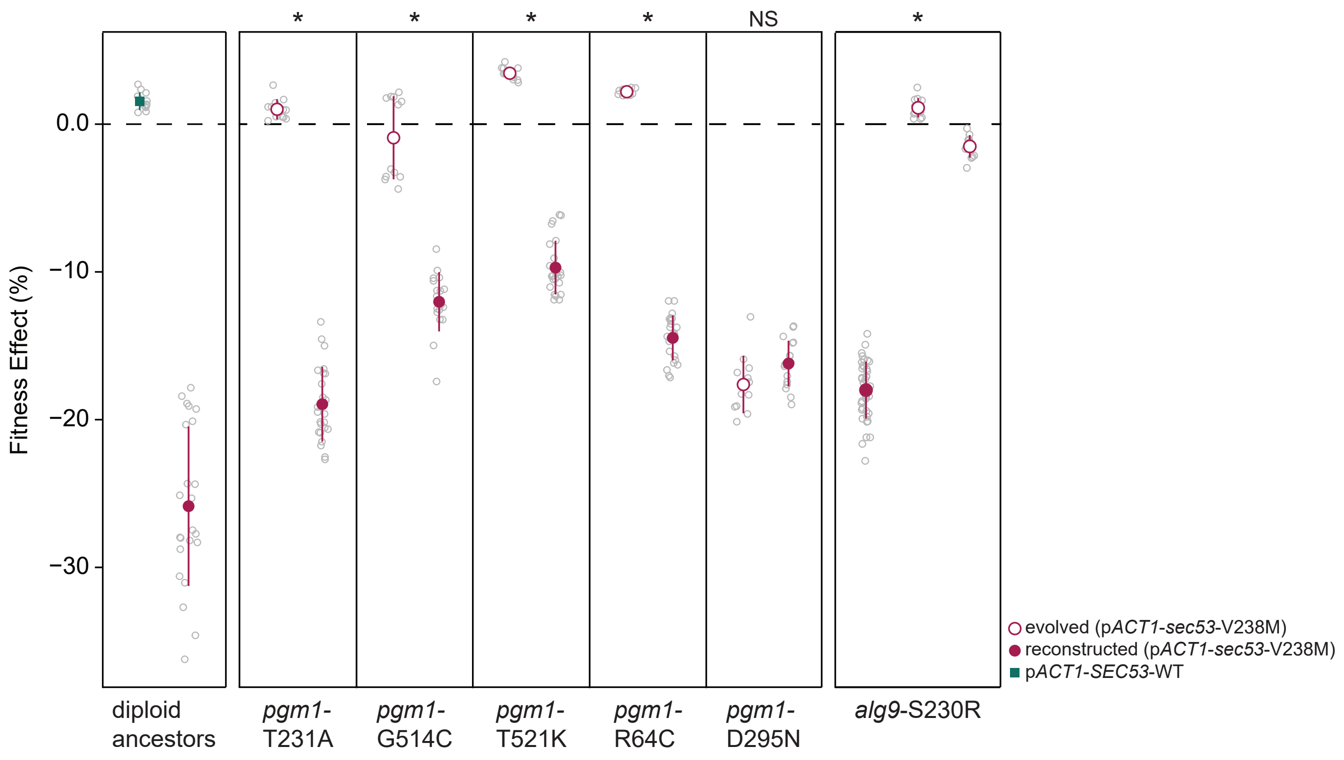


**Figure 3 — figure supplement 1. Fitness effects of reconstructed mutations and evolved clones containing those same mutations.** Average fitness effects and standard deviations of reconstructed heterozygous *pgm1* and *alg9* mutations (closed circles) compared to evolved clones containing those mutations plus 3-9 additional mutations (open circles). Replicate measurements are plotted as gray circles. Note there are two clones from different populations with an *alg9*-S230R mutation. Asterisks (*) represent statically significant differences between fitness effects of evolved clones and fitness effects of reconstructed clones (Welch's modified t-test): *pgm1*-T231A ($p<0.0001$, $t=36.1$, $df=28.9$); *pgm1*-G514C ($p<0.0001$, $t=11.8$, $df=18.4$); *pgm1*-T521K ($p<0.0001$, $t=33.8$, $df=27.3$); *pgm1*-R64C ($p<0.0001$, $t=52.9$, $df=24.3$); *pgm1*-D295N ($p=0.05$, $t=2.10$, $df=19.9$); *alg9*-S230R (left) ($p<0.0001$, $t=53.7$, $df=49.7$); *alg9*-S230R (right) ($p<0.0001$, $t=44.1$, $df=46.1$).
