## Supplemental Data 6 for "Experimental evolution of phosphomannomutase-deficient yeast reveals compensatory mutations in a phosphoglucomutase"

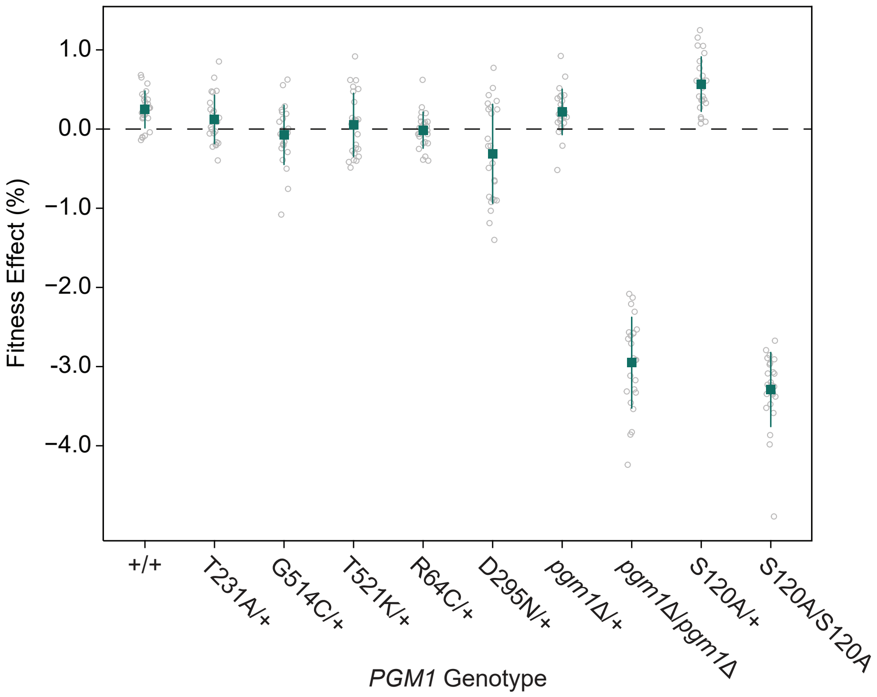


**Figure 3 — figure supplement 2. Fitness effects of *pgm1* mutations in the *SEC53*-WT background.** Average fitness effects and standard deviations of constructed *pgm1* mutations in the diploid p*ACT1*-*SEC53*-WT background. Replicate measurements are plotted as gray circles. Plus signs (+) indicate wild-type alleles.
