## Supplemental Data 7 for "Experimental evolution of phosphomannomutase-deficient yeast reveals compensatory mutations in a phosphoglucomutase"

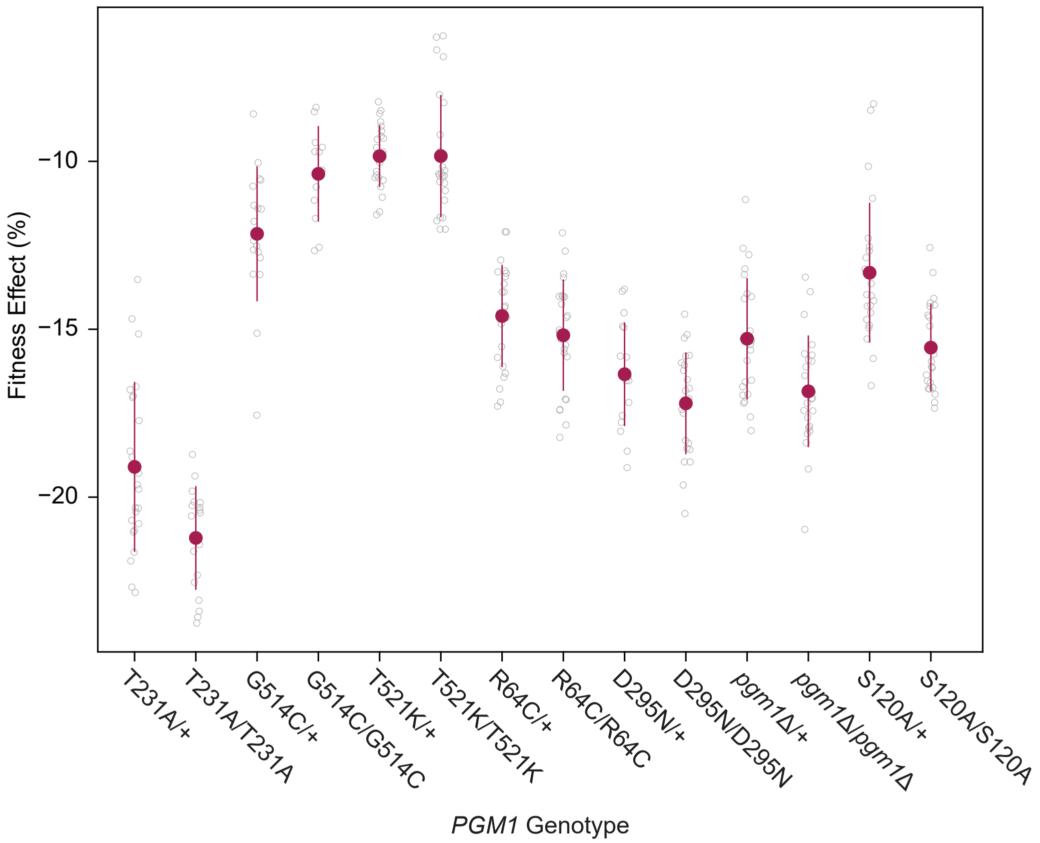


**Figure 3 — figure supplement 3. Fitness effect of homozygous pgm1 mutations.** Average fitness effects and standard deviations of reconstructed *pgm1* mutations in the diploid p*ACT1*-*sec53*-V238M background as heterozygous (mutation/+) or homozygous (mutation/mutation) alleles. Replicate measurements are plotted as gray circles. Comparison of heterozygous to homozygous allele fitness effects (Welch's modified t-test): *pgm1*-T231A (p=0.002, *t*=3.35, *df*=38.6); *pgm1*-G514C (p=0.008, *t*=2.85, *df*=27.8); *pgm1*-T521K (p=0.996, *t*=0.005, *df*=34.0); *pgm1*-R64C (p=0.219, *t*=1.24, *df*=45.7); *pgm1*-T231A (p=0.077, *t*=1.84, *df*=36.4); *pgm1*Δ (p=0.003, *t*=3.12, *df*=45.7); *pgm1*-S120A (p<0.0001, *t*=4.45, *df*=38.6).
