## Supplemental Data 8 for "Experimental evolution of phosphomannomutase-deficient yeast reveals compensatory mutations in a phosphoglucomutase"

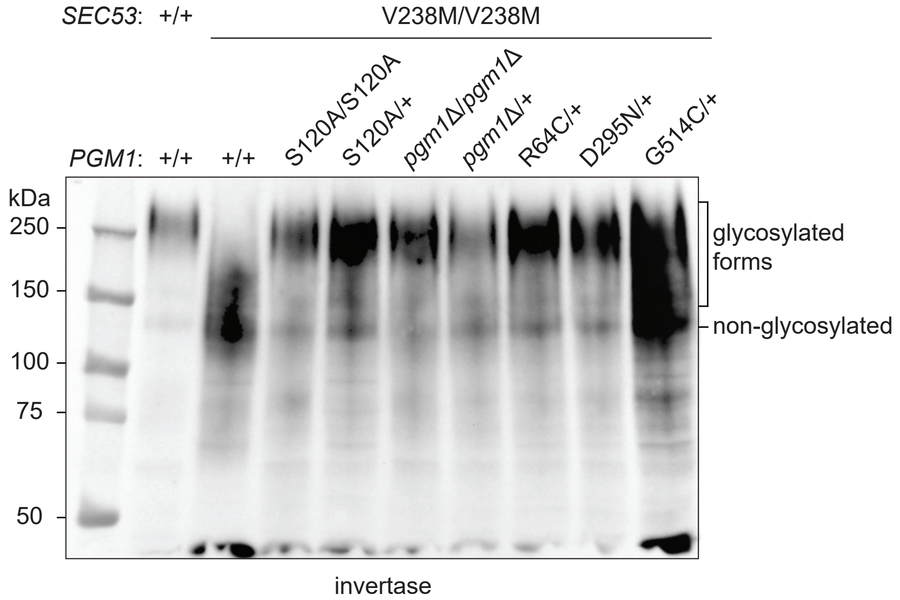


**Figure 3 — figure supplement 4. Effects of *pgm1* mutations on invertase glycosylation in the p*ACT1-sec53*-V238M background.** Western blot of invertase from reconstructed *sec53*/*pgm1* strains.
