## Supplemental Data 9 for "Experimental evolution of phosphomannomutase-deficient yeast reveals compensatory mutations in a phosphoglucomutase"

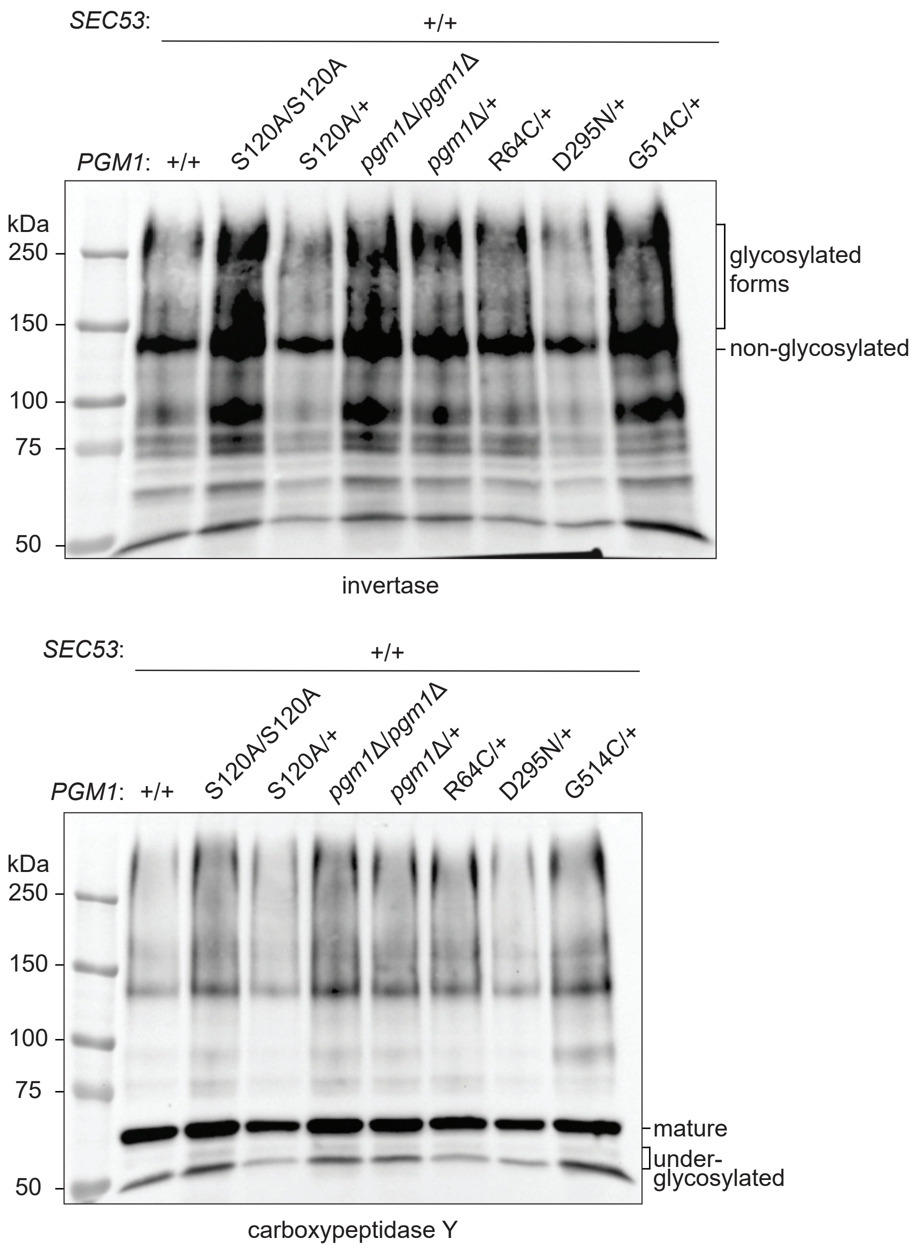


**Figure 3 — figure supplement 5. Effects of *pgm1* mutations on invertase and CPY glycosylation in the p*ACT1-SEC53*-WT background.** Western blots of invertase and CPY from constructed *SEC53*/*pgm1* strains.
