## Supplemental Data 10 for "Experimental evolution of phosphomannomutase-deficient yeast reveals compensatory mutations in a phosphoglucomutase"

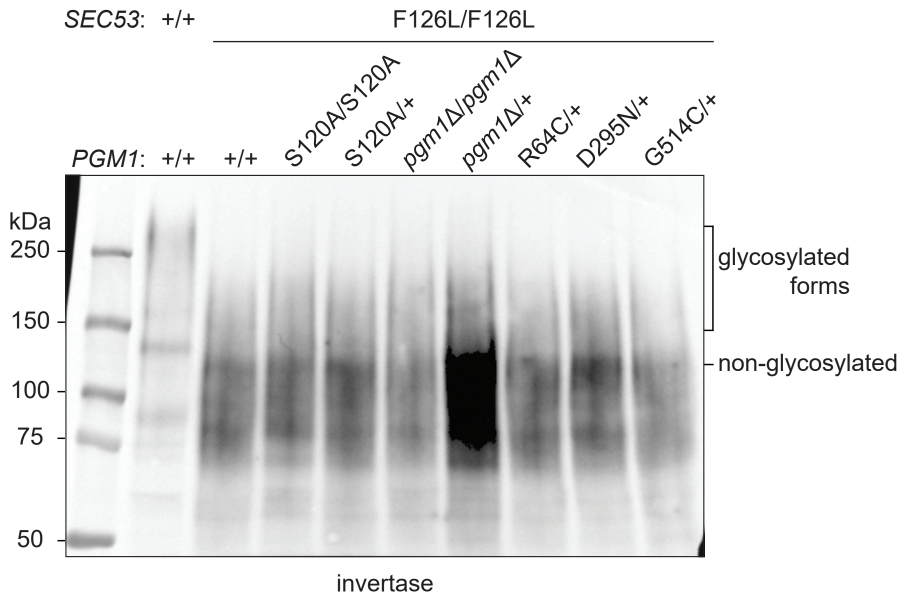


**Figure 3 — figure supplement 6. Effects of *pgm1* mutations on invertase glycosylation in the p*ACT1*-*sec53*-F126L background.** Western blot of invertase from reconstructed sec53/pgm1 strains.
