## Supplemental Data 11 for "Experimental evolution of phosphomannomutase-deficient yeast reveals compensatory mutations in a phosphoglucomutase"

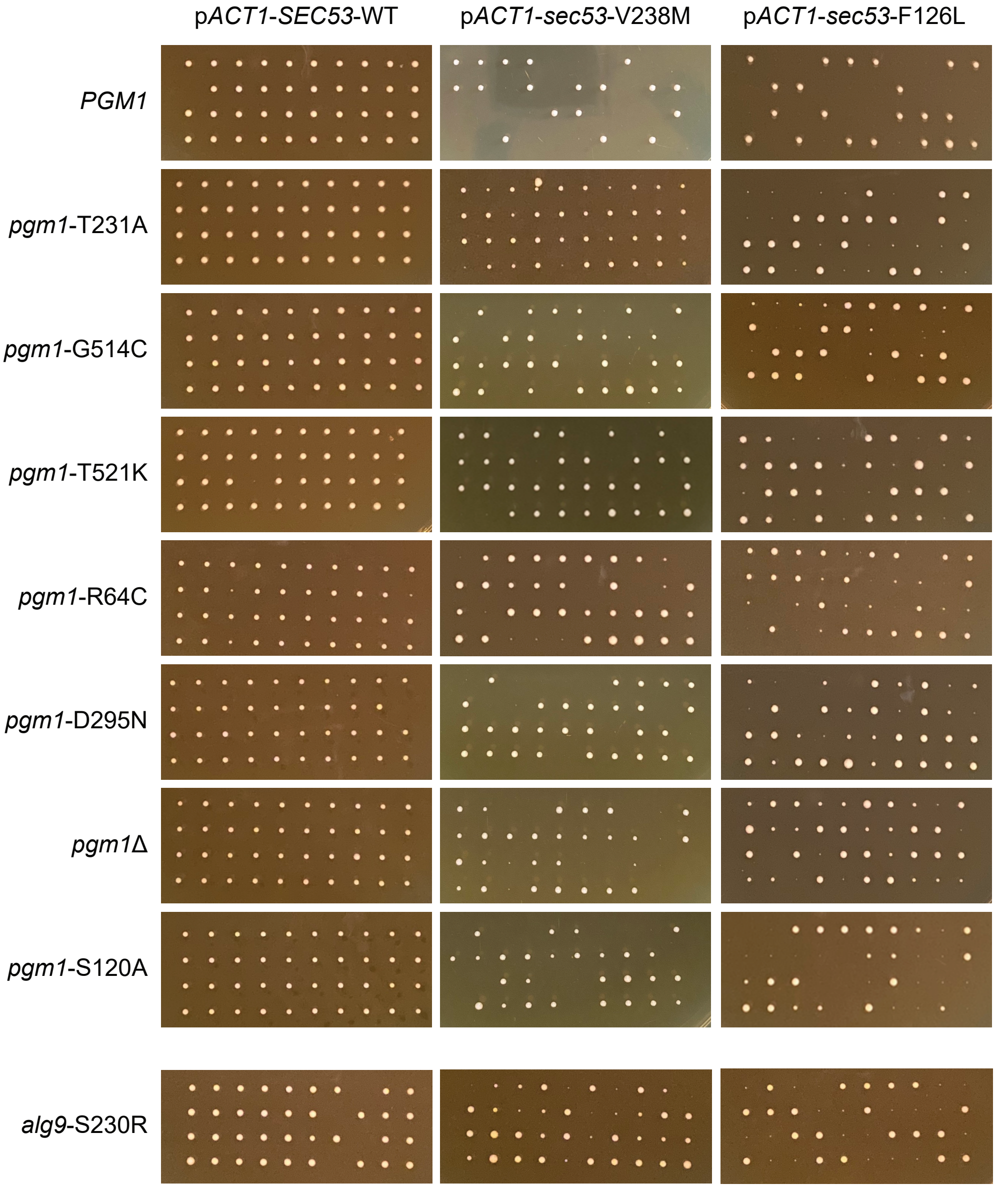


**Figure 4 — figure supplement 1. Tetrad dissections of *PGM1*/*pgm1* and *ALG9*/*alg9* strains.** 10 tetrads were dissected for each strain. Spores from the same tetrad are grouped vertically. The parental diploid strains contained heterozygous *SEC53*/*sec53* alleles and heterozygous *PGM1*/*pgm1* or *ALG9*/*alg9* alleles.
