## Supplemental Data 12 for "Experimental evolution of phosphomannomutase-deficient yeast reveals compensatory mutations in a phosphoglucomutase"

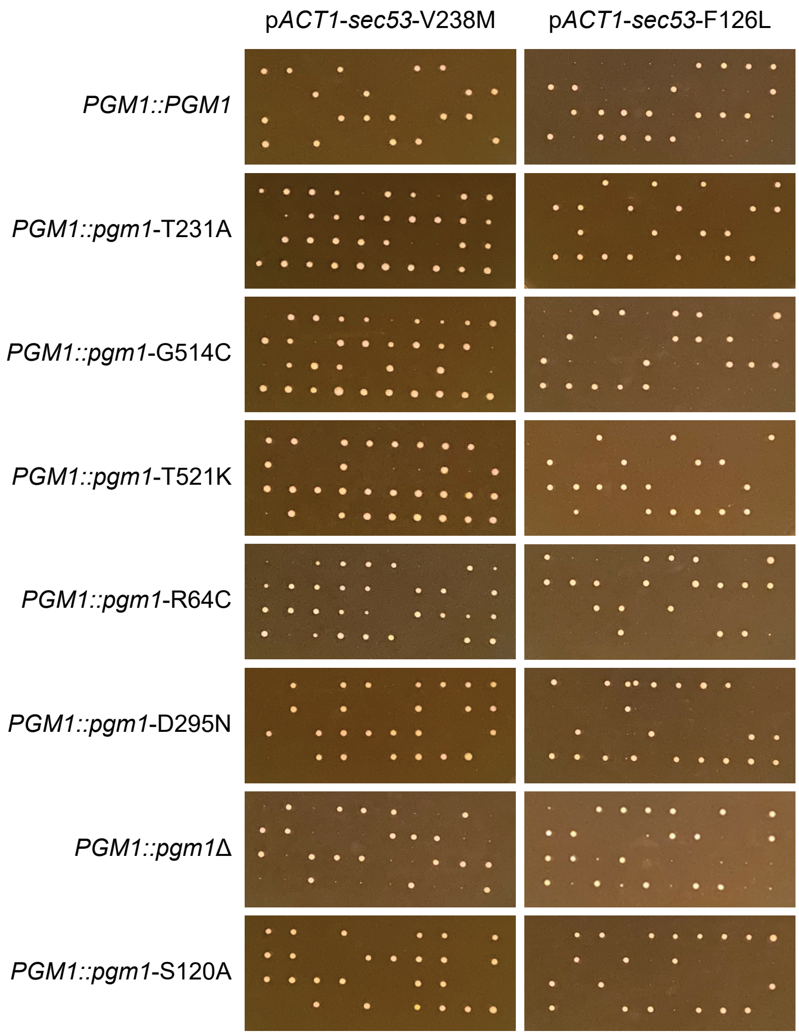


**Figure 4 — figure supplement 2. Tetrad dissections of *PGM1*-linked strains.** 10 tetrads were dissected for each strain. Spores from the same tetrad are grouped vertically. The parental diploid strains contained heterozygous *SEC53*/*sec53* alleles and 4 copies of *PGM1*; a wild-type copy of *PGM1* was introduced next to the endogenous *PGM1* ORF. Each haploid spore, then, contained either two wild-type copies (*PGM1*::*PGM1*) or a wild-type and mutant copy (*PGM1*::*pgm1*).
