## Supplemental Data 13 for "Experimental evolution of phosphomannomutase-deficient yeast reveals compensatory mutations in a phosphoglucomutase"

### coupled spectrophotometric phosphoglucomutase assay

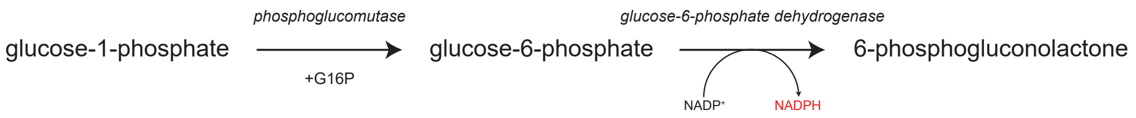


### $^{31}$P-NMR phosphomannomutase assay

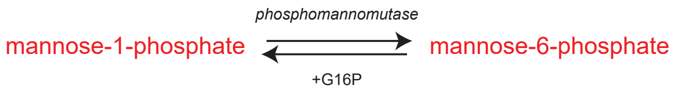


### coupled spectrophotometric bisphosphatase assay

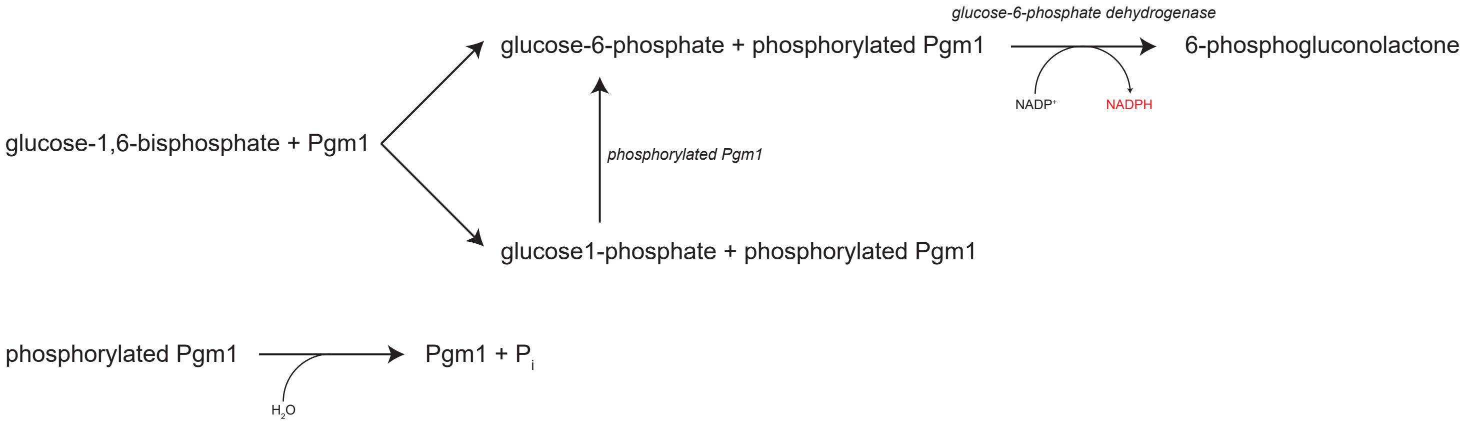


### $^{31}$P-NMR phosphoglucomutase assay

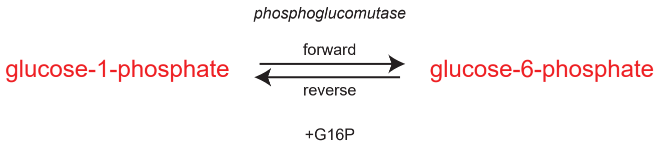


**Figure 5 — figure supplement 1. Enzymatic assays.** Diagrams of the enzymatic reactions assayed in this study.
