## Supplemental Data 14 for "Experimental evolution of phosphomannomutase-deficient yeast reveals compensatory mutations in a phosphoglucomutase"

| Protein | Relative Activity (compared to G1P 300 μM, G16P 20 μM) | | | $EC_{50}$ for G16P (μM) |
|---|---|---|---|---|
| | G1P 300 μM G16P 20 μM | G1P 300 μM G16P 20 μM +EDTA 10 mM | G1P 300 μM G16P 0 μM | G1P 300 μM |
| Pgm1-WT | 1.00 | 0.015 | <0.01 | 0.86 ± 0.08 |
| Pgm1-T231A | 1.00 | 0.04 | 0.012 | 0.43 ± 0.07 |
| Pgm1-G514C | 1.00 | 0.16 | <0.01 | 2.07 ± 0.13 |
| Pgm1-T521K | 1.00 | 0.04 | n.d. | n.d. |

**Figure 5 — figure supplement 2. Other biochemical properties of mutant Pgm1.** The spectrophotometric-coupled assay was used to measure phosphoglucomutase activity. The concentration at which G16P exerts half of its maximal effect ($EC_{50}$) was obtained by fitting the data (activity vs G16P concentration) to the Michaelis–Menten equation. $EC_{50}$ values ± 95% confidence intervals. Relative activity and G16P $EC_{50}$ of Pgm1-T521K were not determined (n.d.).
