## Supplemental Data 15 for "Experimental evolution of phosphomannomutase-deficient yeast reveals compensatory mutations in a phosphoglucomutase"

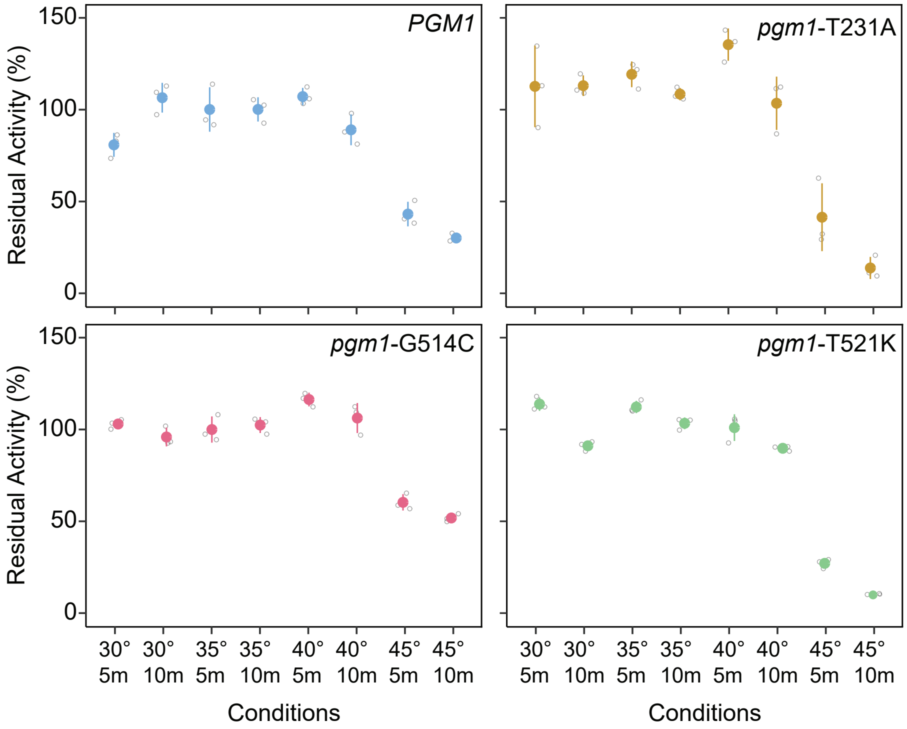


**Figure 5 — figure supplement 3. Thermostabilities of mutant Pgm1.** Average residual activity and standard deviations of recombinant Pgm1 enzymes determined after incubation at the conditions described on the x-axis. Replicate measurements plotted as gray circles. Proteins were incubated in the presence BSA (0.1 mg/ml).
