## Supplemental Data 16 for "Experimental evolution of phosphomannomutase-deficient yeast reveals compensatory mutations in a phosphoglucomutase"

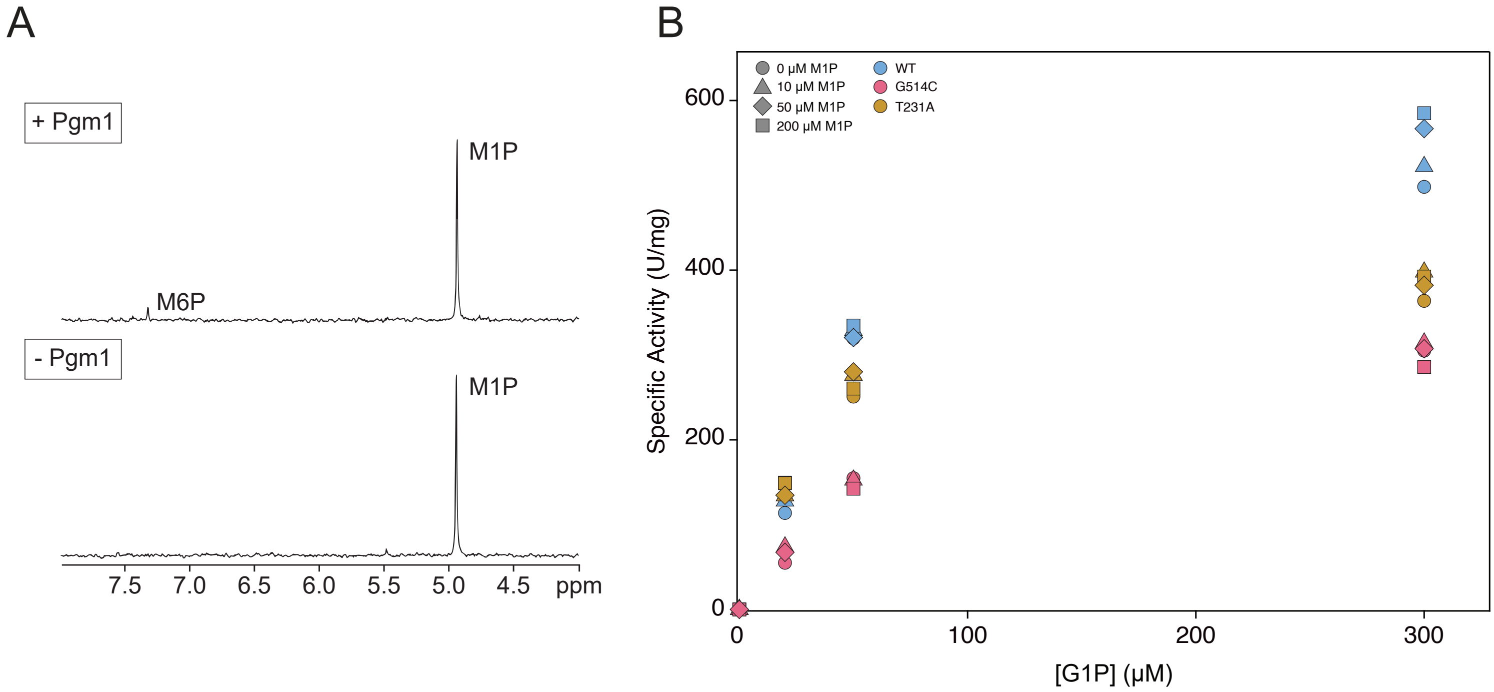


**Figure 5 — figure supplement 4. Phosphomannomutase activity of mutant Pgm1.** (A) Representative phosphomannomutase assay by $^{31}P$-NMR spectroscopy. Pgm1 was incubated with 1mM M1P and 20 µM G16P for known periods of time. The amount of M1P or M6P was measured by integrating the area of the signals and comparing them to creatine phosphate, added as an internal standard. (B) Spectrophotometric phosphoglucomutase assays were conducted at different concentrations of G1P and M1P. Shapes indicate varying concentrations of M1P and colors indicate Pgm1 variants.
