## Supplemental Data 17 for "Experimental evolution of phosphomannomutase-deficient yeast reveals compensatory mutations in a phosphoglucomutase"

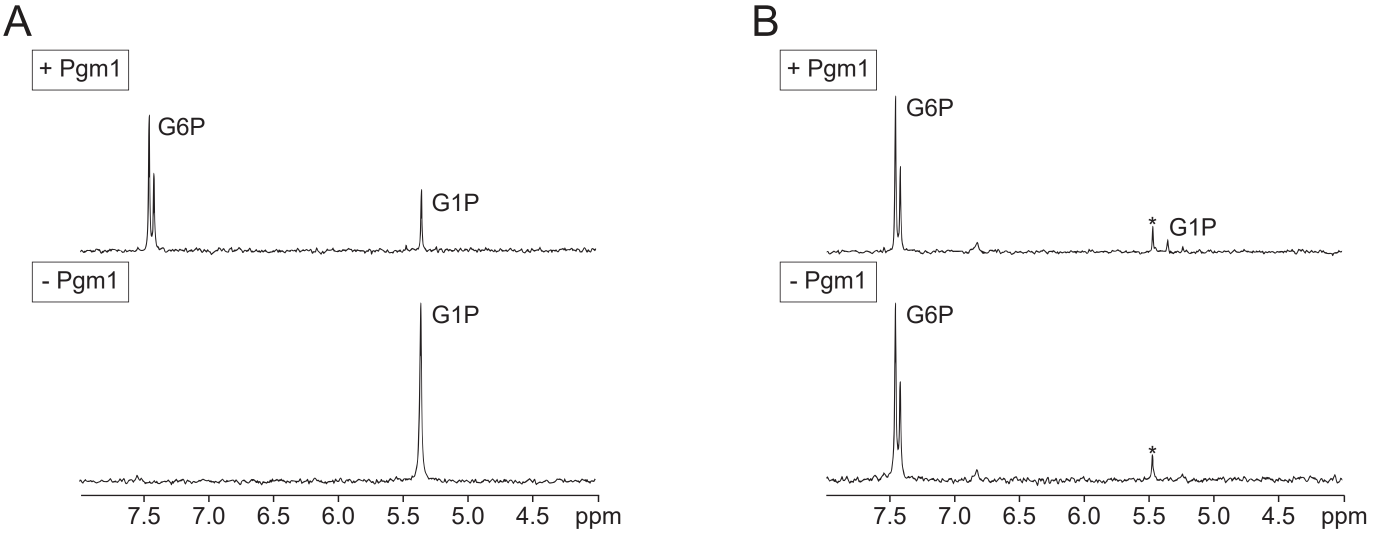


**Figure 5 — figure supplement 5. Representative forward/reverse phosphoglucomutase assay.** Equal amounts of Pgm1 were incubated with G1P (A) or G6P (B) in the presence of G16P. The amount of the substrates and products were measured by integrating the area of the signals and comparing them to creatine phosphate, added as an internal standard. Asterisks (*) indicate $P_i$ contaminants.
