## Supplementary figures and images for "Experimental evolution of phosphomannomutase-deficient yeast reveals compensatory mutations in a phosphoglucomutase"

### 1x_Wildtype_01.png

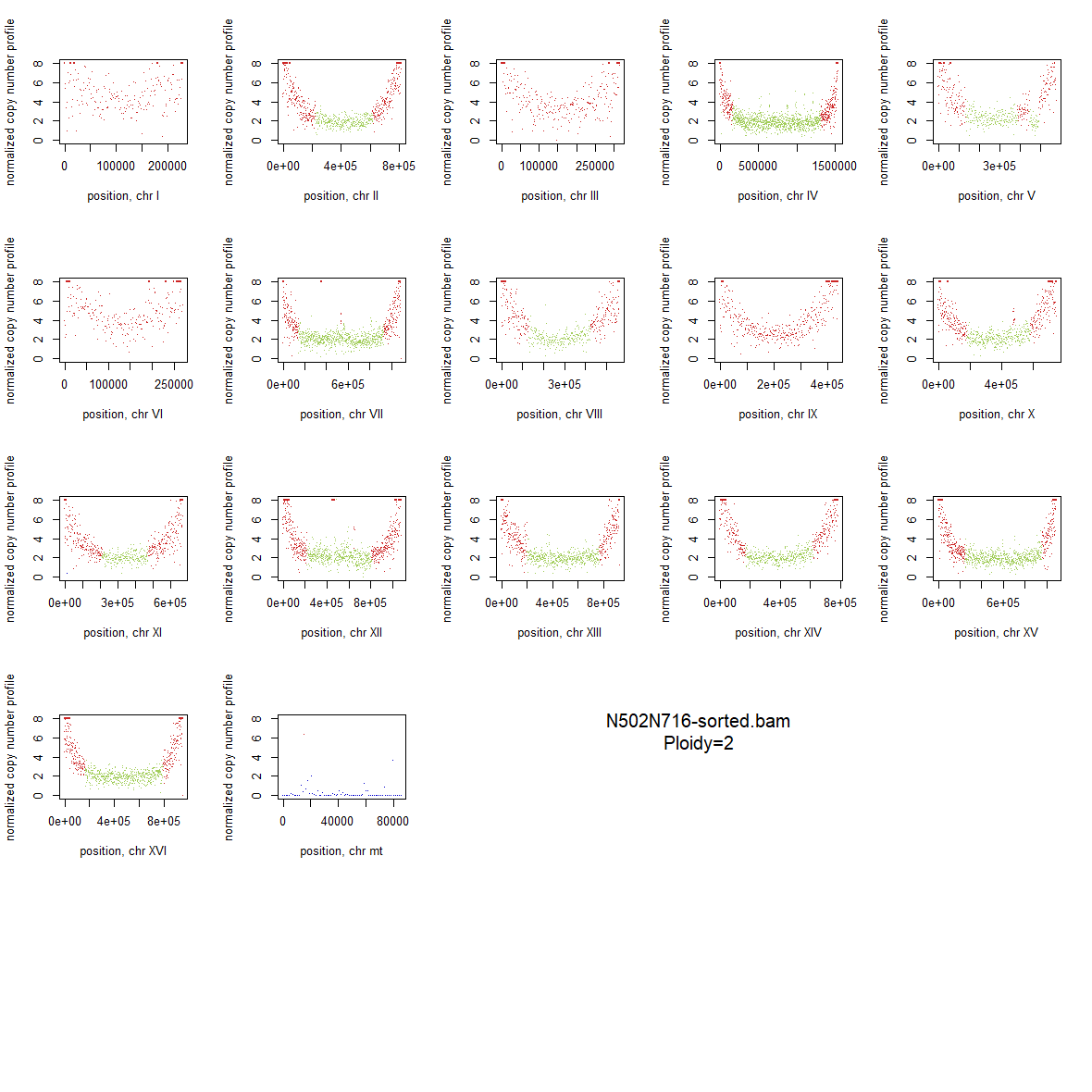

### 1x_Wildtype_02.png

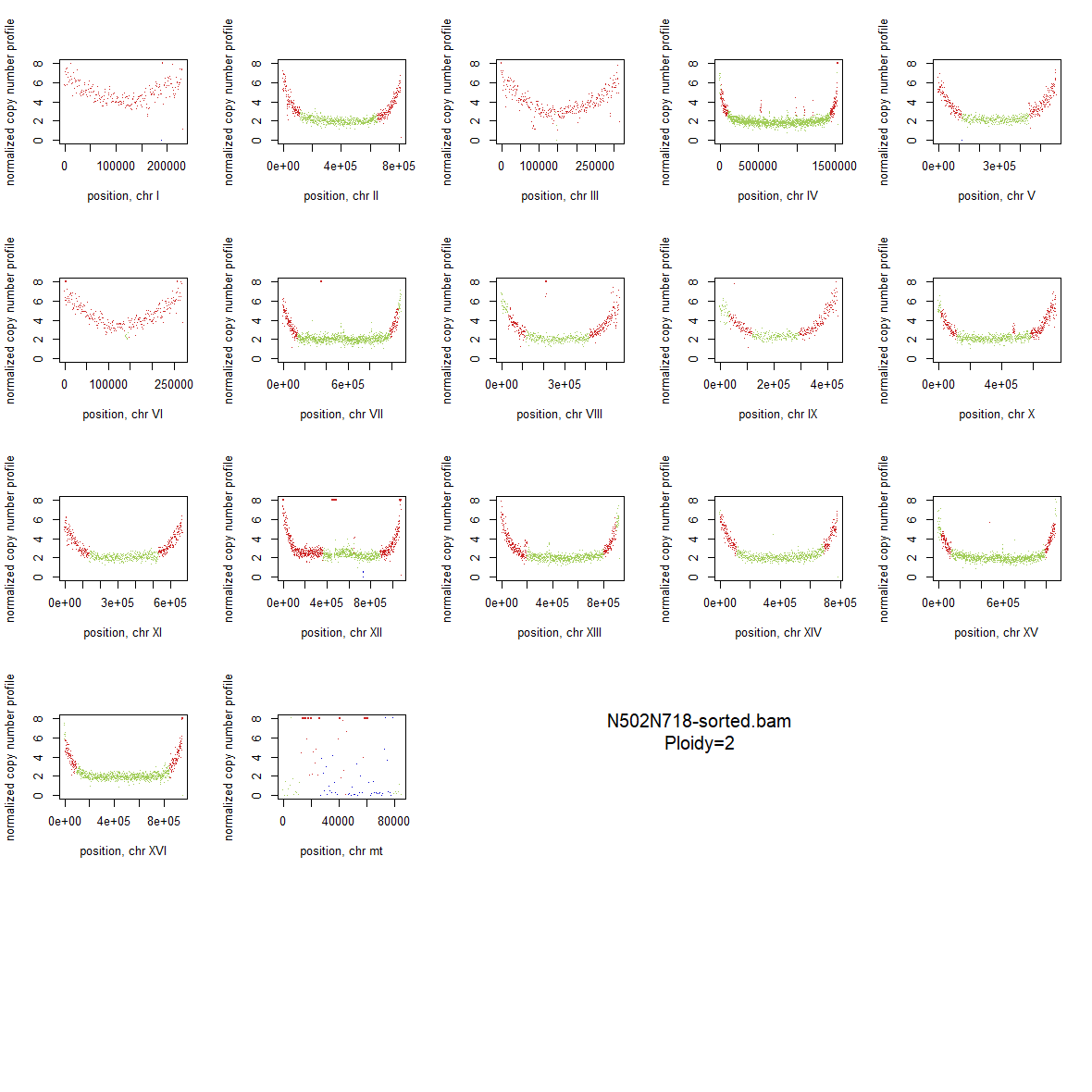

### 1x_Wildtype_03.png

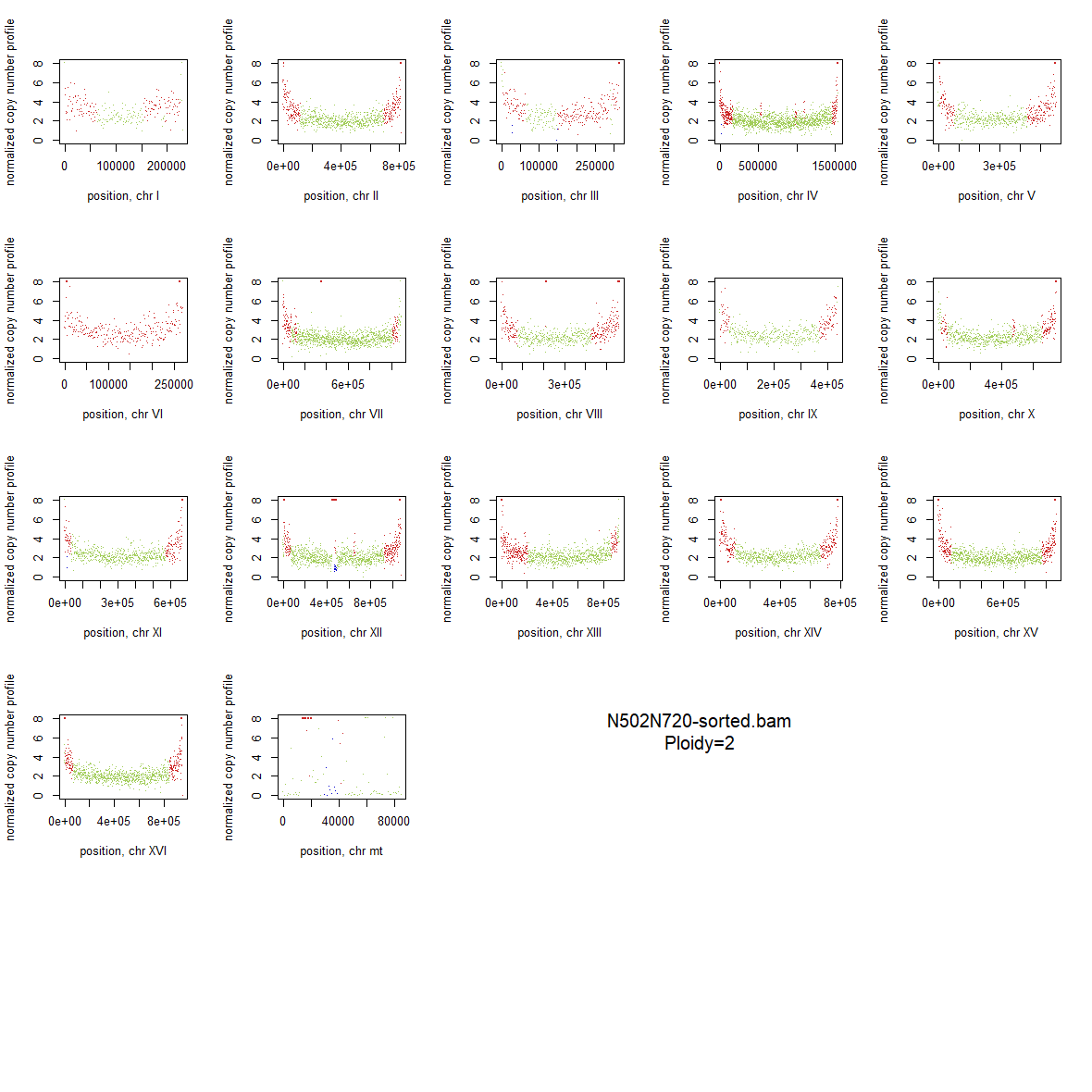

### 1x_Wildtype_04.png

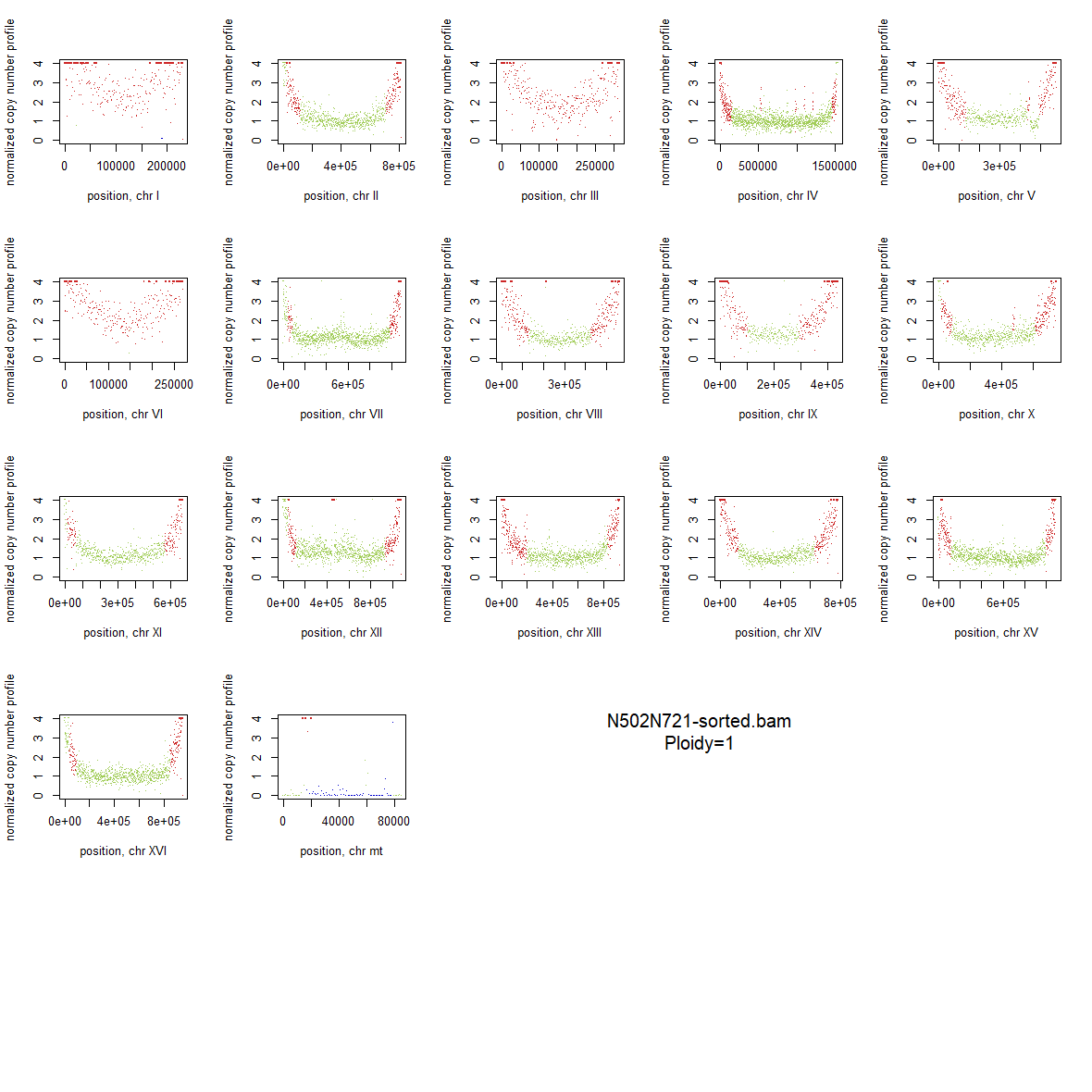

### 1x_Wildtype_05.png

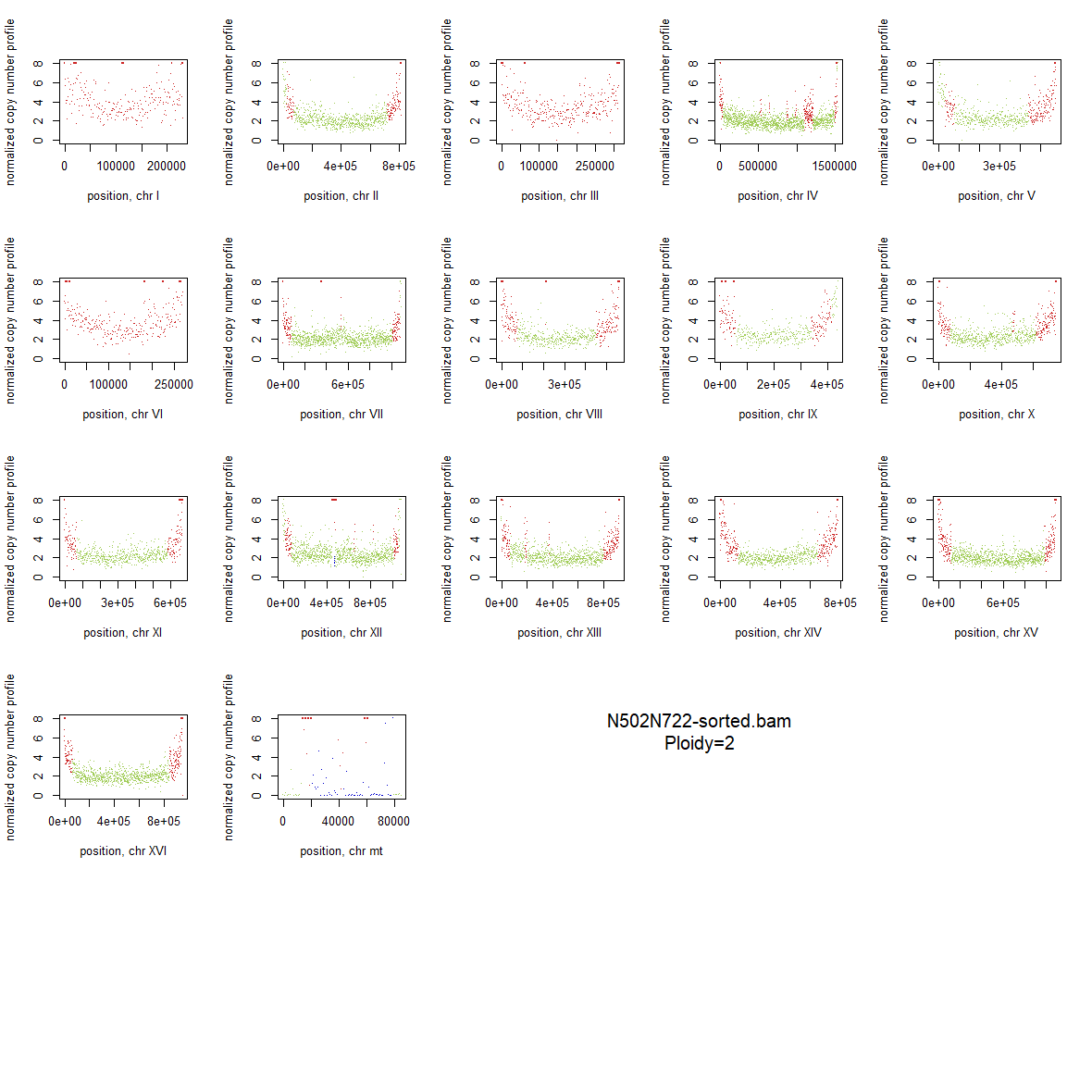

### 1x_Wildtype_06.png

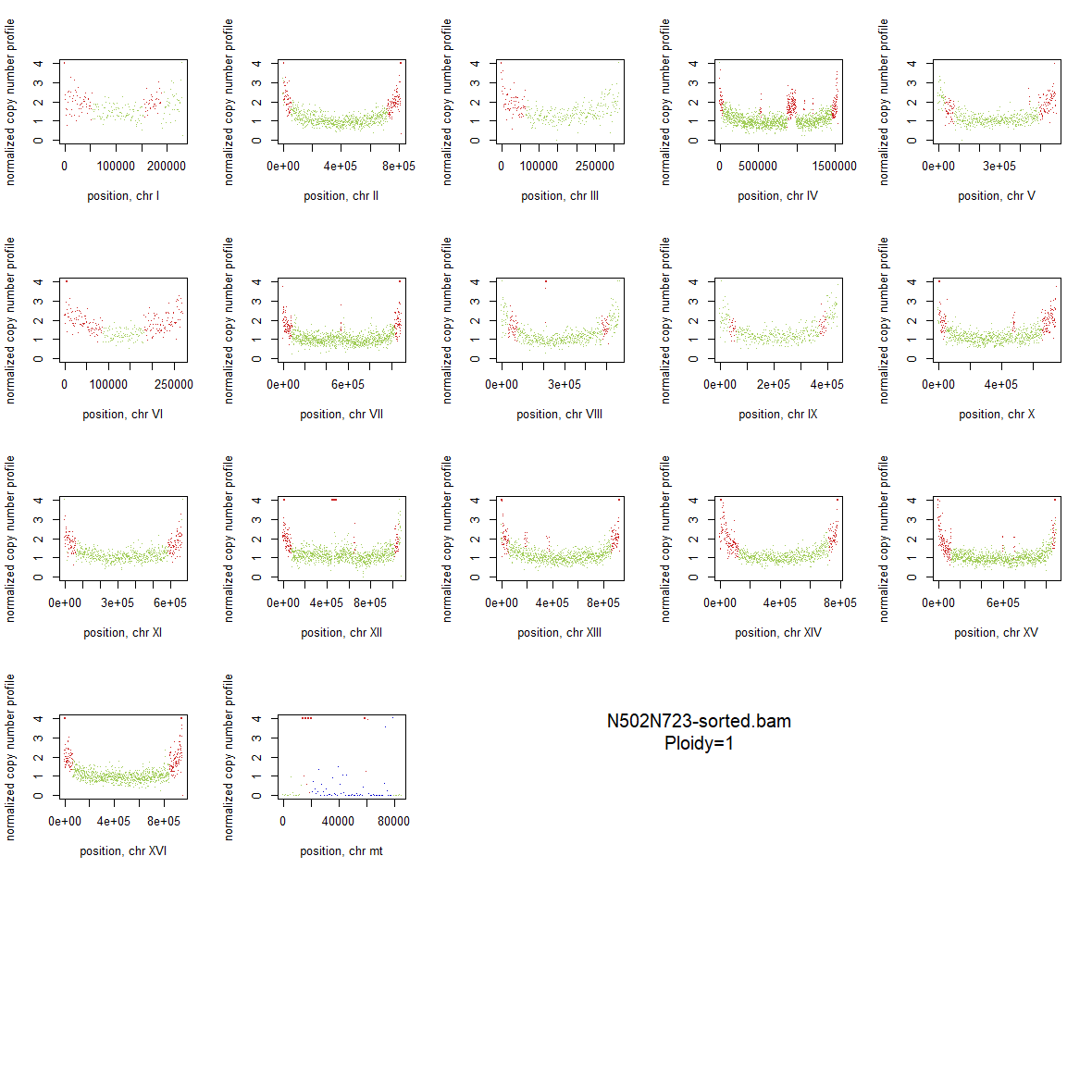

### 1x_Wildtype_07.png

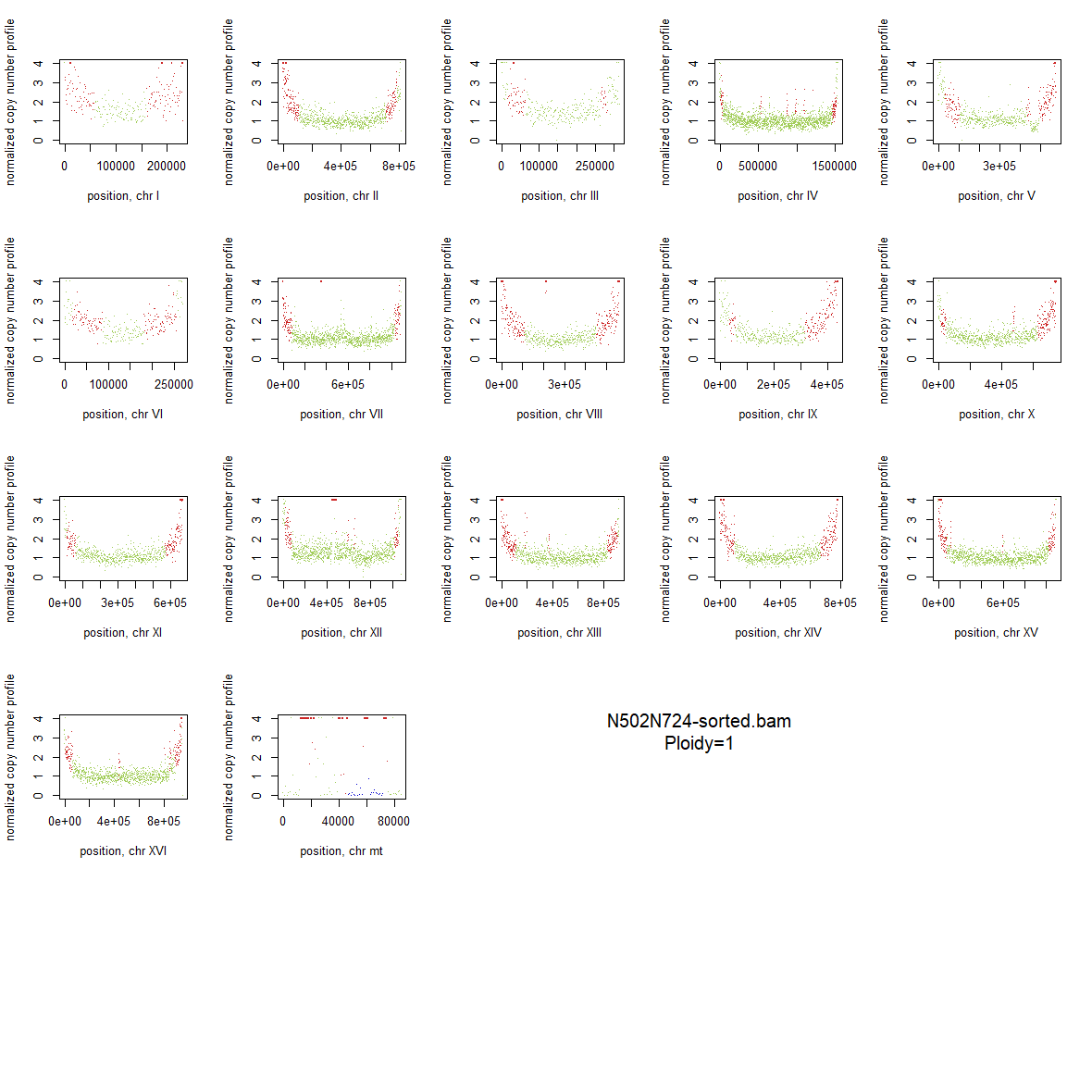

### 1x_Wildtype_08.png

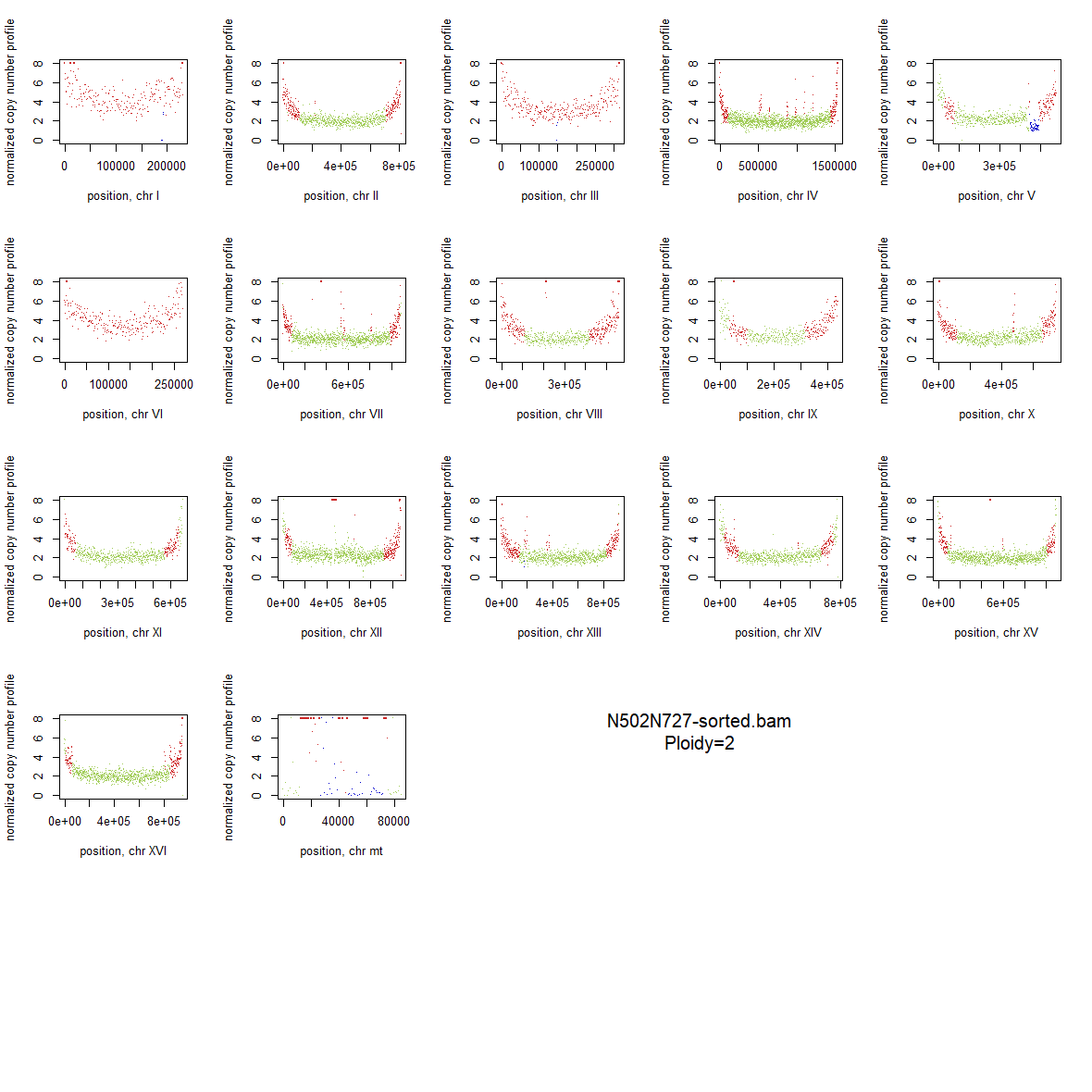

### 1x_Wildtype_09.png

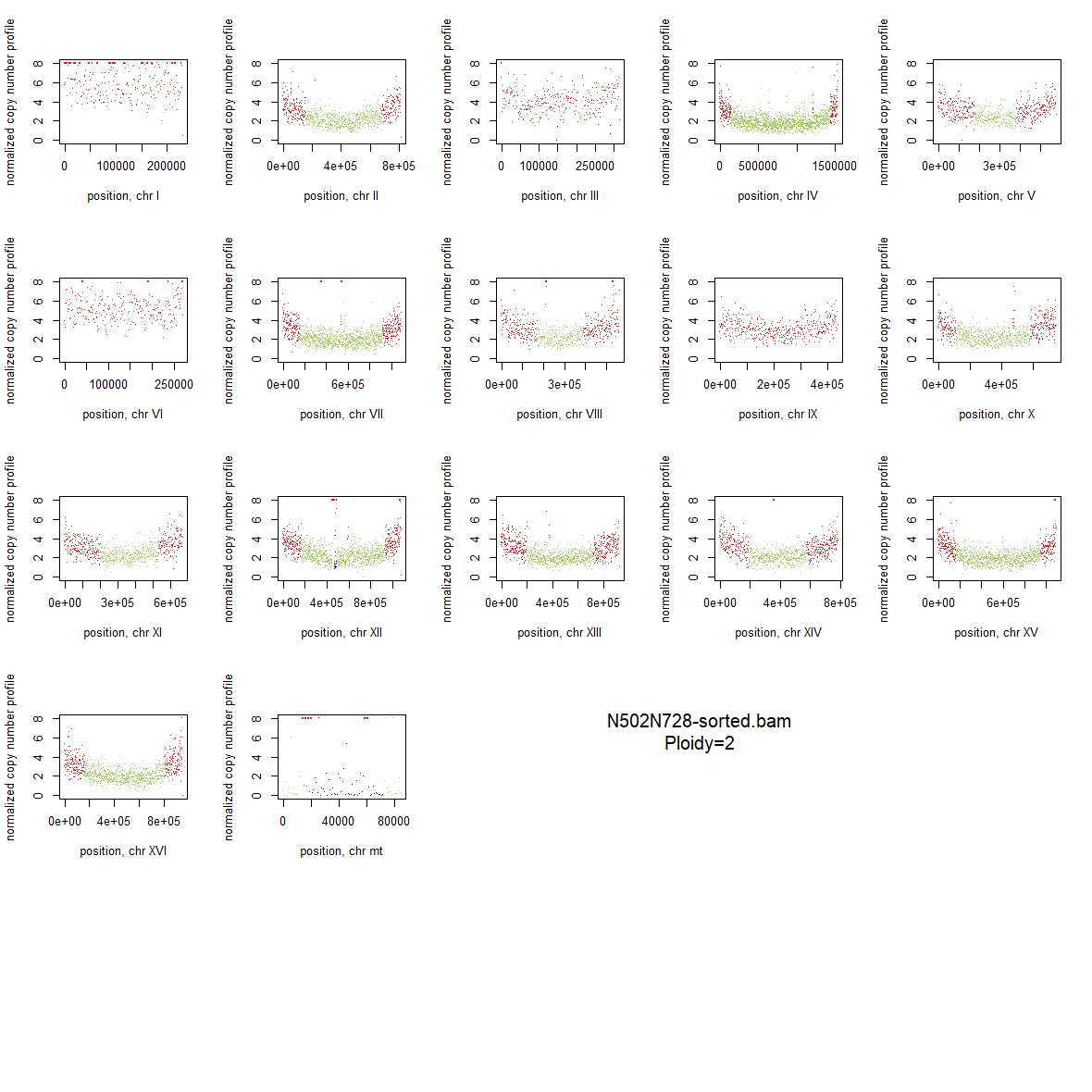

### 1x_Wildtype_10.png

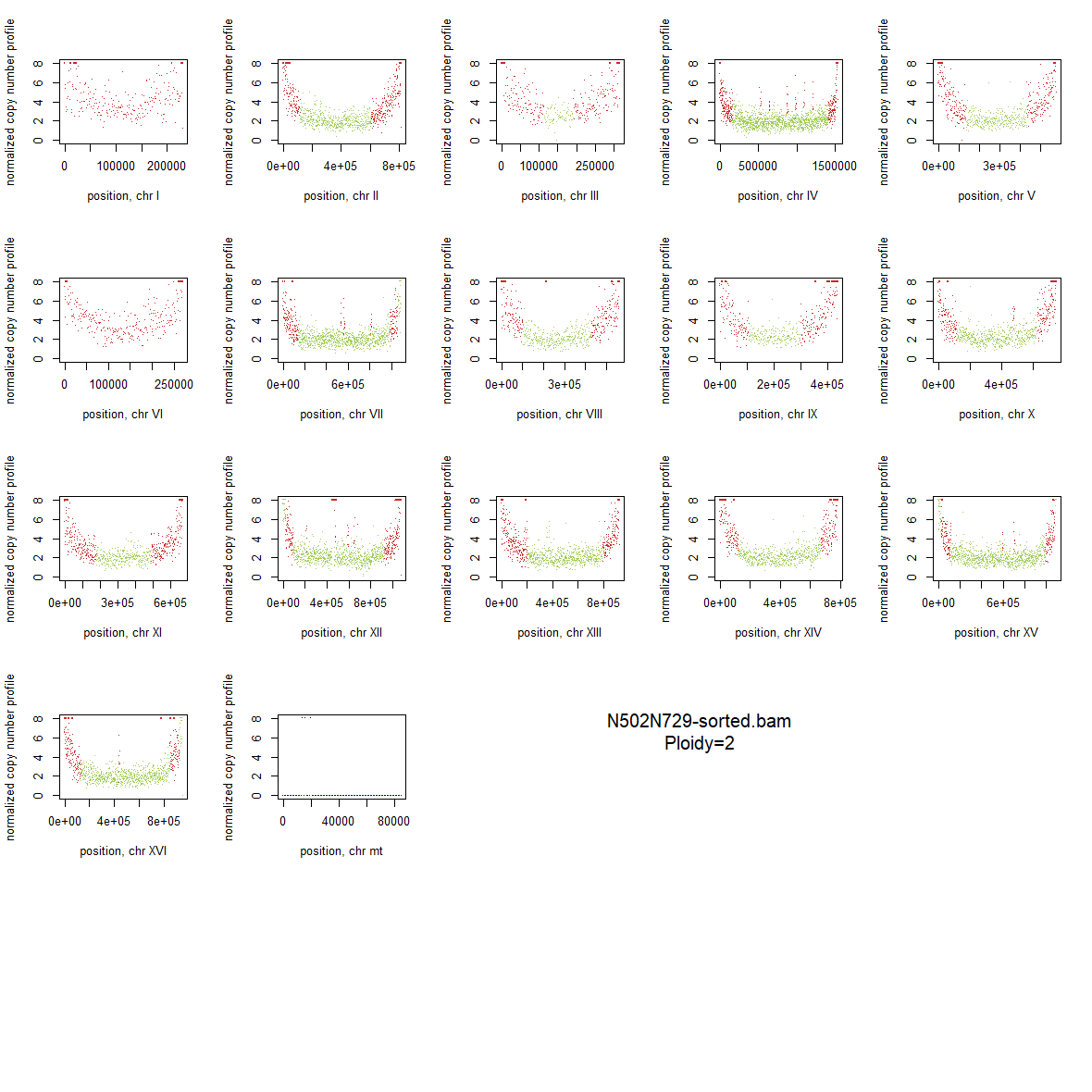

### 1x_Wildtype_11.png

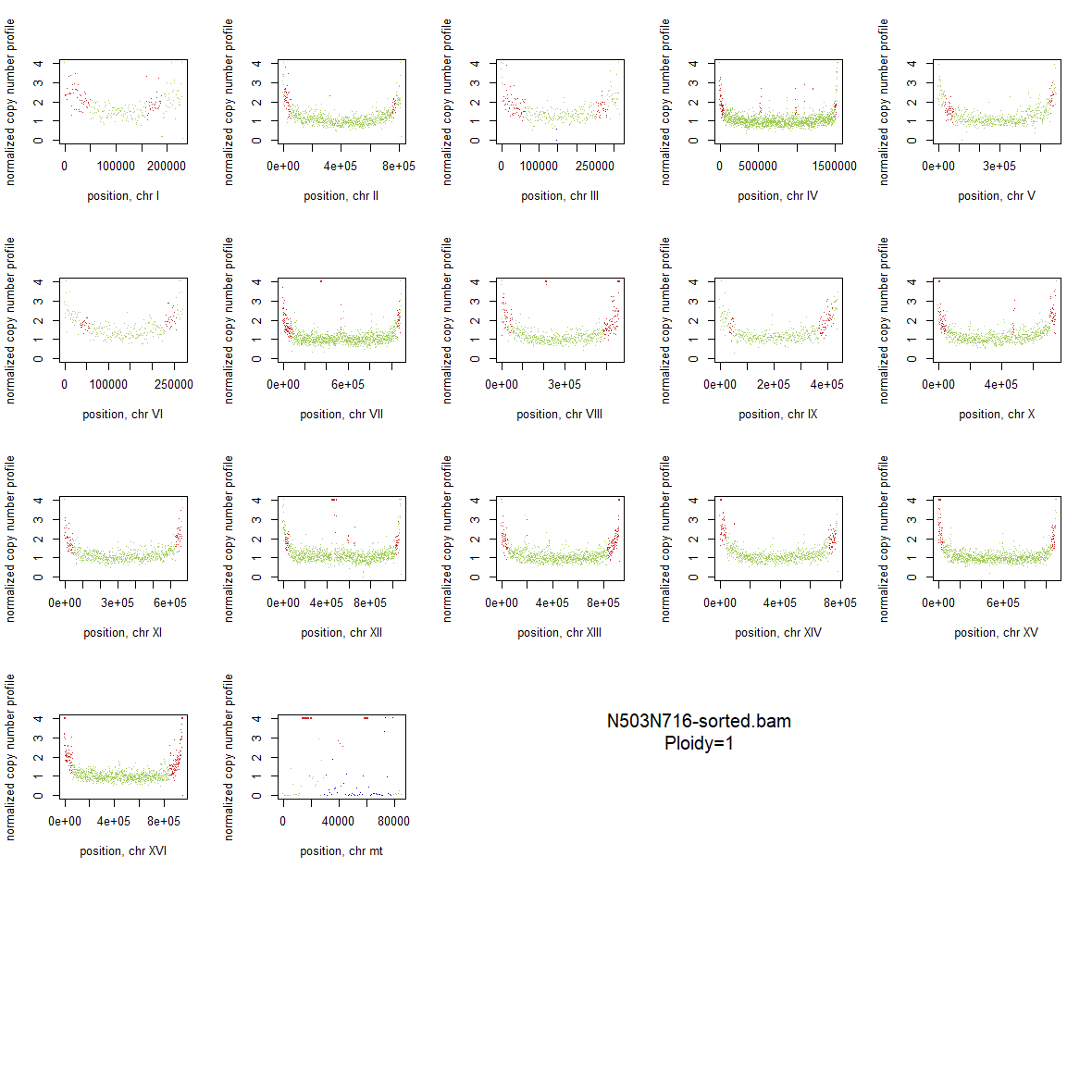

### 1x_Wildtype_12.png

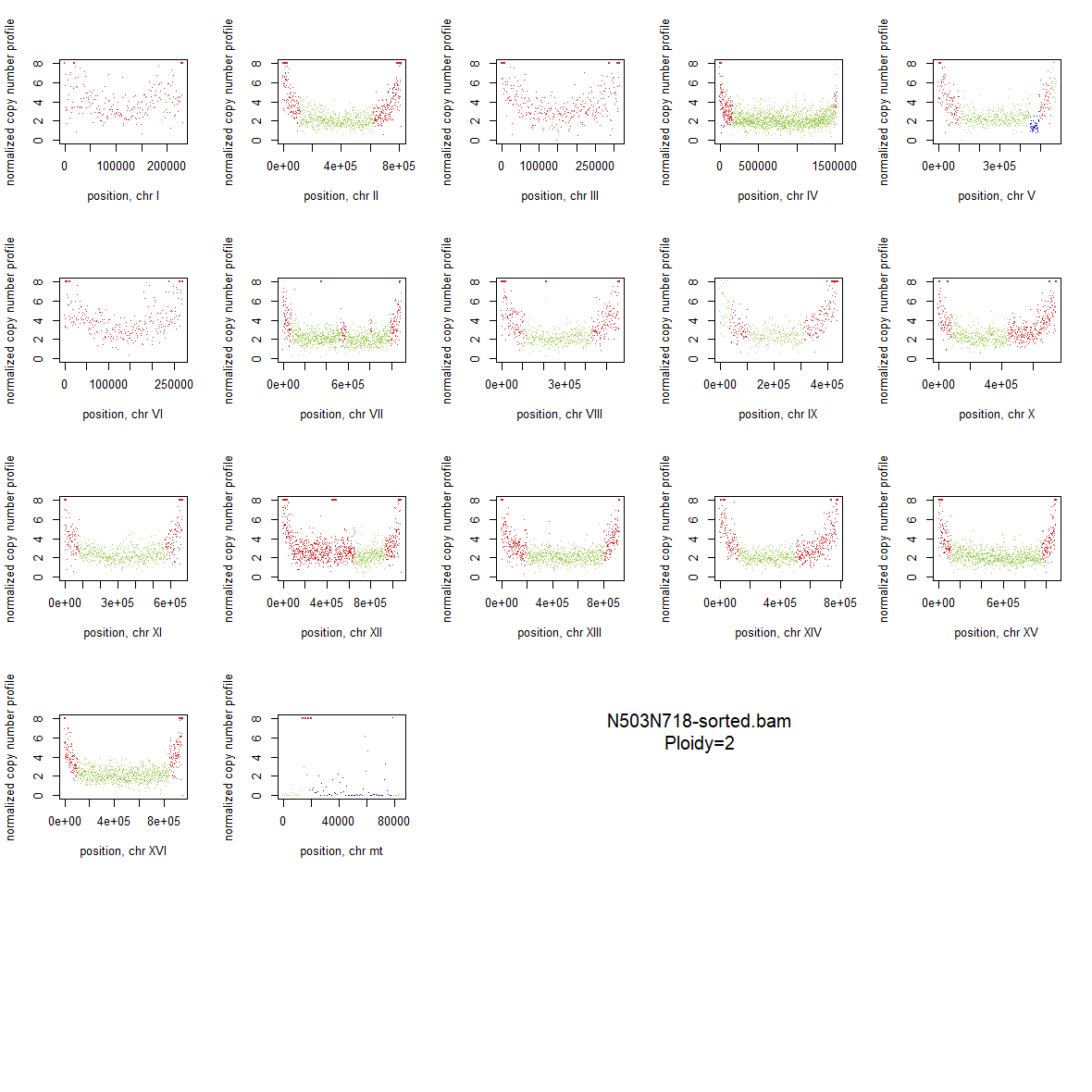

### 1x_Wildtype_13.png

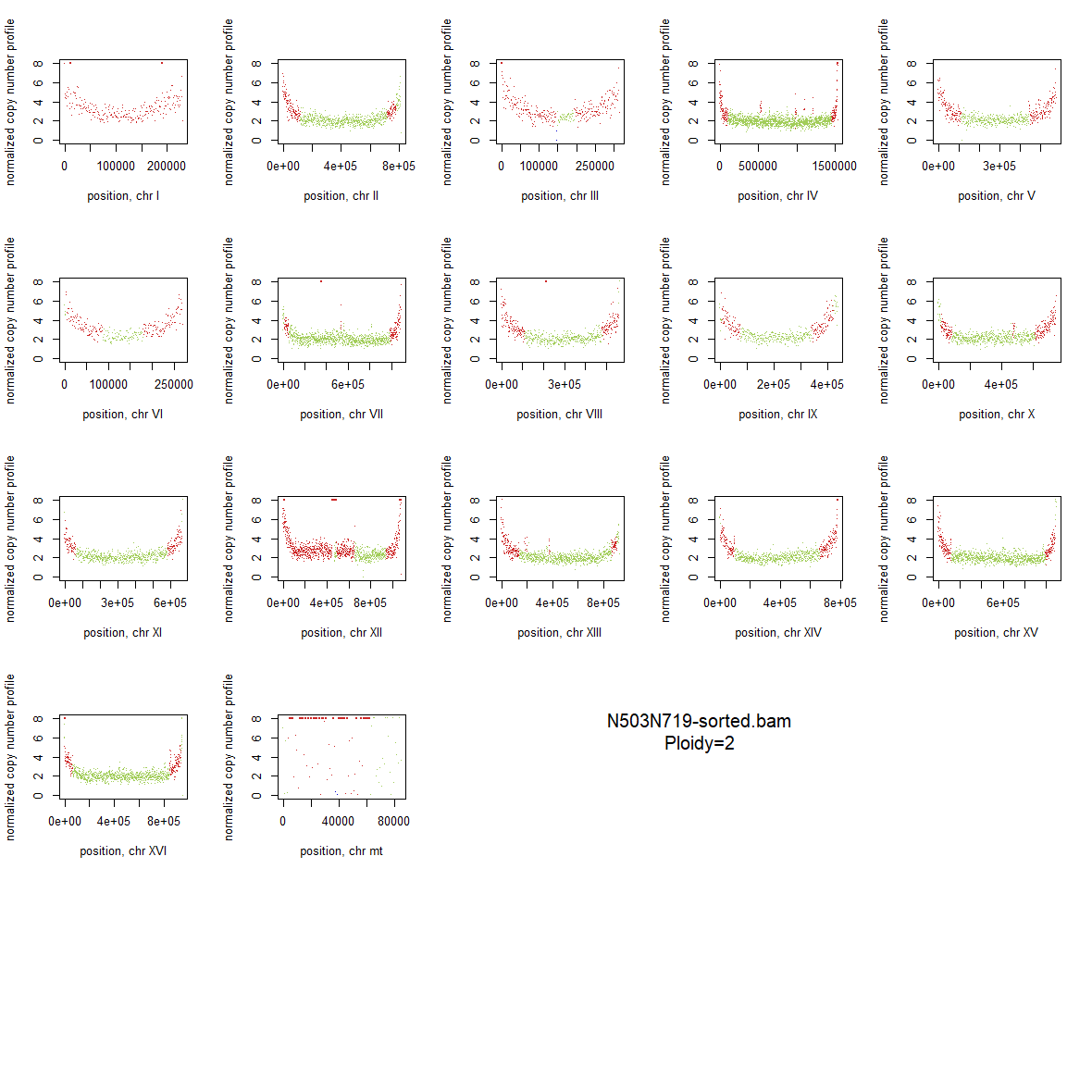

### 1x_Wildtype_14.png

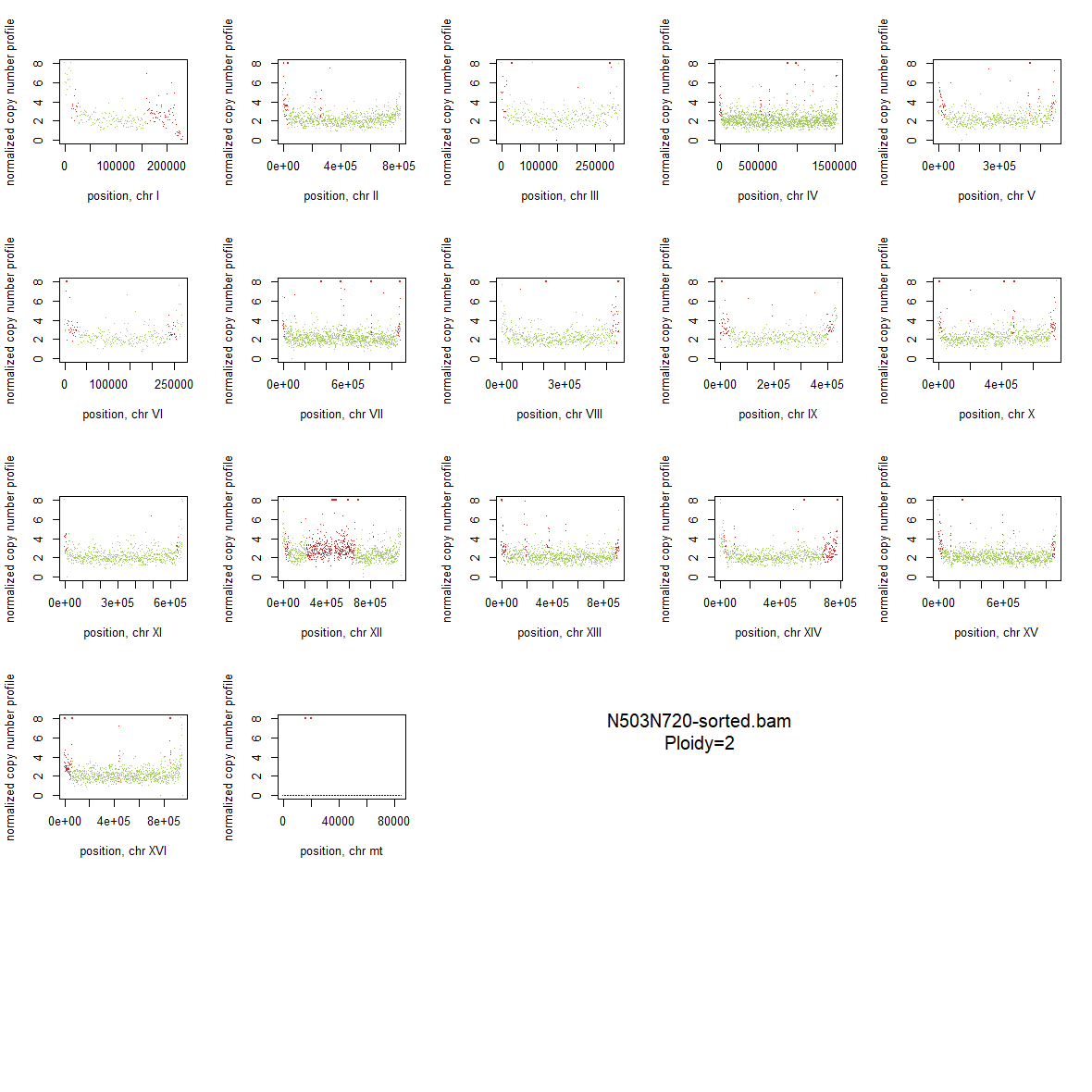

### 1x_Wildtype_15.png

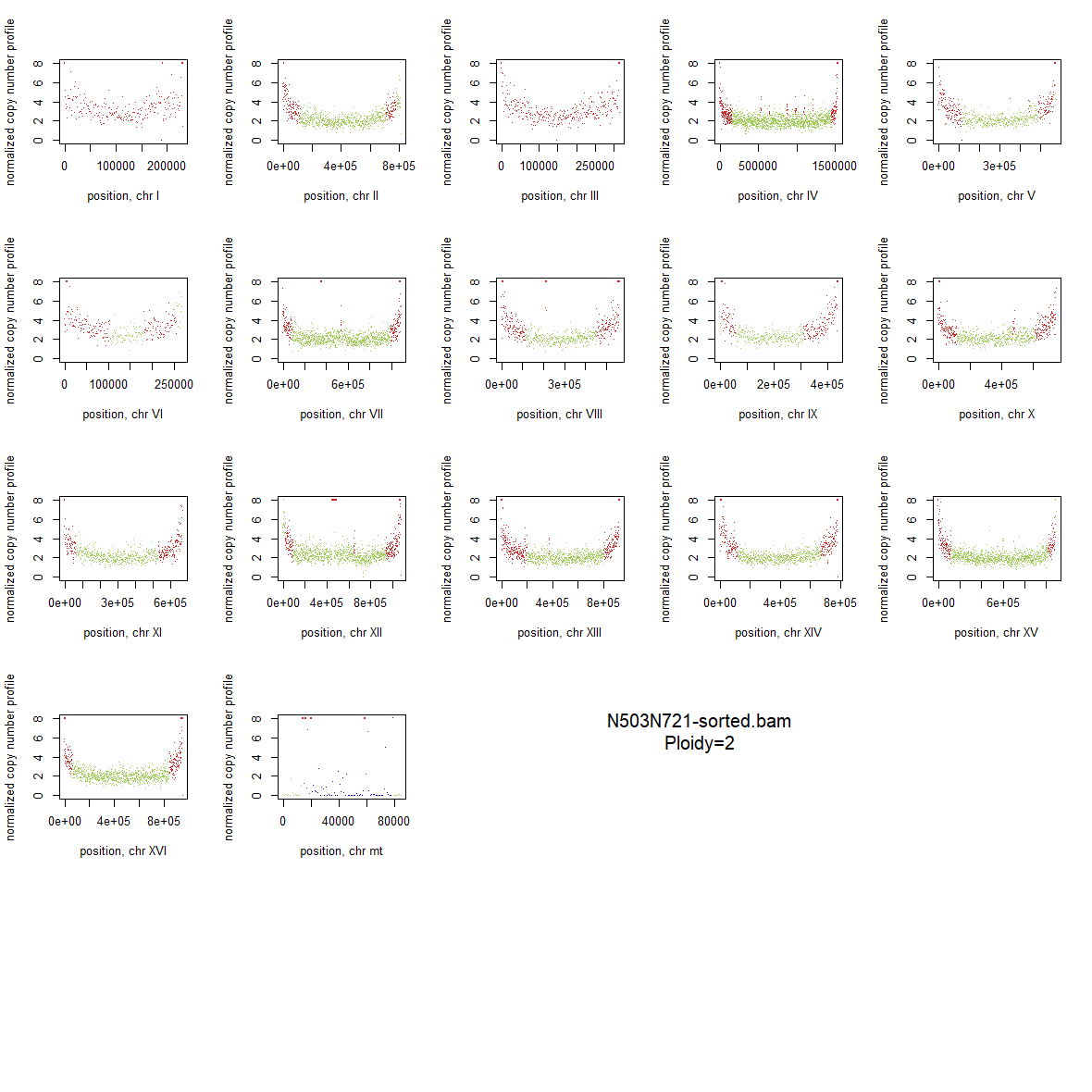

### 1x_Wildtype_16.png

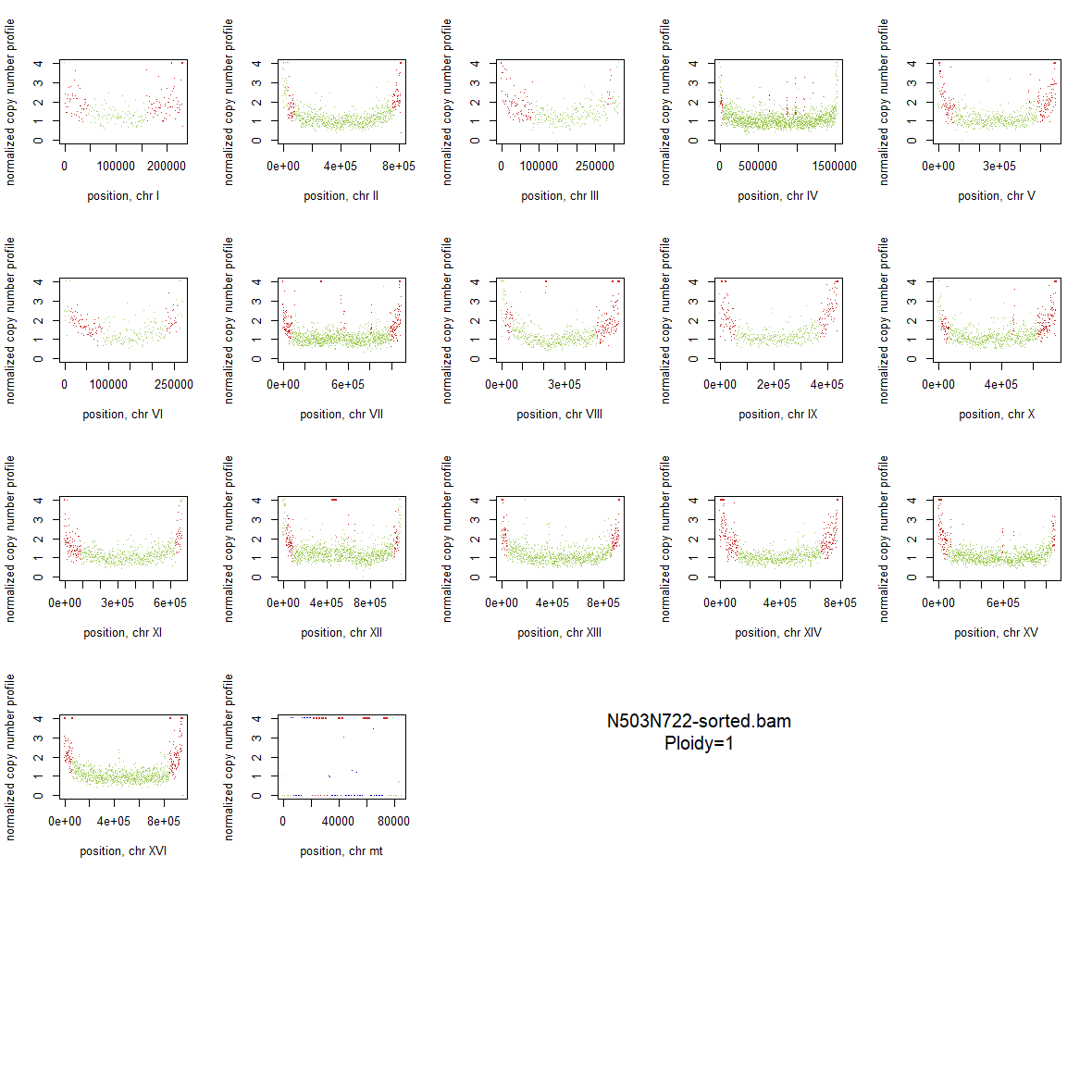

### 1x_Wildtype_17.png

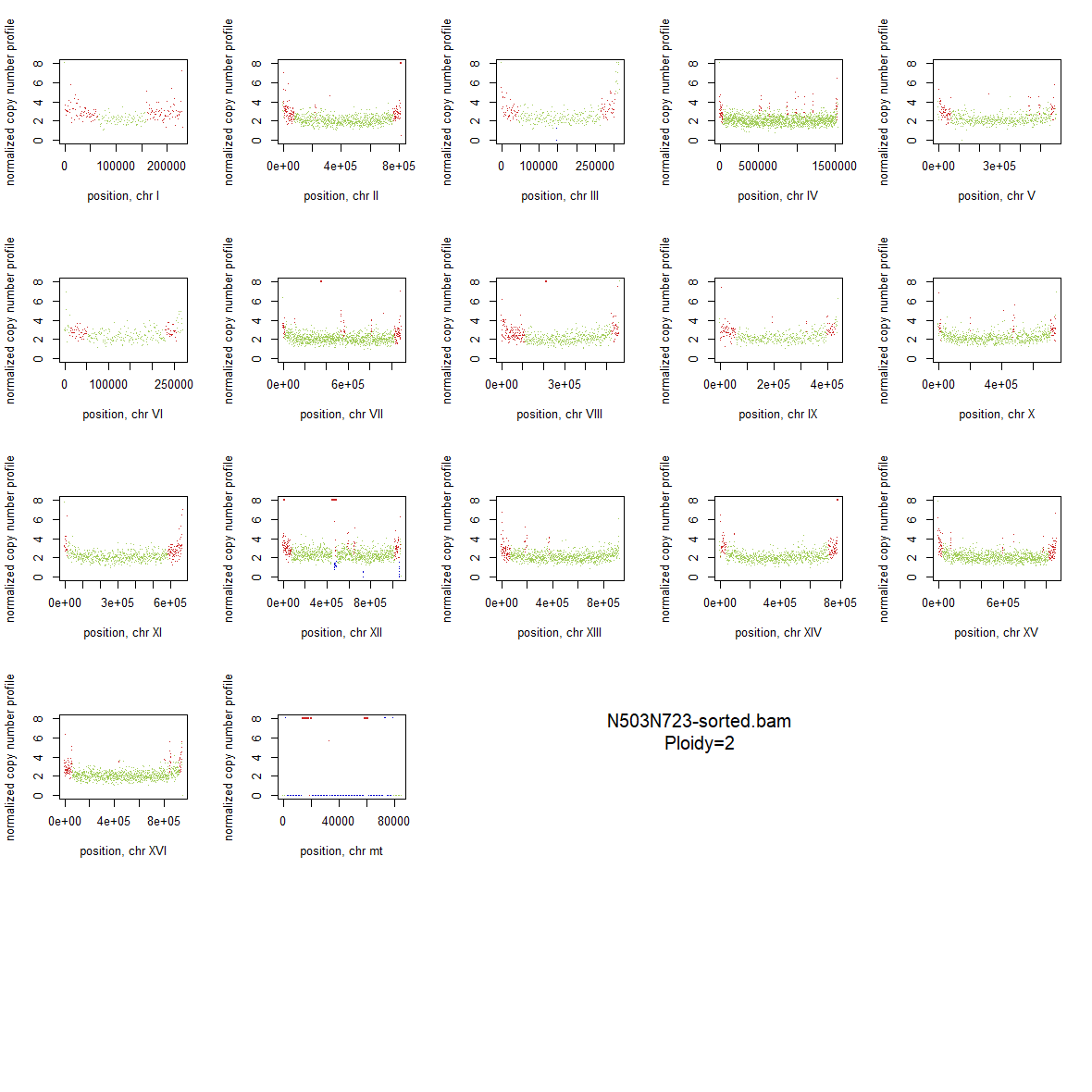

### 1x_Wildtype_18.png

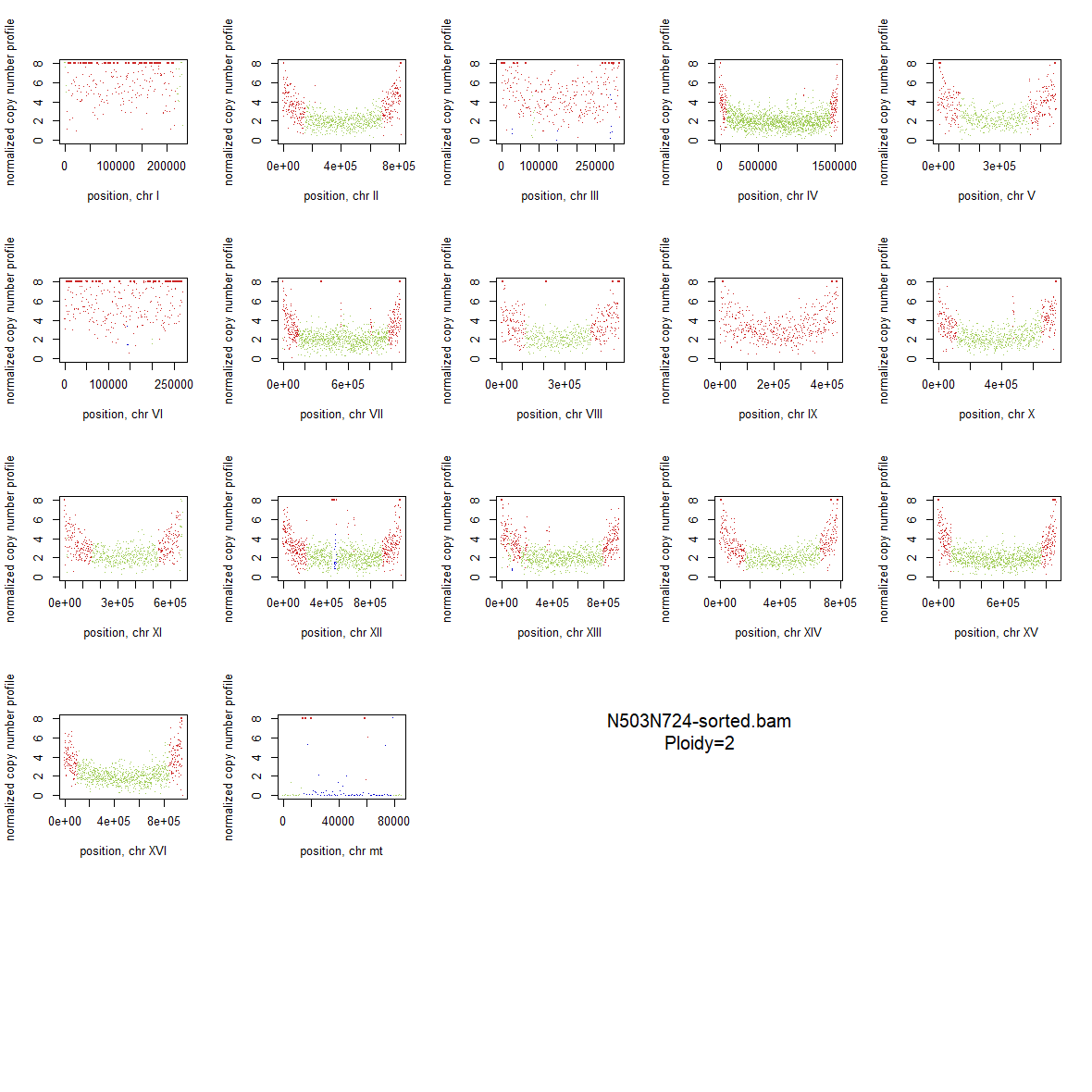

### 1x_Wildtype_19.png

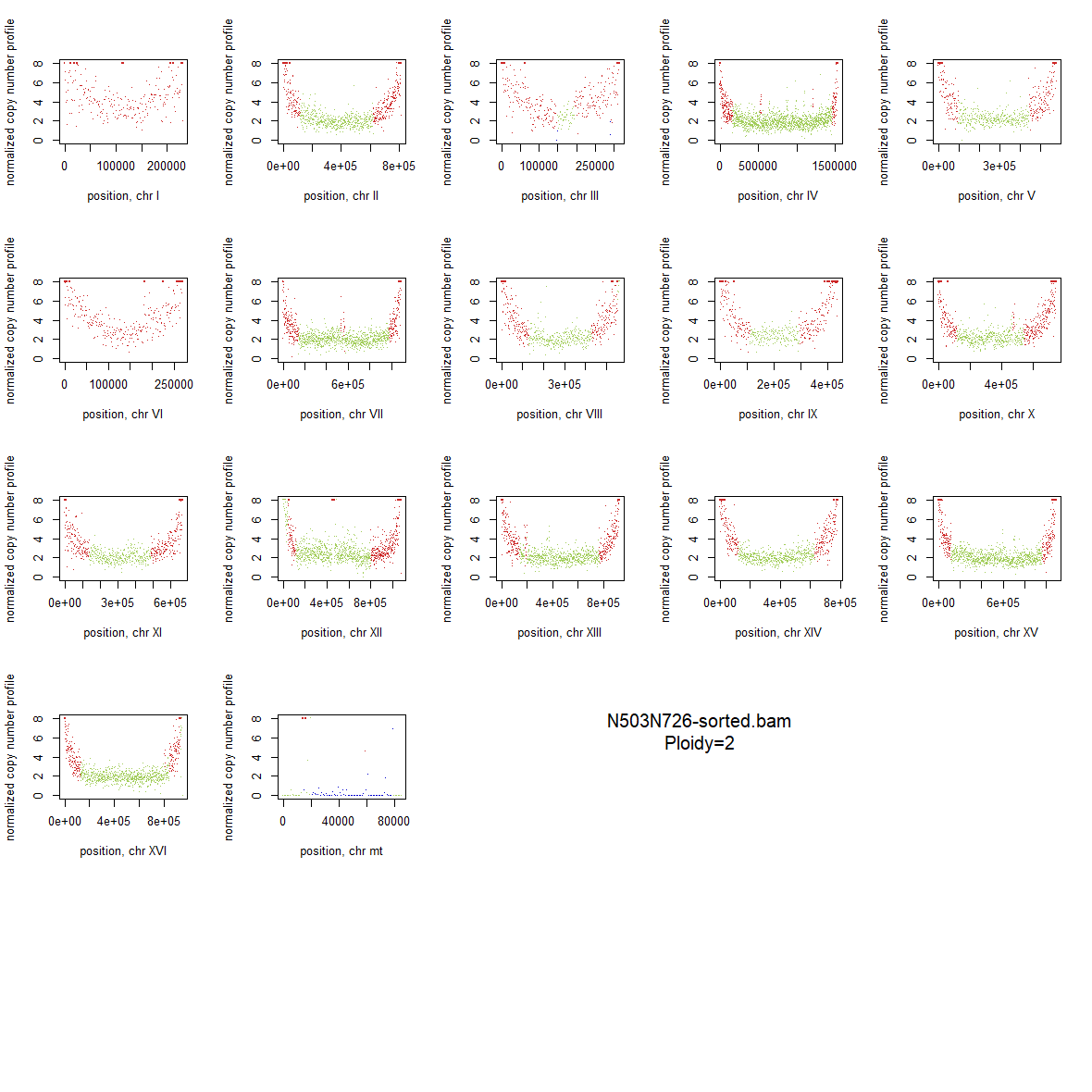

### 1x_Wildtype_20.png

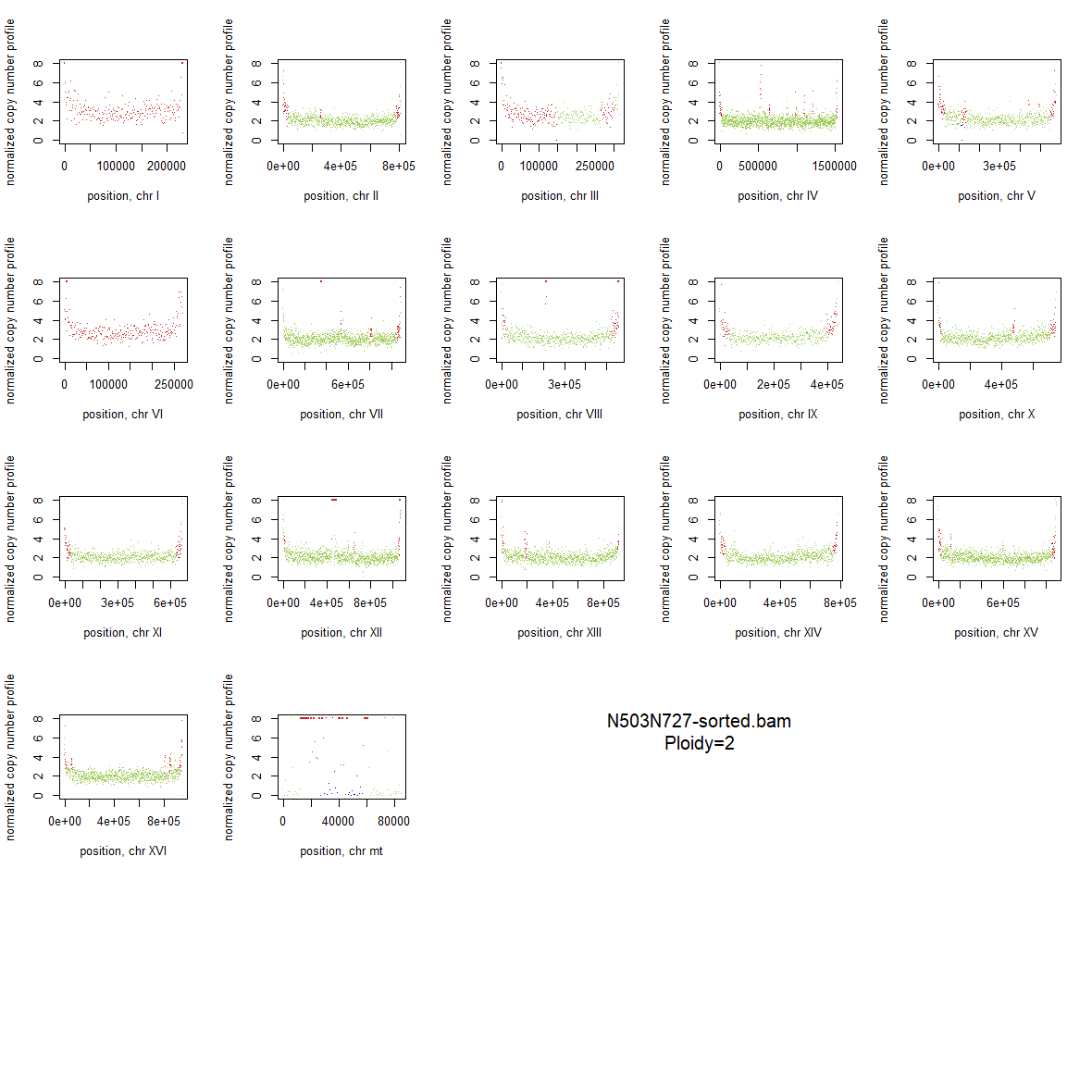

### 1x_Wildtype_21.png

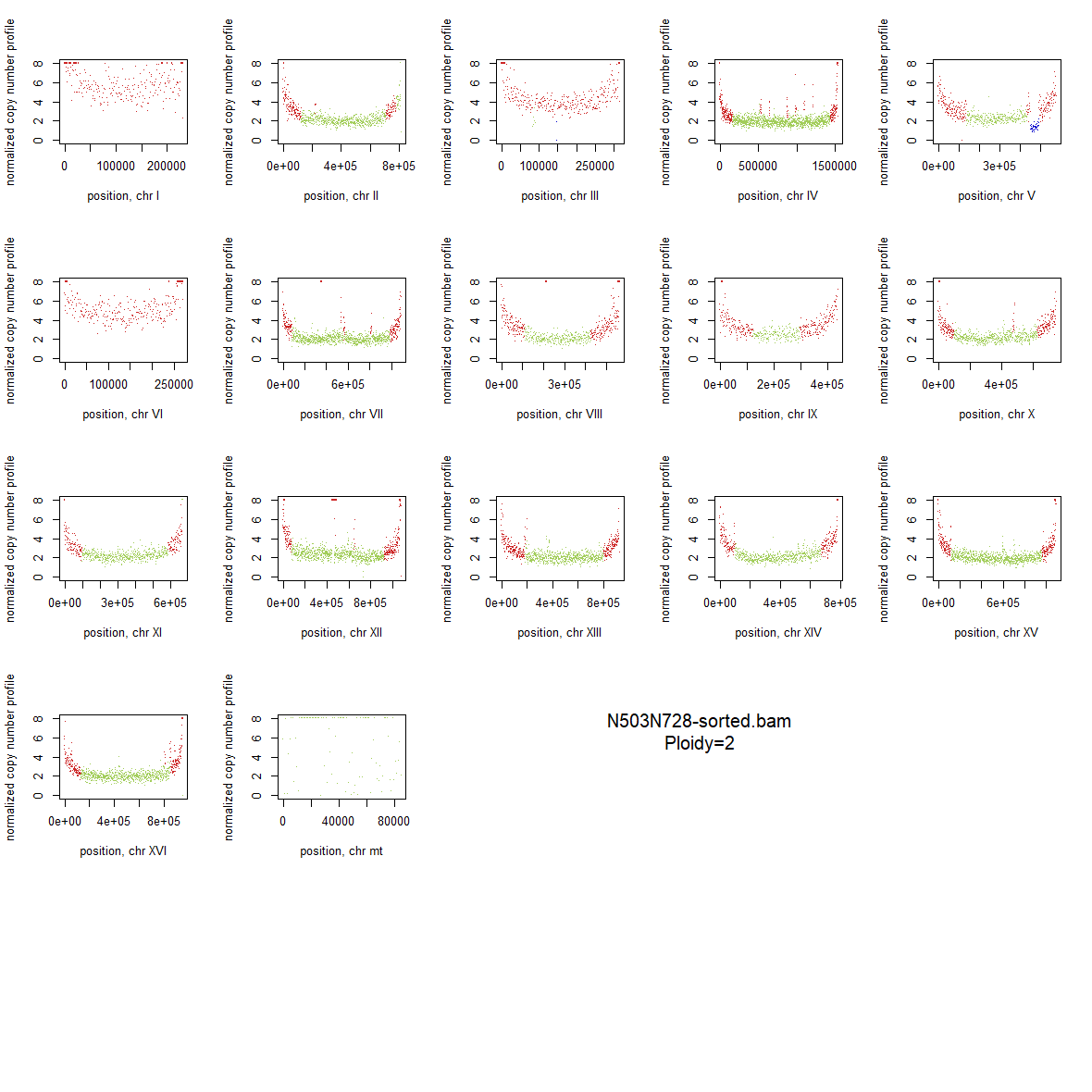

### 1x_Wildtype_22.png

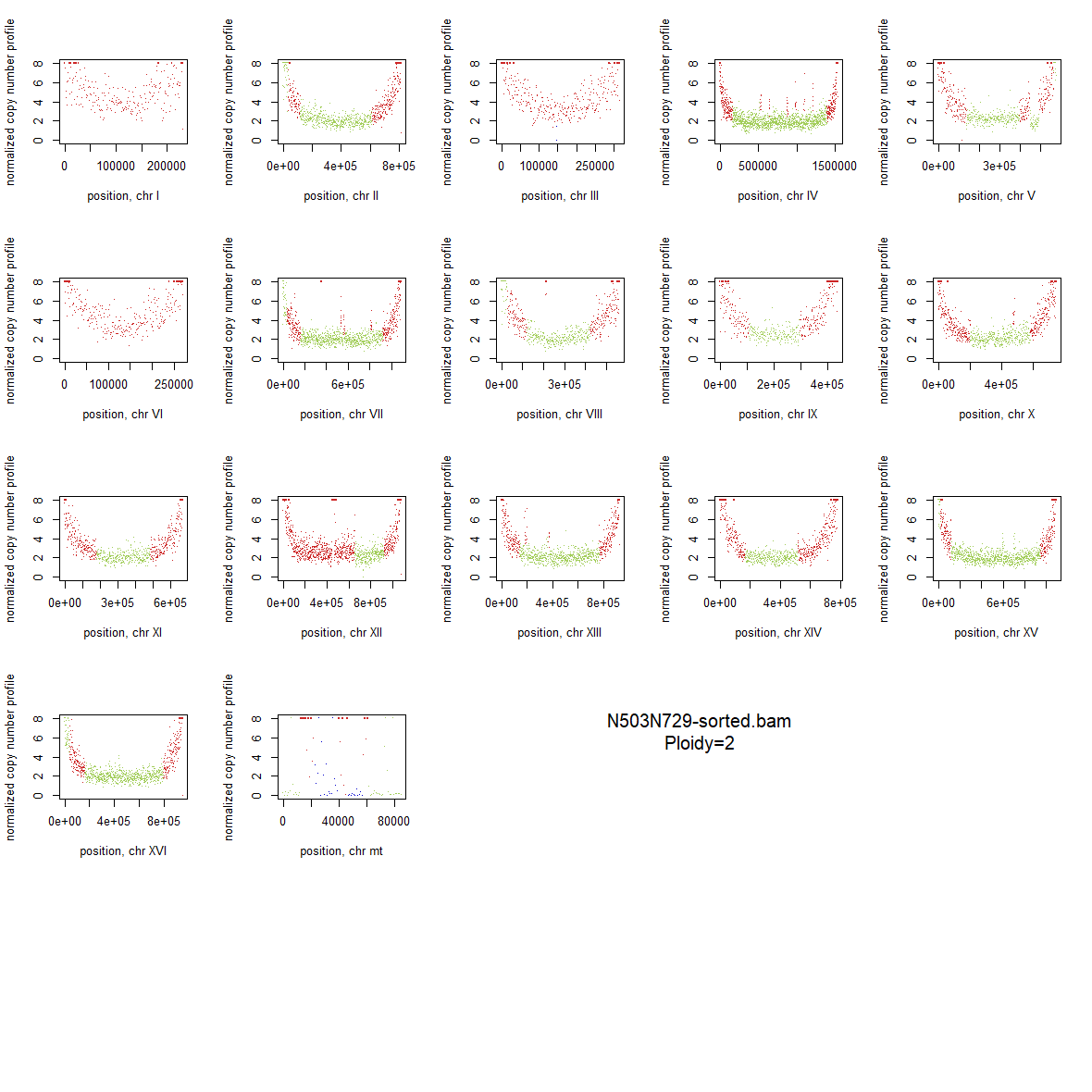

### 1x_Wildtype_23.png

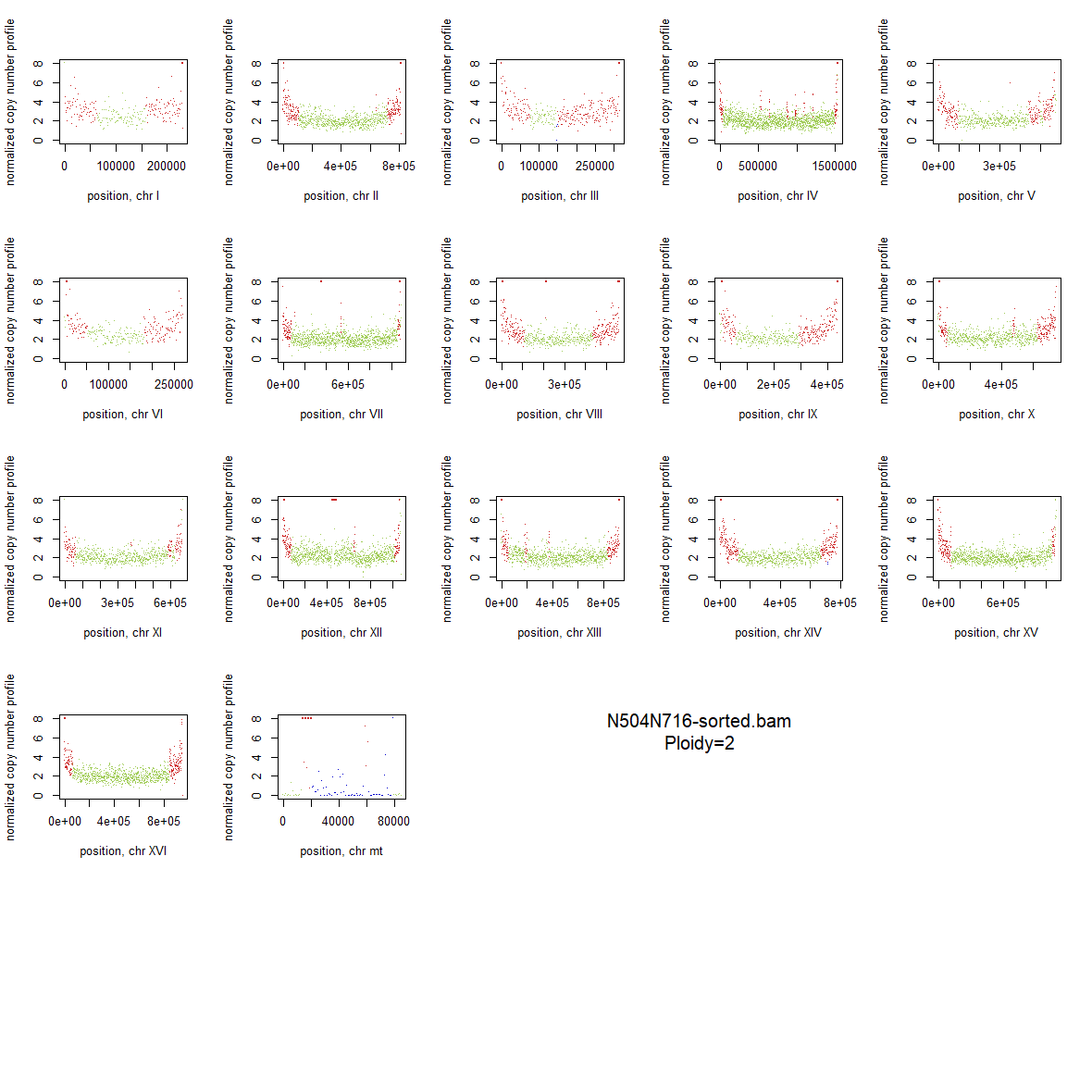

### 1x_Wildtype_24.png

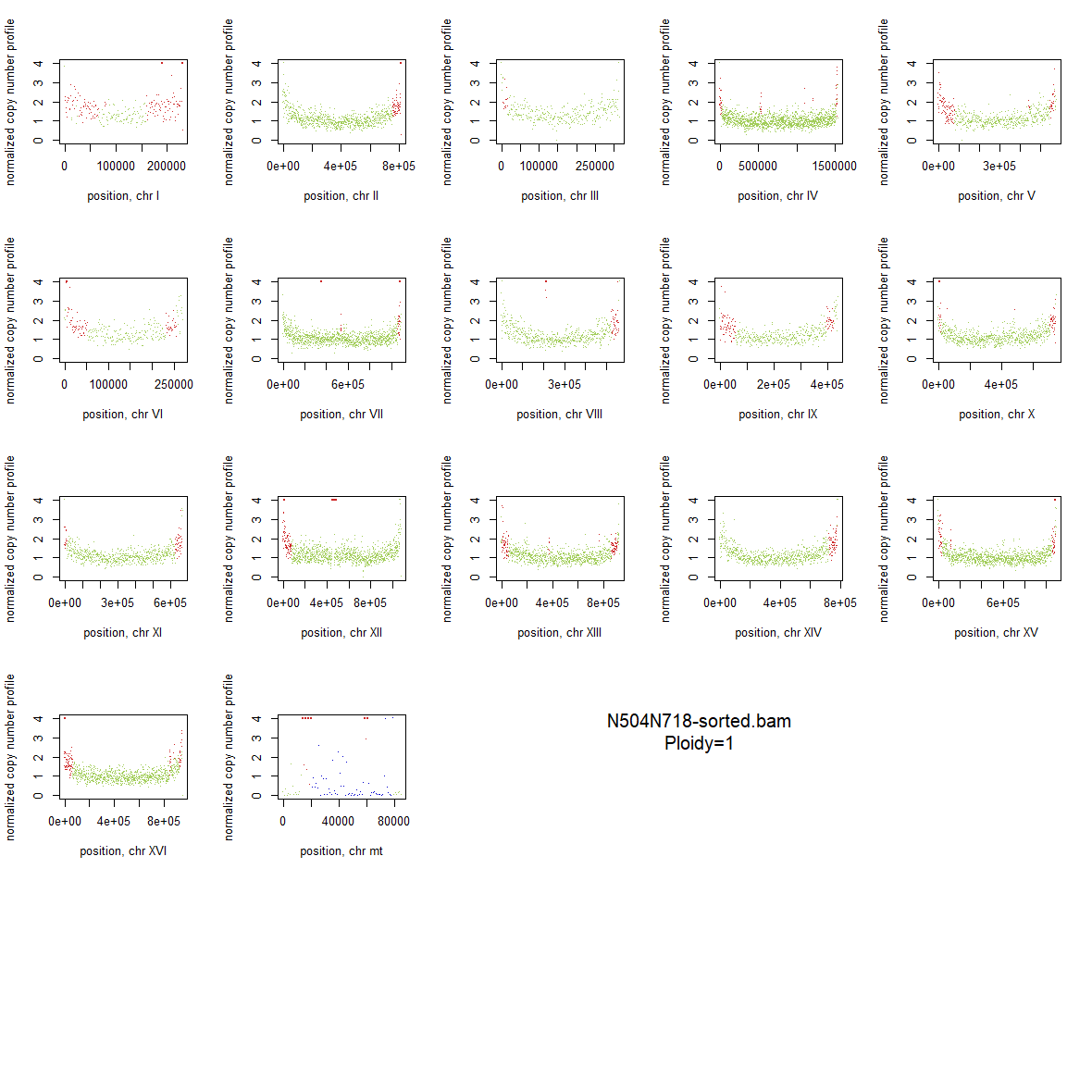

### 1x_Wildtype_25.png

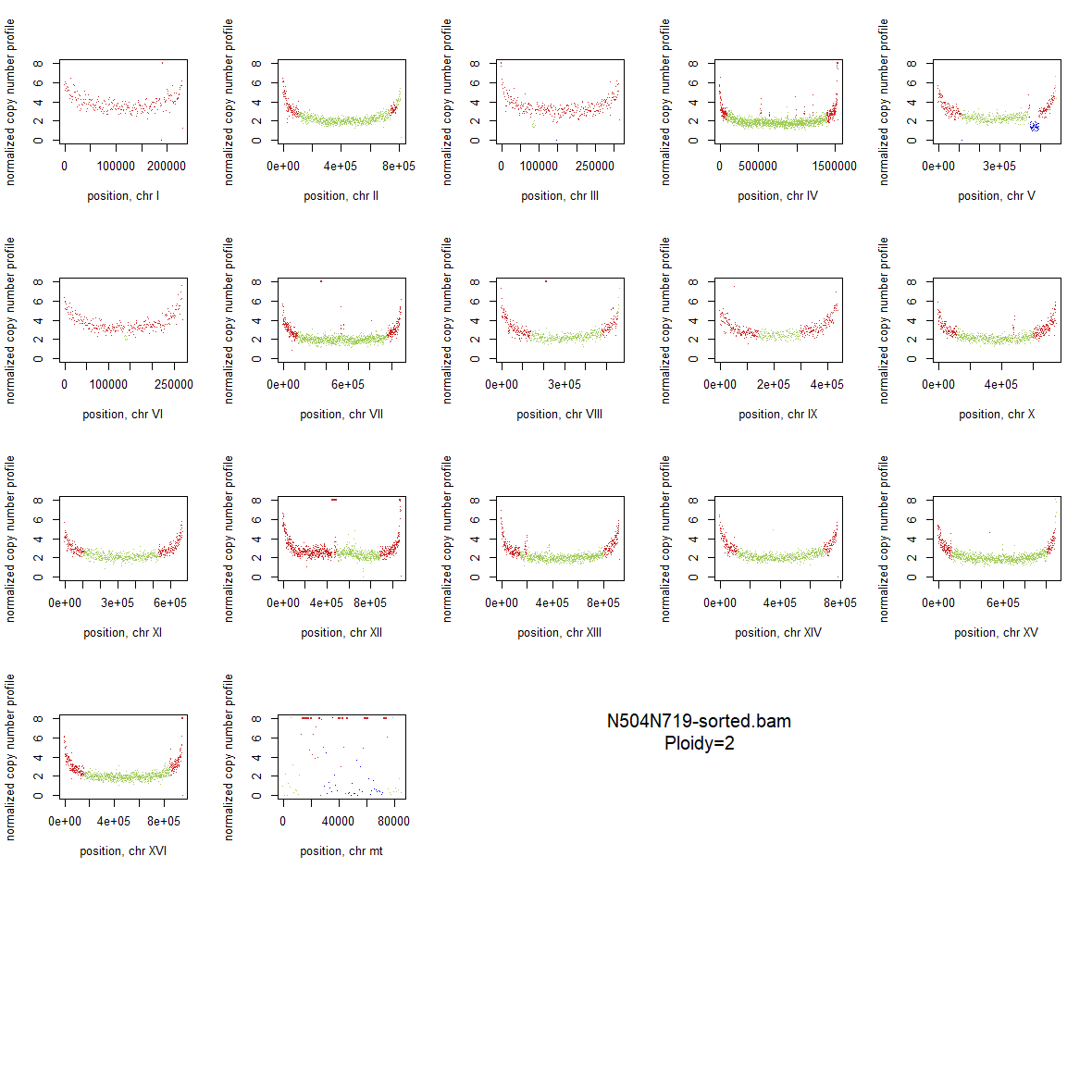

### 1x_Wildtype_26.png

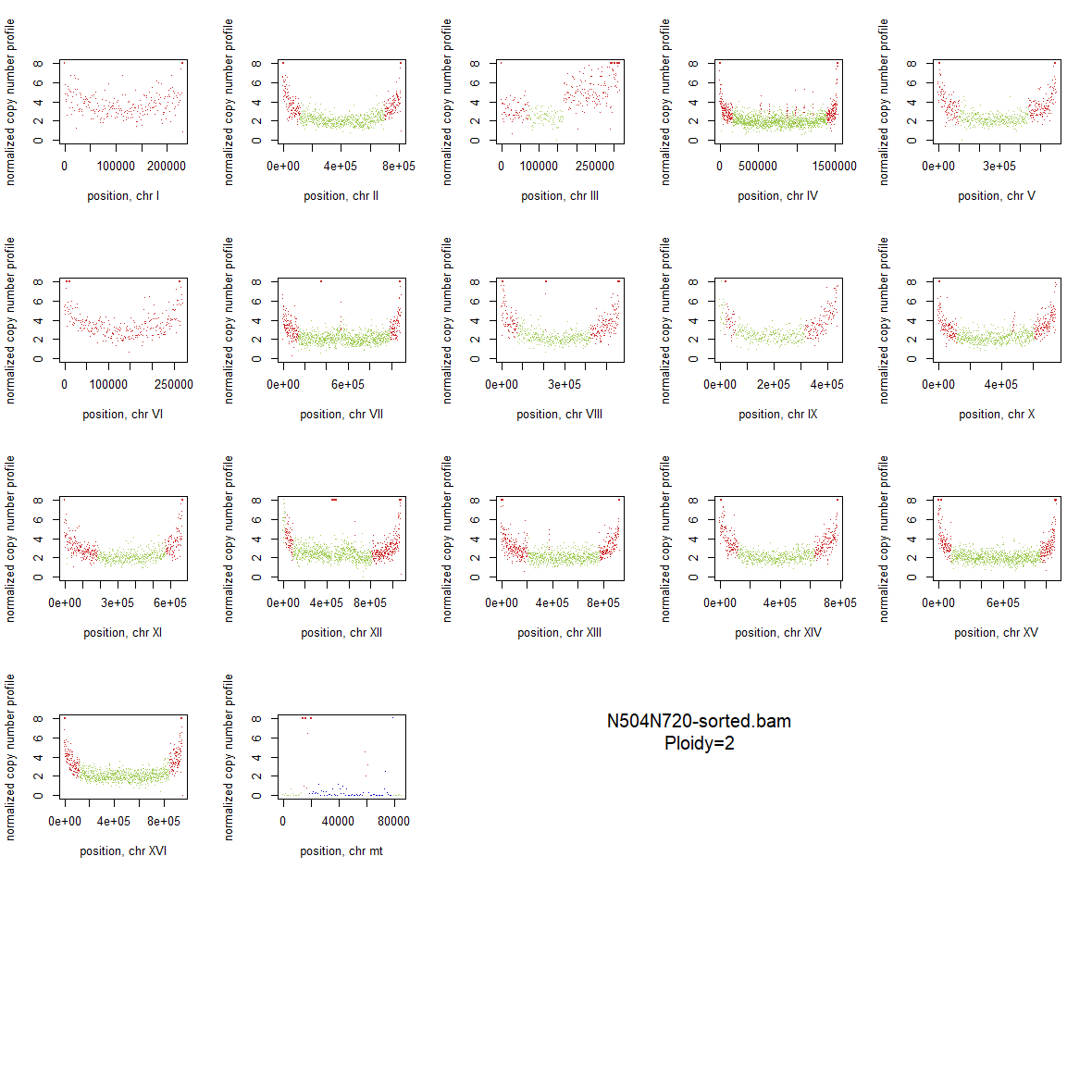

### 1x_Wildtype_27.png

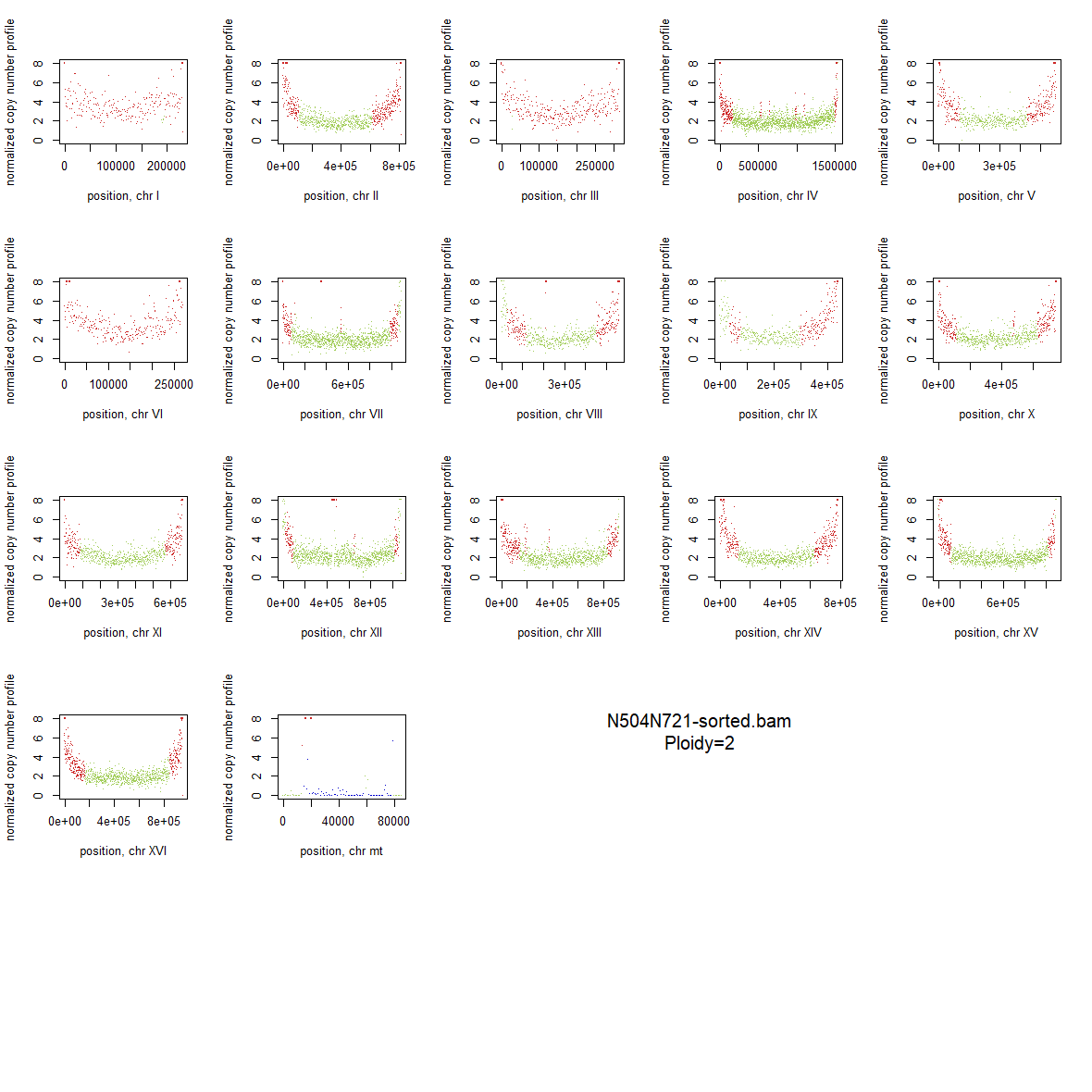

### 1x_Wildtype_28.png

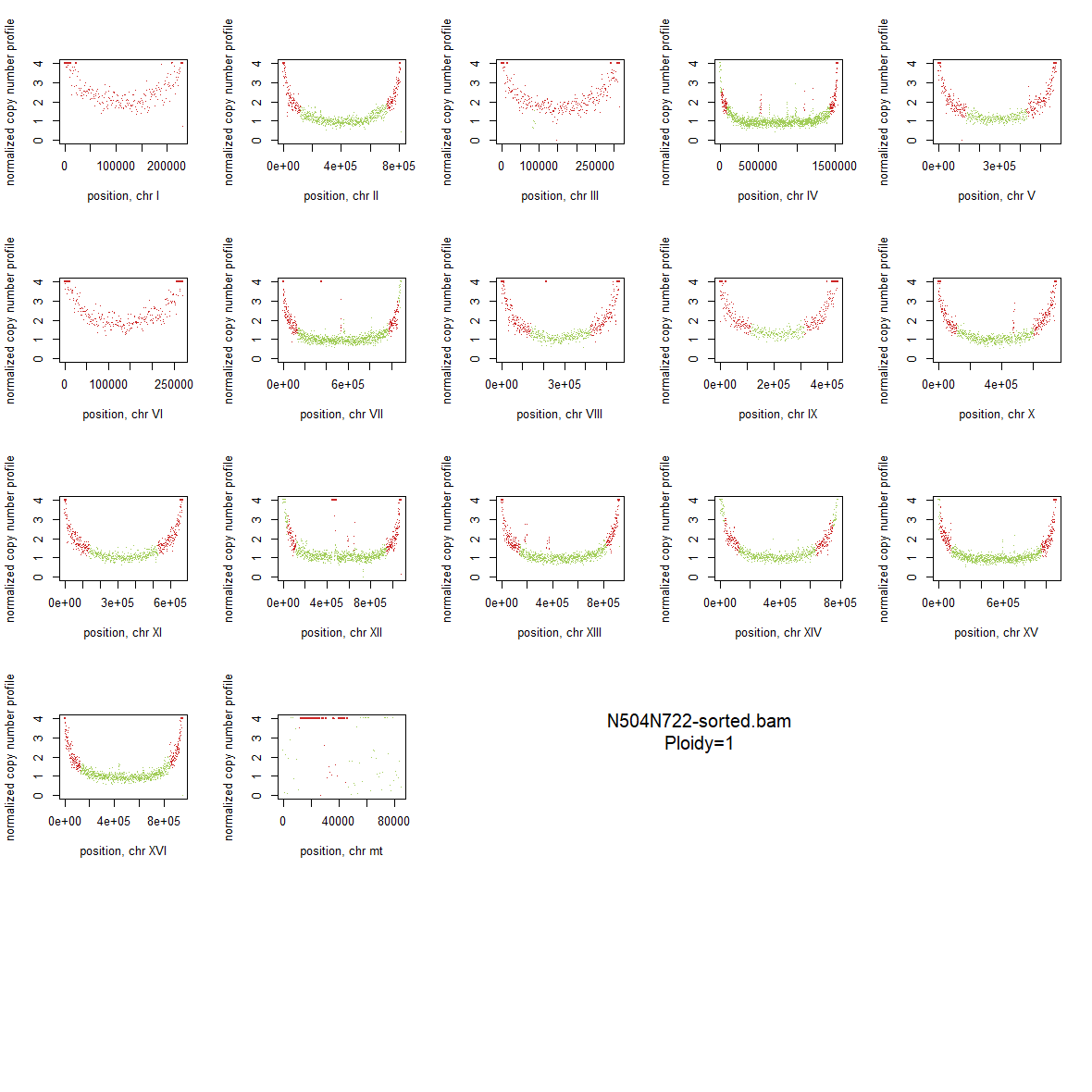

### 1x_Wildtype_29.png

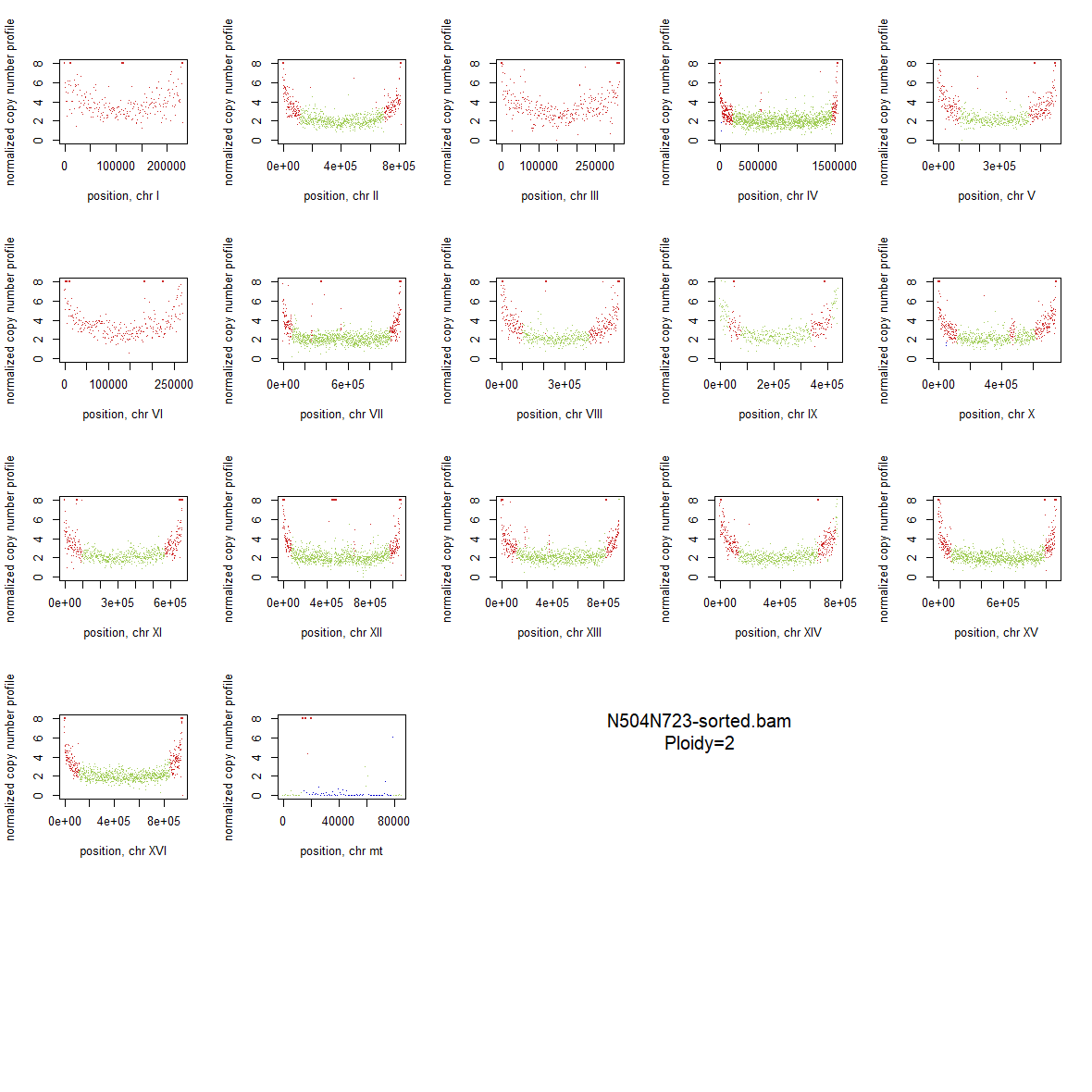

### 1x_Wildtype_30.png

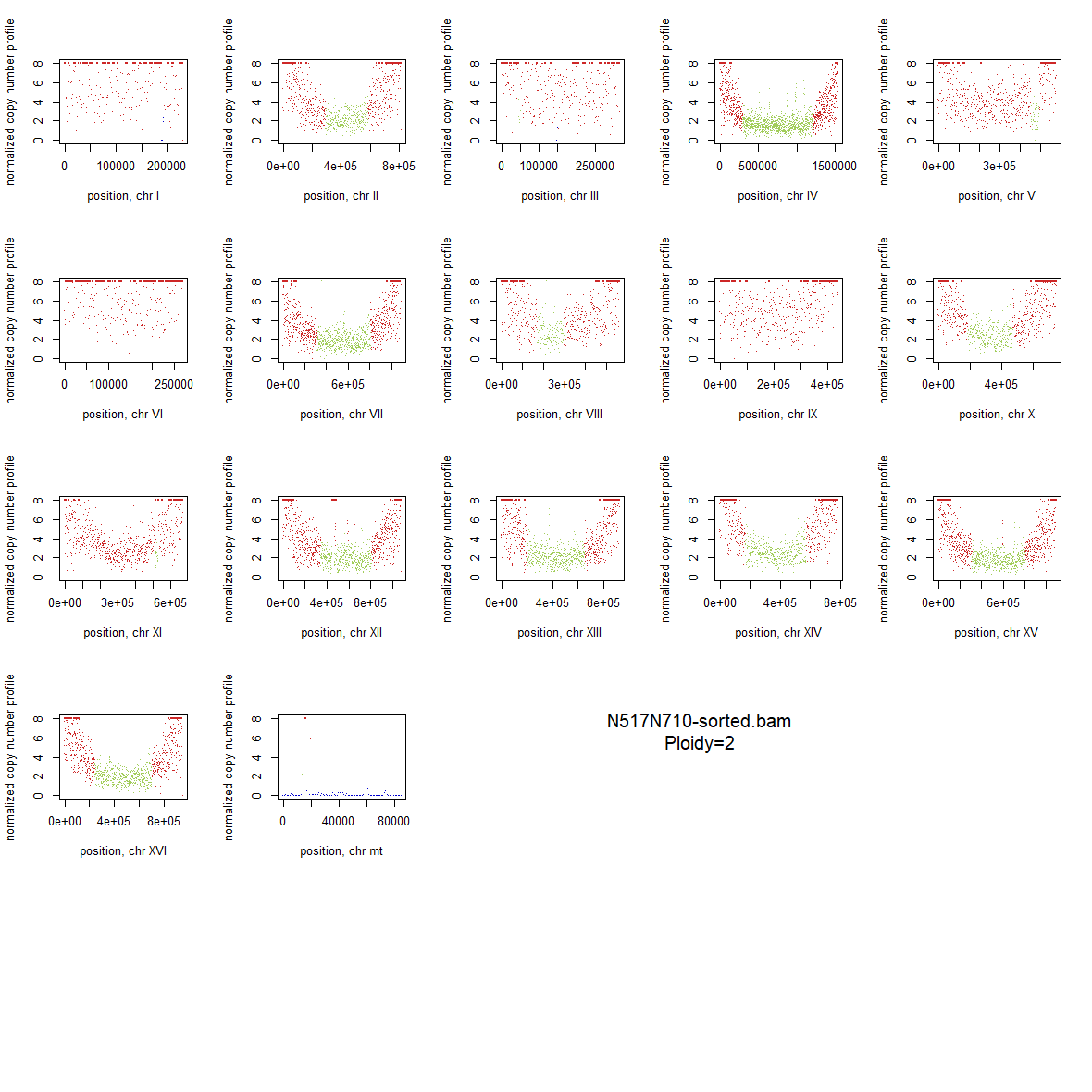

### 1x_Wildtype_31.png

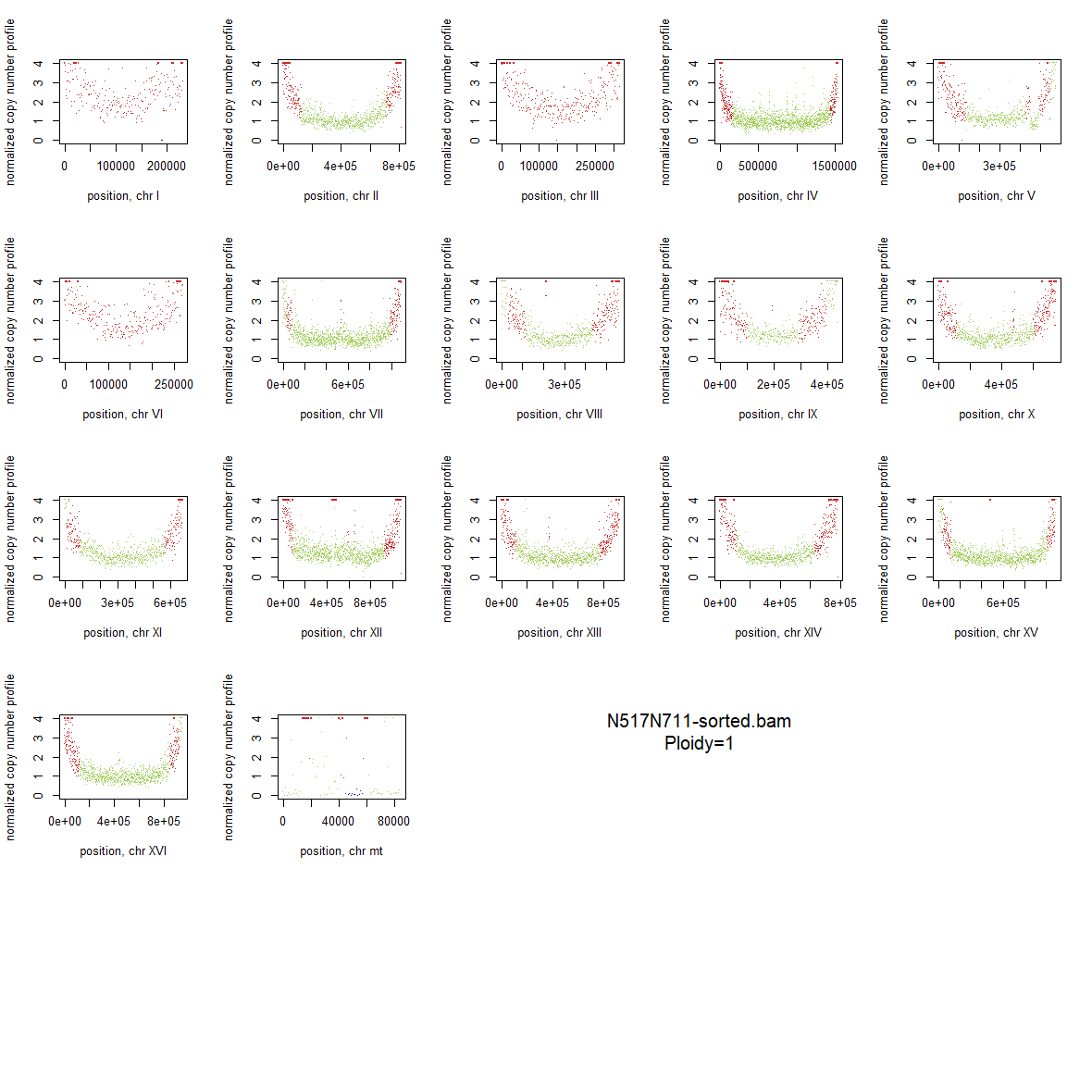

### 1x_Wildtype_32.png

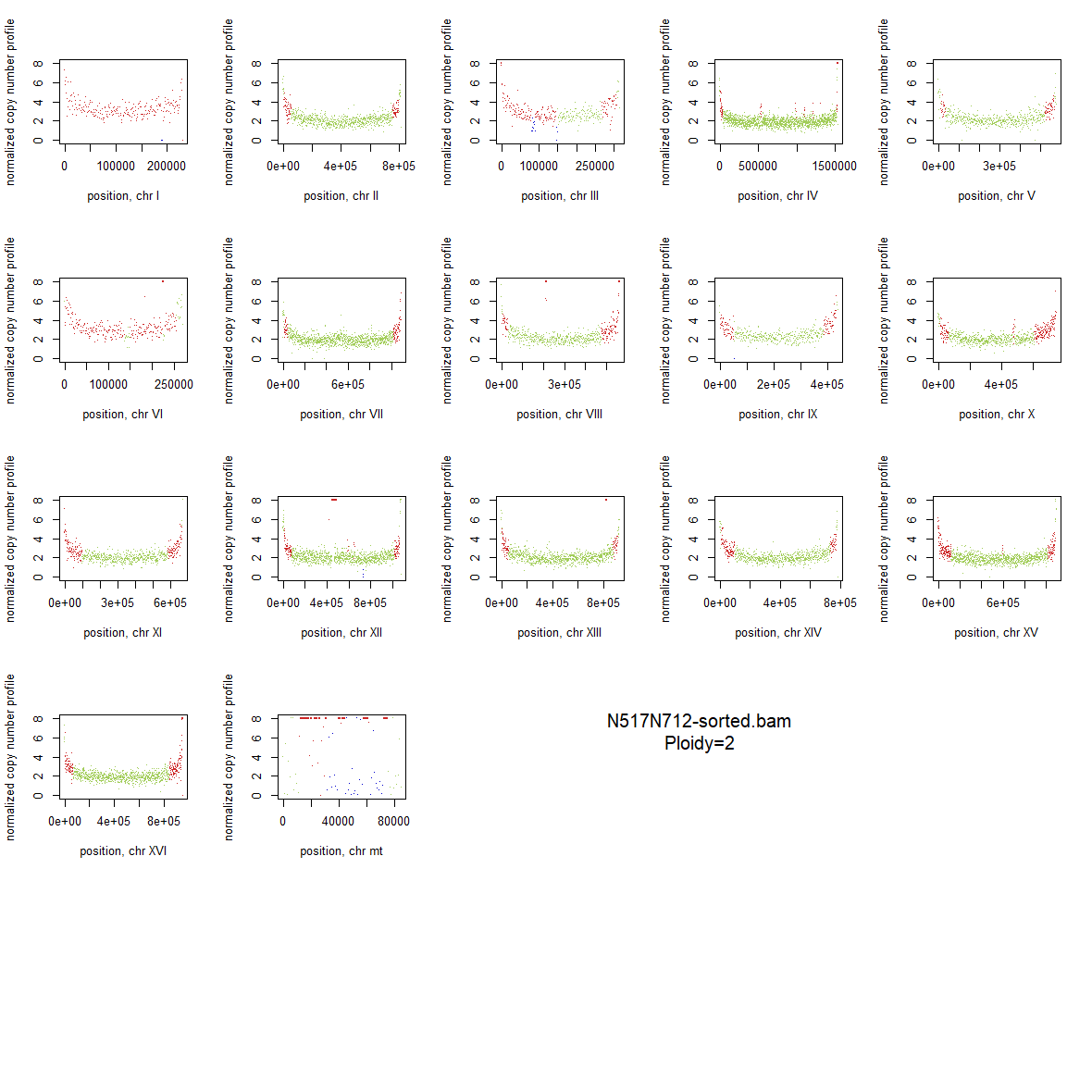

### 2x_F126L_01.png

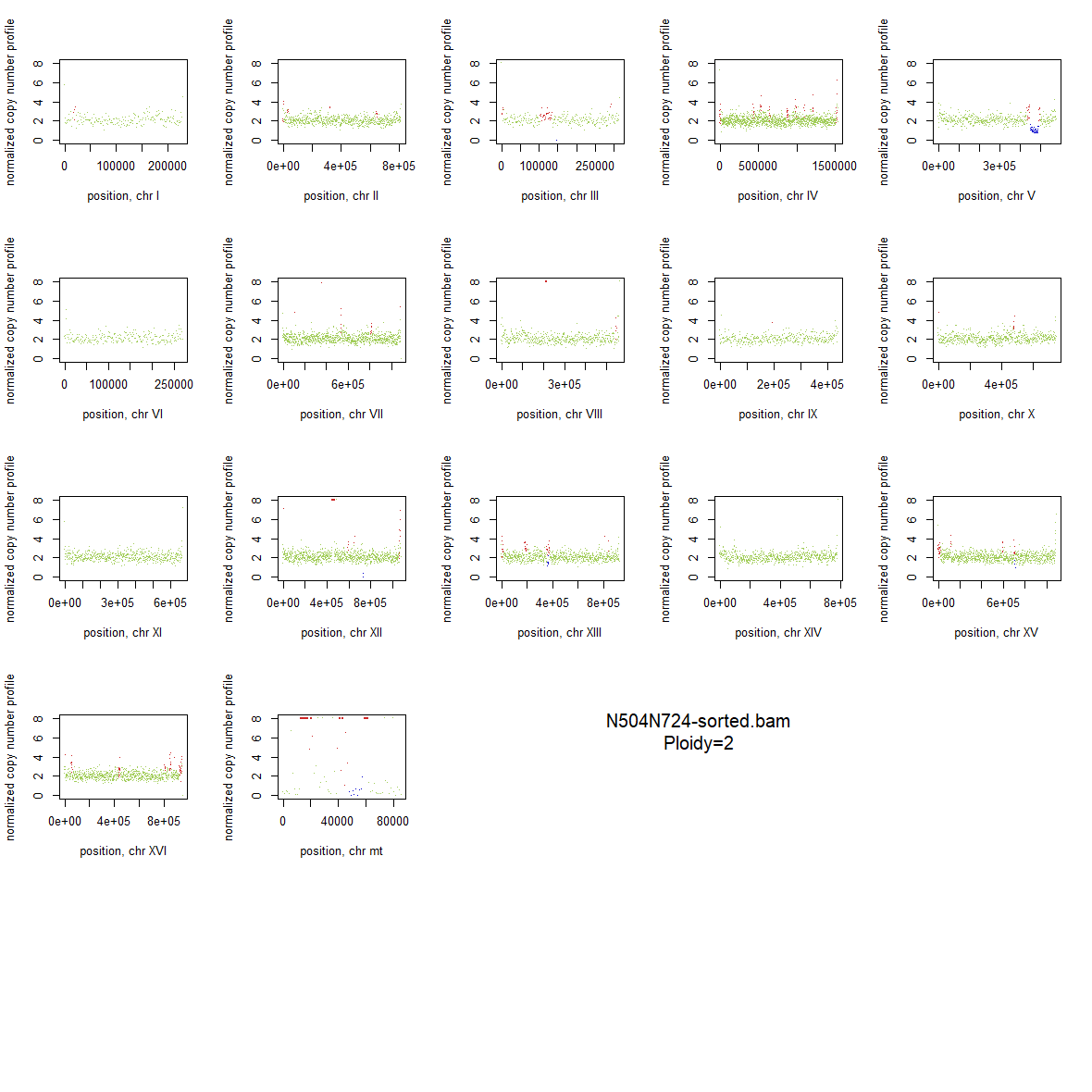

### 2x_F126L_02.png

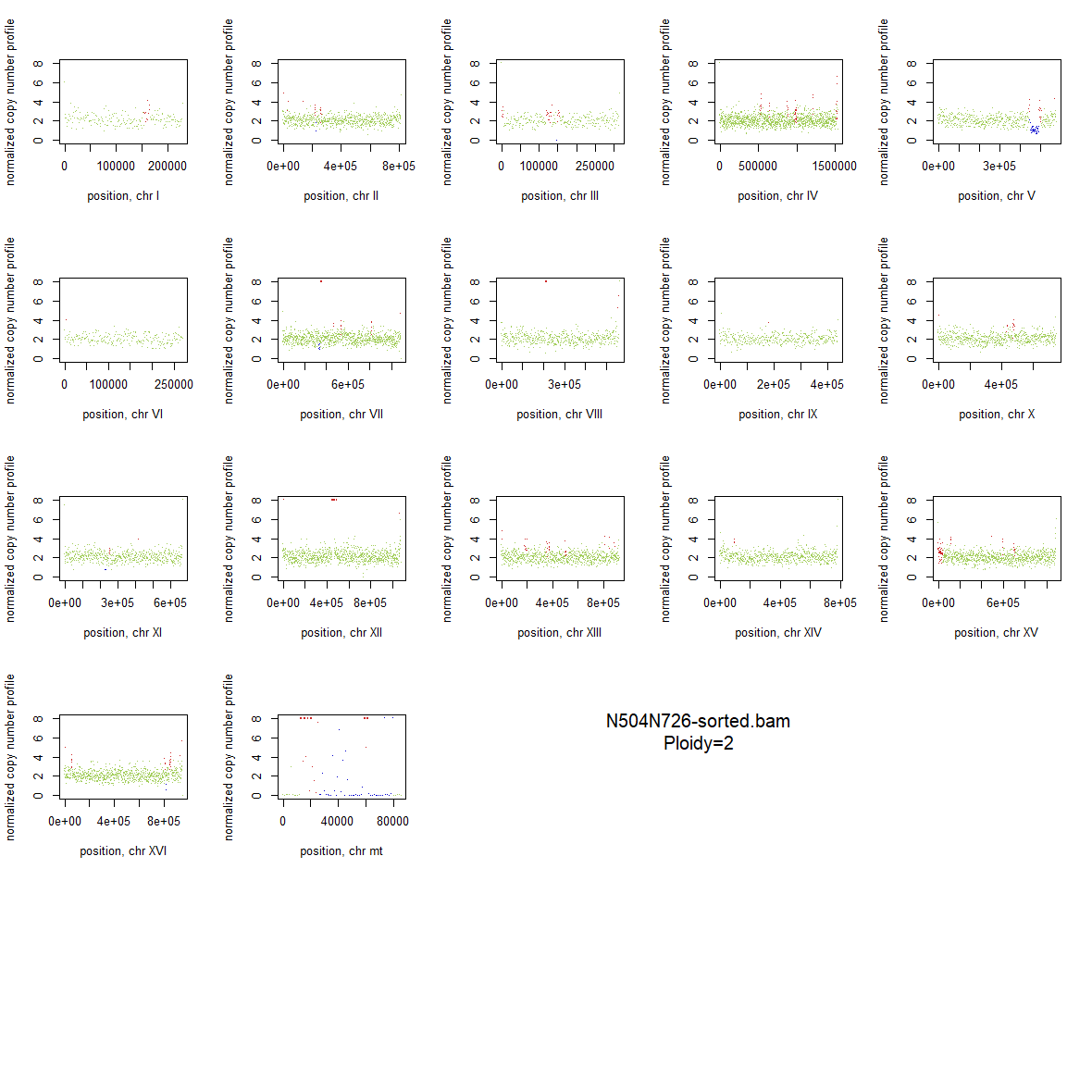

### 2x_F126L_03.png

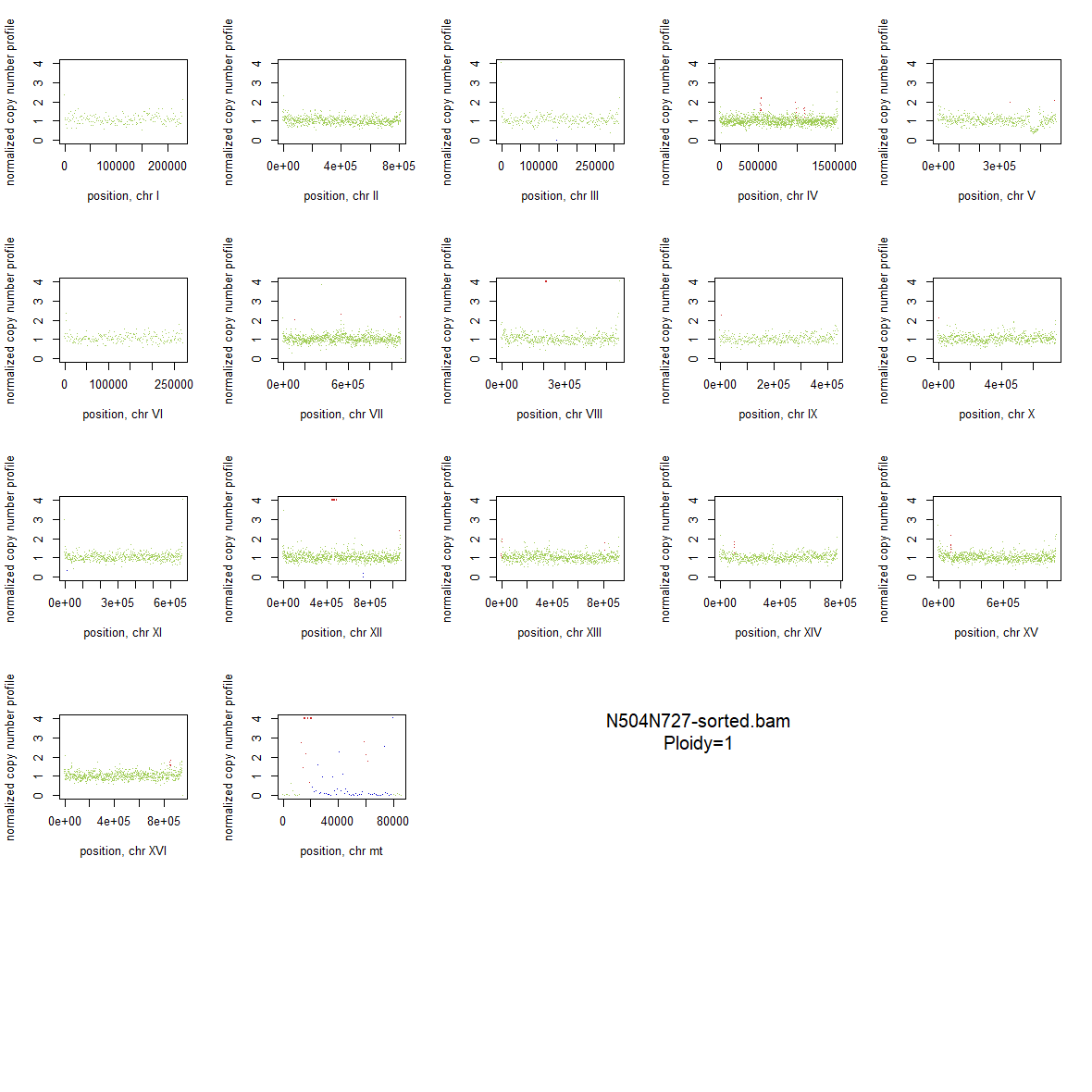

### 2x_F126L_04.png

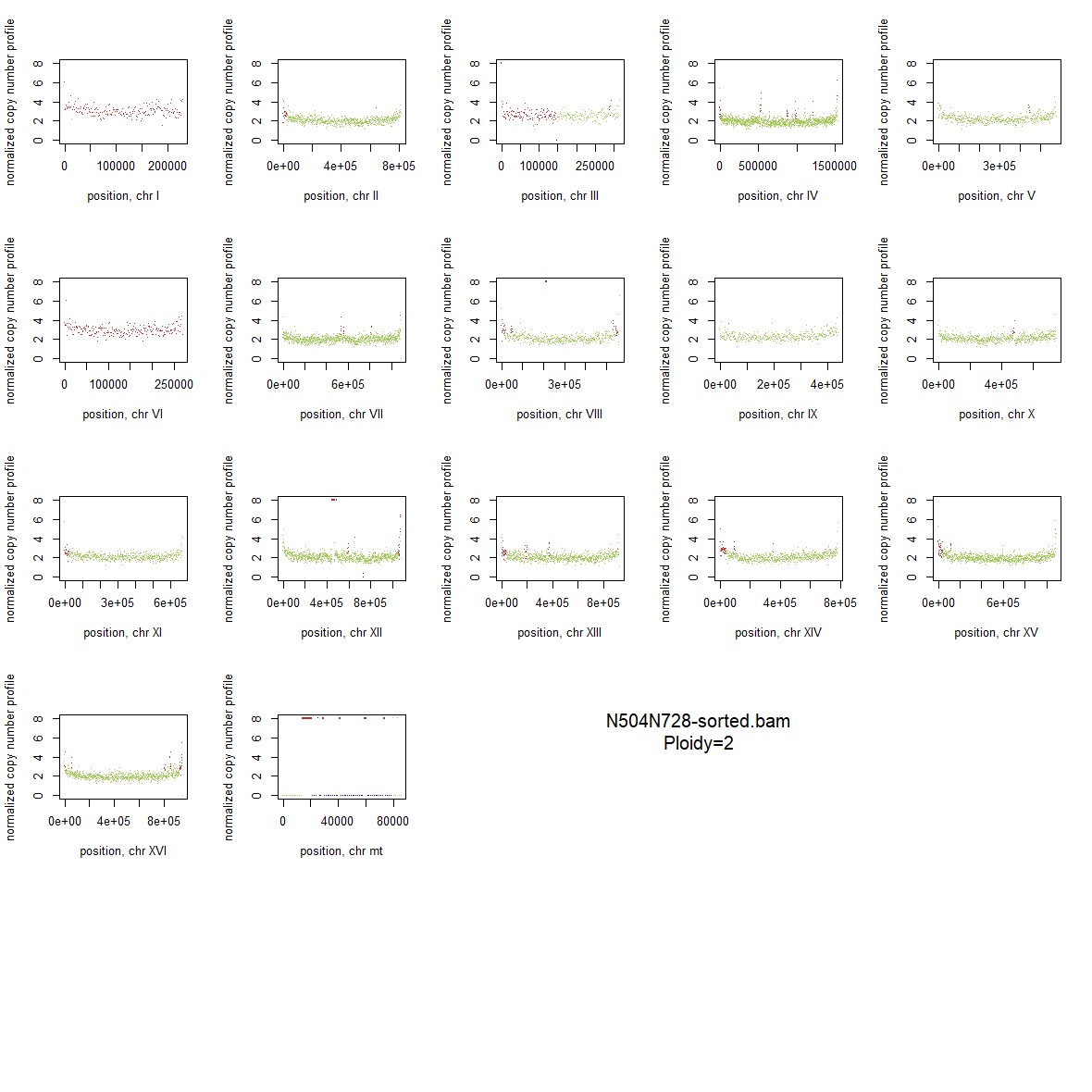

### 2x_F126L_05.png

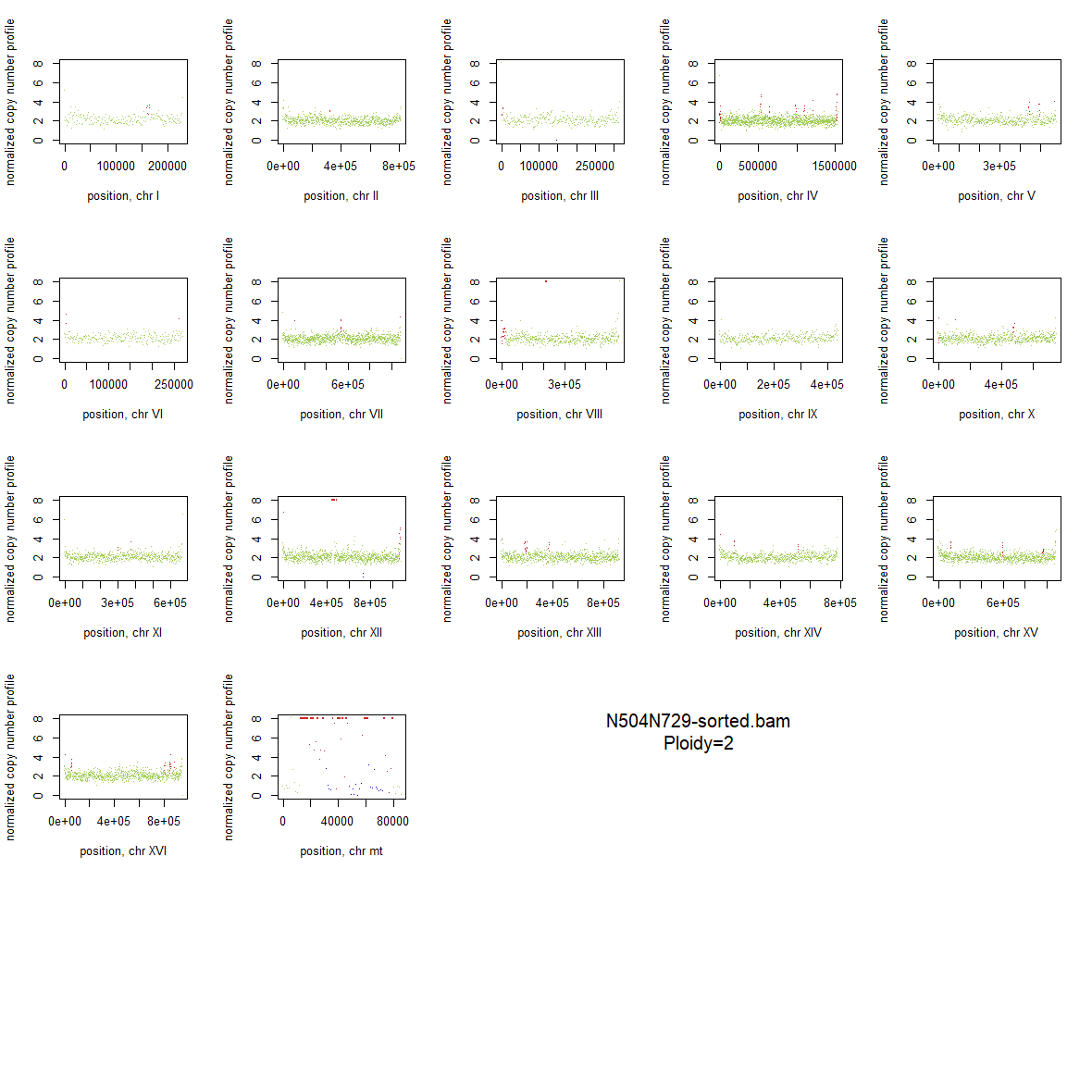

### 2x_F126L_06.png

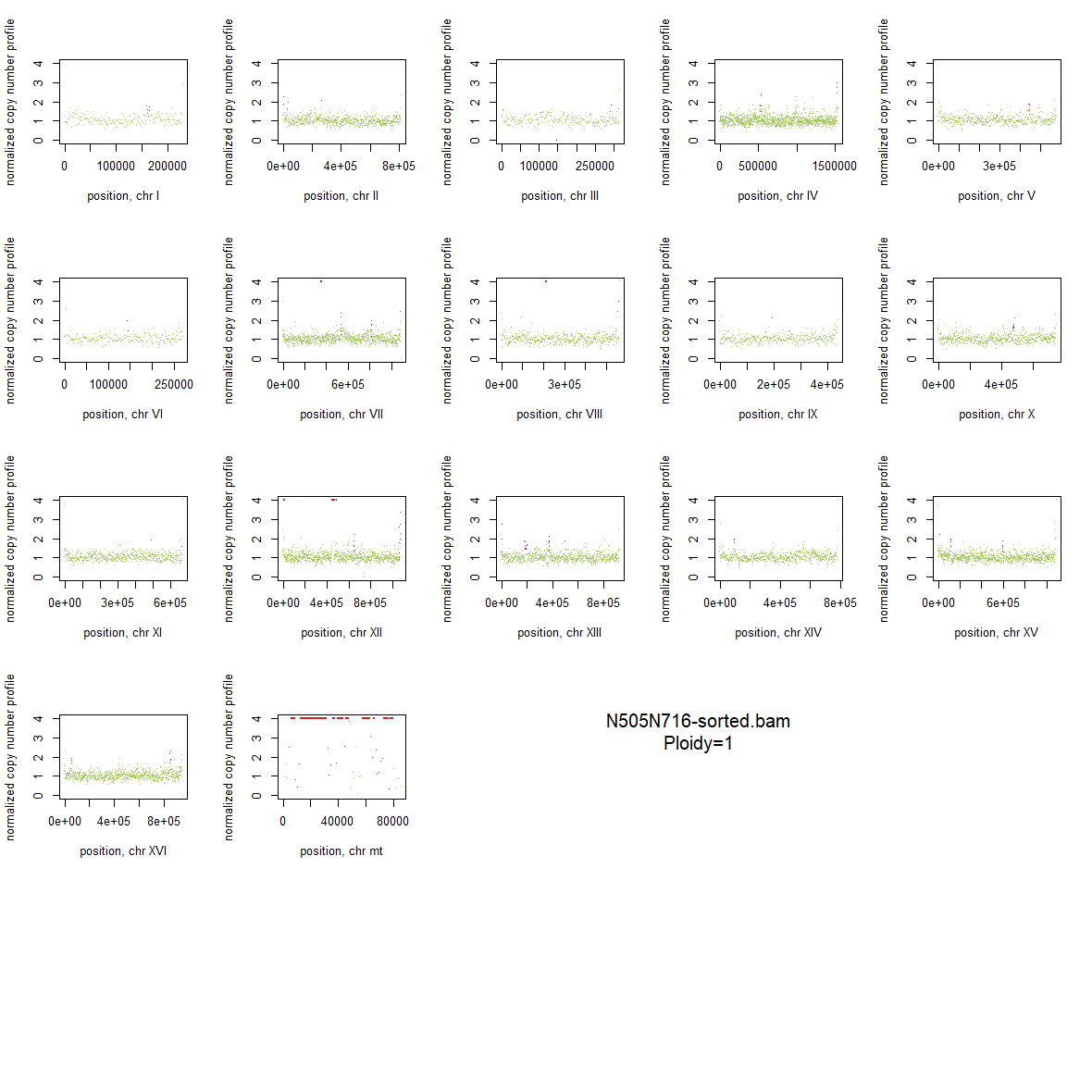

### 2x_F126L_07.png

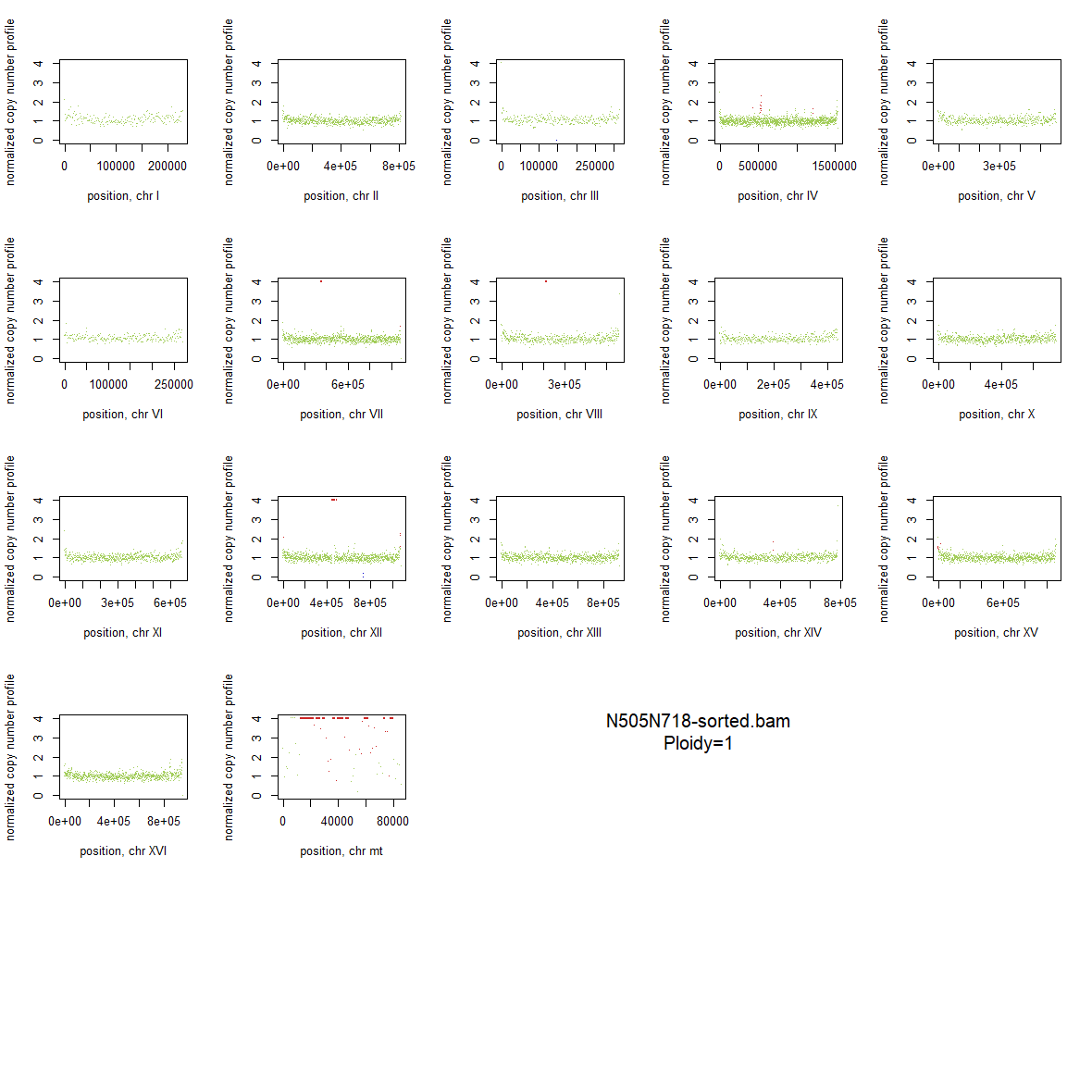

### 2x_F126L_08.png

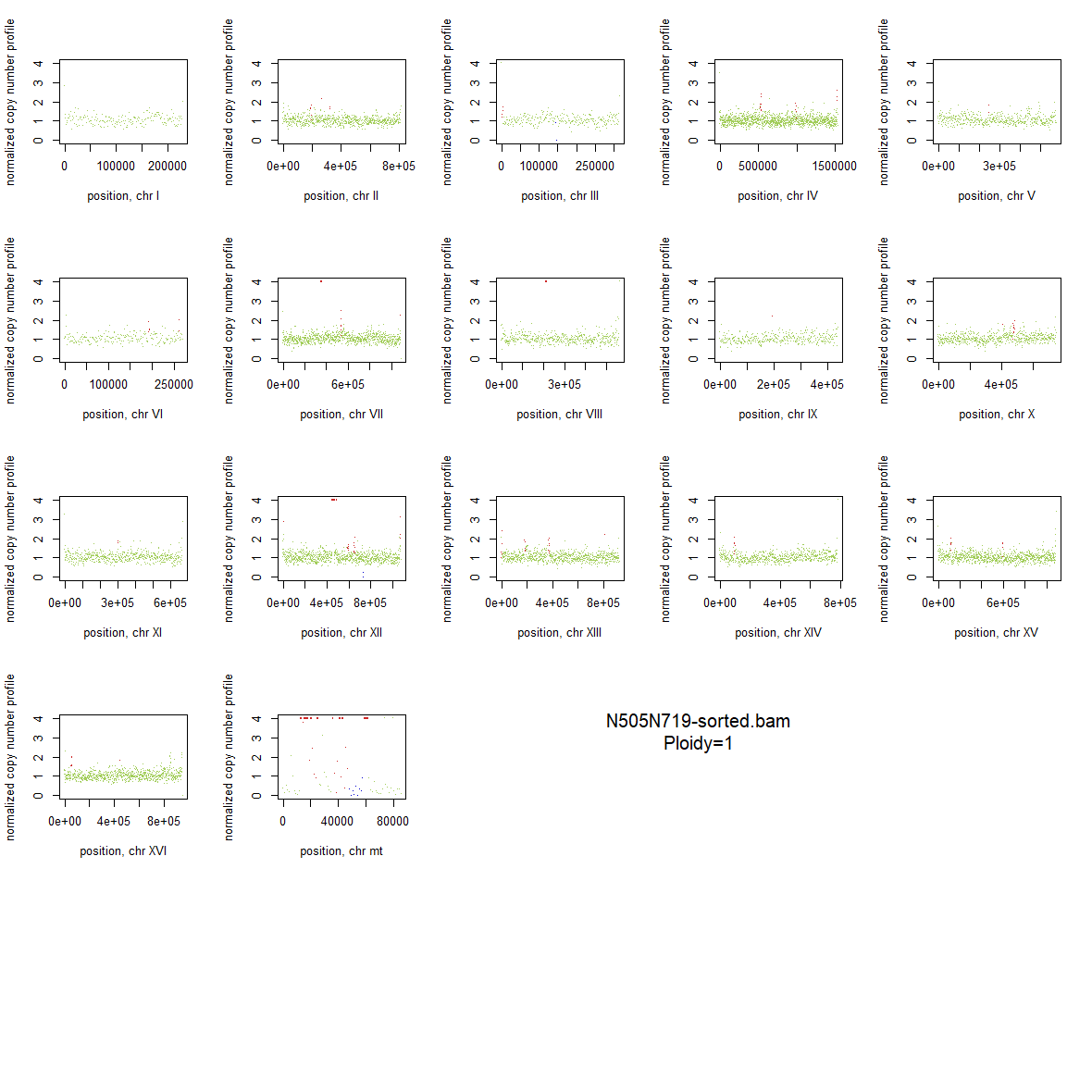

### 2x_F126L_09.png

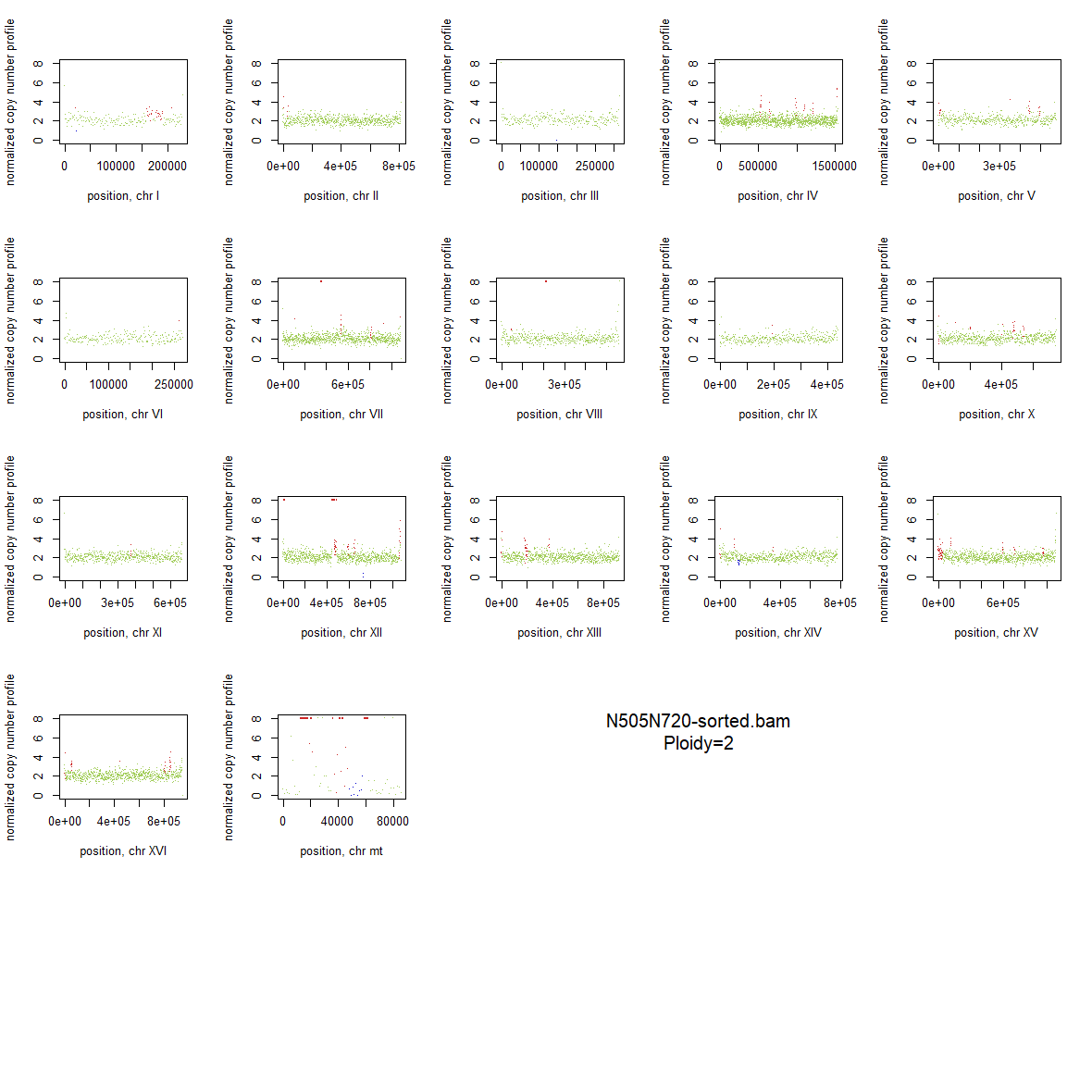

### 2x_F126L_10.png

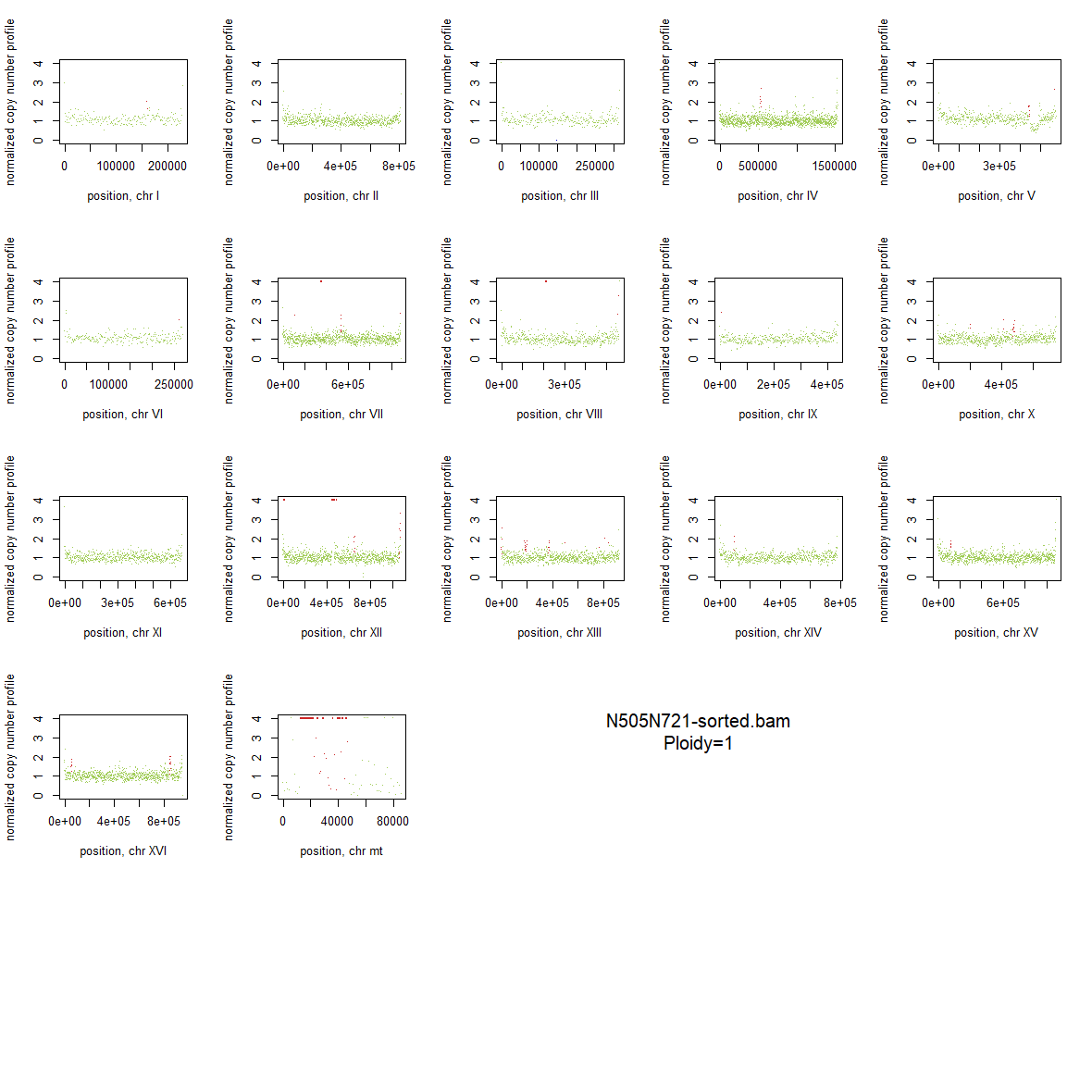

### 2x_F126L_11.png

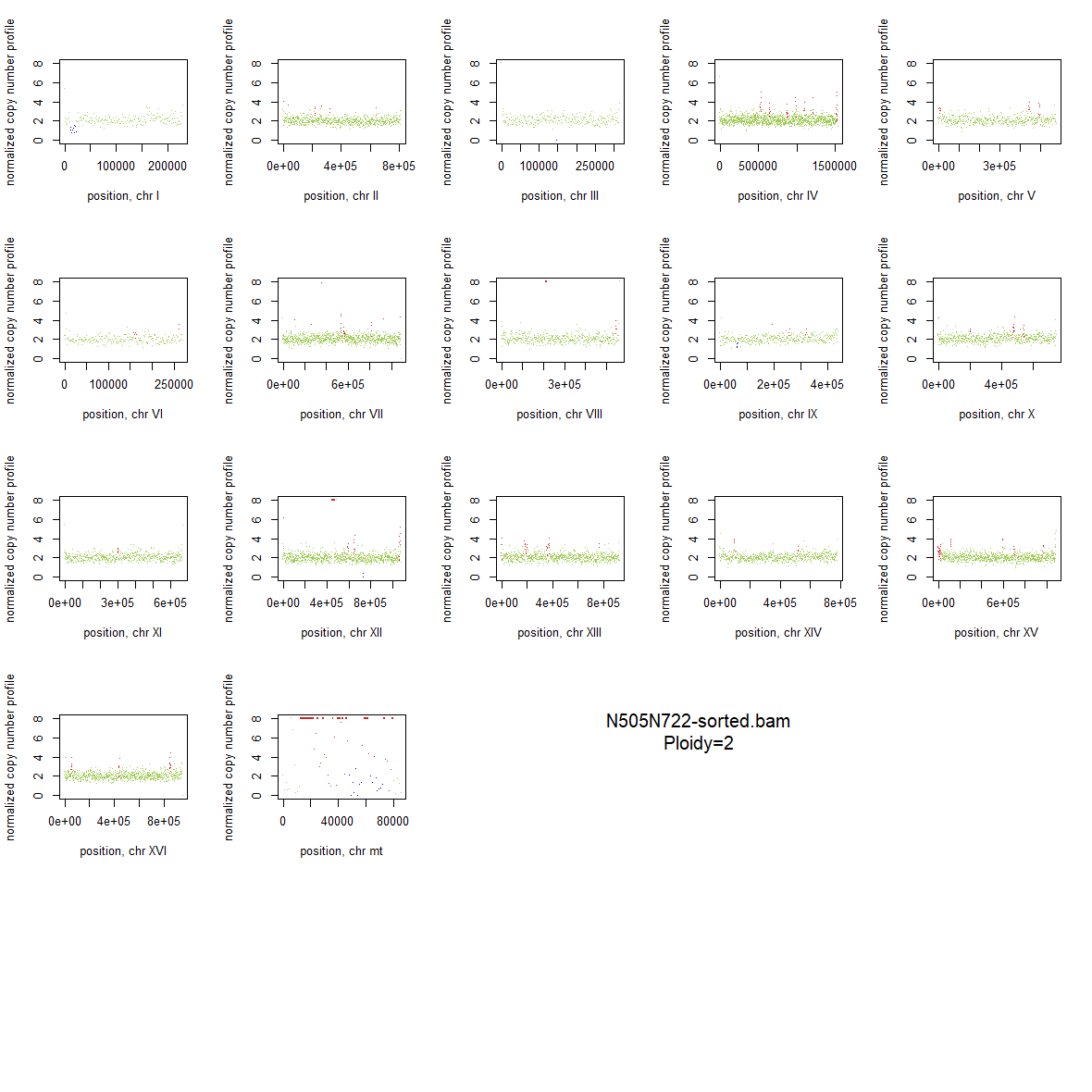

### 2x_F126L_12.png

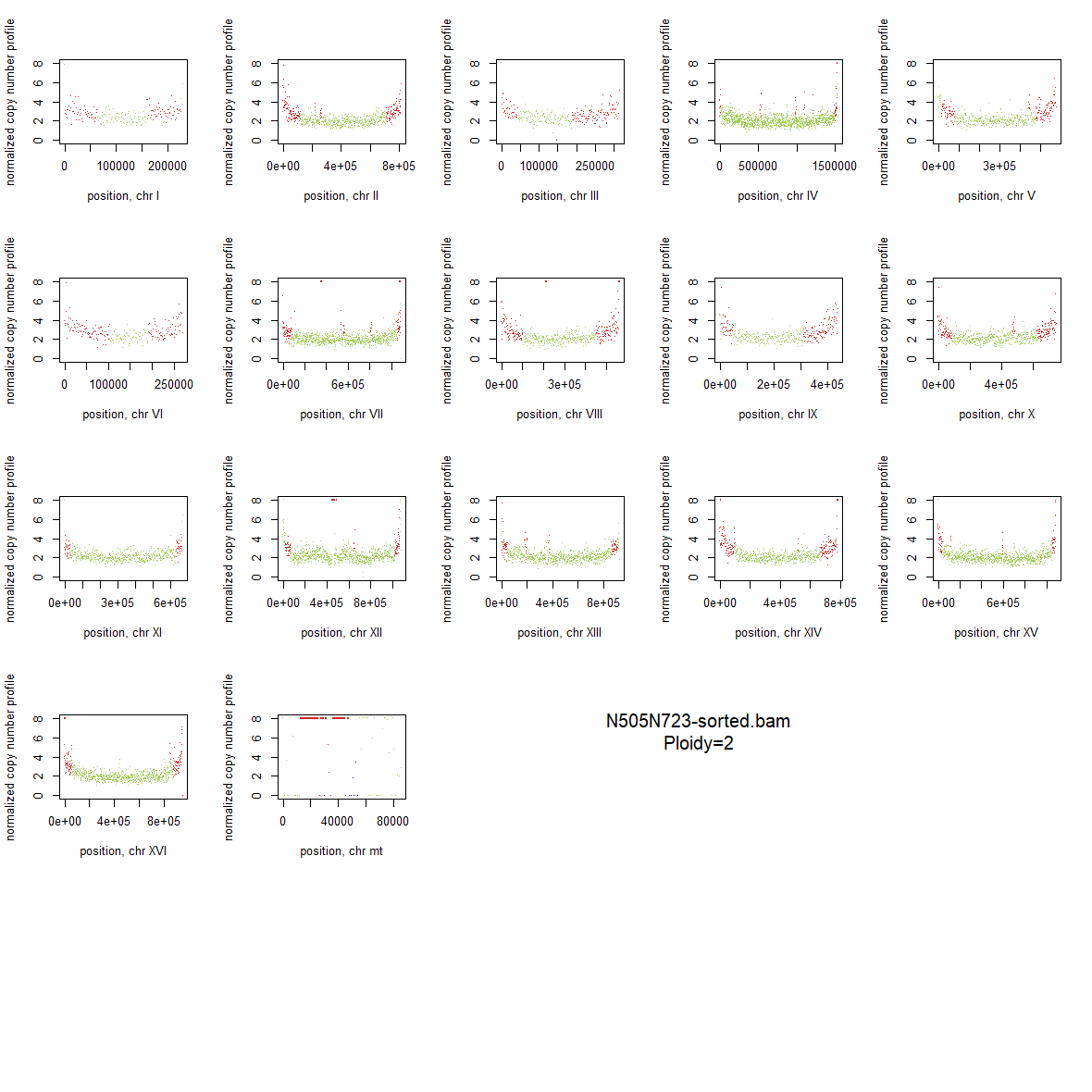

### 2x_F126L_13.png

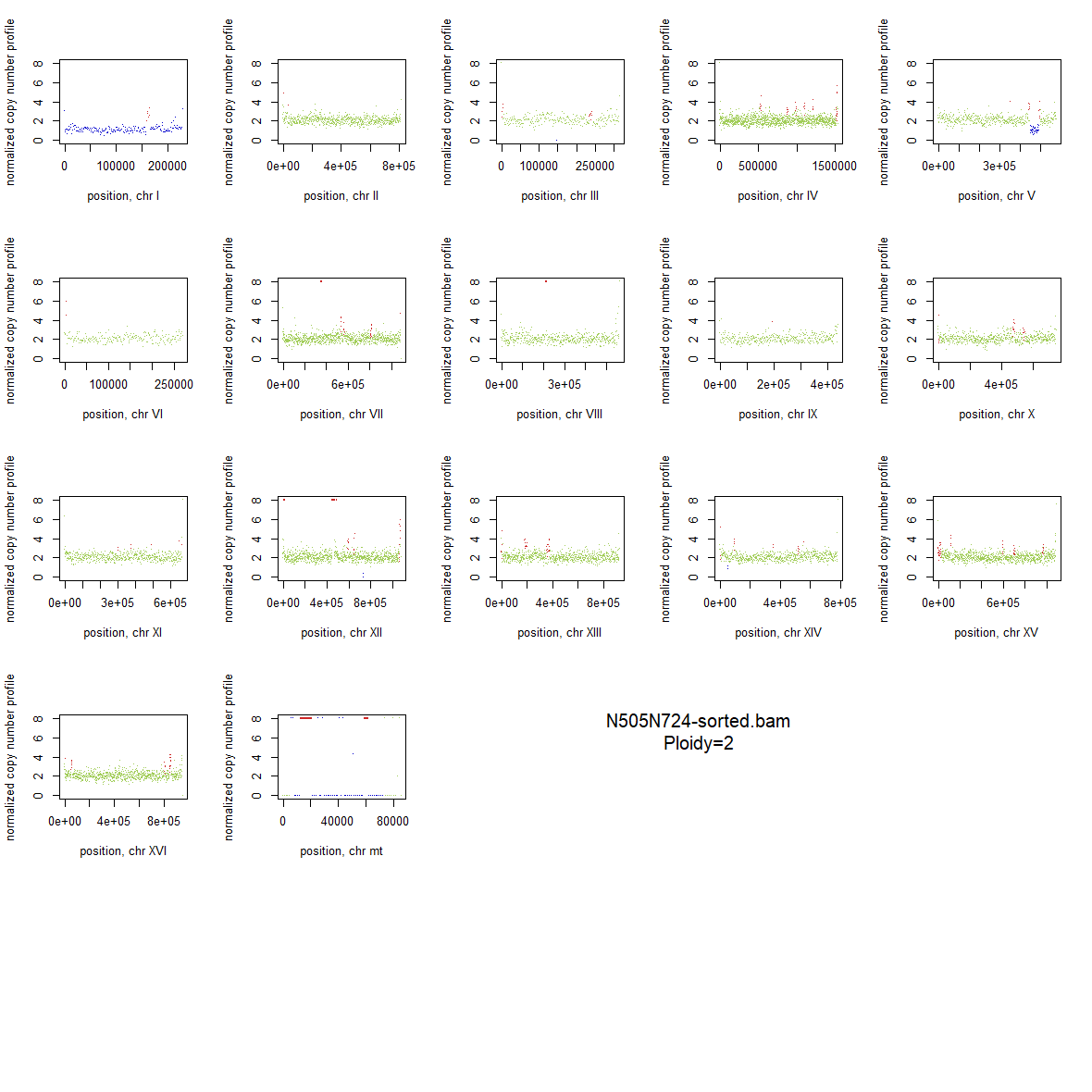

### 2x_F126L_14.png

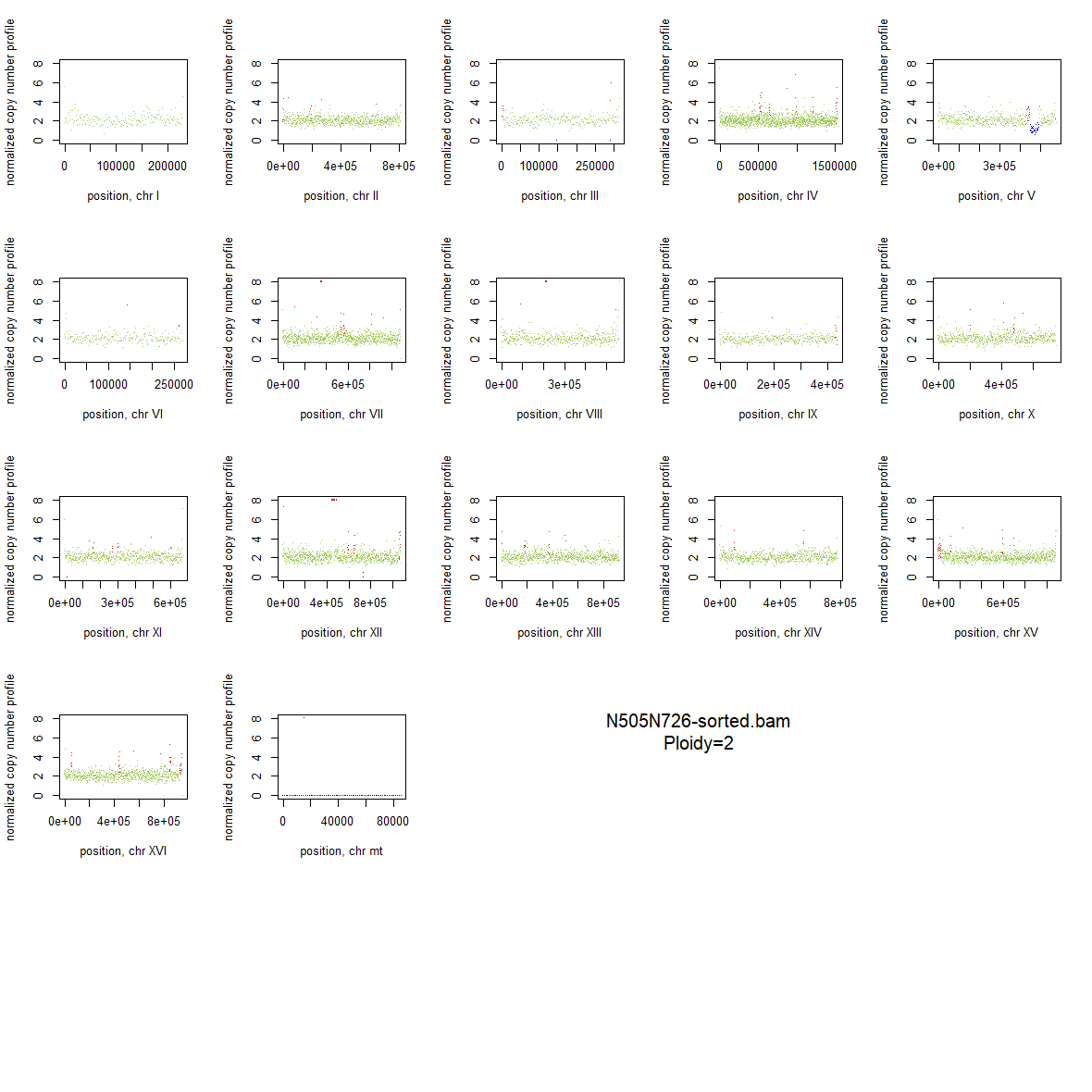

### 2x_F126L_15.png

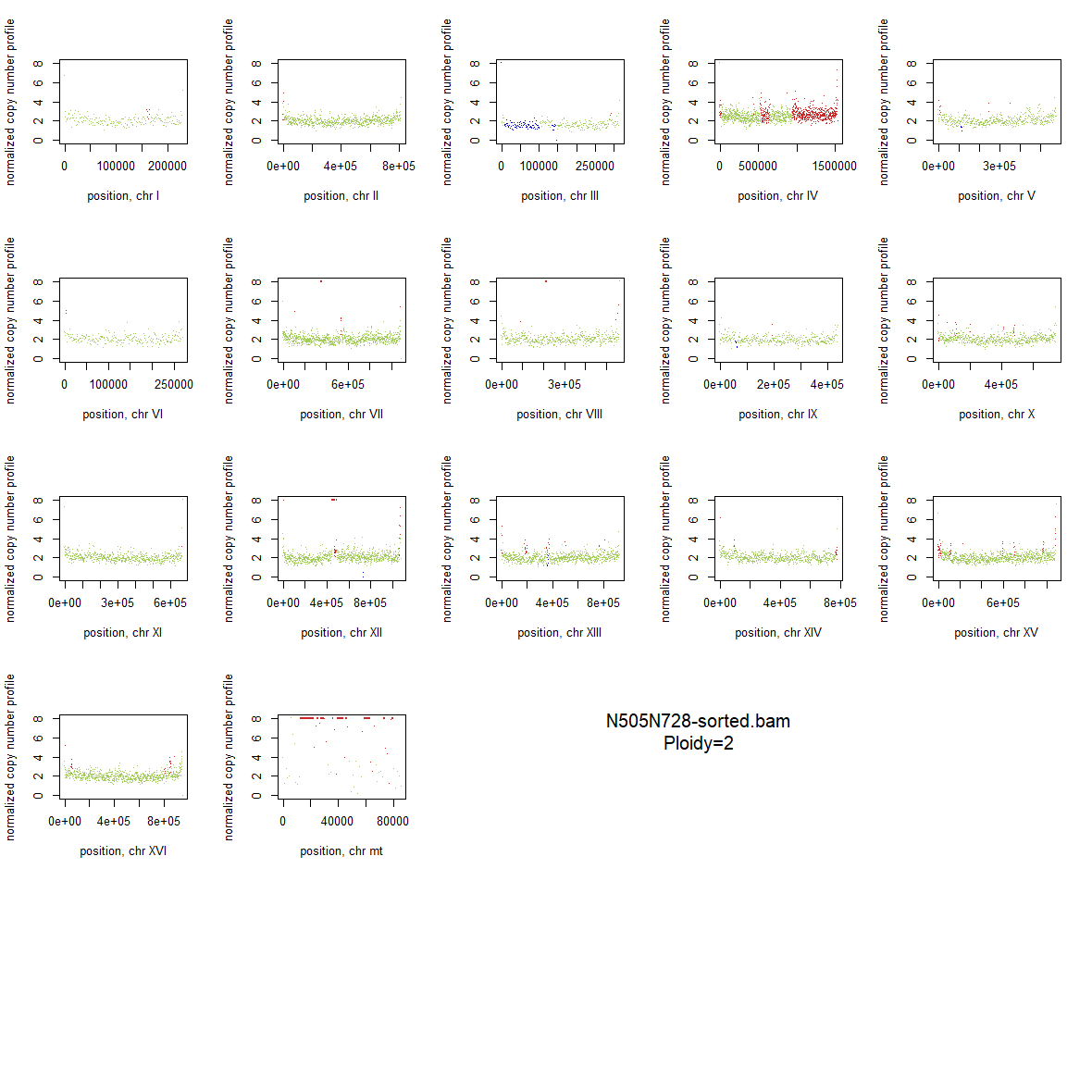

### 2x_F126L_16.png

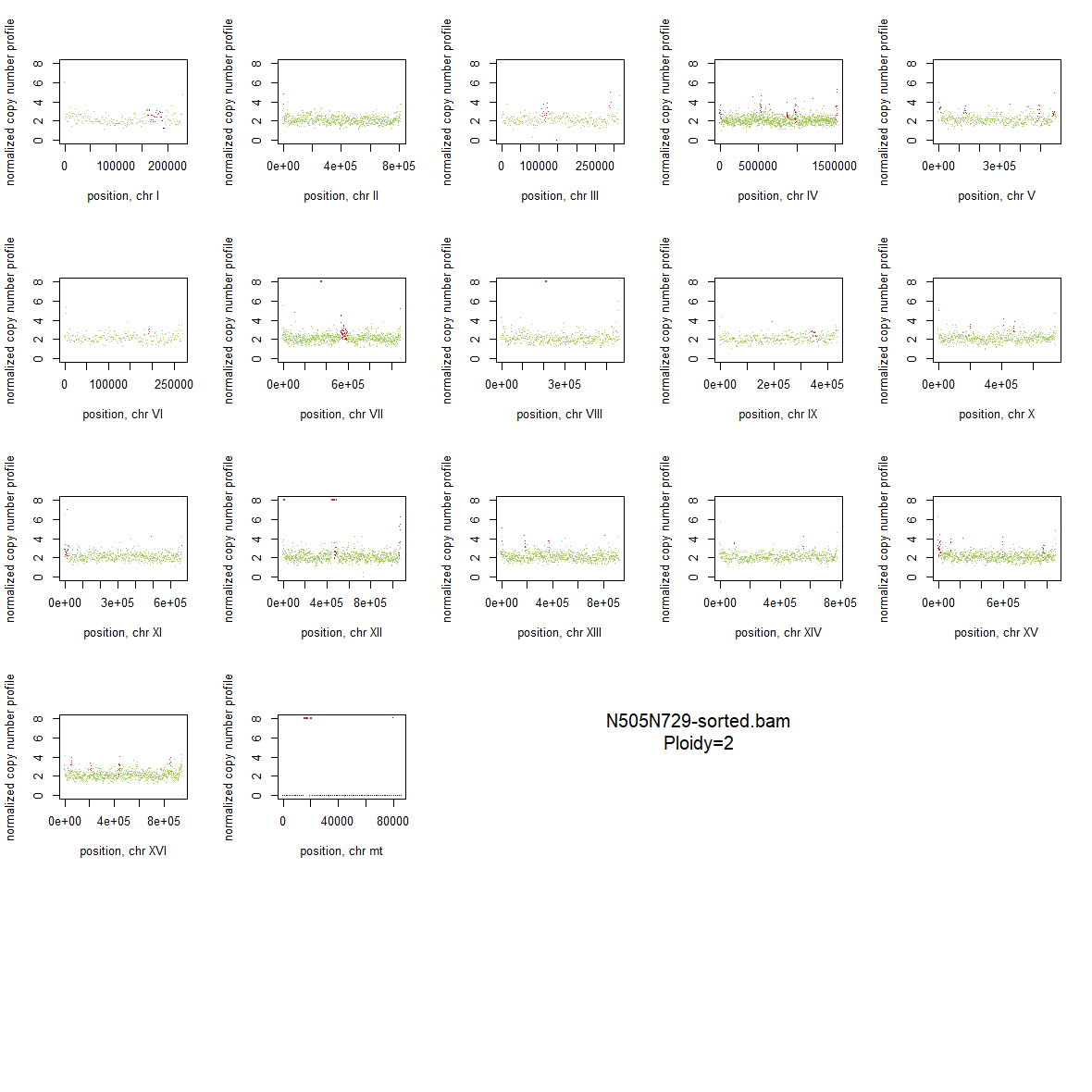

### 2x_F126L_17.png

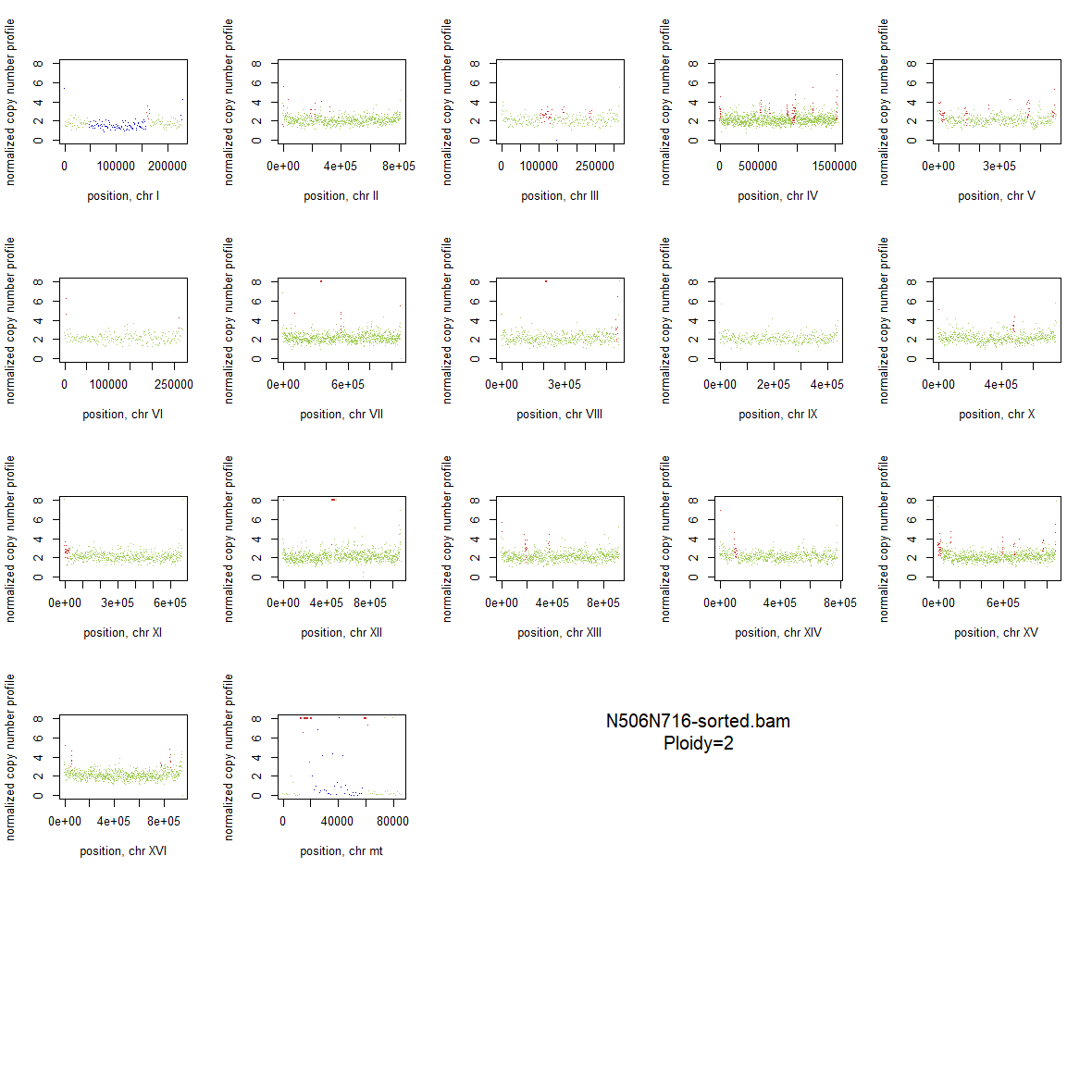

### 2x_F126L_18.png

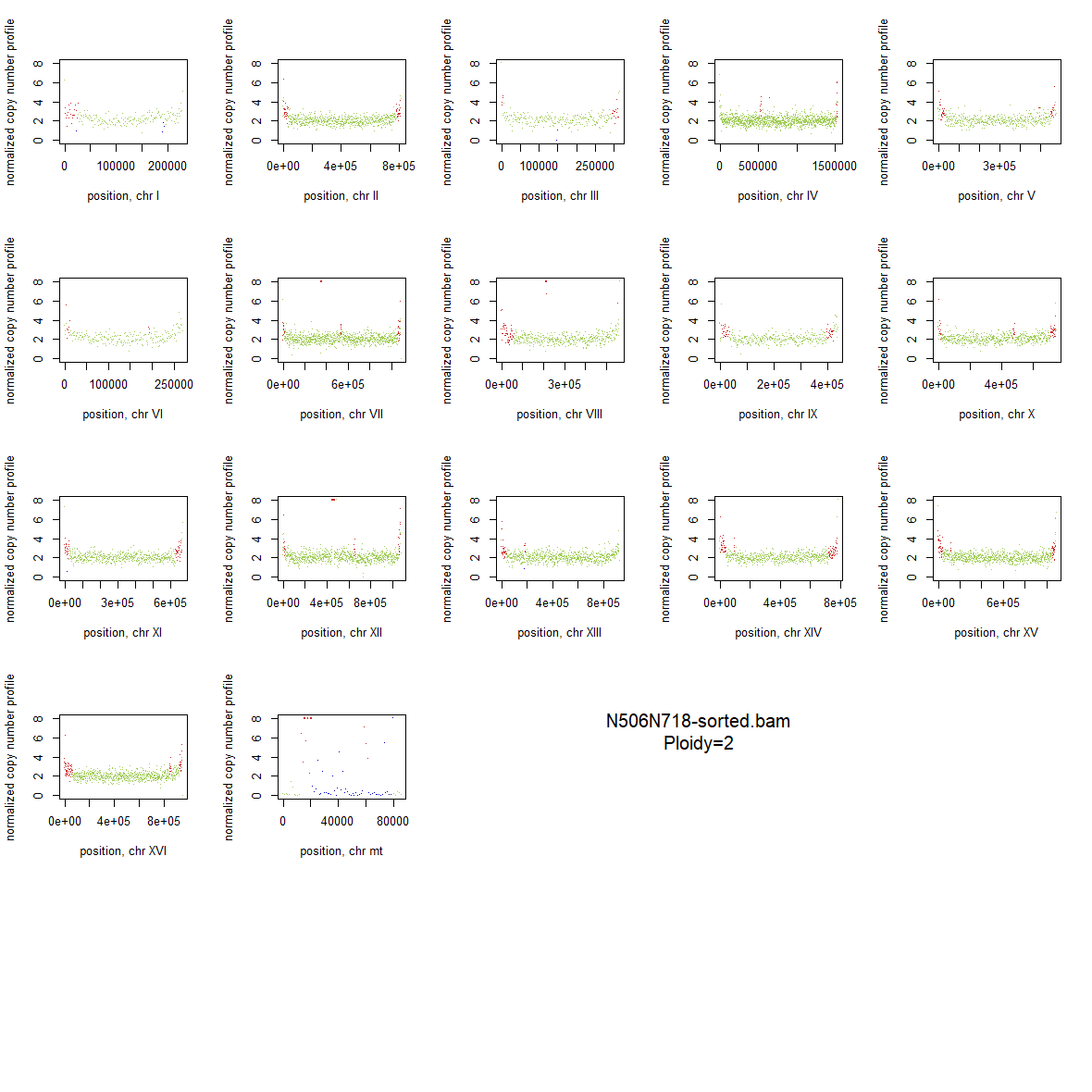

### 2x_F126L_19.png

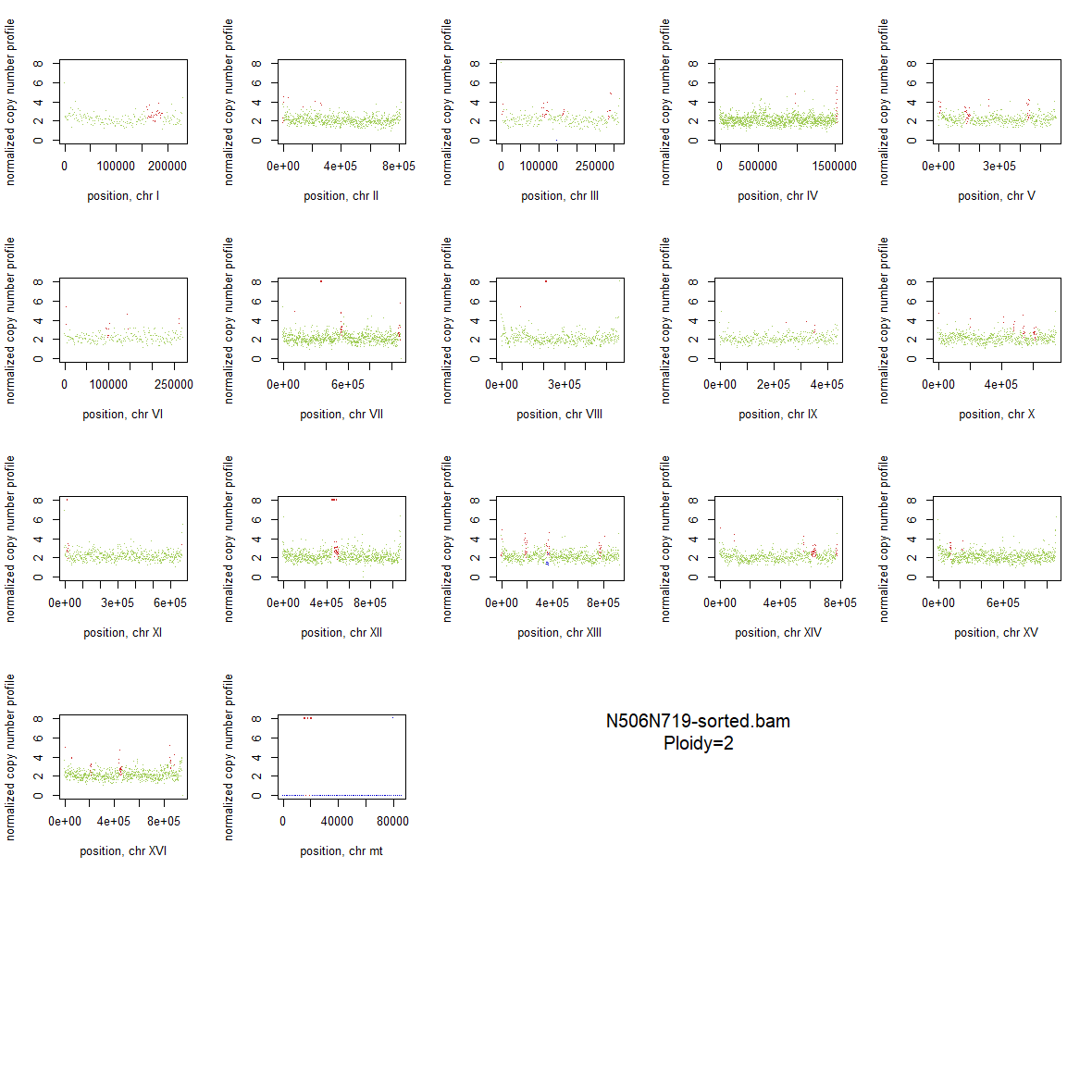

### 2x_F126L_20.png

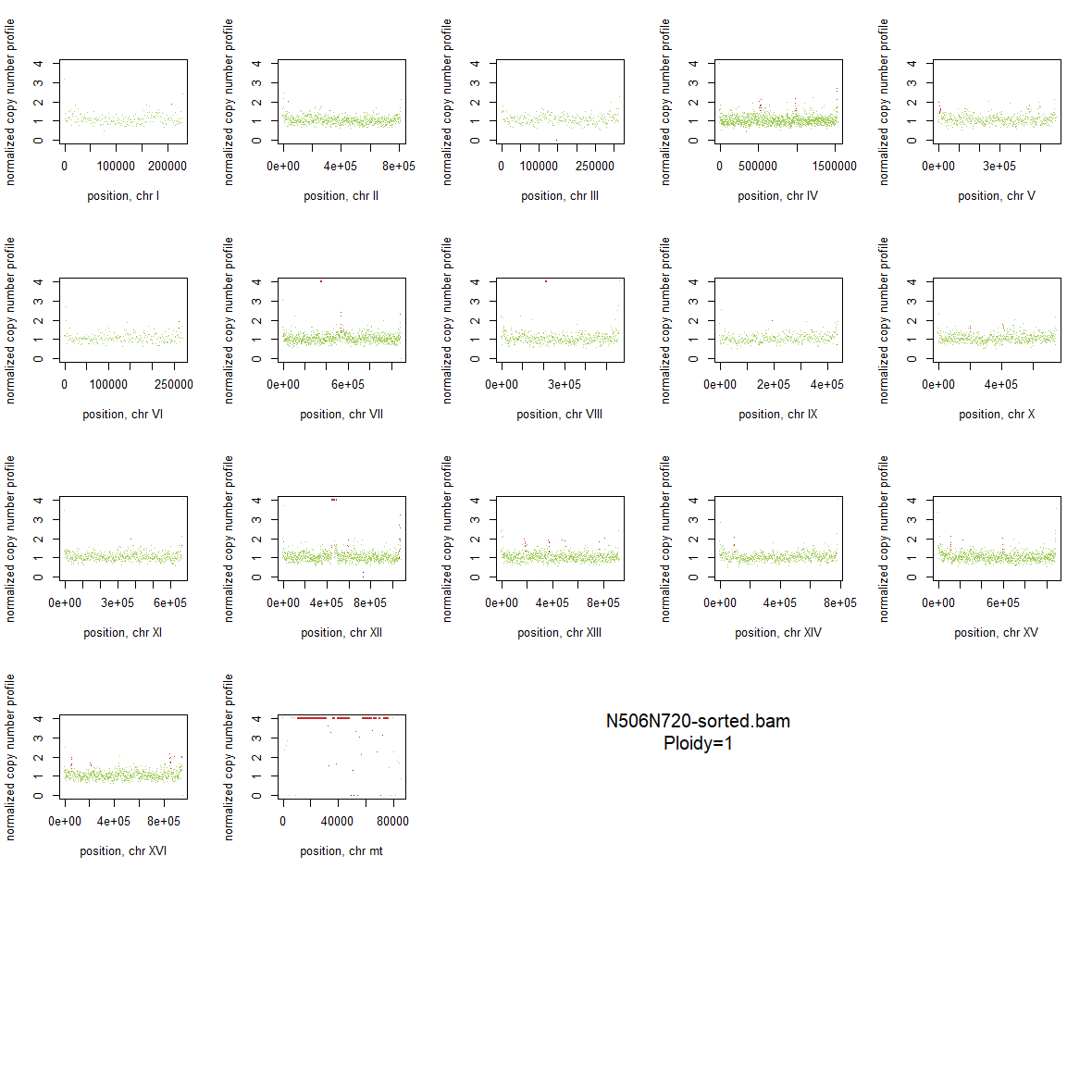

### 2x_F126L_21.png

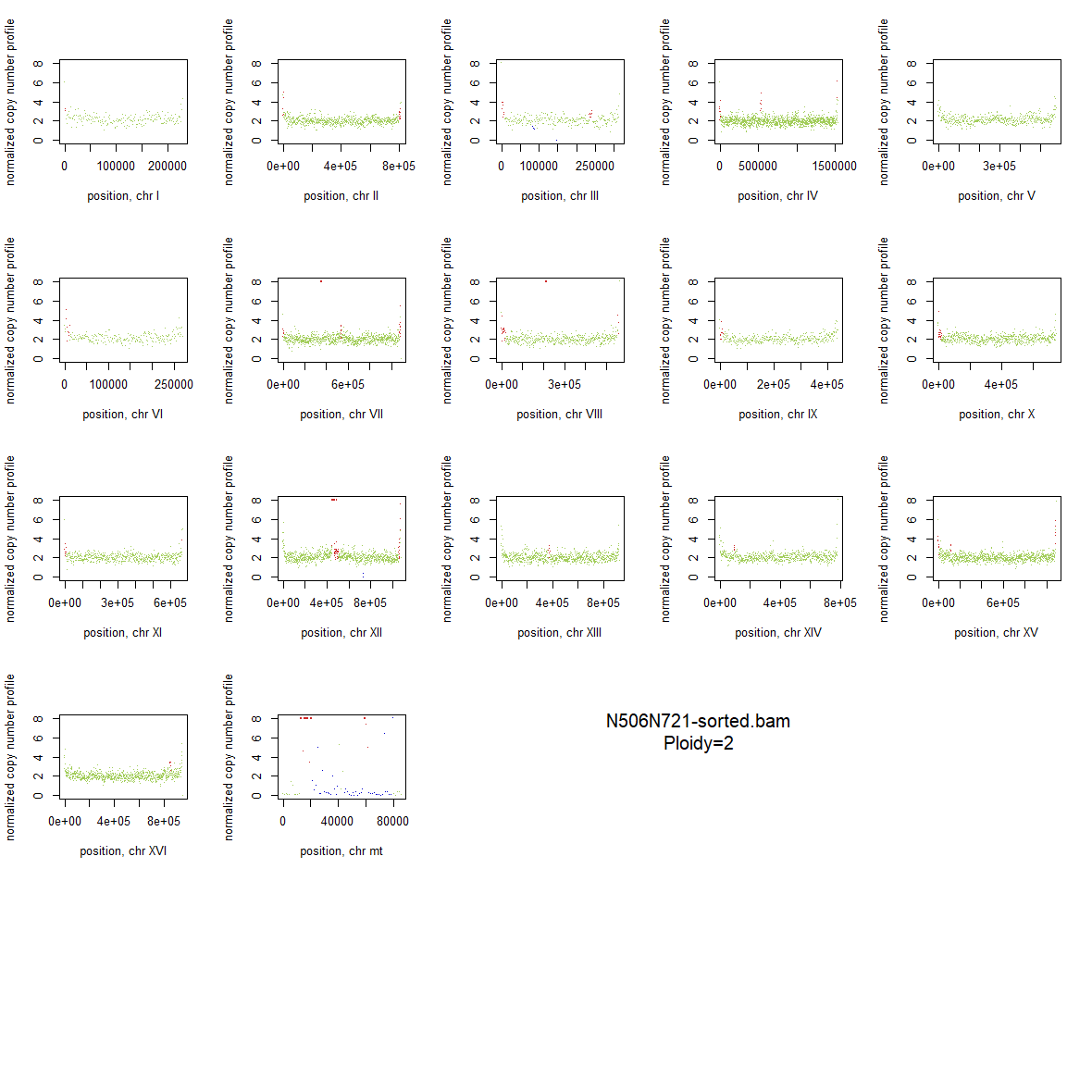

### 2x_F126L_22.png

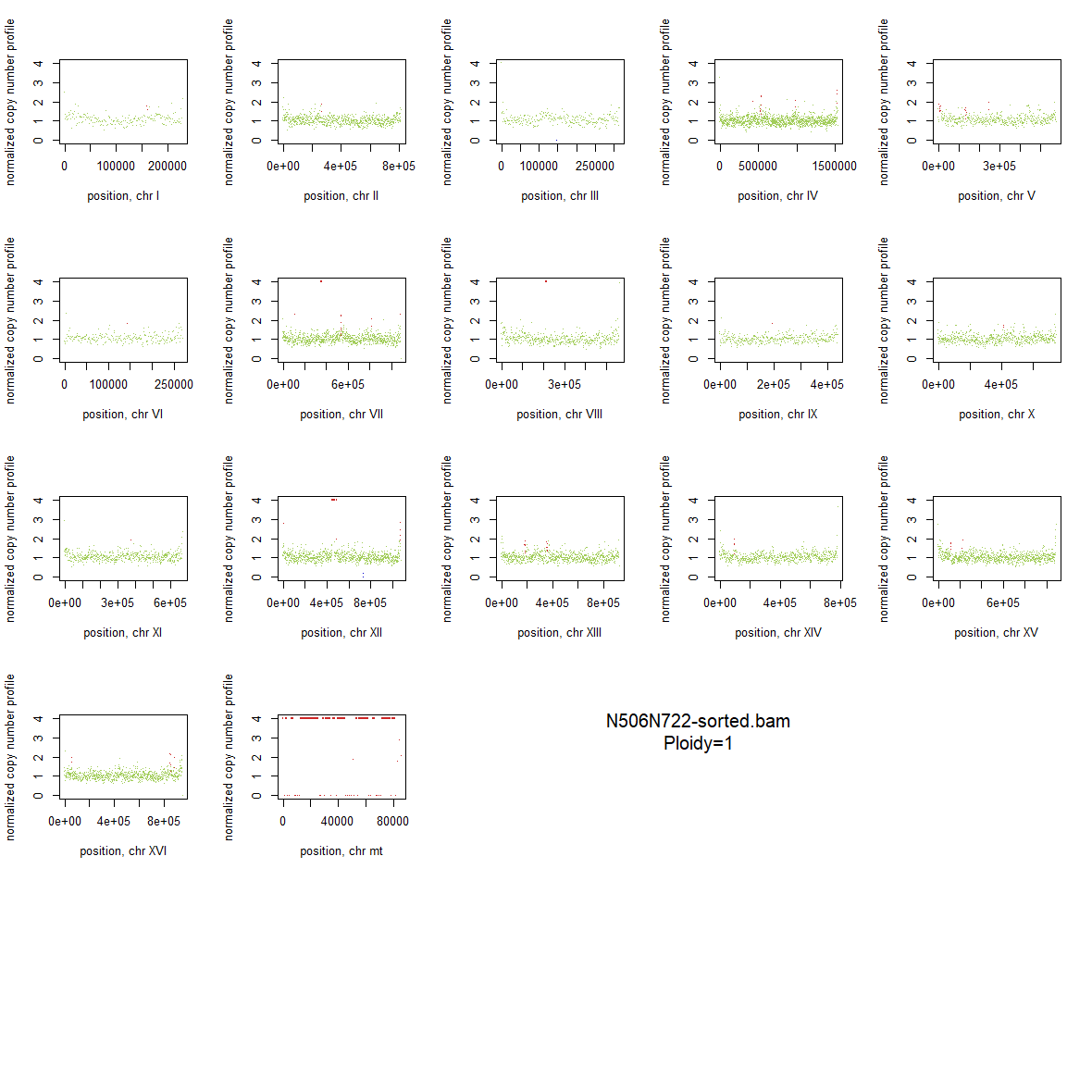

### 2x_F126L_23.png

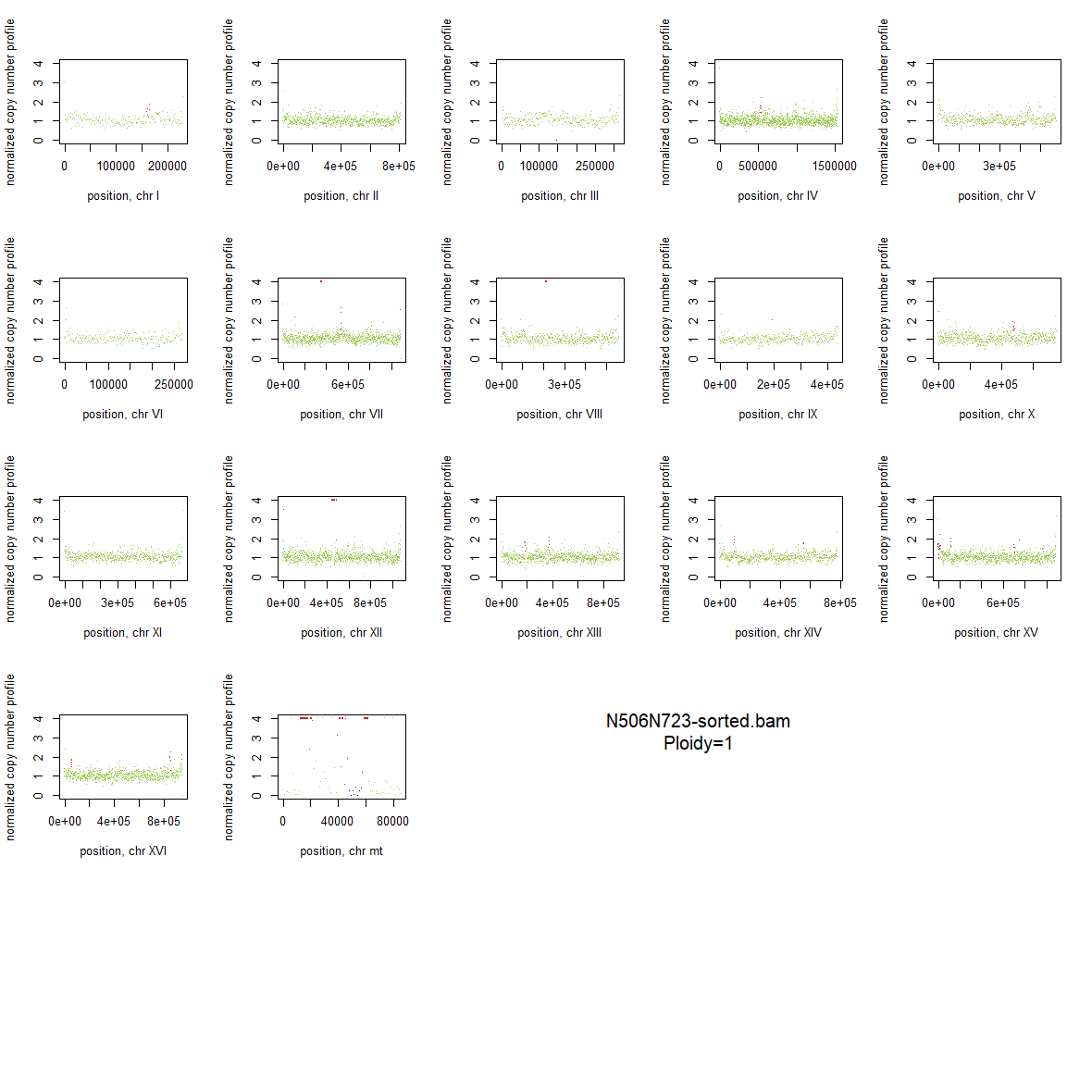

### 2x_F126L_24.png

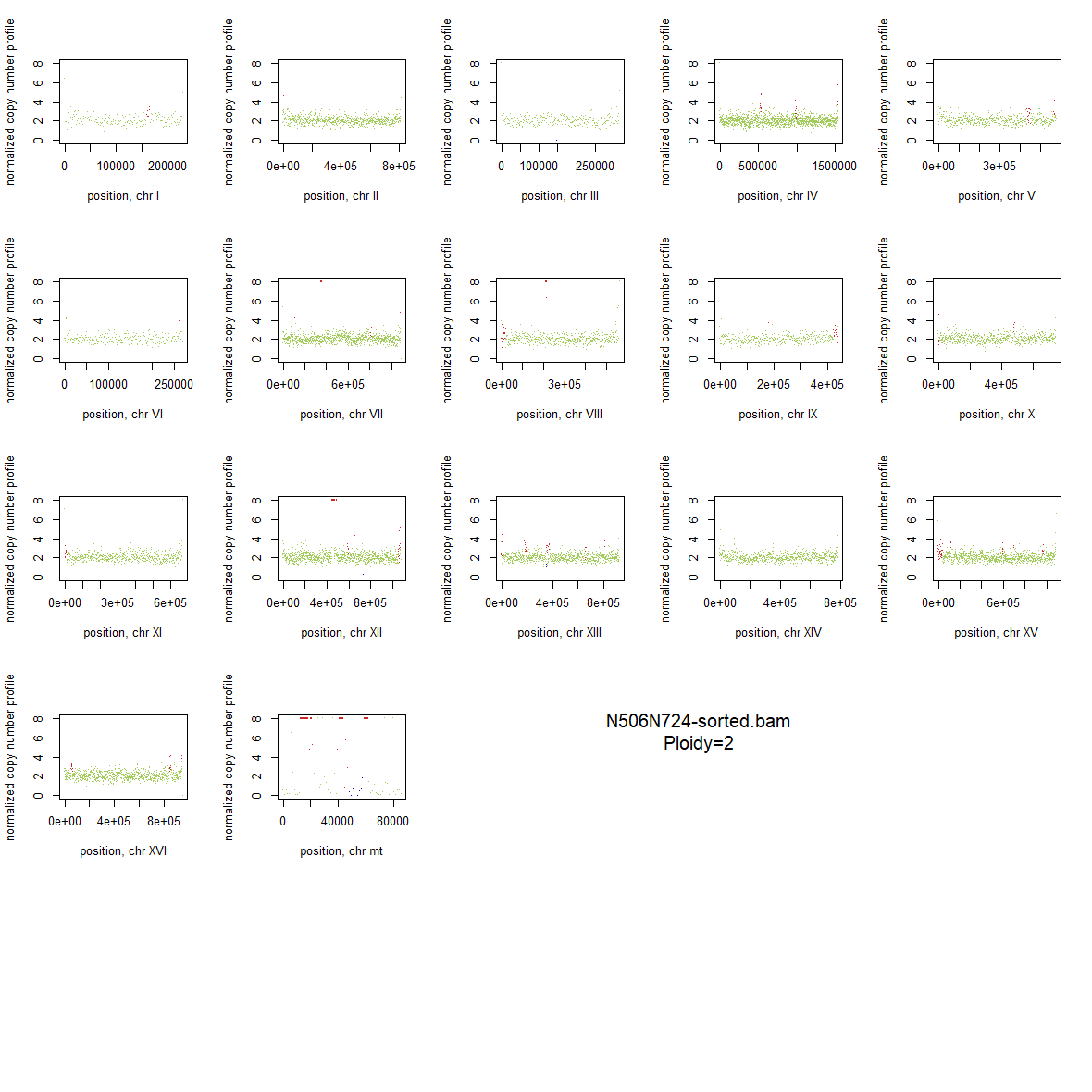

### 2x_F126L_25.png

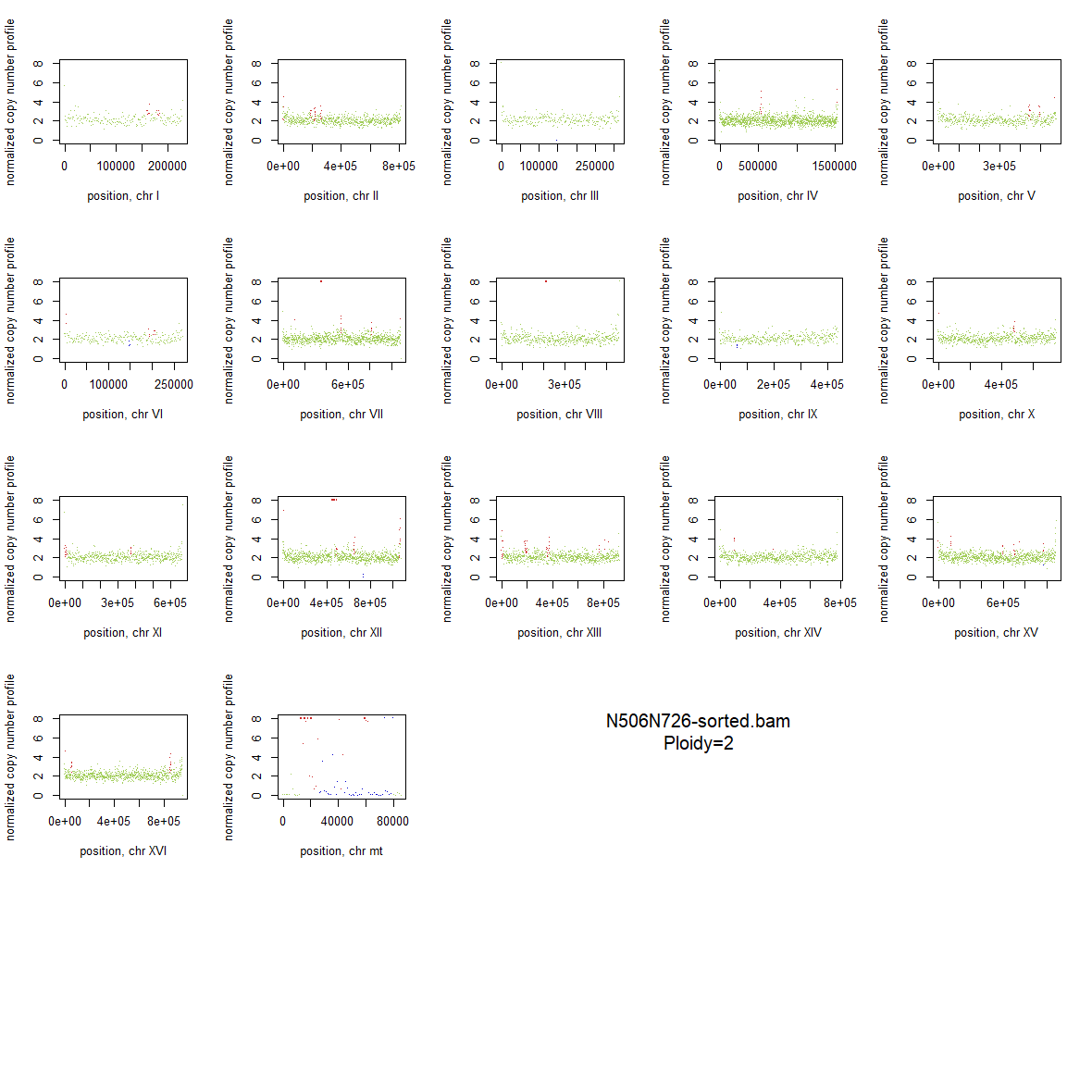

### 2x_F126L_26.png

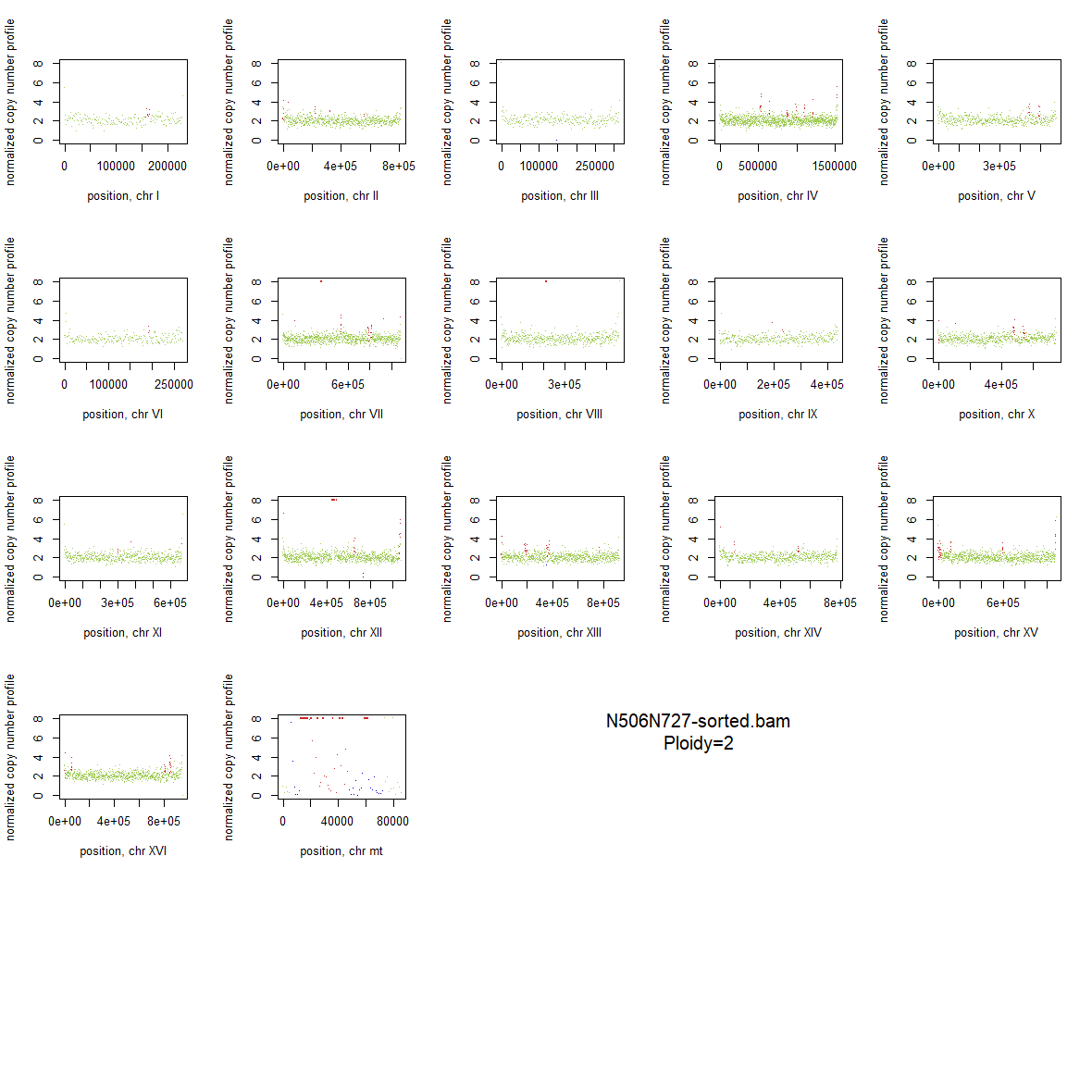

### 2x_F126L_27.png

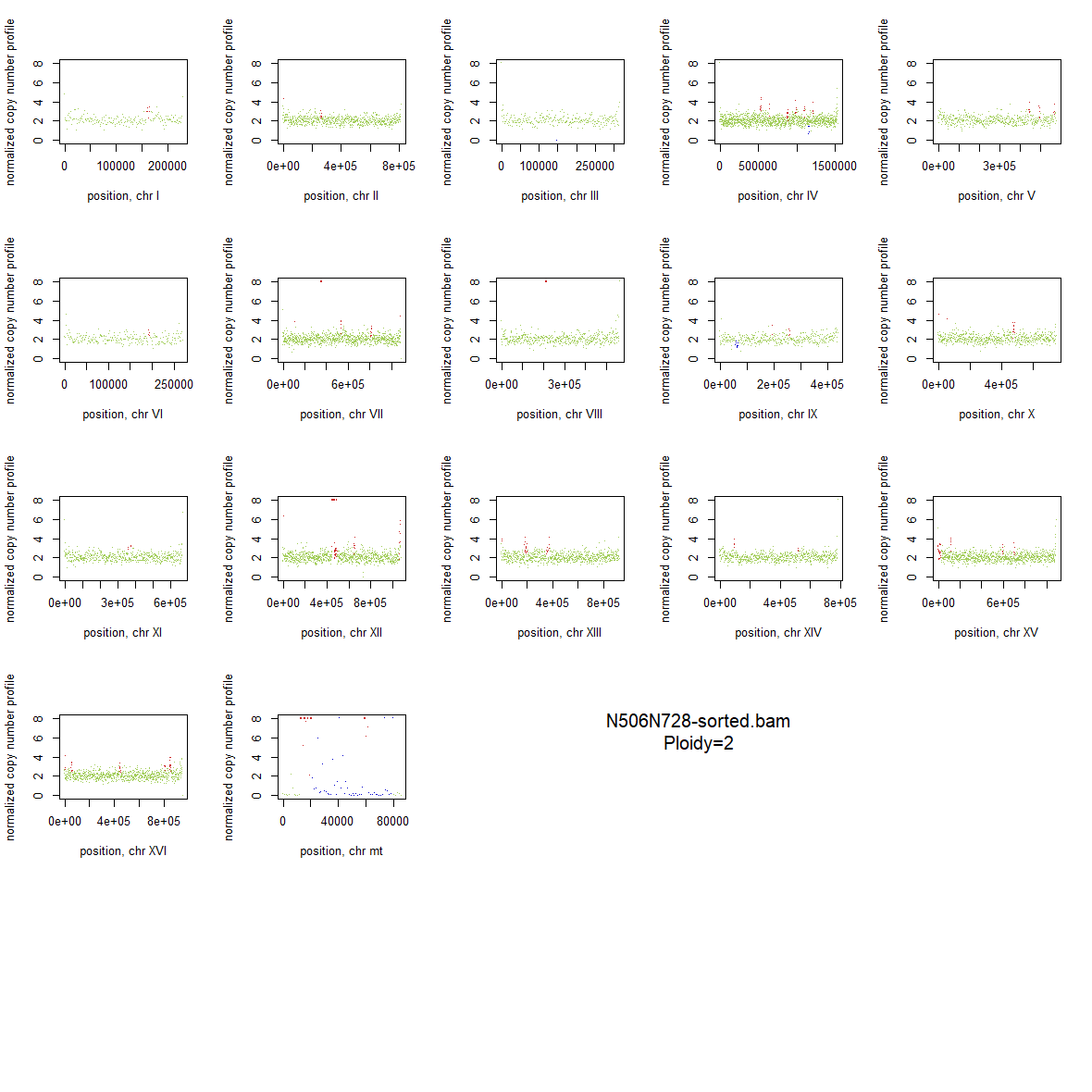

### 2x_F126L_28.png

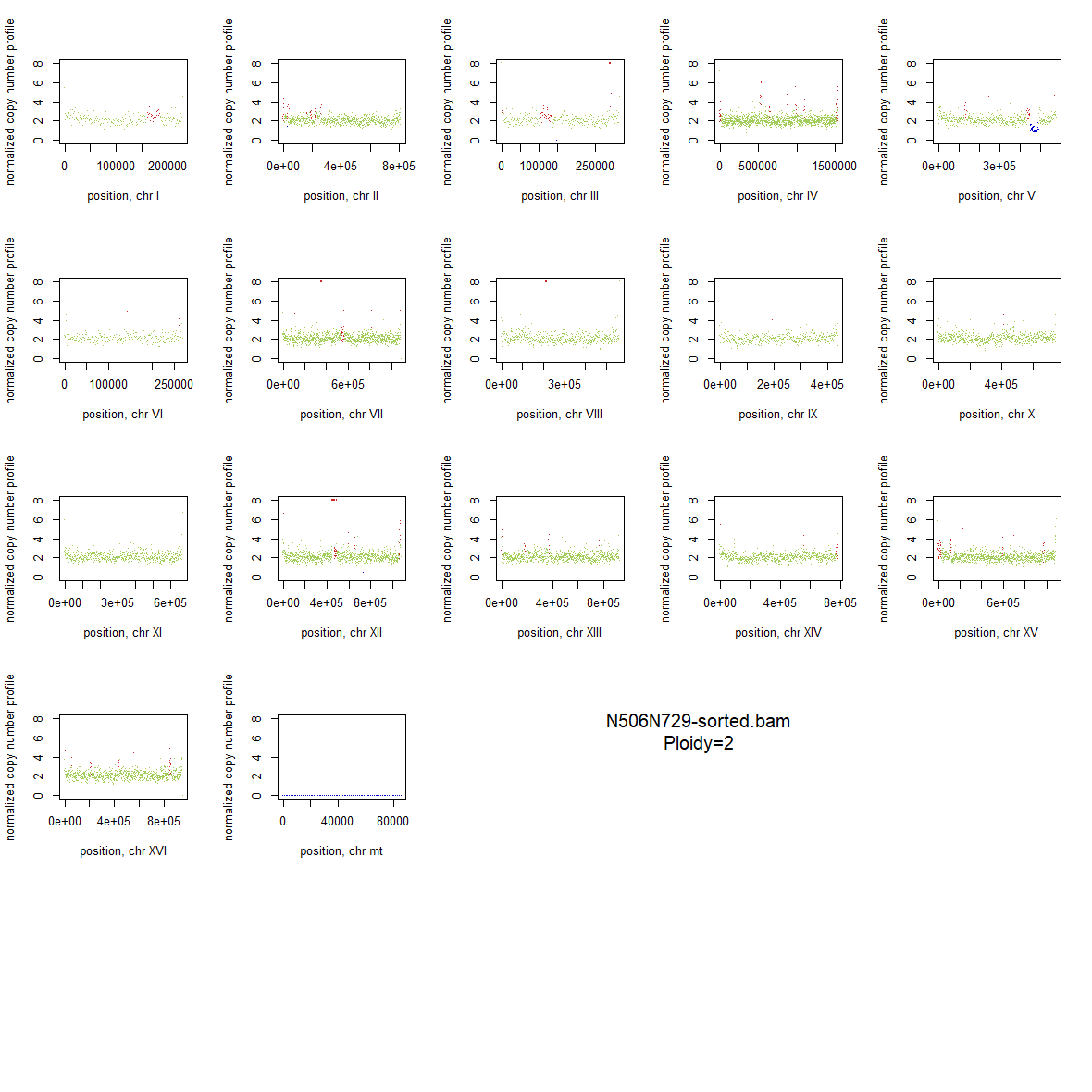

### 2x_F126L_29.png

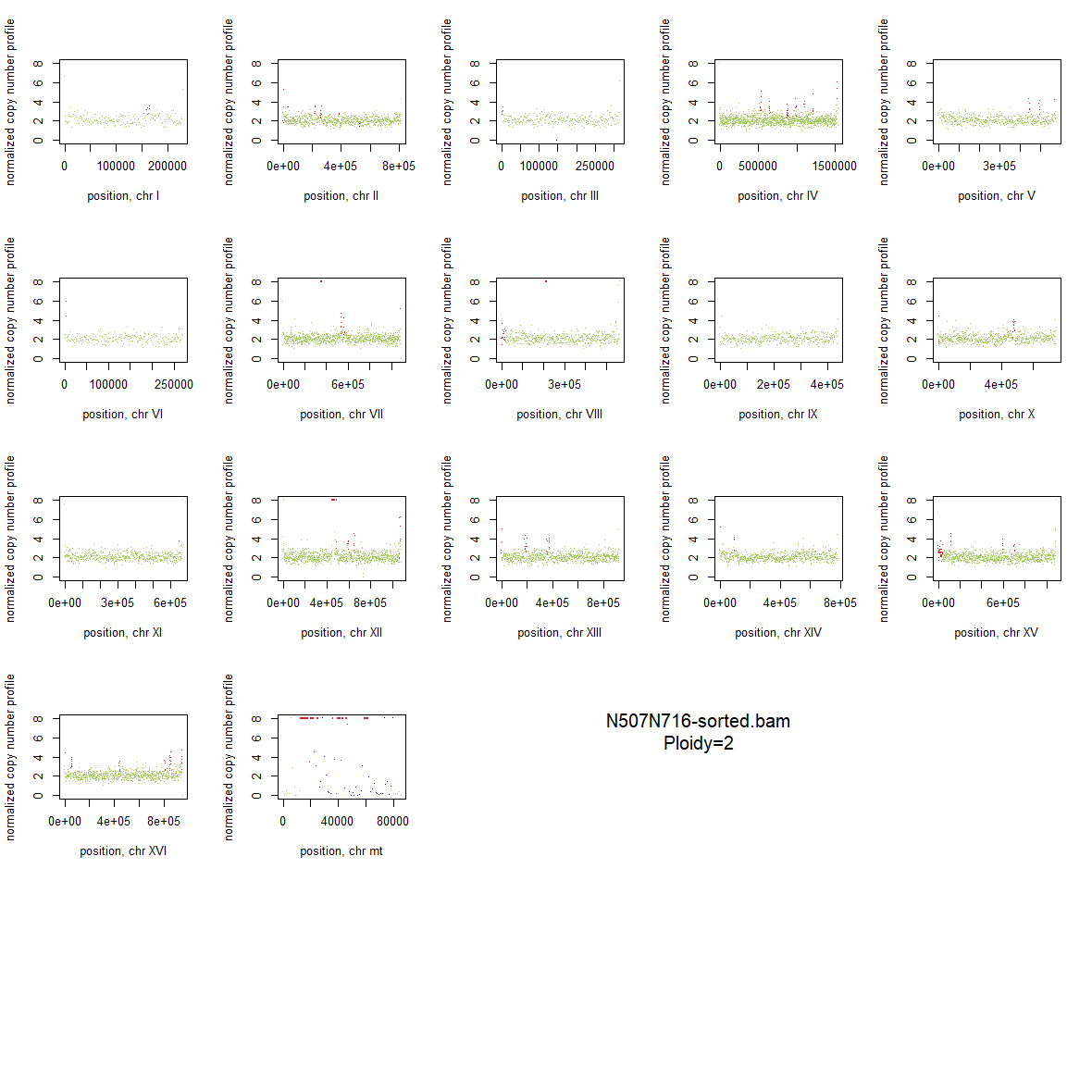

### 2x_F126L_30.png

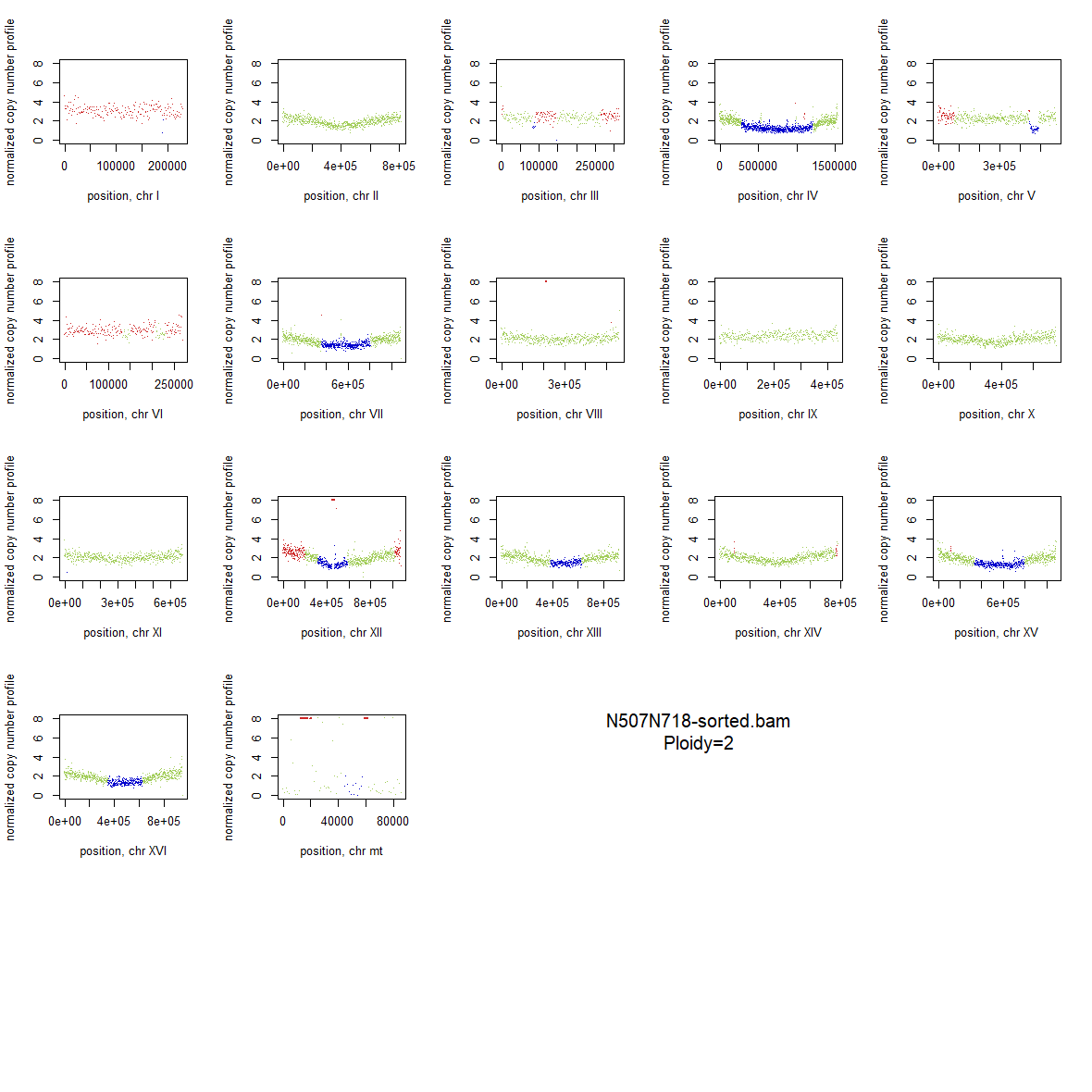

### 2x_F126L_31.png

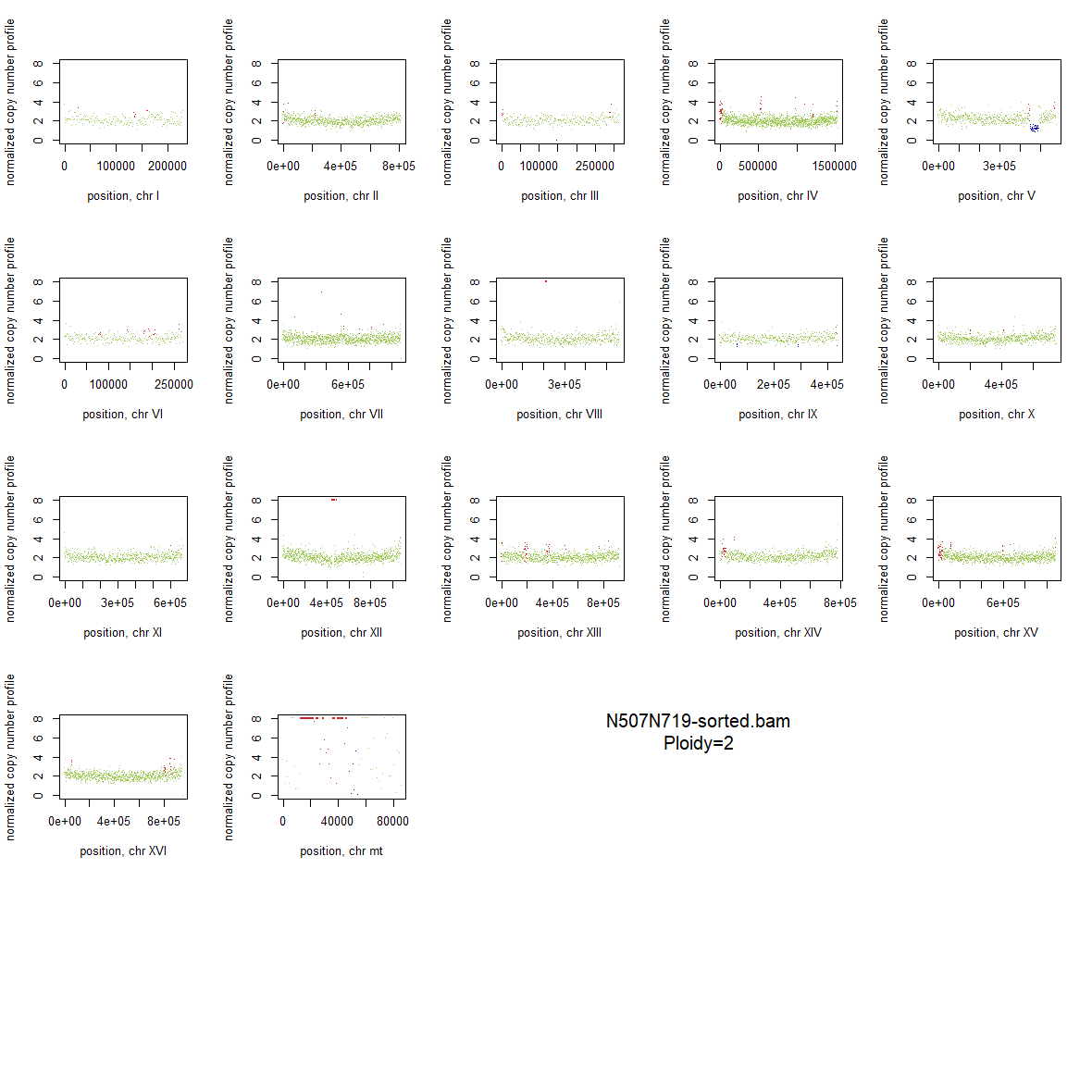

### 2x_F126L_32.png

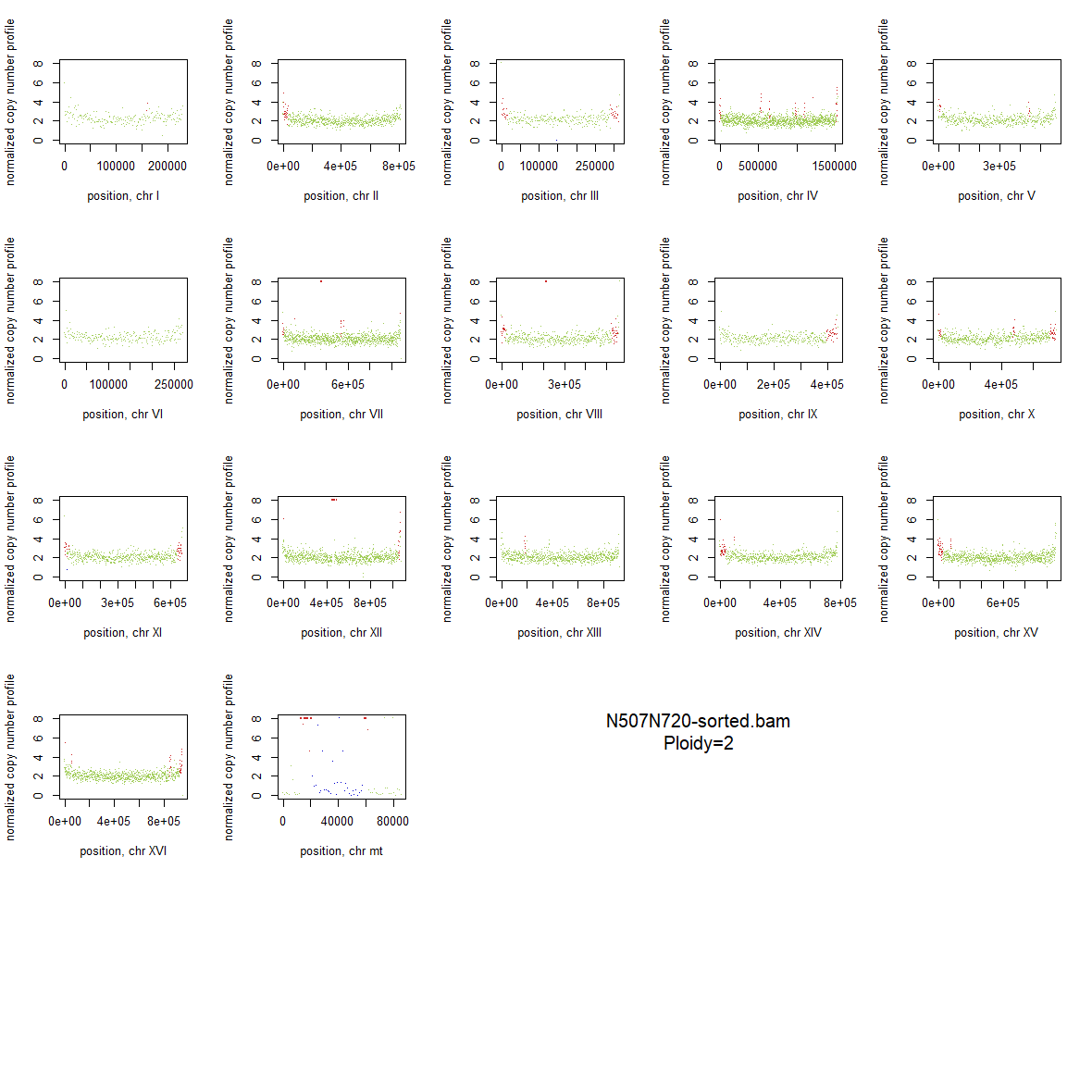

### 2x_F126L_33.png

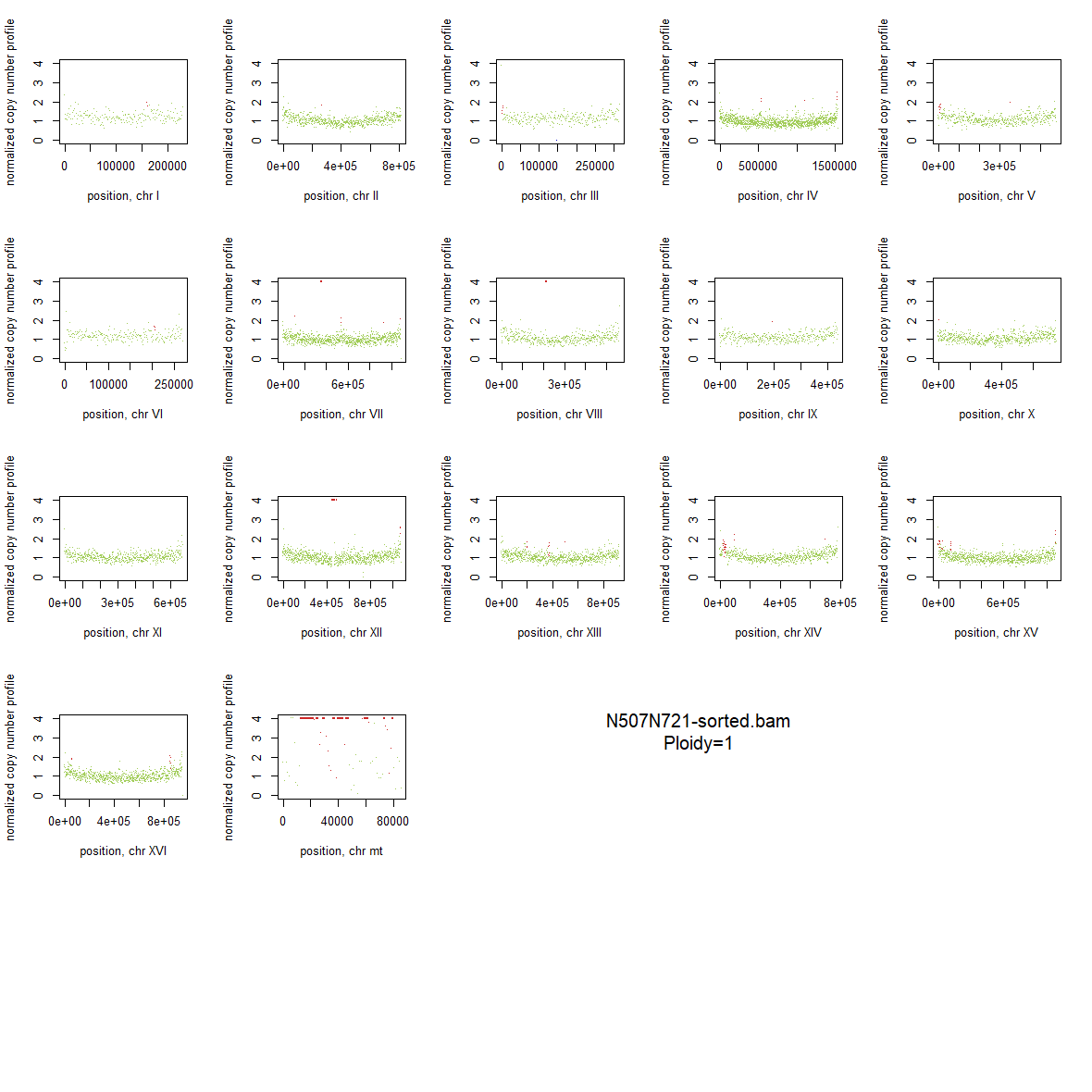

### 2x_F126L_34.png

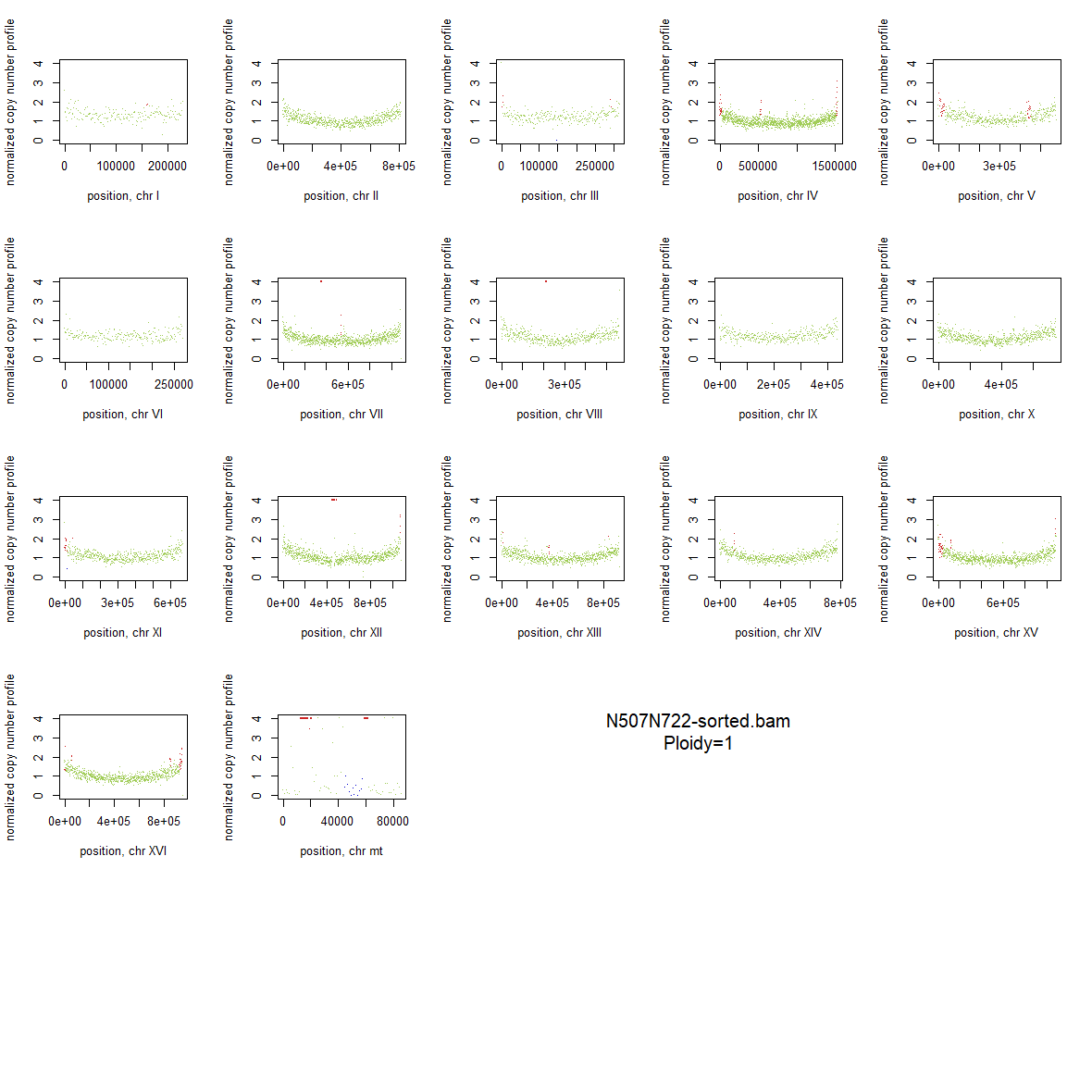

### 2x_F126L_35.png

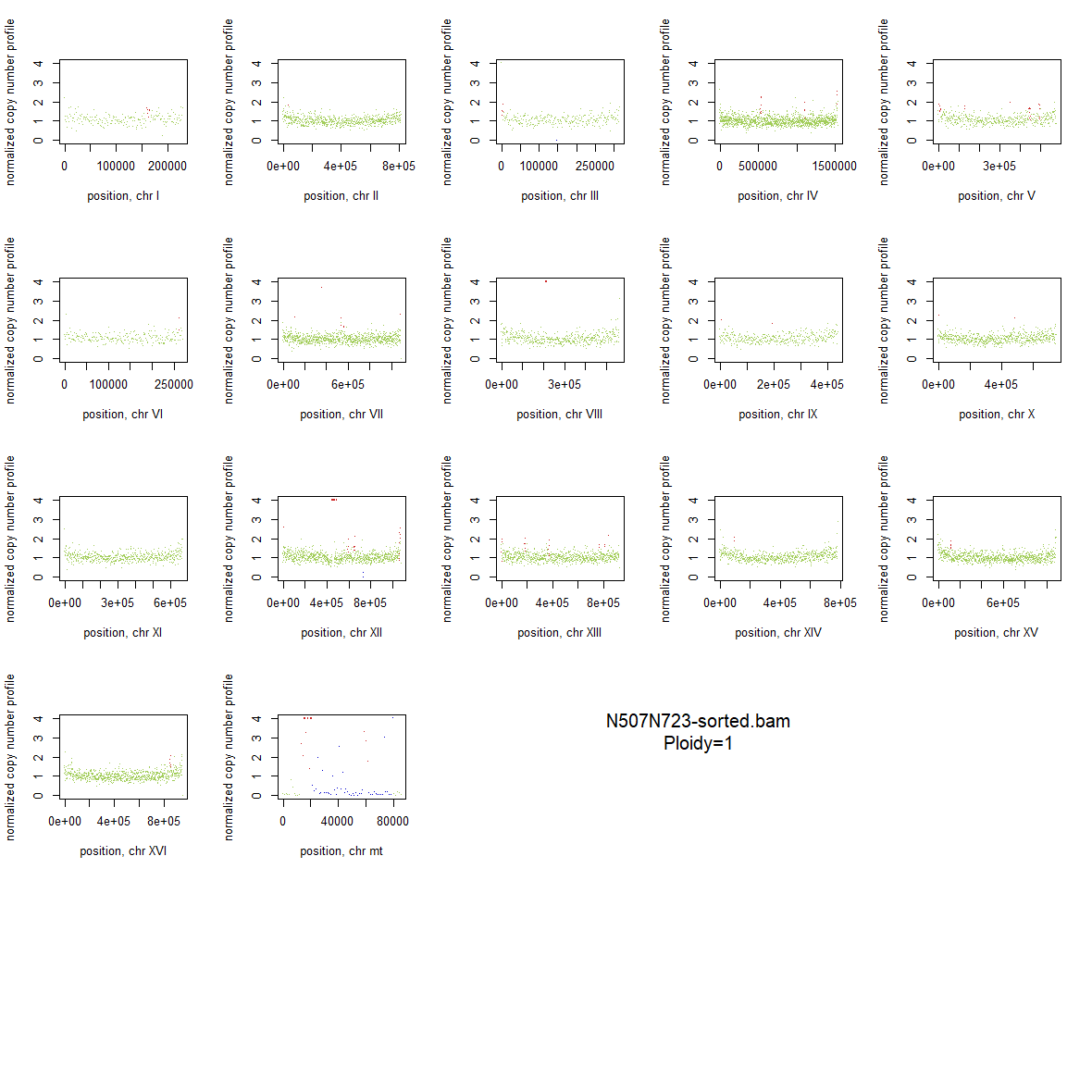

### 2x_F126L_36.png

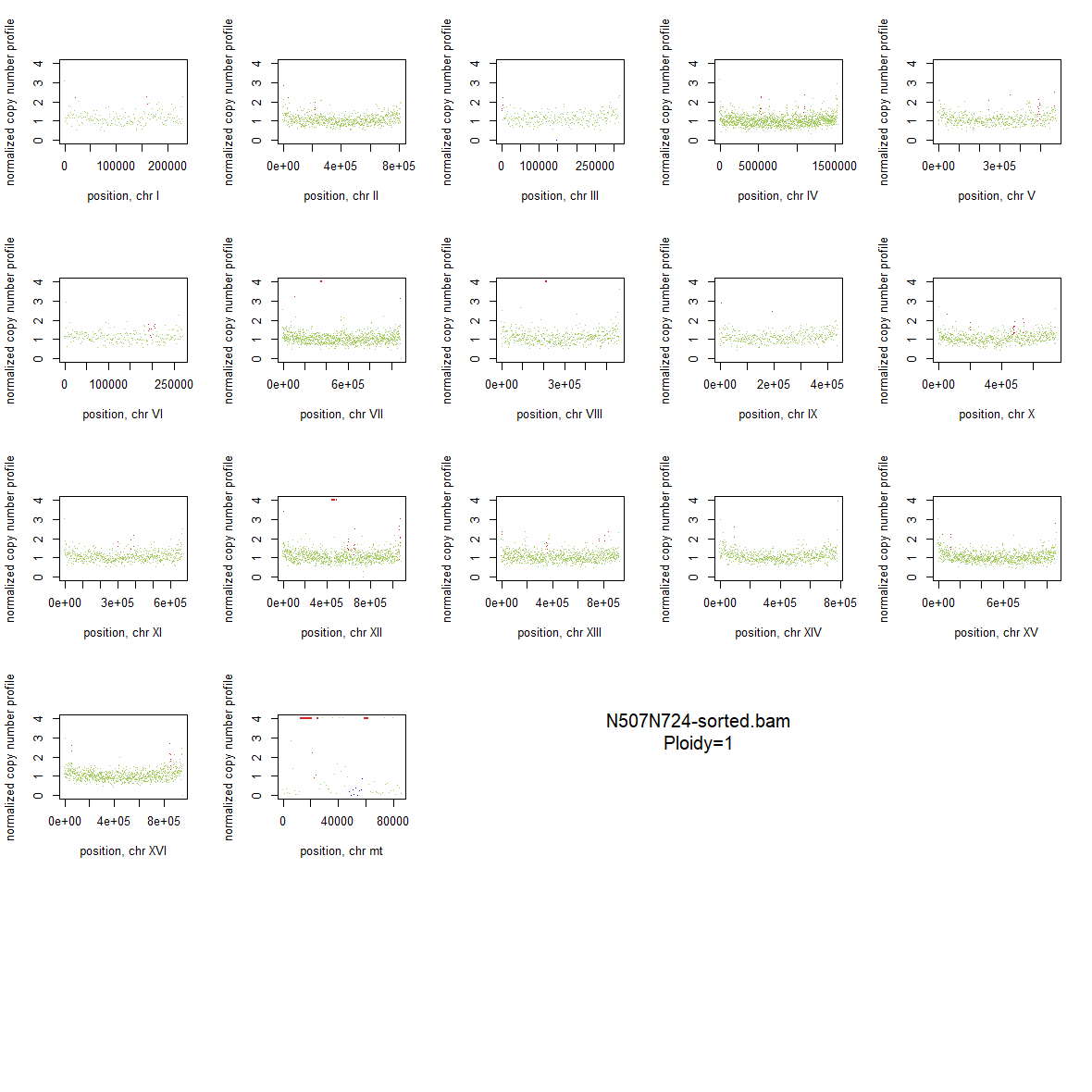

### 2x_Wildtype_10.png

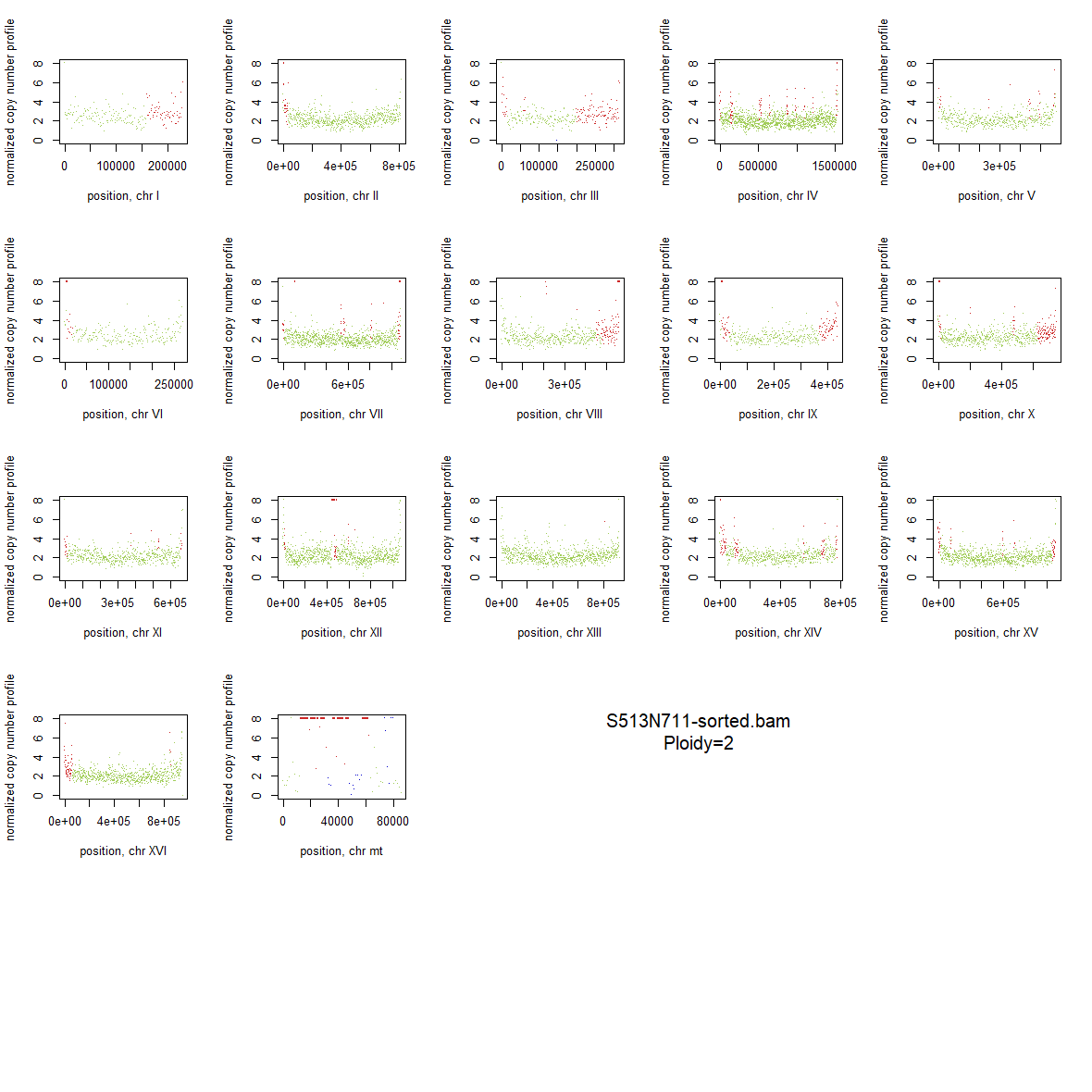

### 2x_Wildtype_11.png

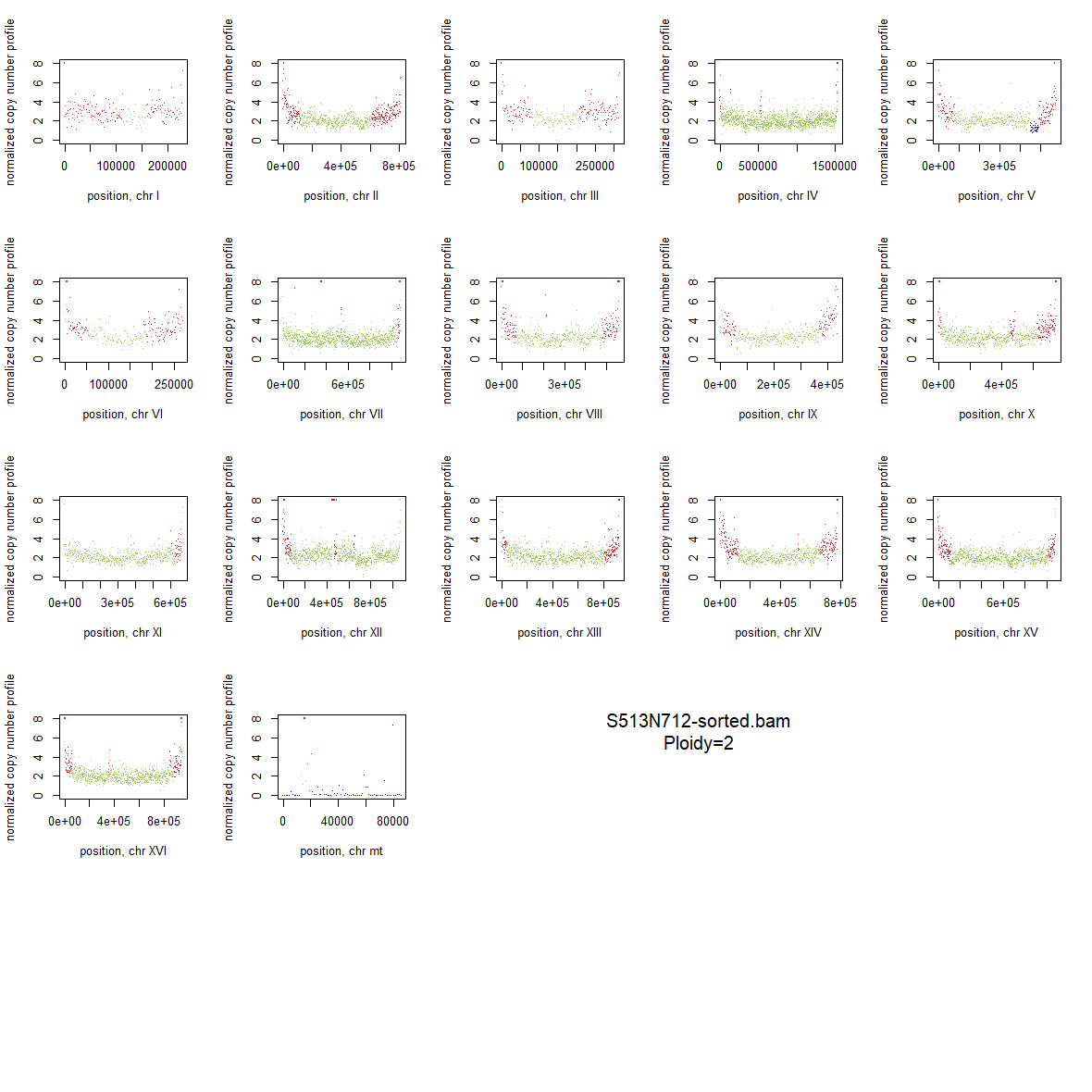

### blotB_V238M_CPY.tif

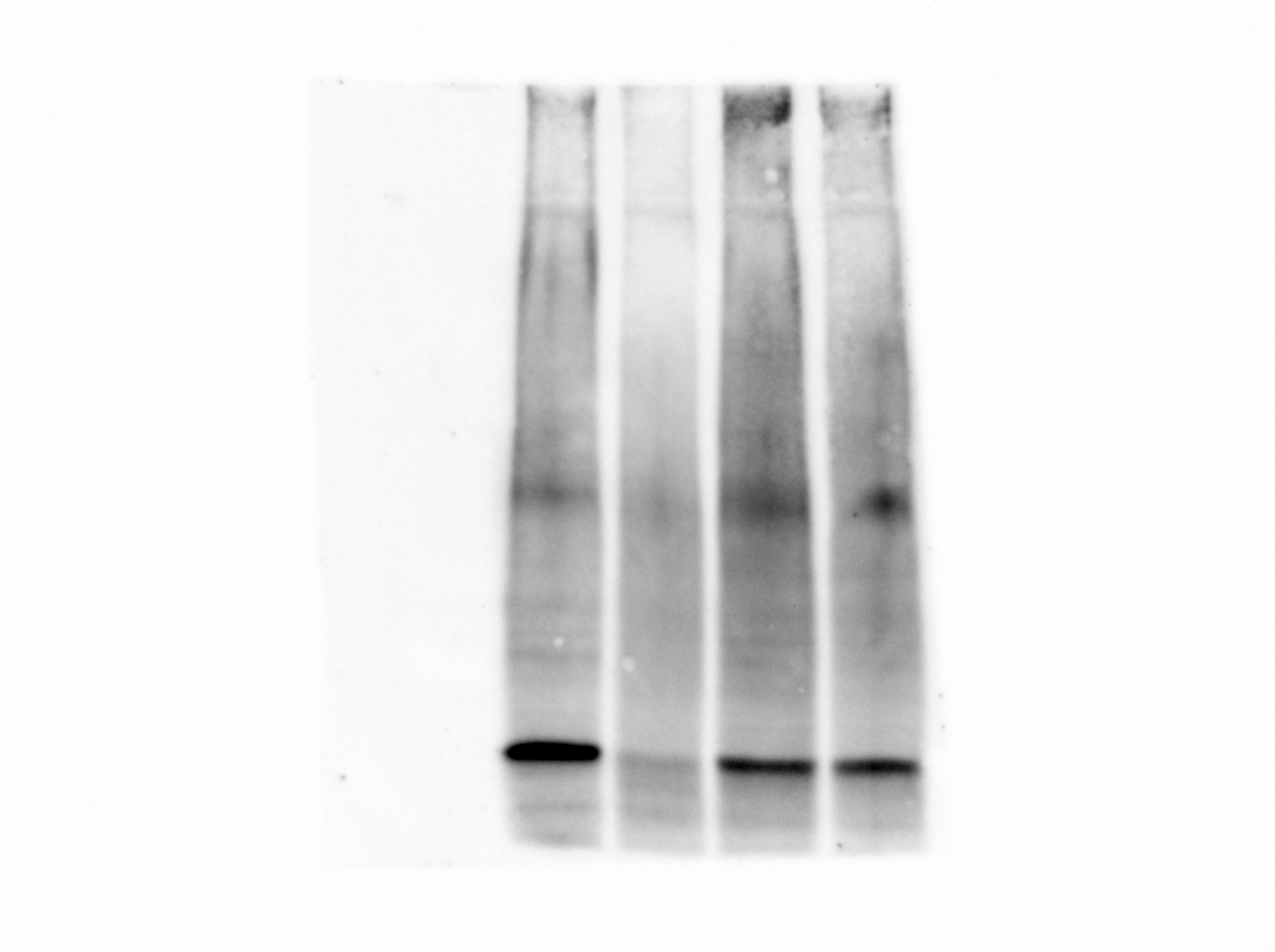

### blotC_V238M_invertase.tif

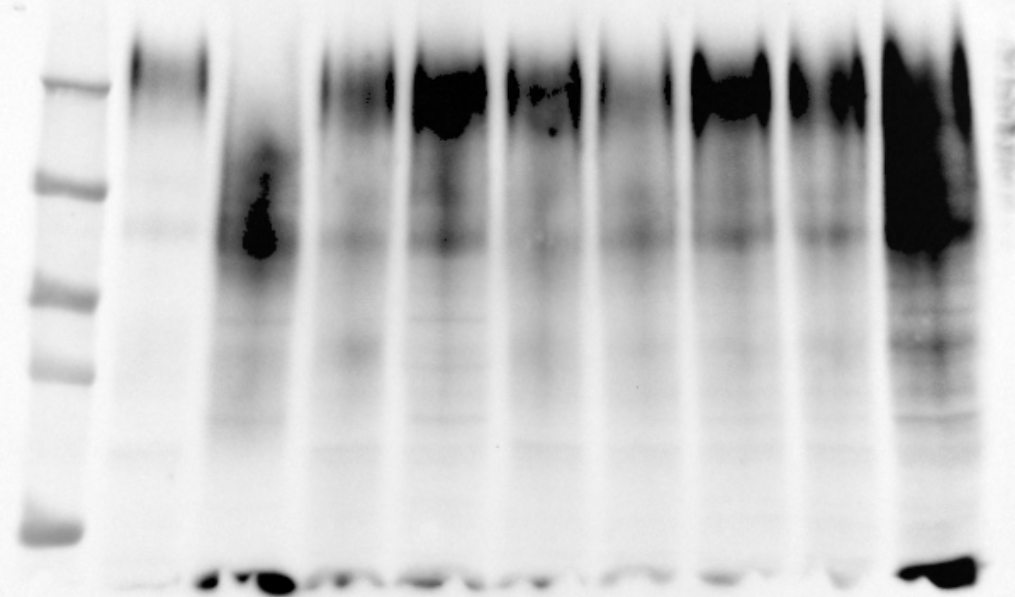

### blotD_WT_invertase.tif

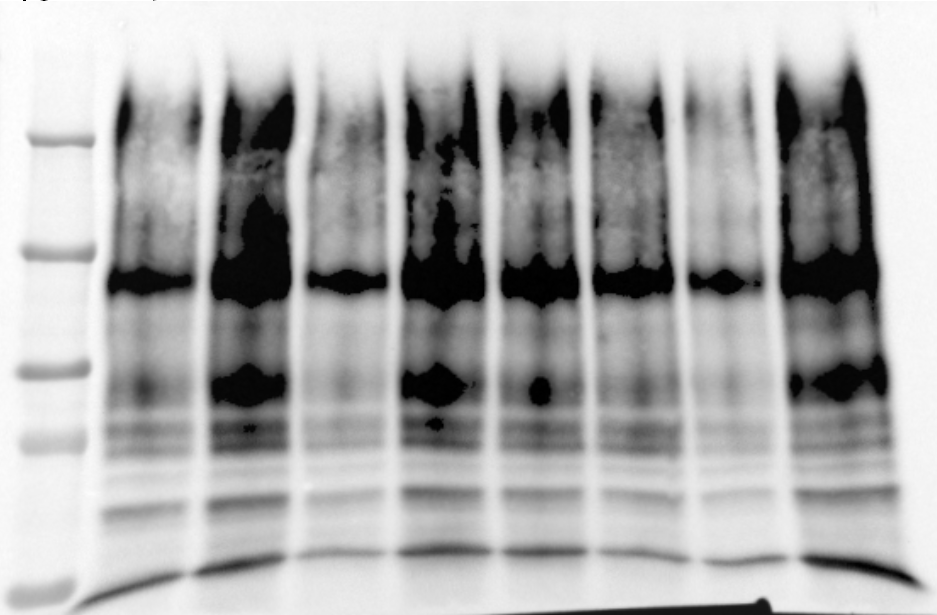

### blotE_WT_CPY.tif

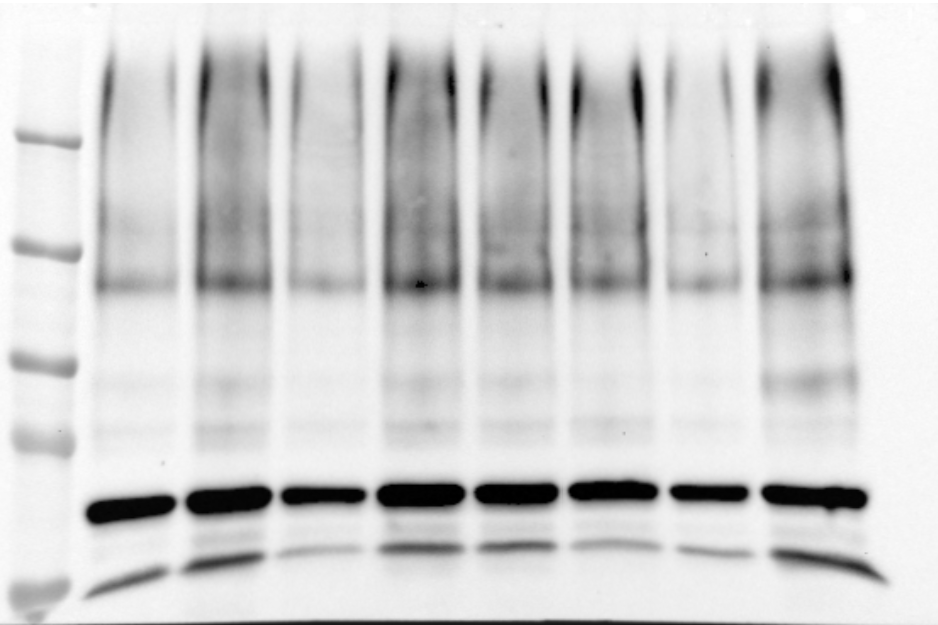

### blotF_F126L_invertase.tif

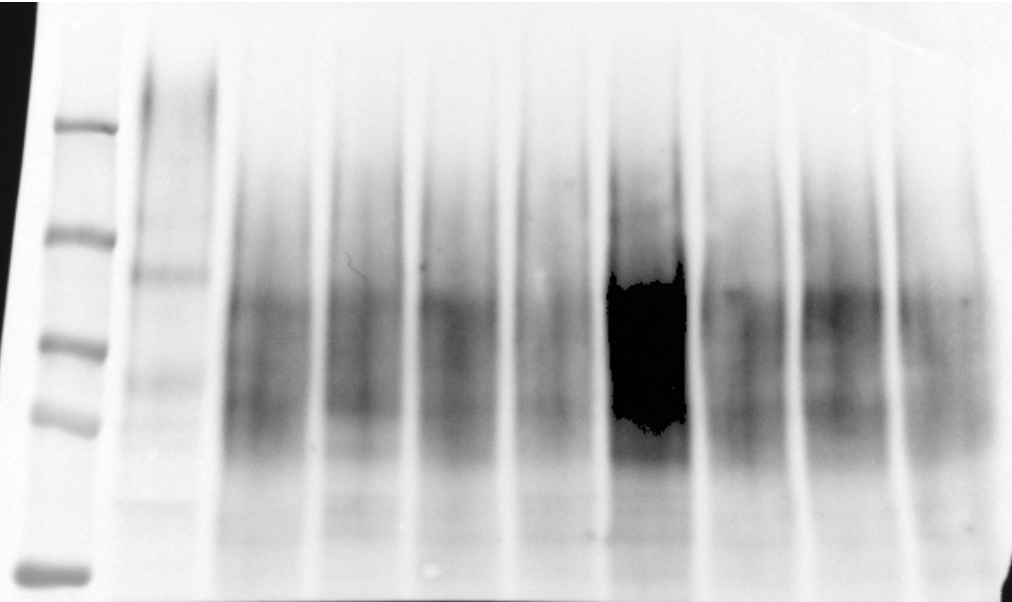

### ponceauA_V238M_invertase2.tif

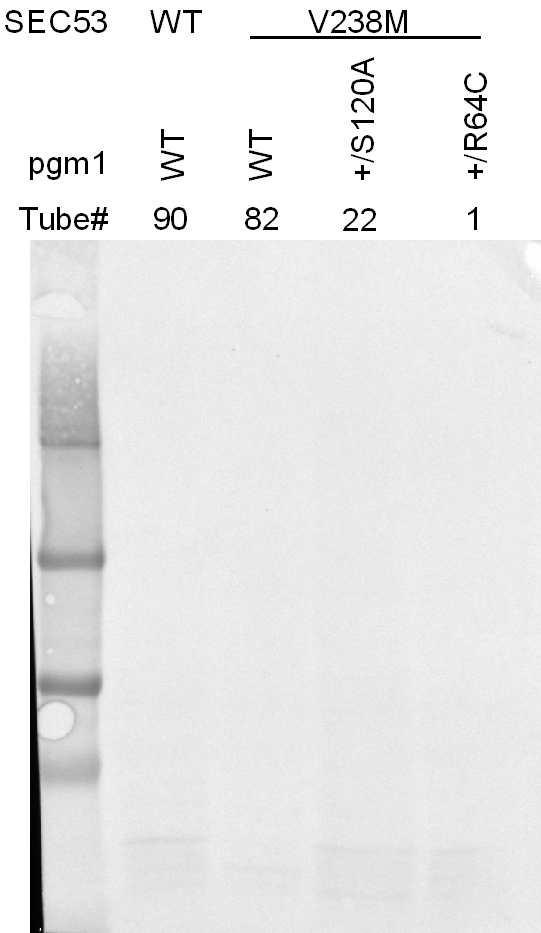

### ponceauA_V238M_invertase.tif

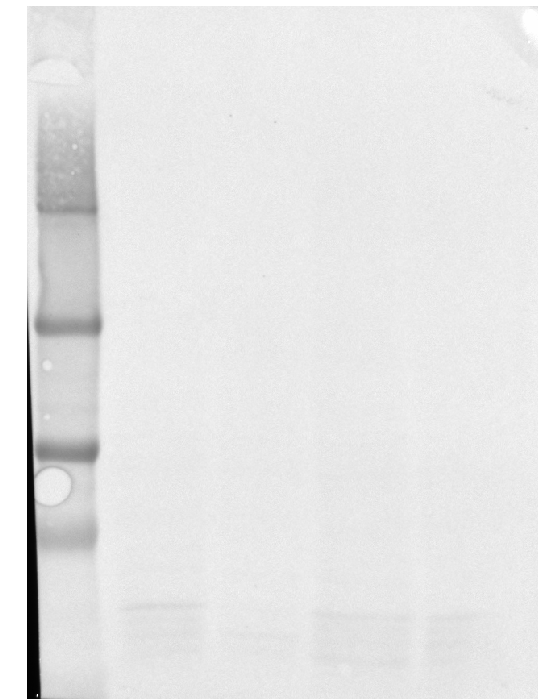

### ponceauB_V238M_CPY2.tif

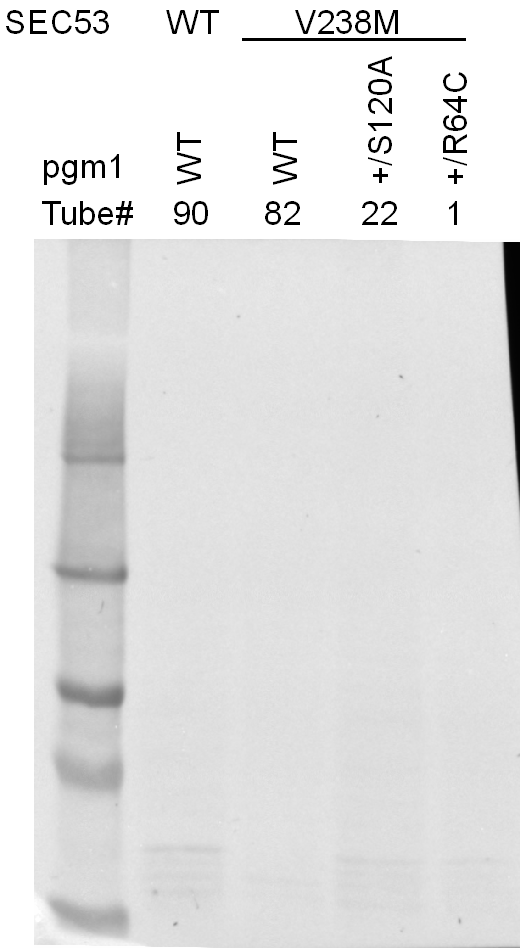

### ponceauB_V238M_CPY.tif

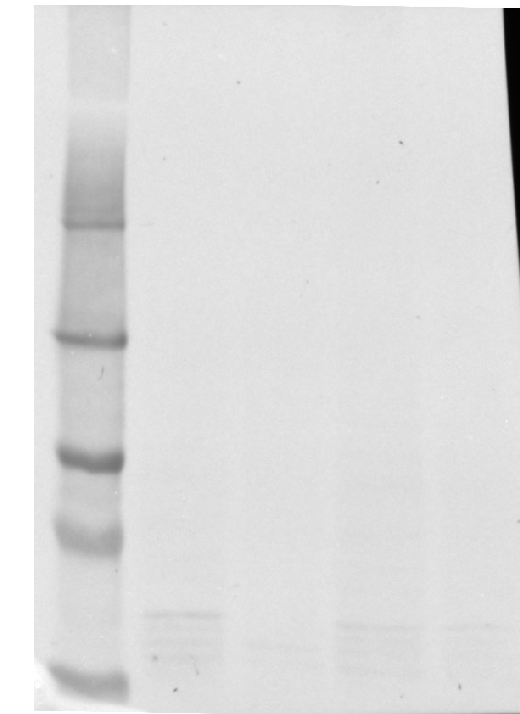

### ponceauC_V238M_invertase2.tif

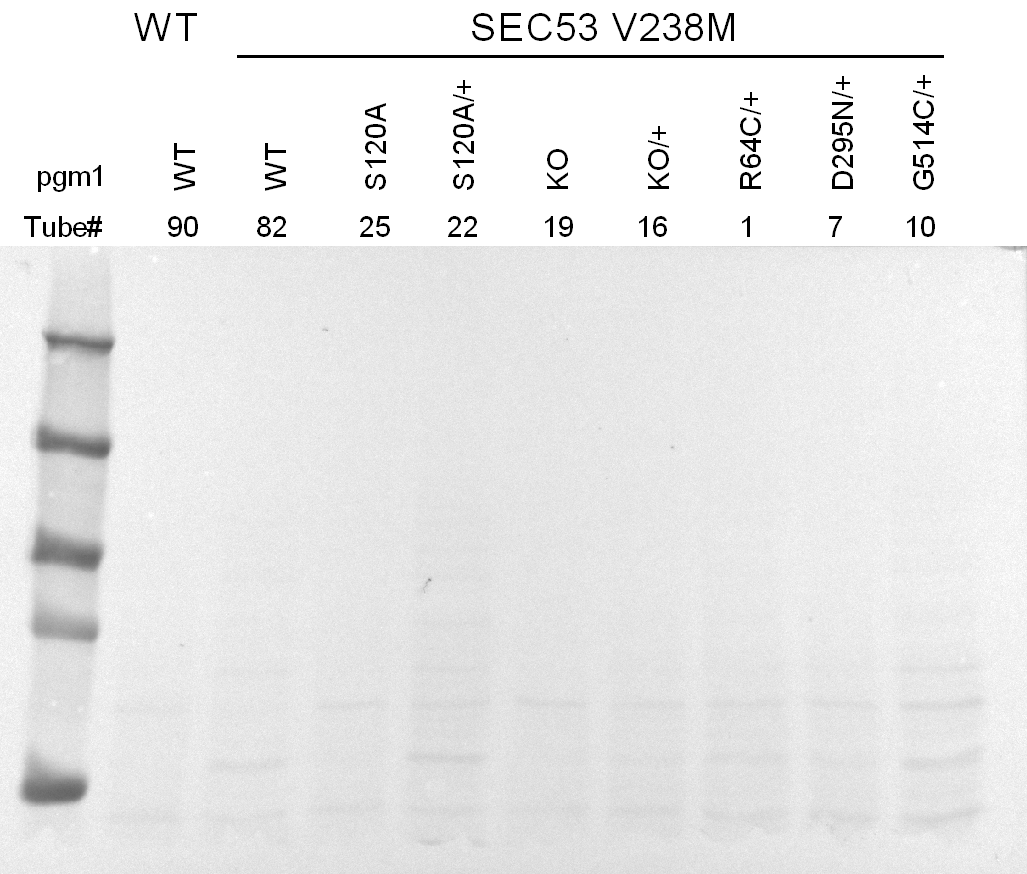

### ponceauC_V238M_invertase.tif

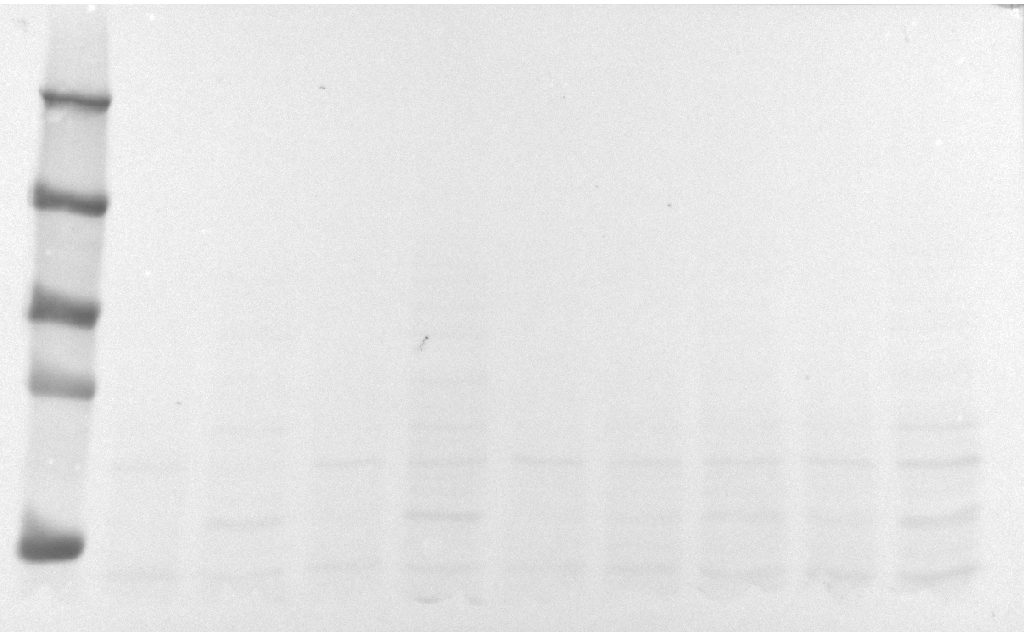

### ponceauD_WT_invertase2.tif

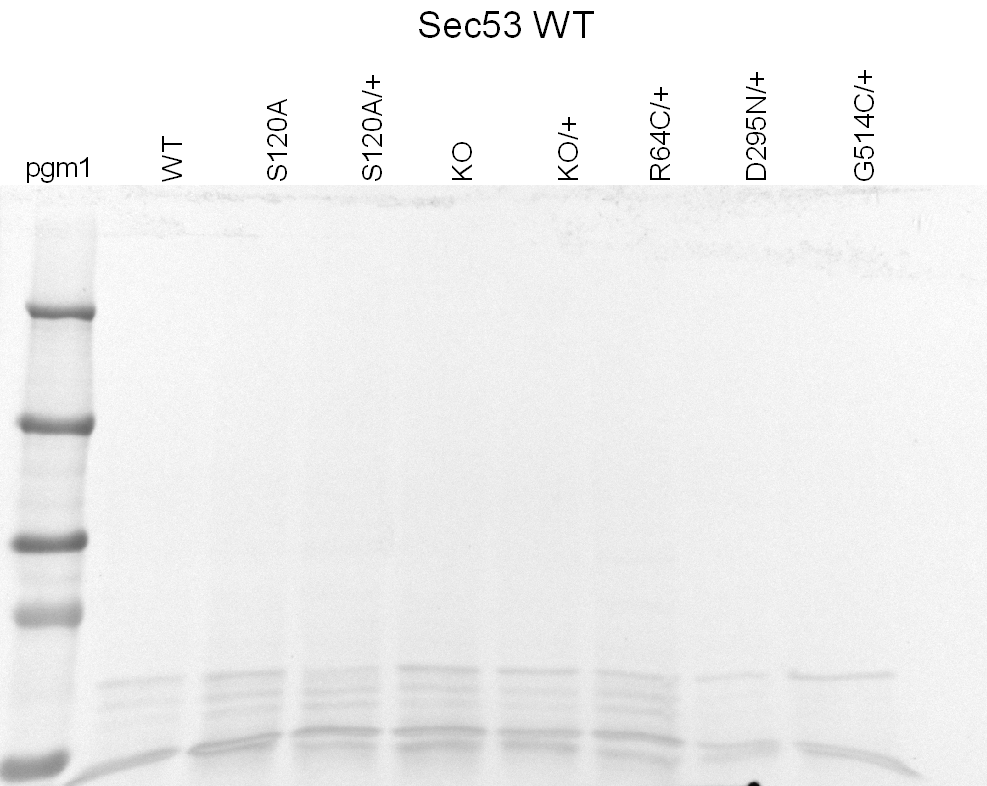

### ponceauD_WT_invertase.tif

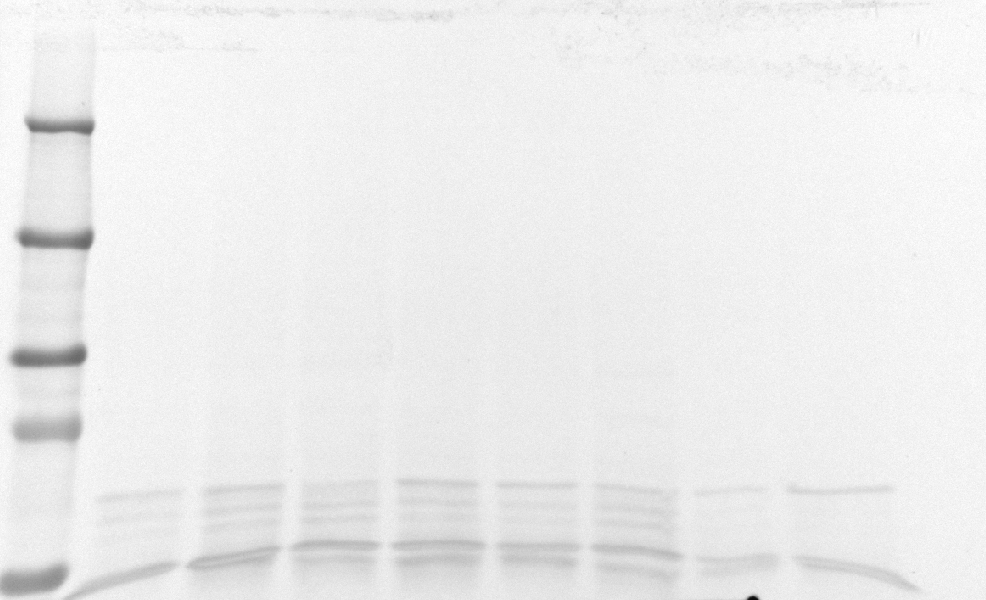

### ponceauE_WT_CPY2.tif

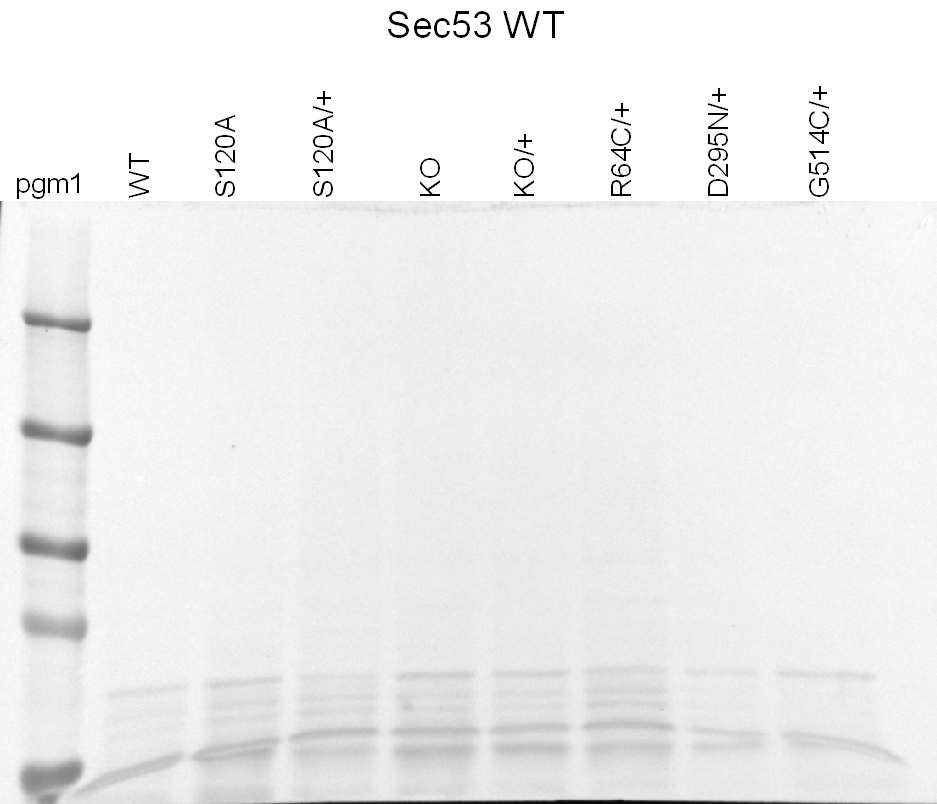

### ponceauE_WT_CPY.tif

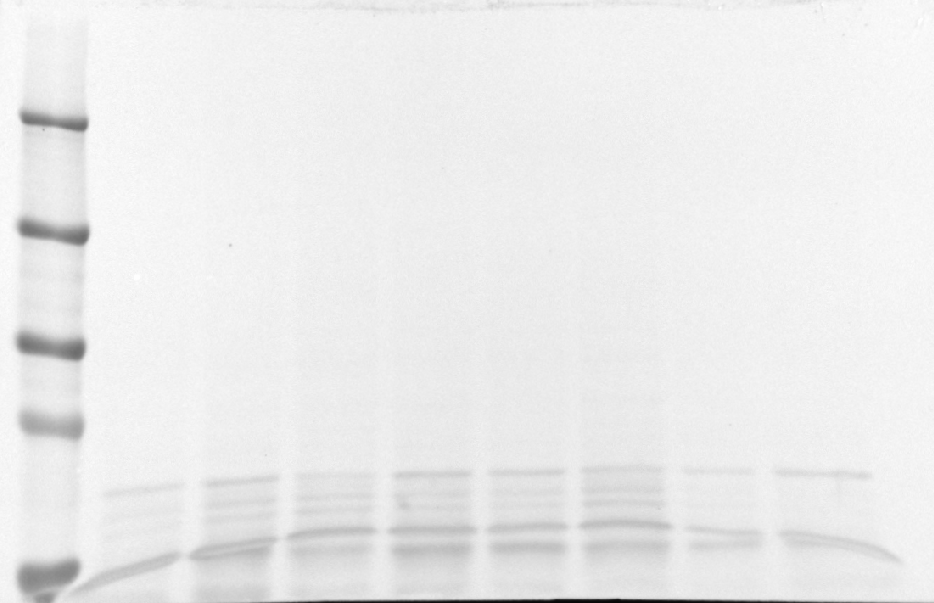

### ponceauF_F126L_invertase2.tif

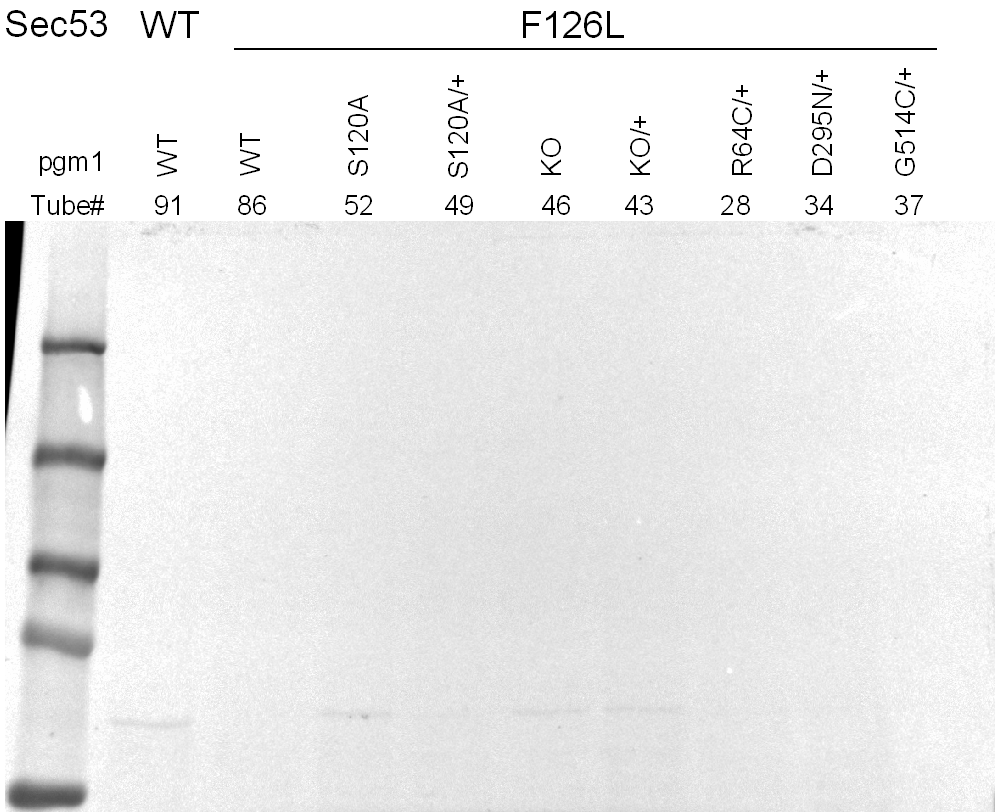

### ponceauF_F126L_invertase.tif

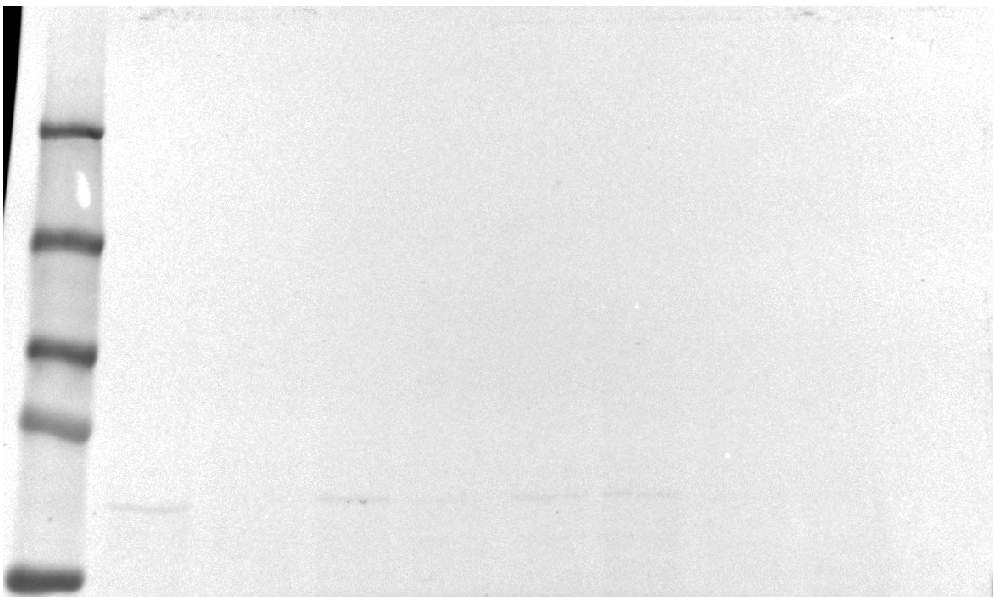
