## Supplementary figures and images for "Experimental evolution of phosphomannomutase-deficient yeast reveals compensatory mutations in a phosphoglucomutase"

### 2x_V238M_01.png

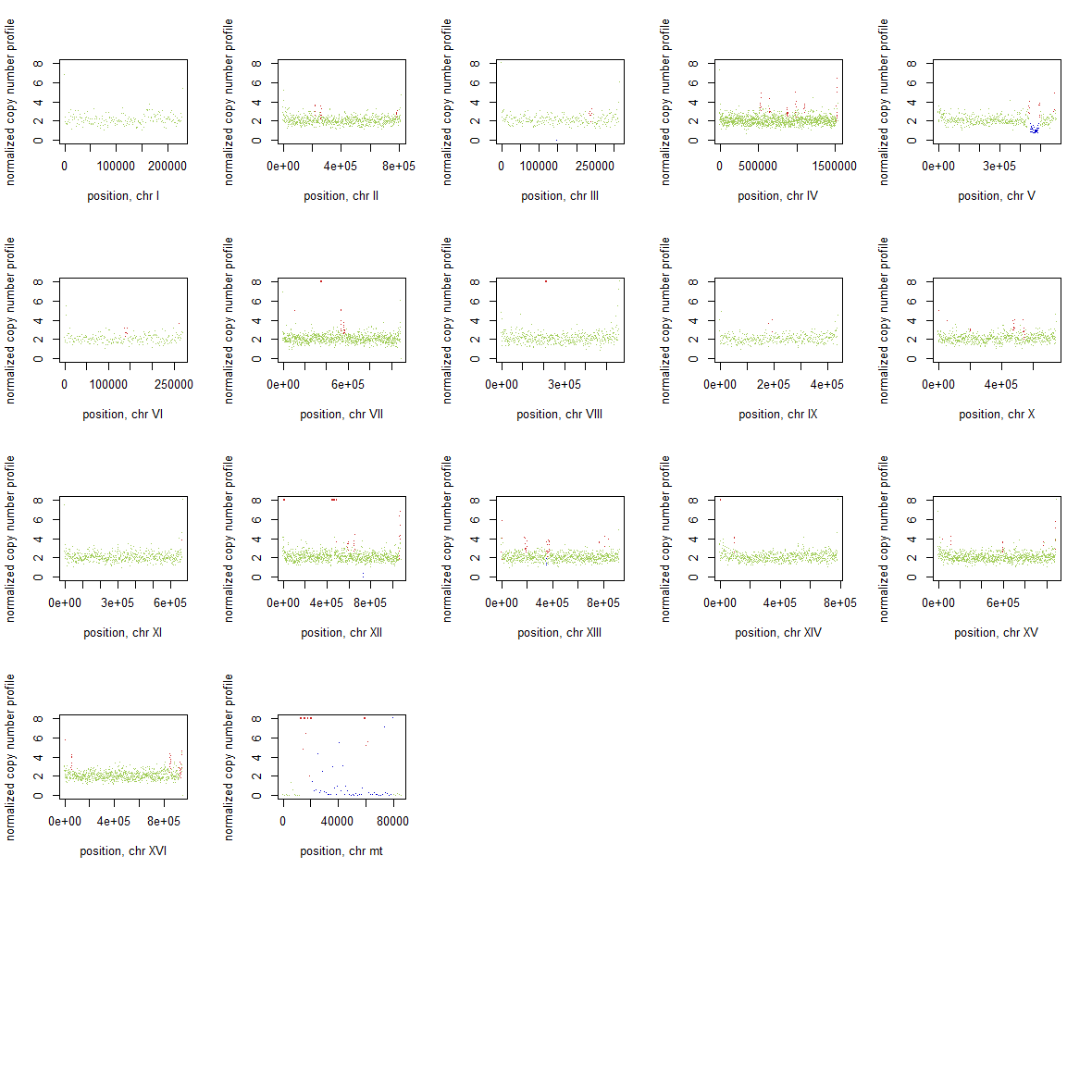

### 2x_V238M_02.png

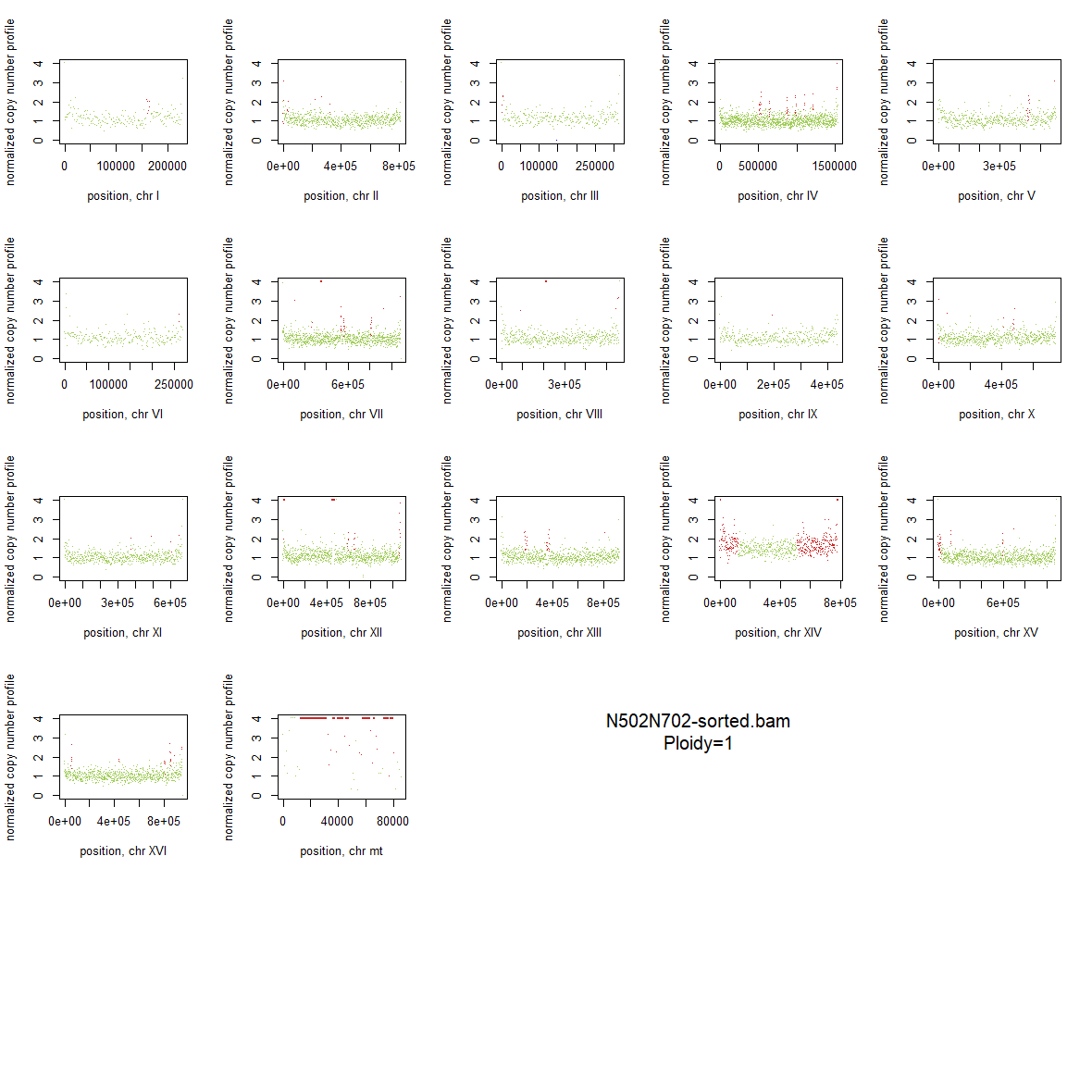

### 2x_V238M_03.png

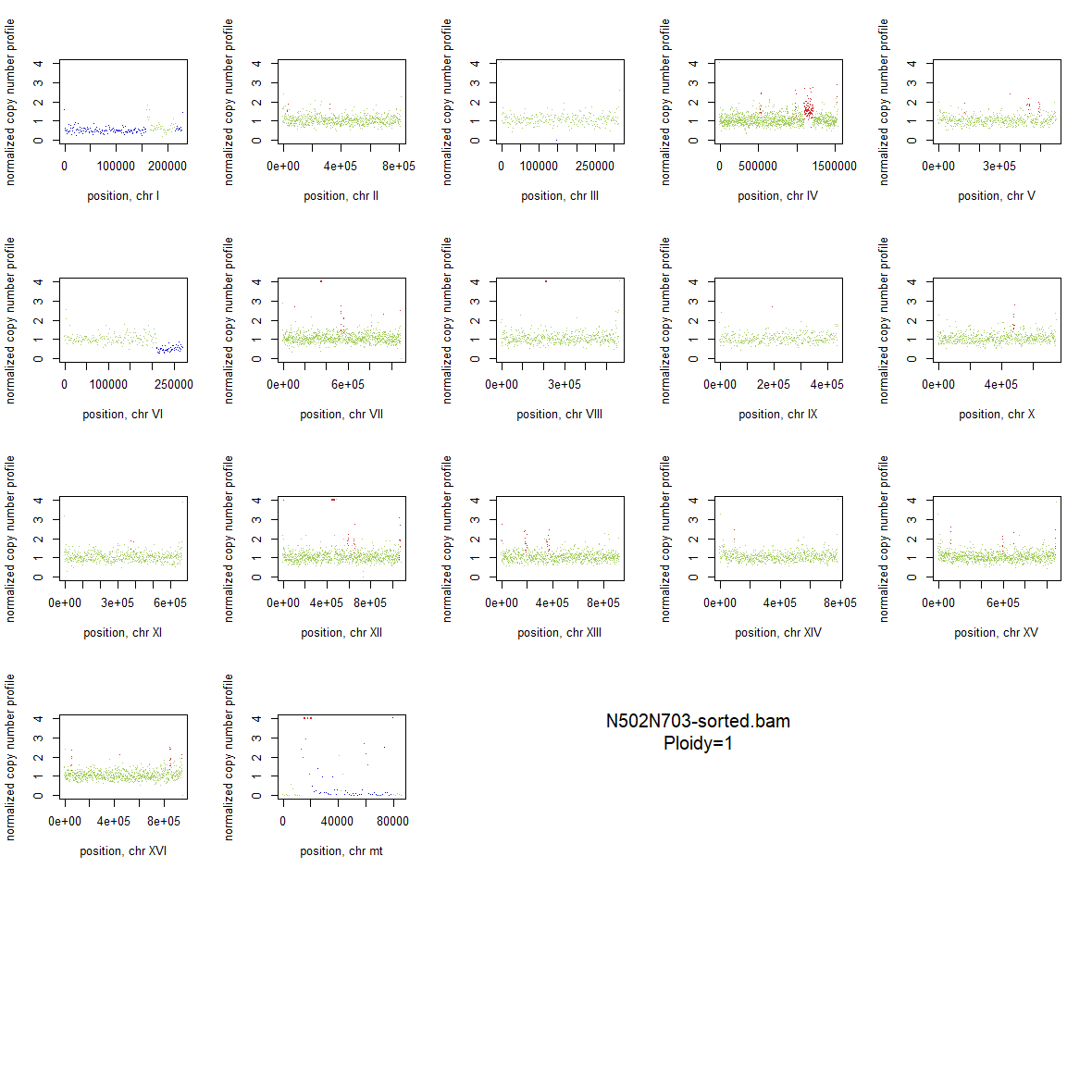

### 2x_V238M_04.png

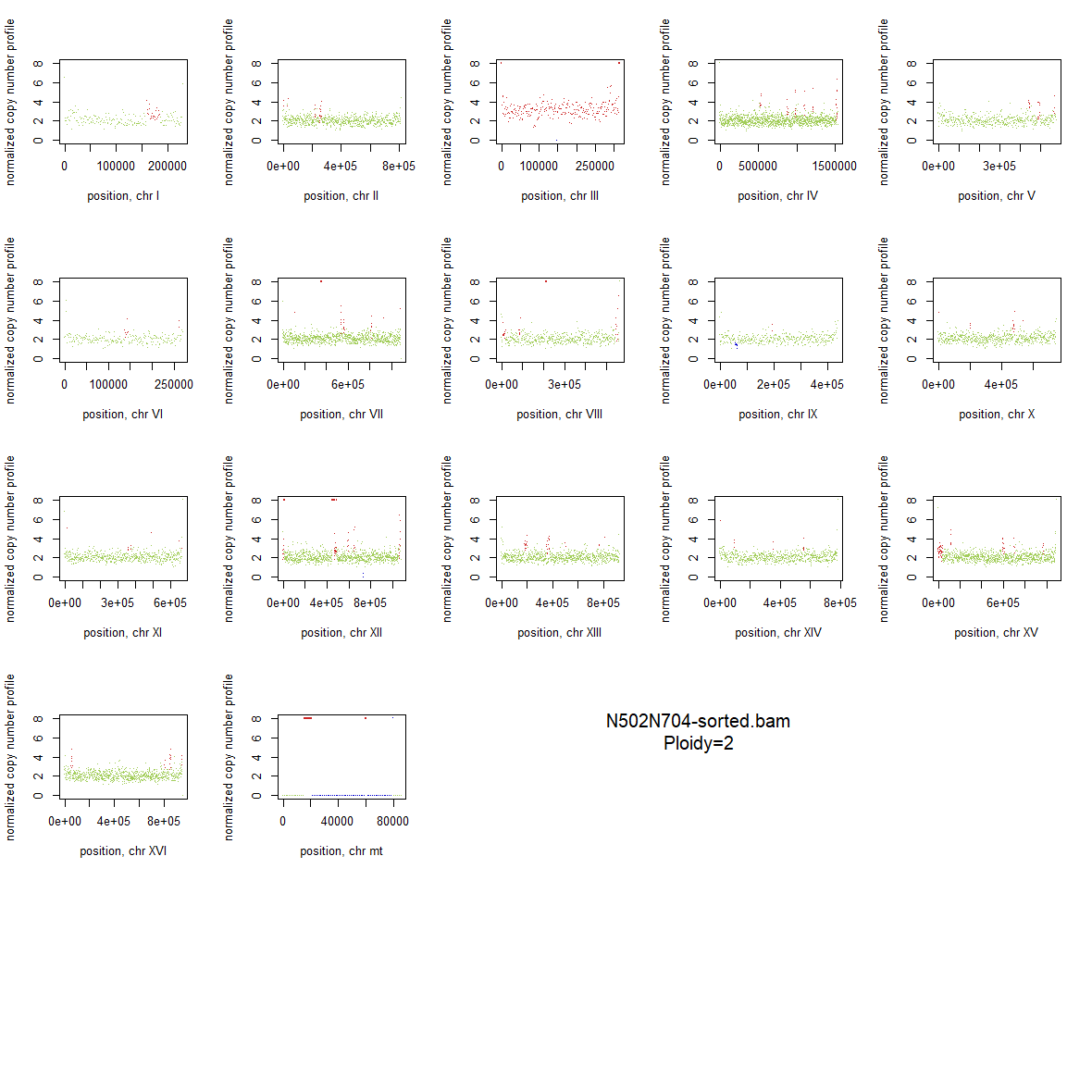

### 2x_V238M_05.png

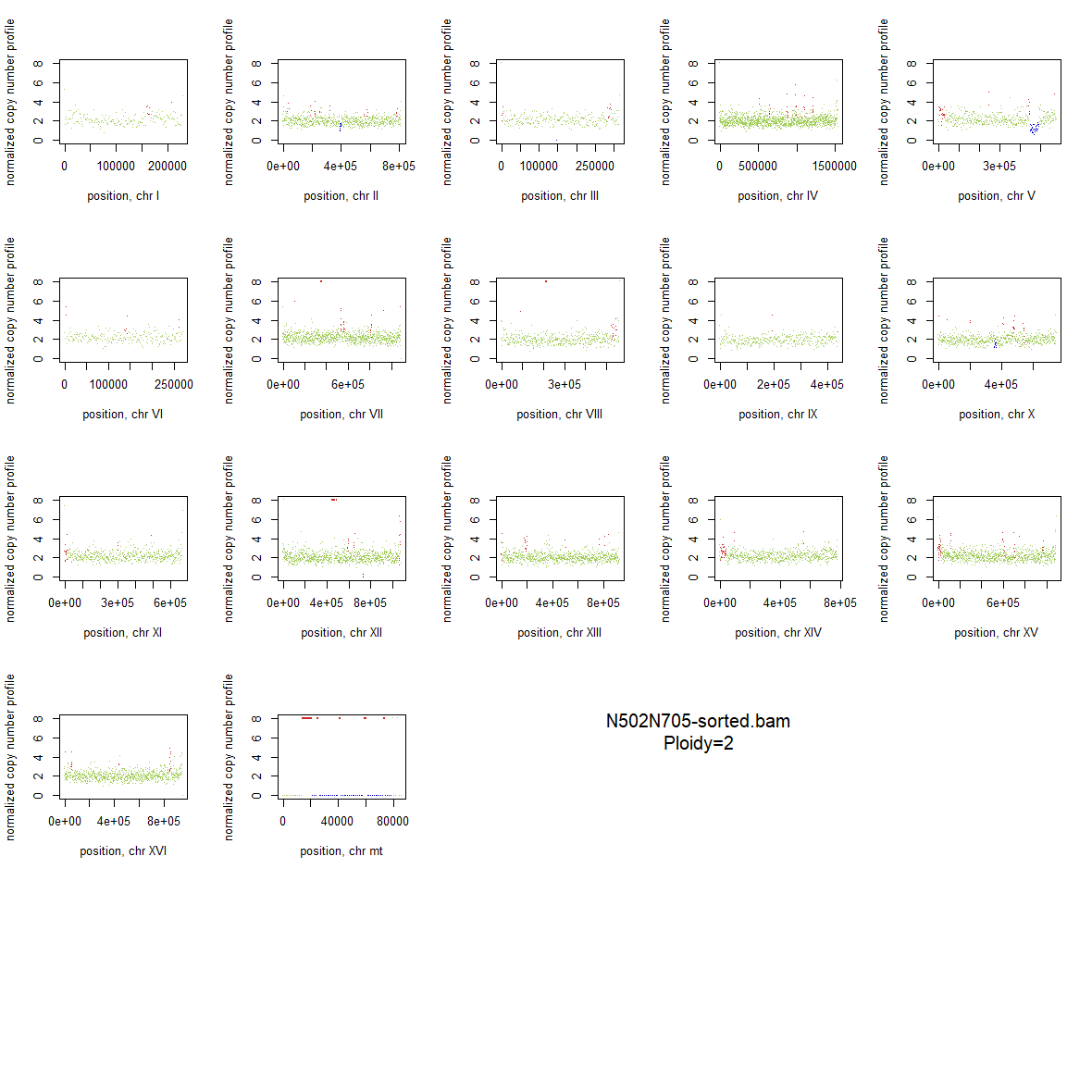

### 2x_V238M_06.png

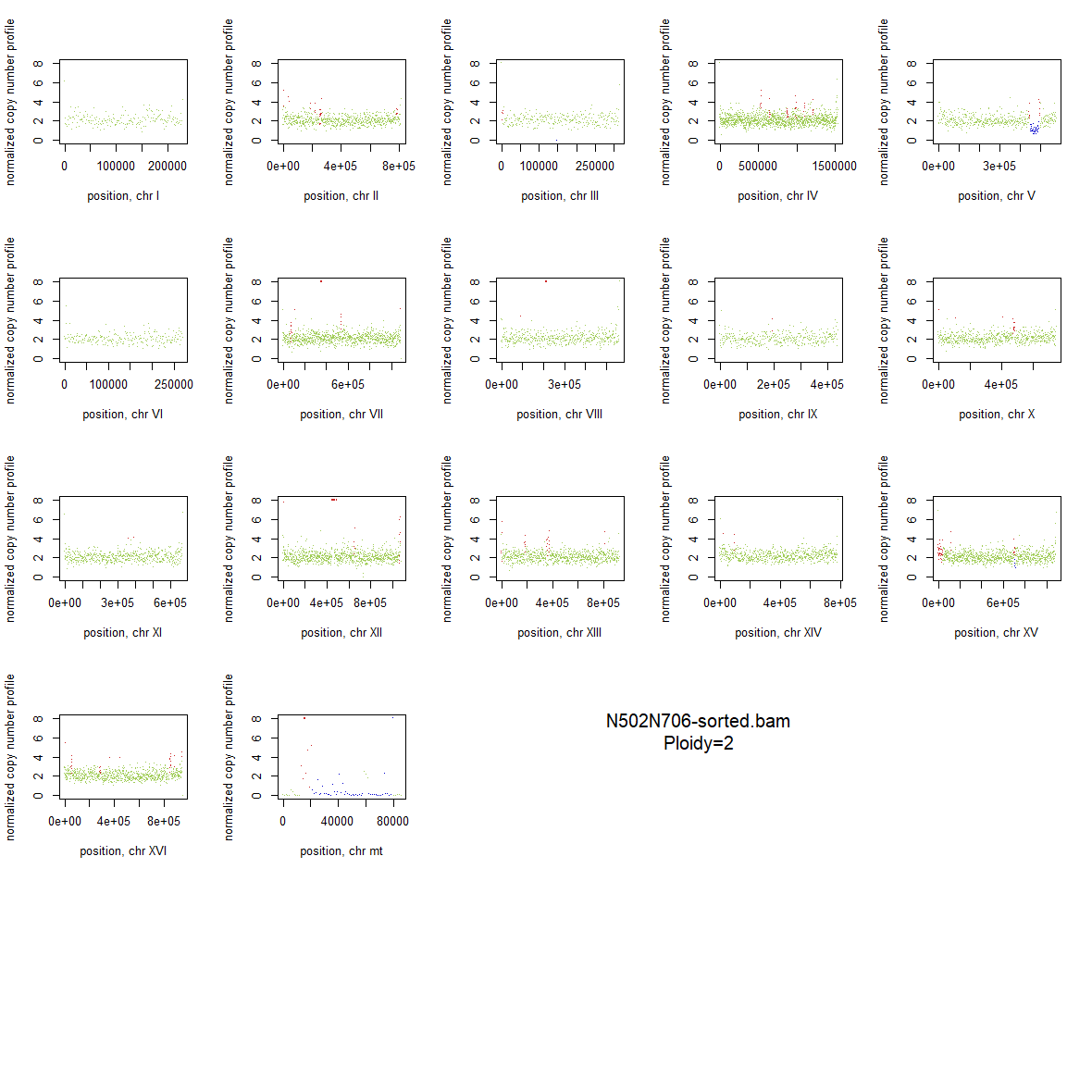

### 2x_V238M_07.png

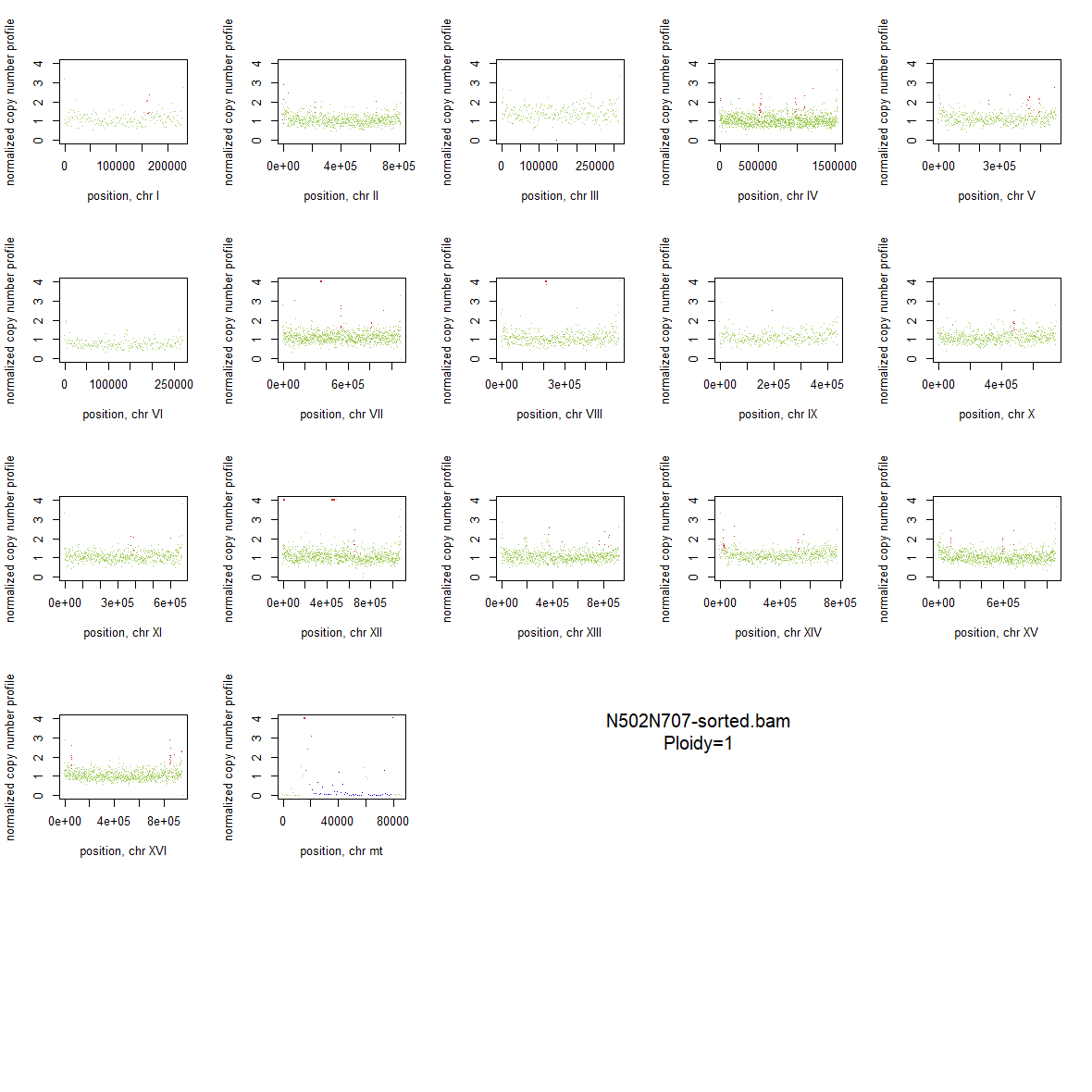

### 2x_V238M_08.png

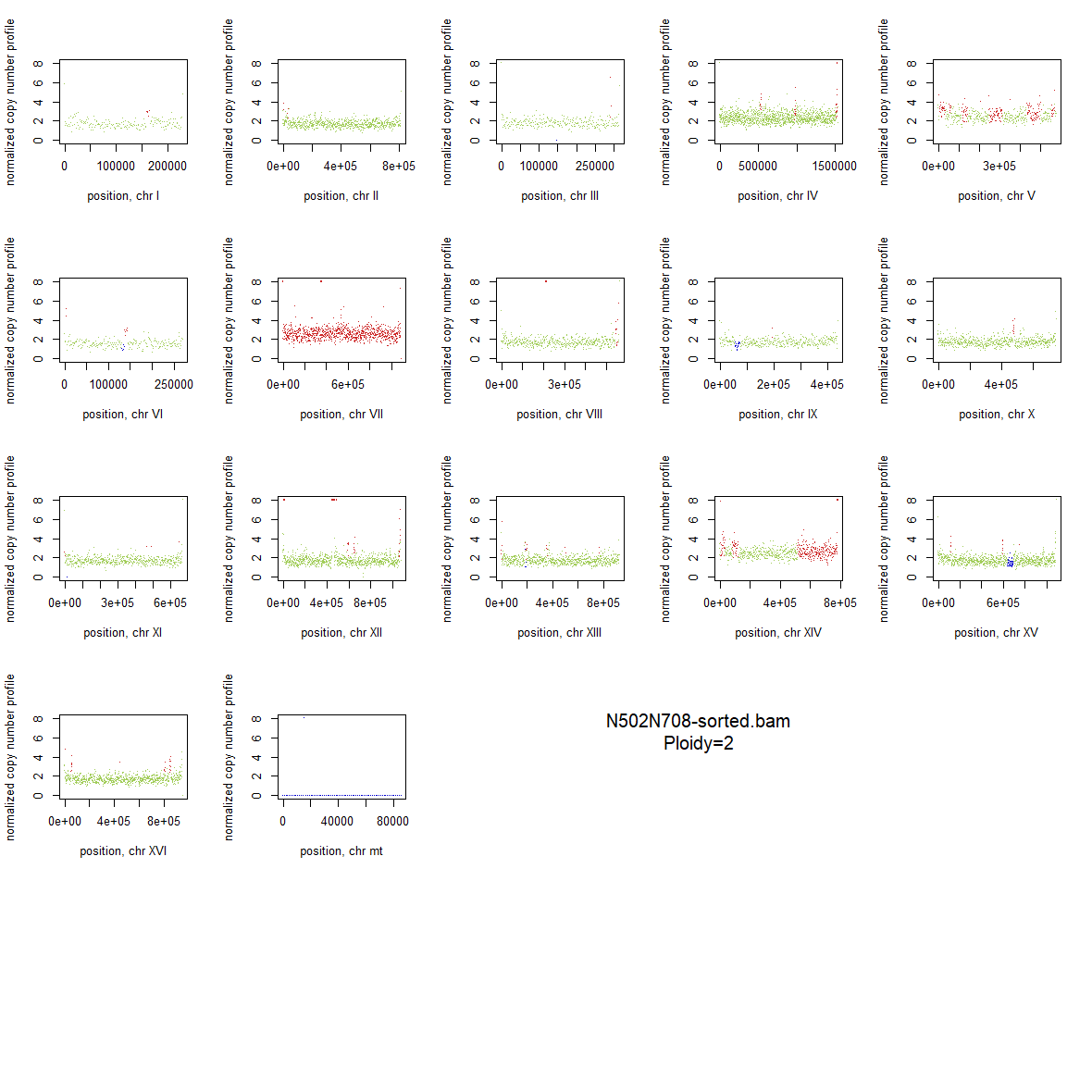

### 2x_V238M_09.png

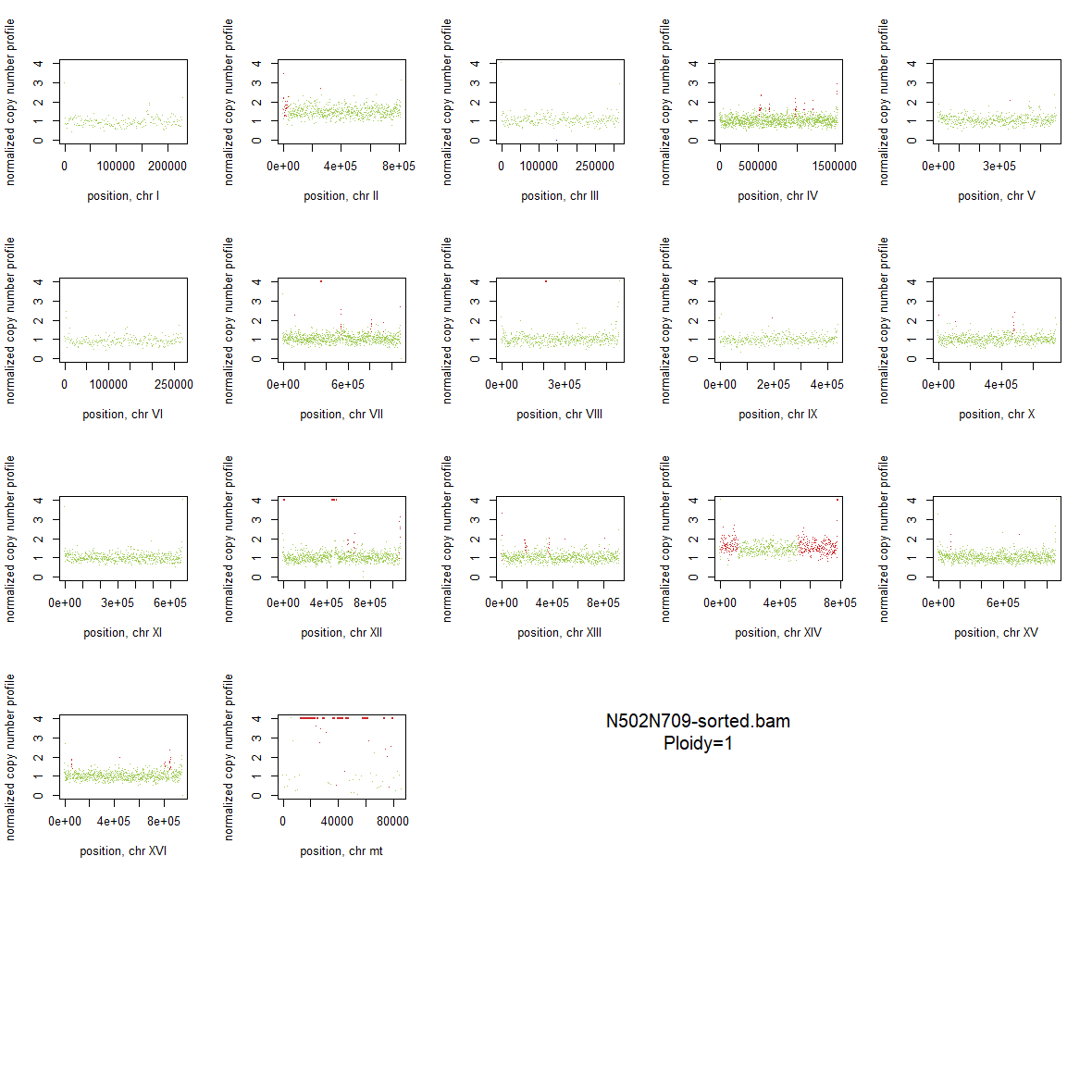

### 2x_V238M_10.png

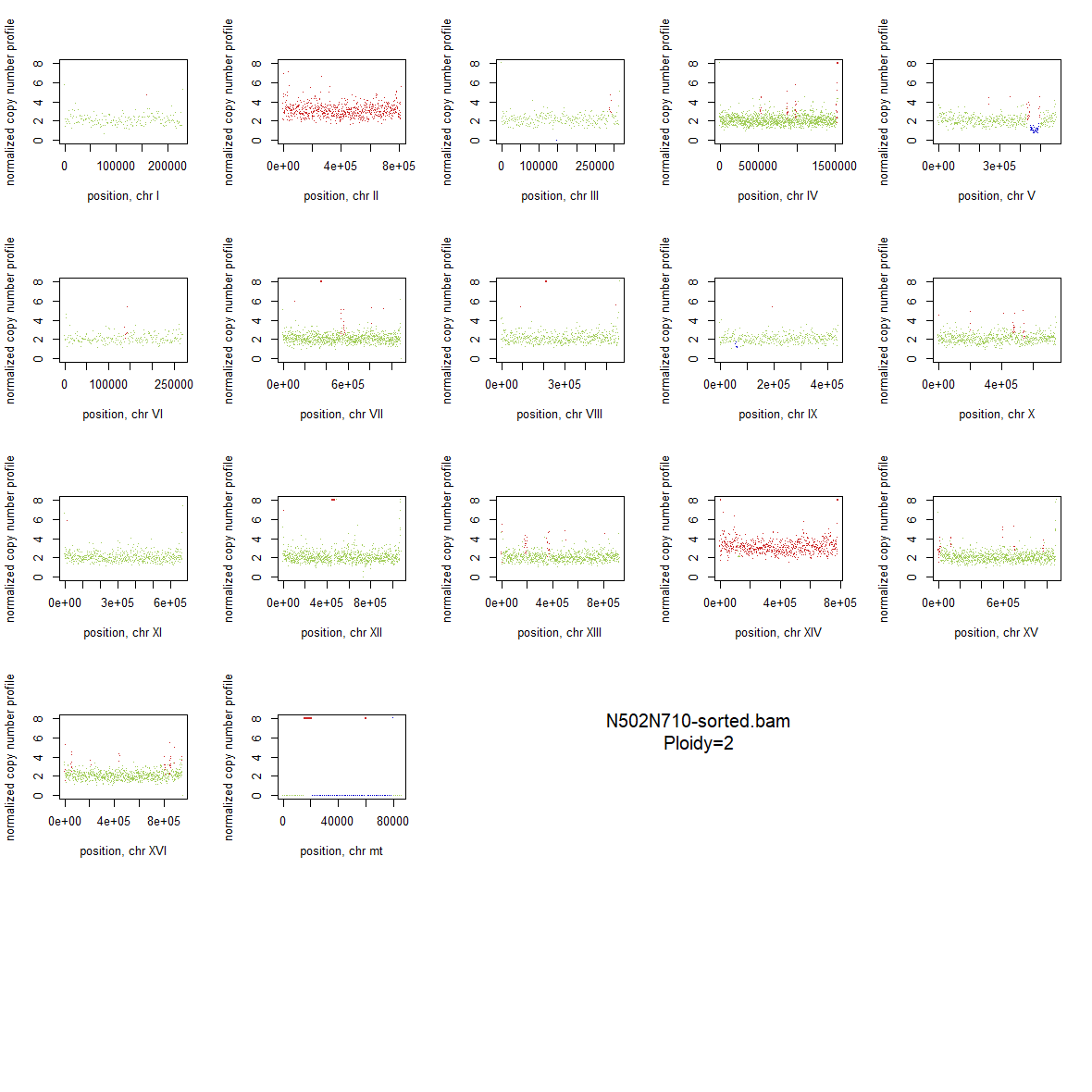

### 2x_V238M_11.png

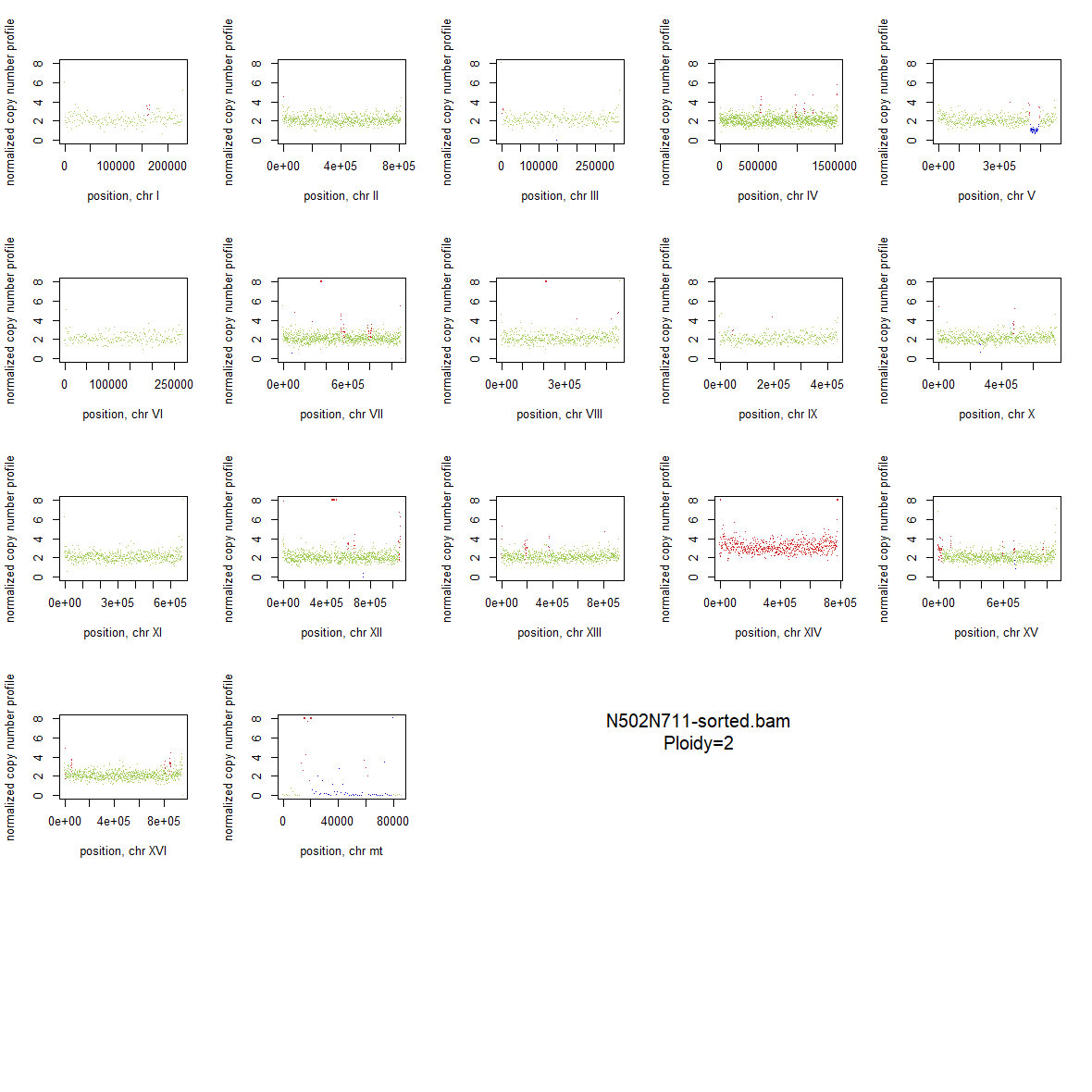

### 2x_V238M_12.png

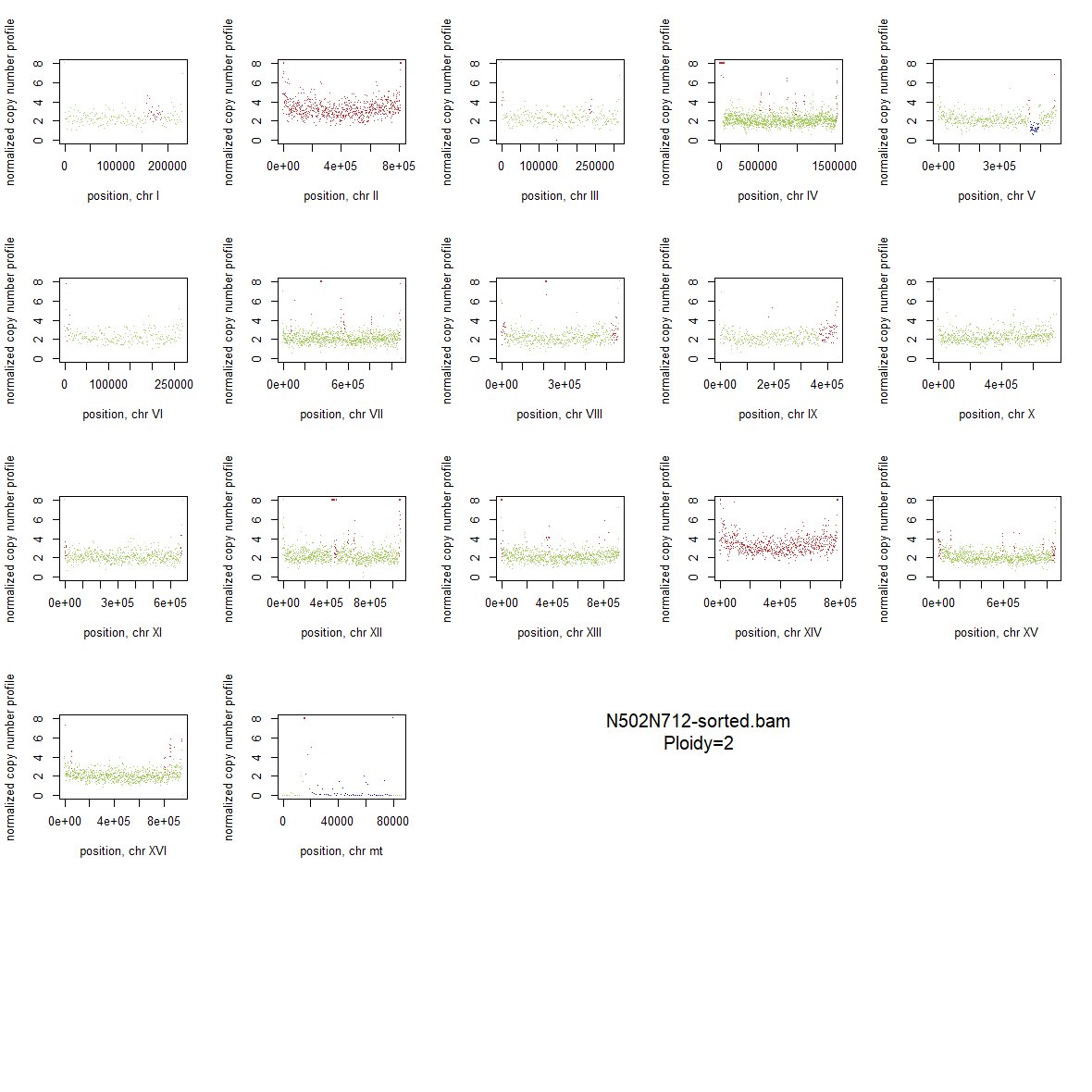

### 2x_V238M_13.png

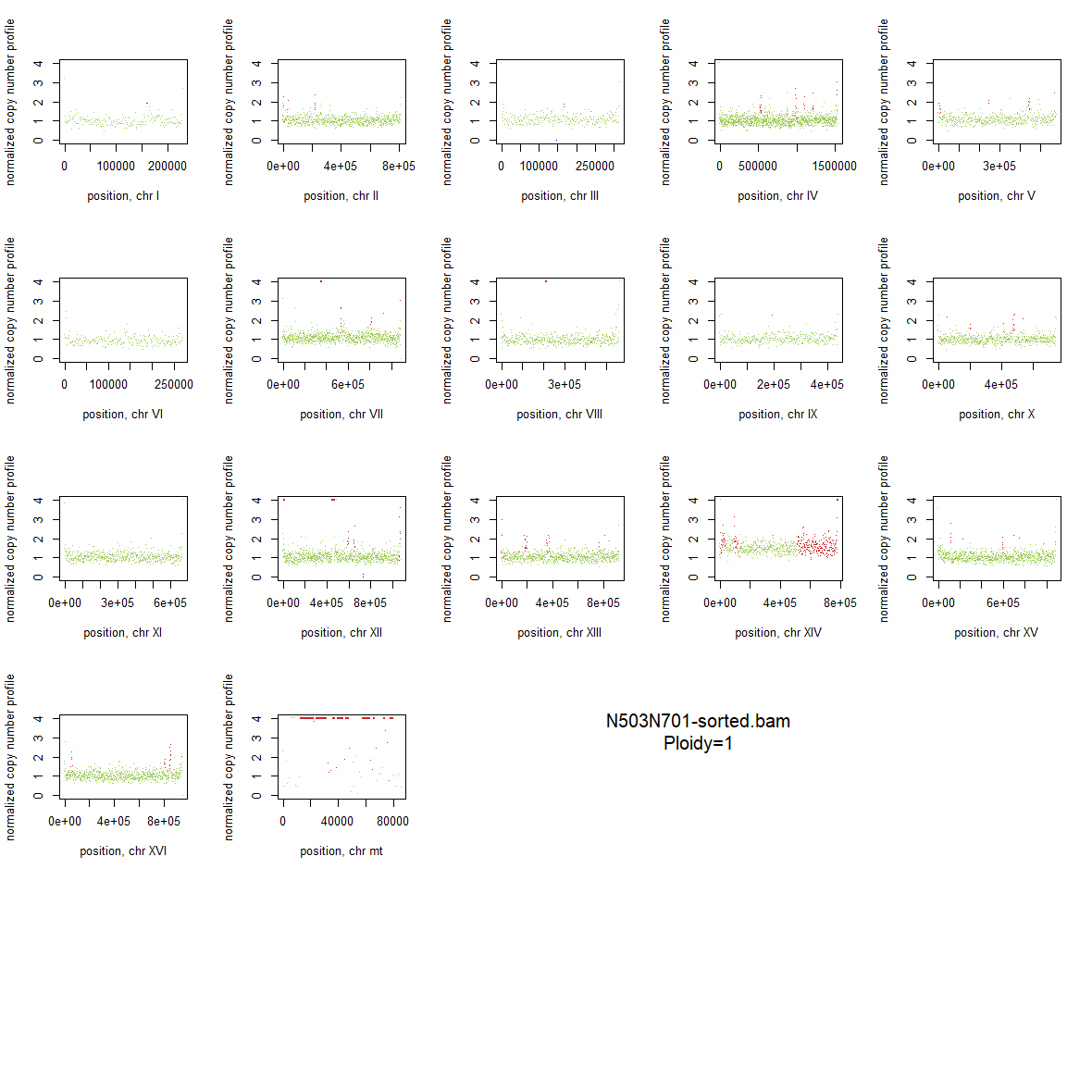

### 2x_V238M_14.png

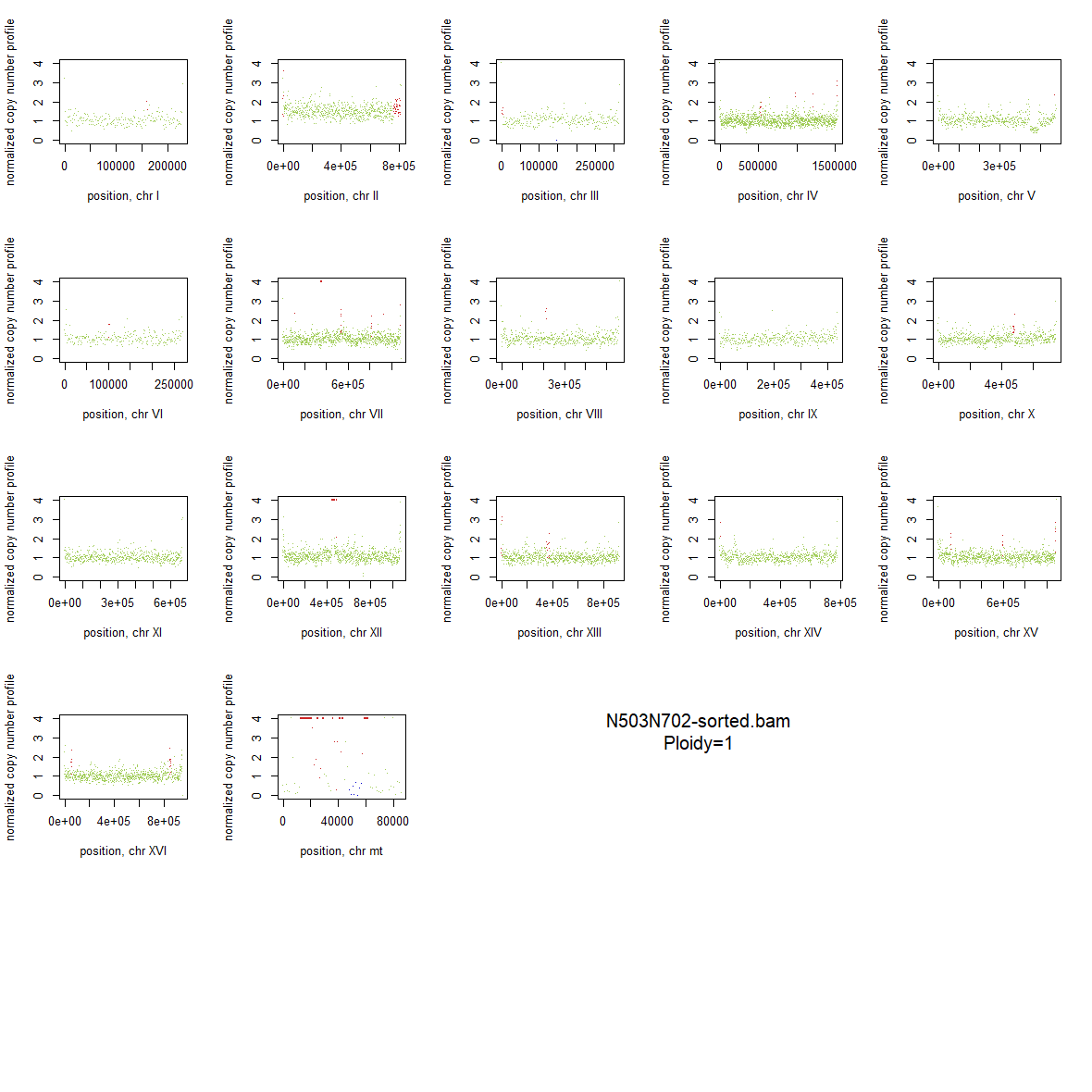

### 2x_V238M_15.png

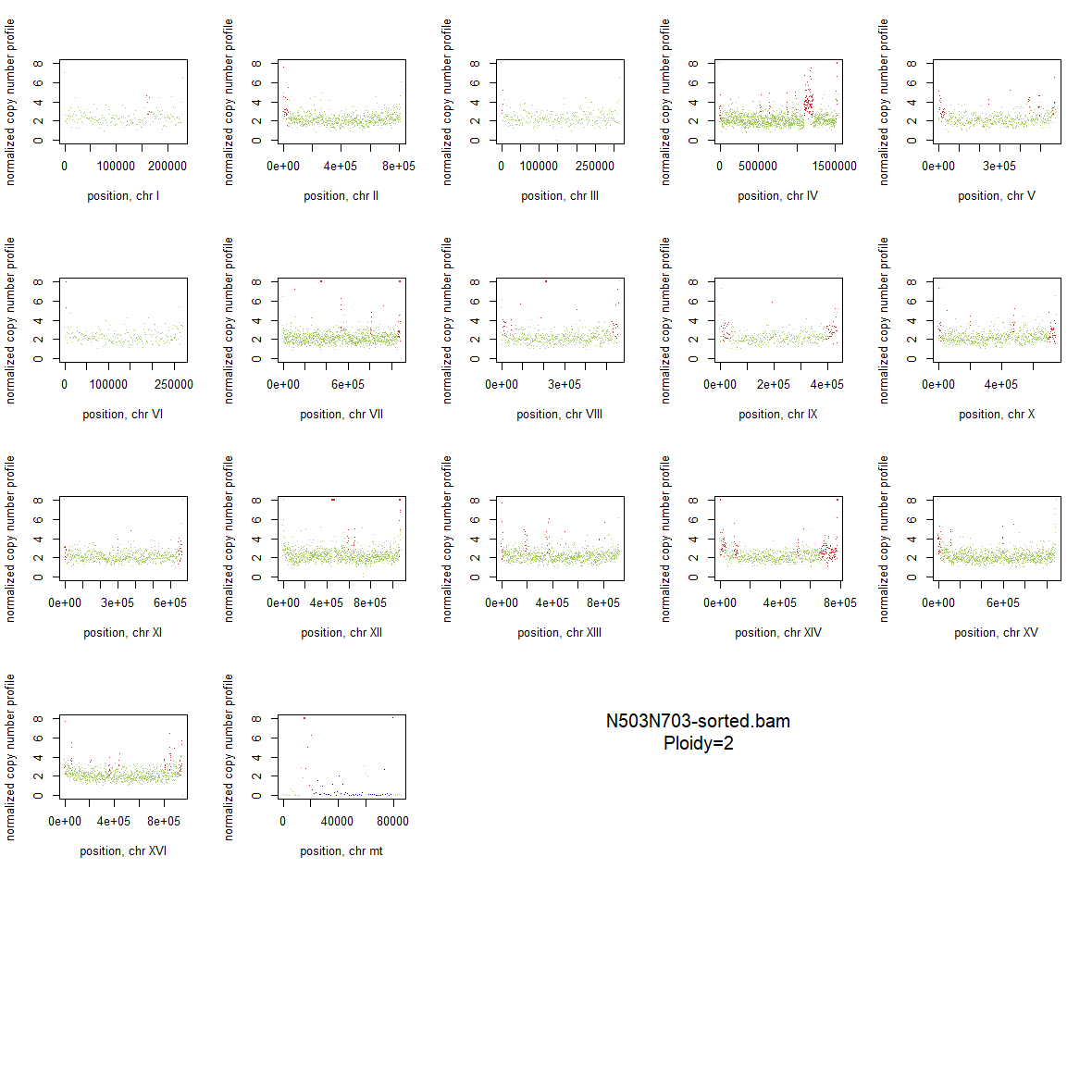

### 2x_V238M_16.png

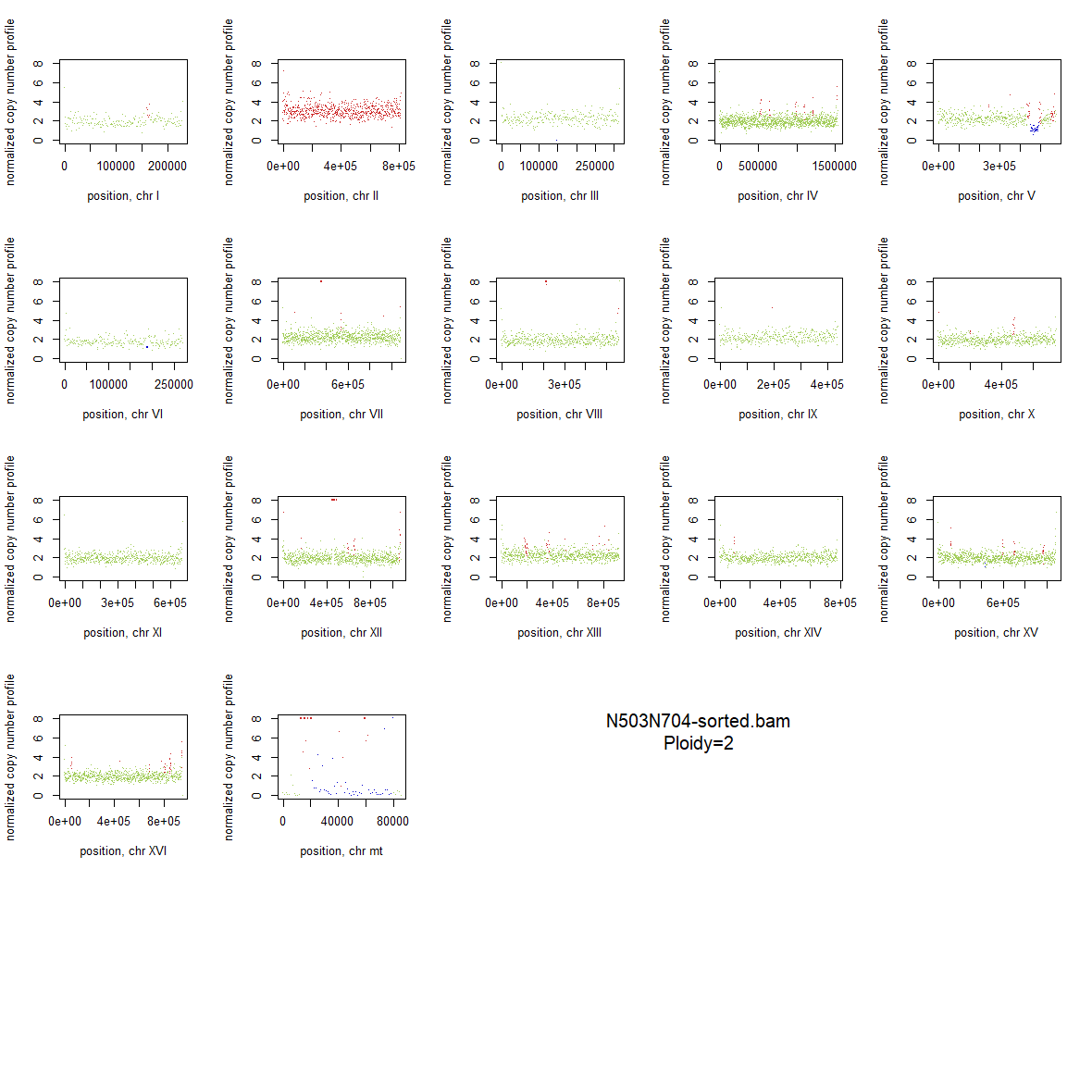

### 2x_V238M_17.png

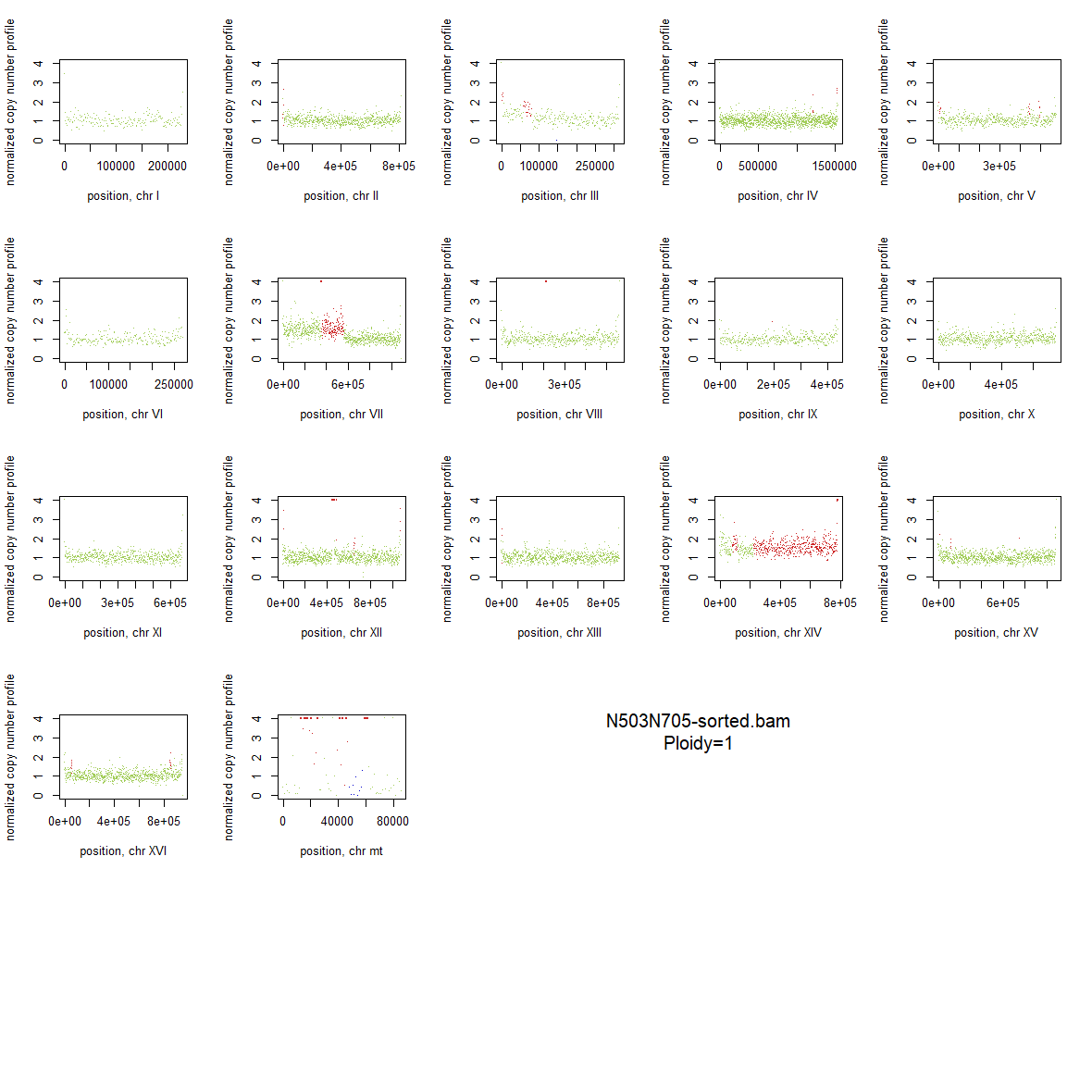

### 2x_V238M_18.png

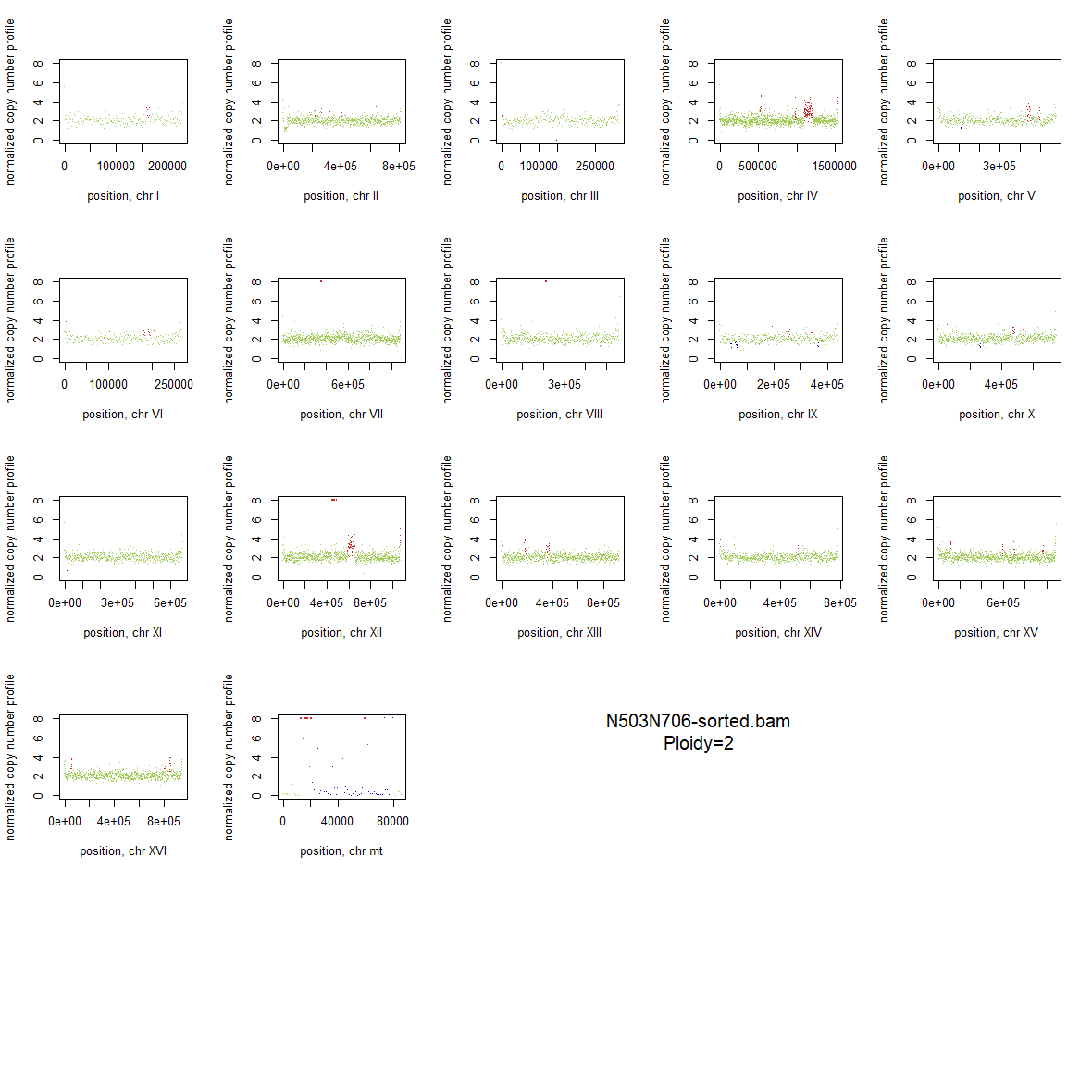

### 2x_V238M_19.png

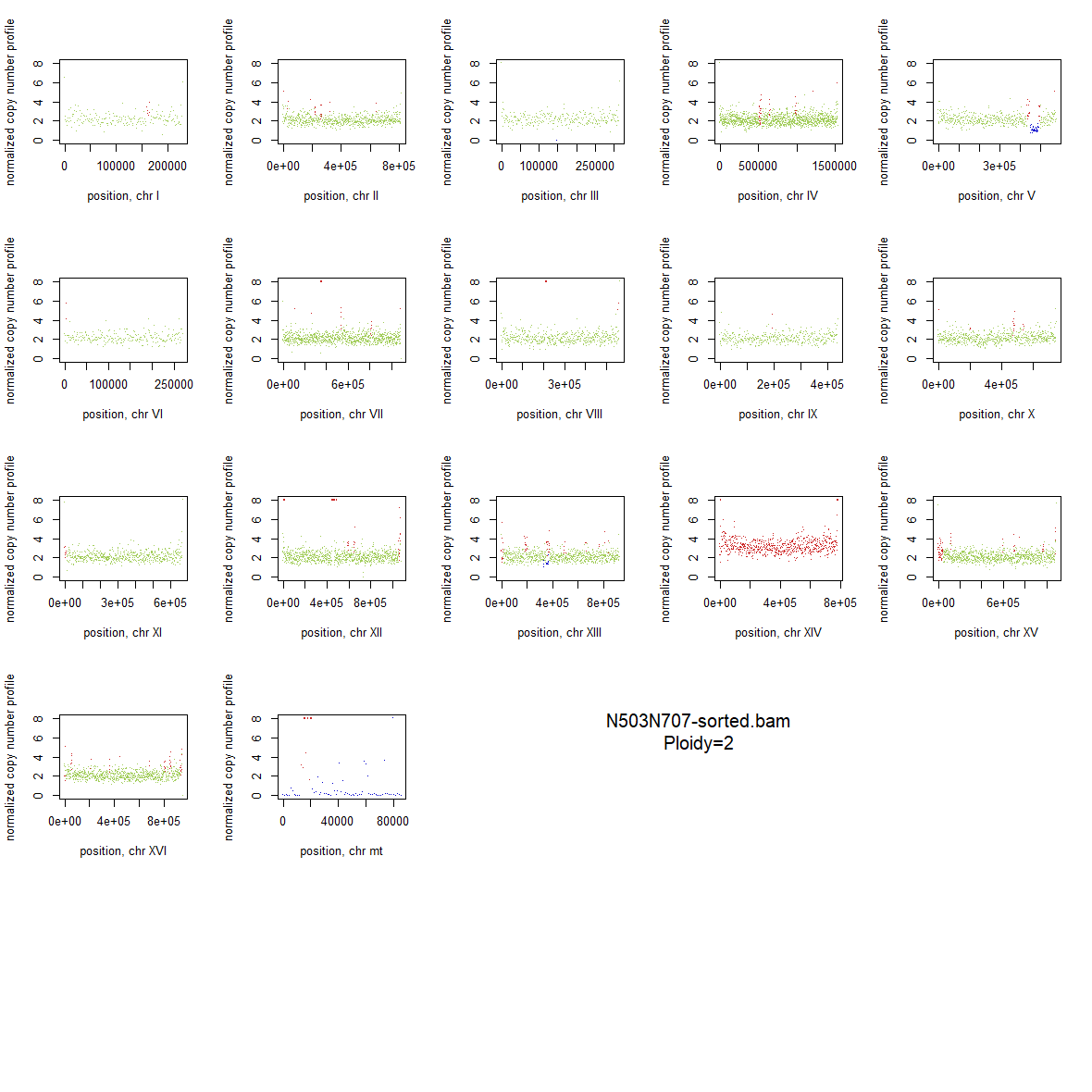

### 2x_V238M_20.png

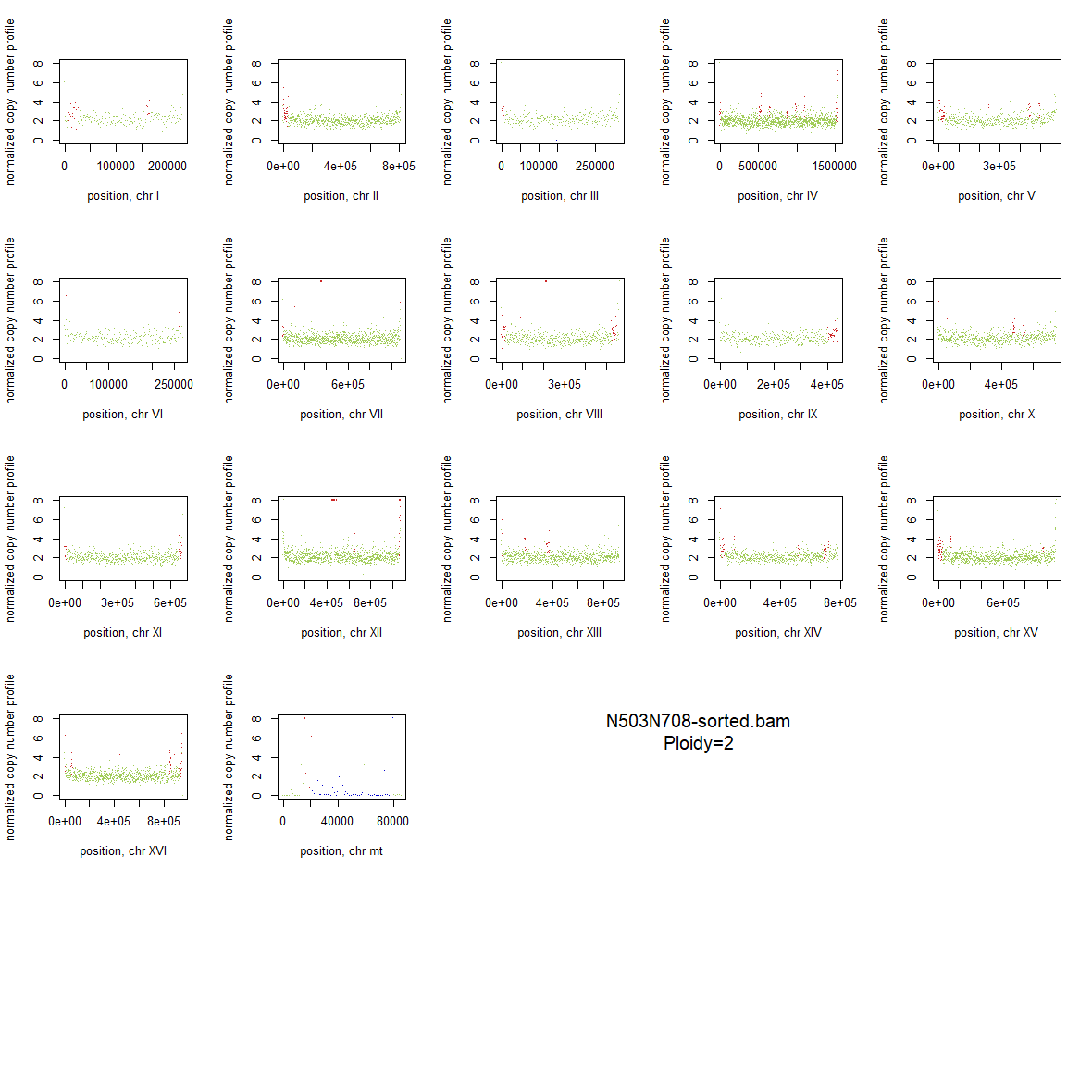

### 2x_V238M_21.png

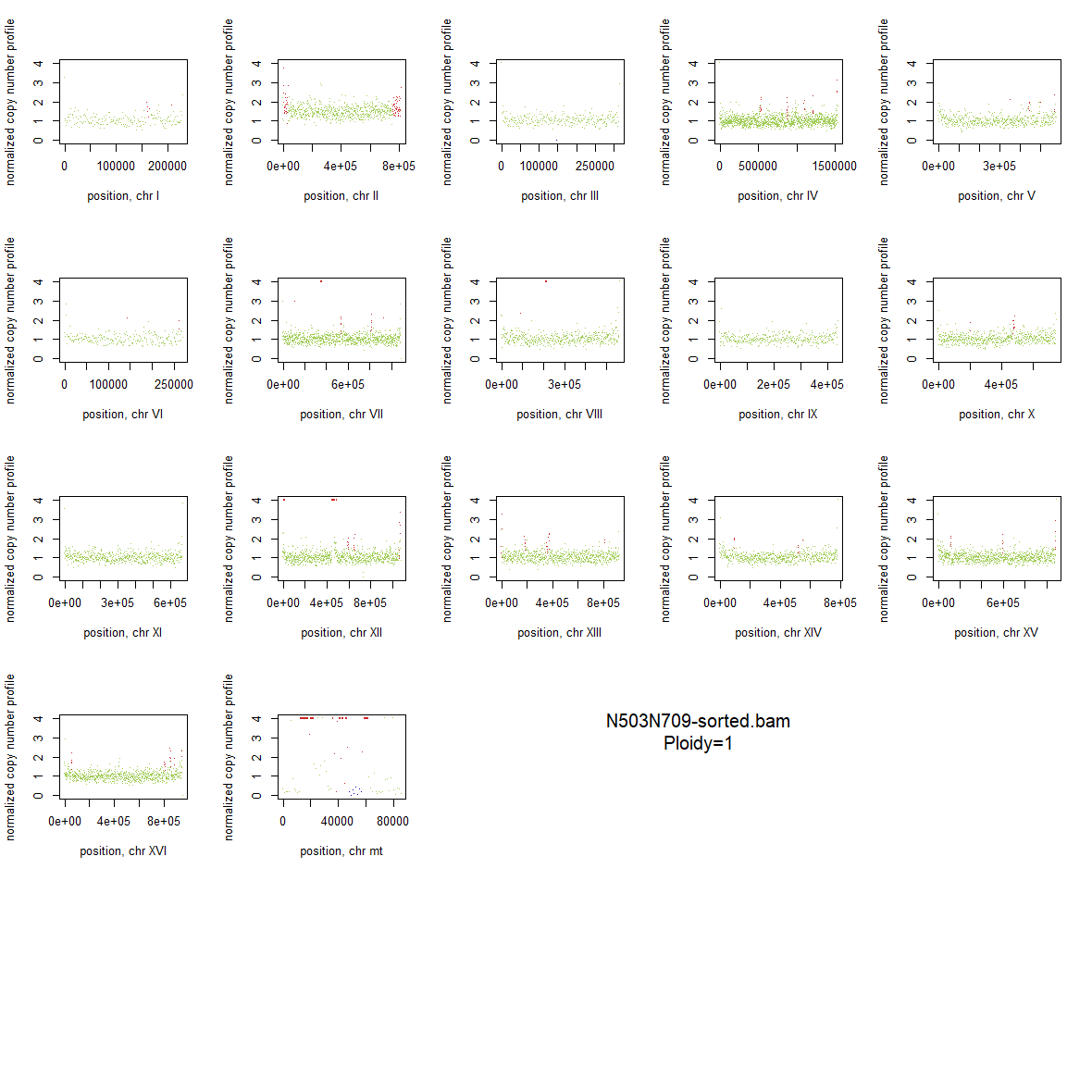

### 2x_V238M_22.png

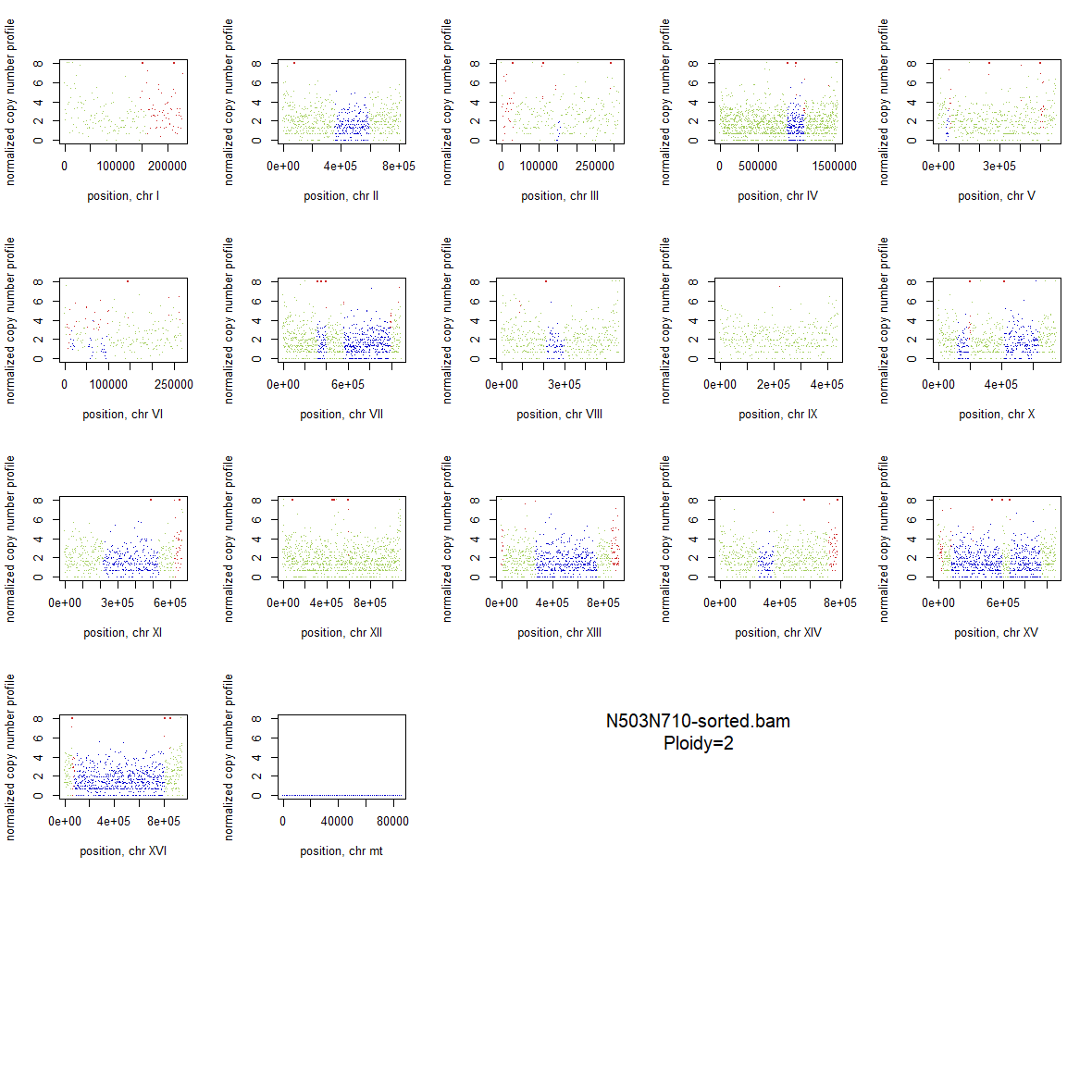

### 2x_V238M_23.png

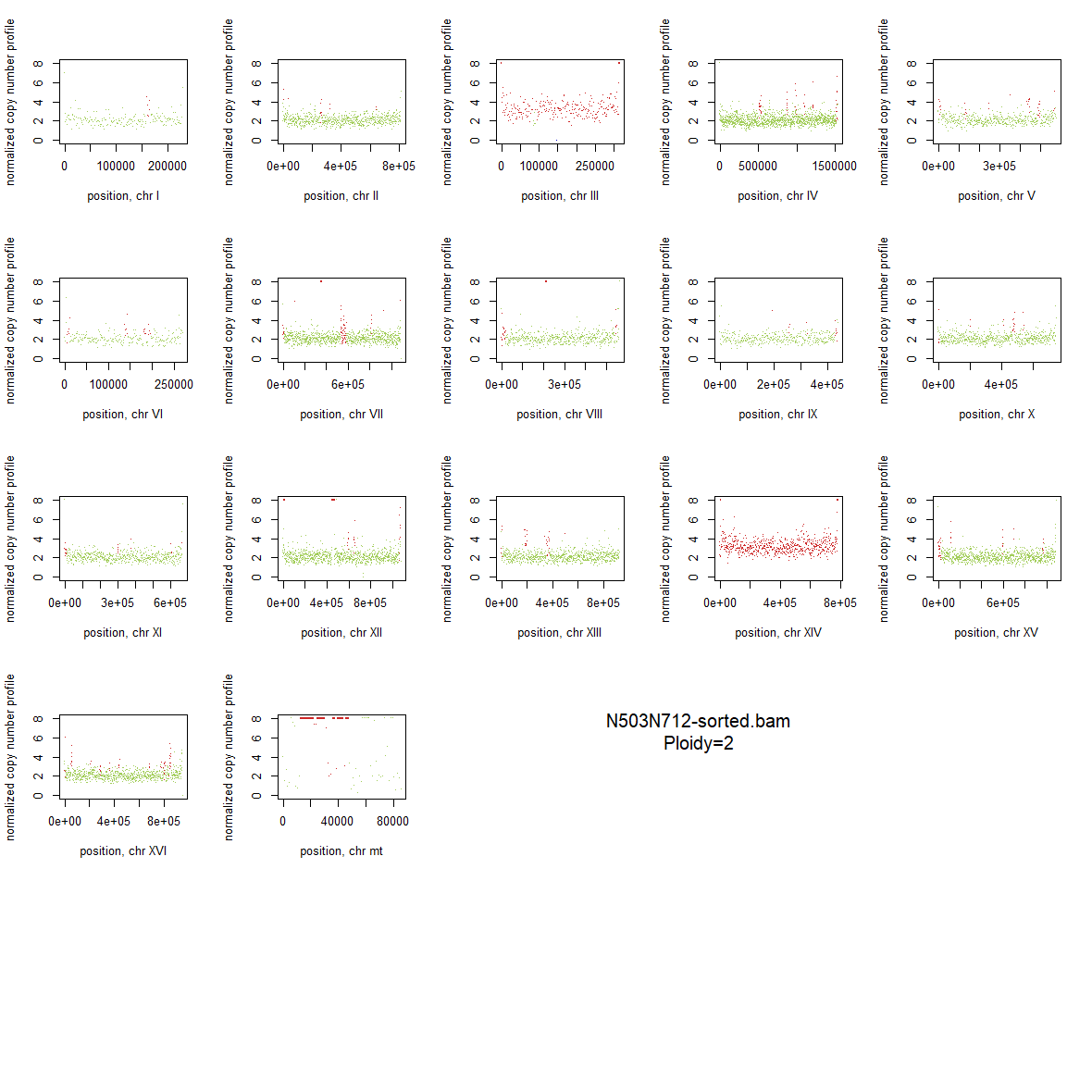

### 2x_V238M_24.png

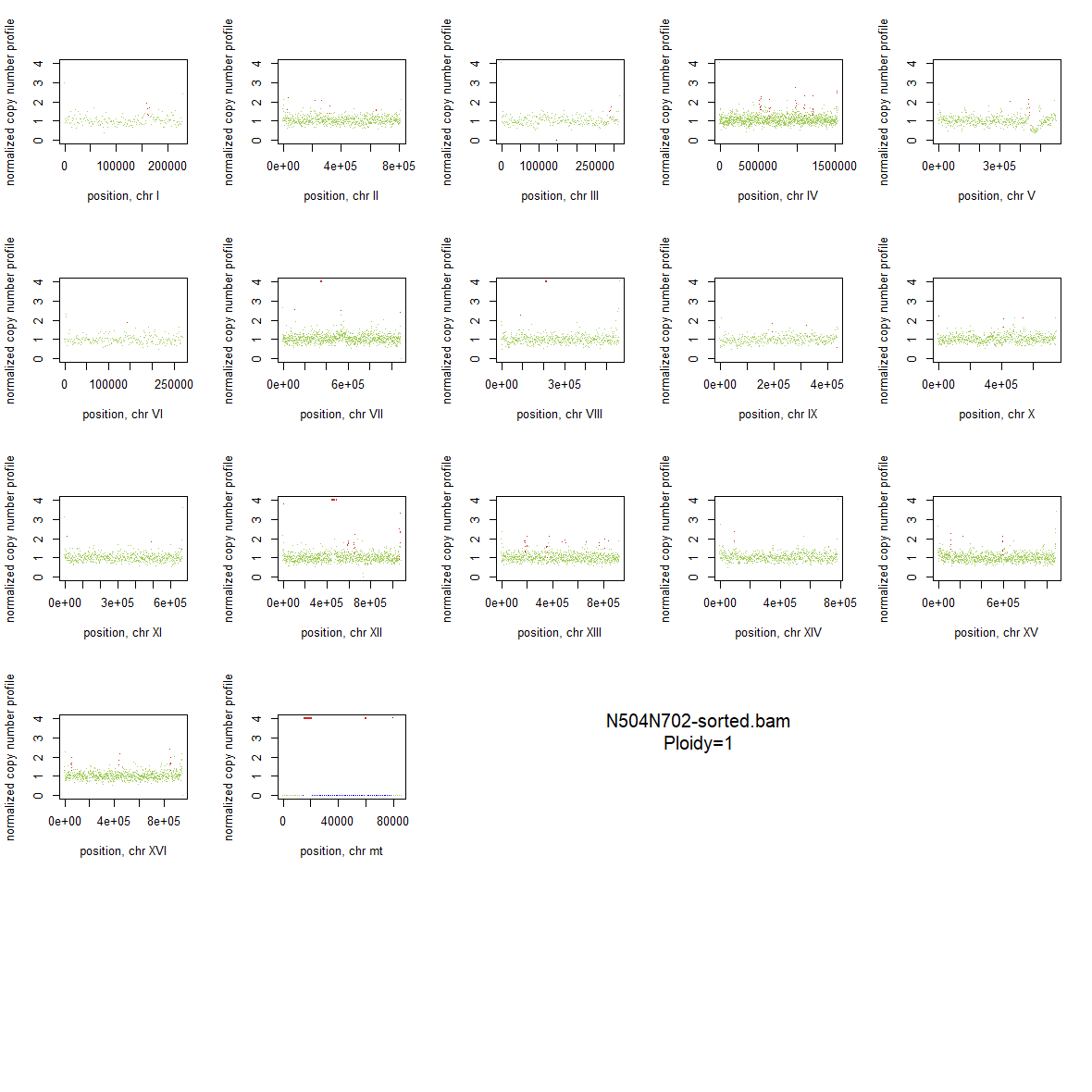

### 2x_V238M_25.png

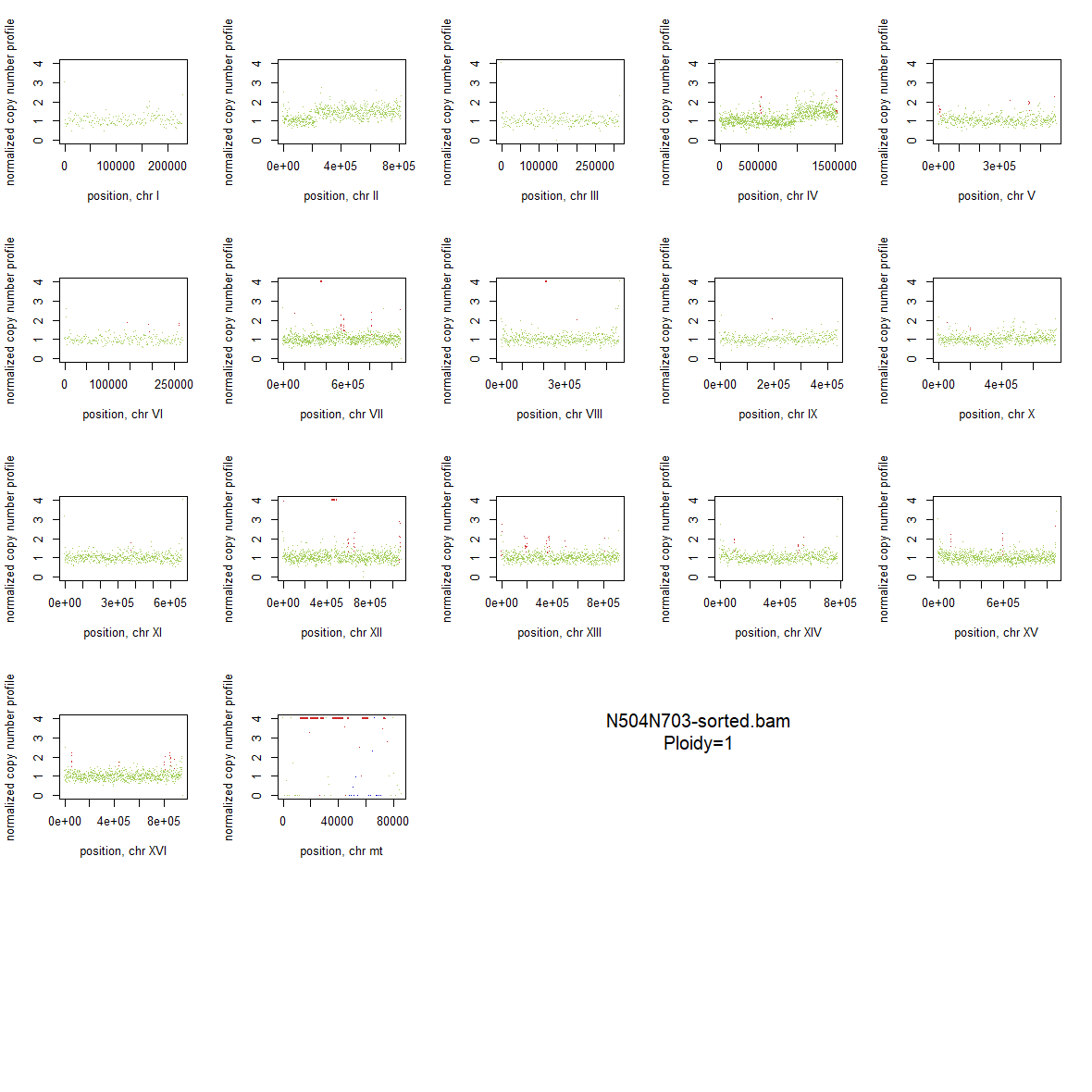

### 2x_V238M_26.png

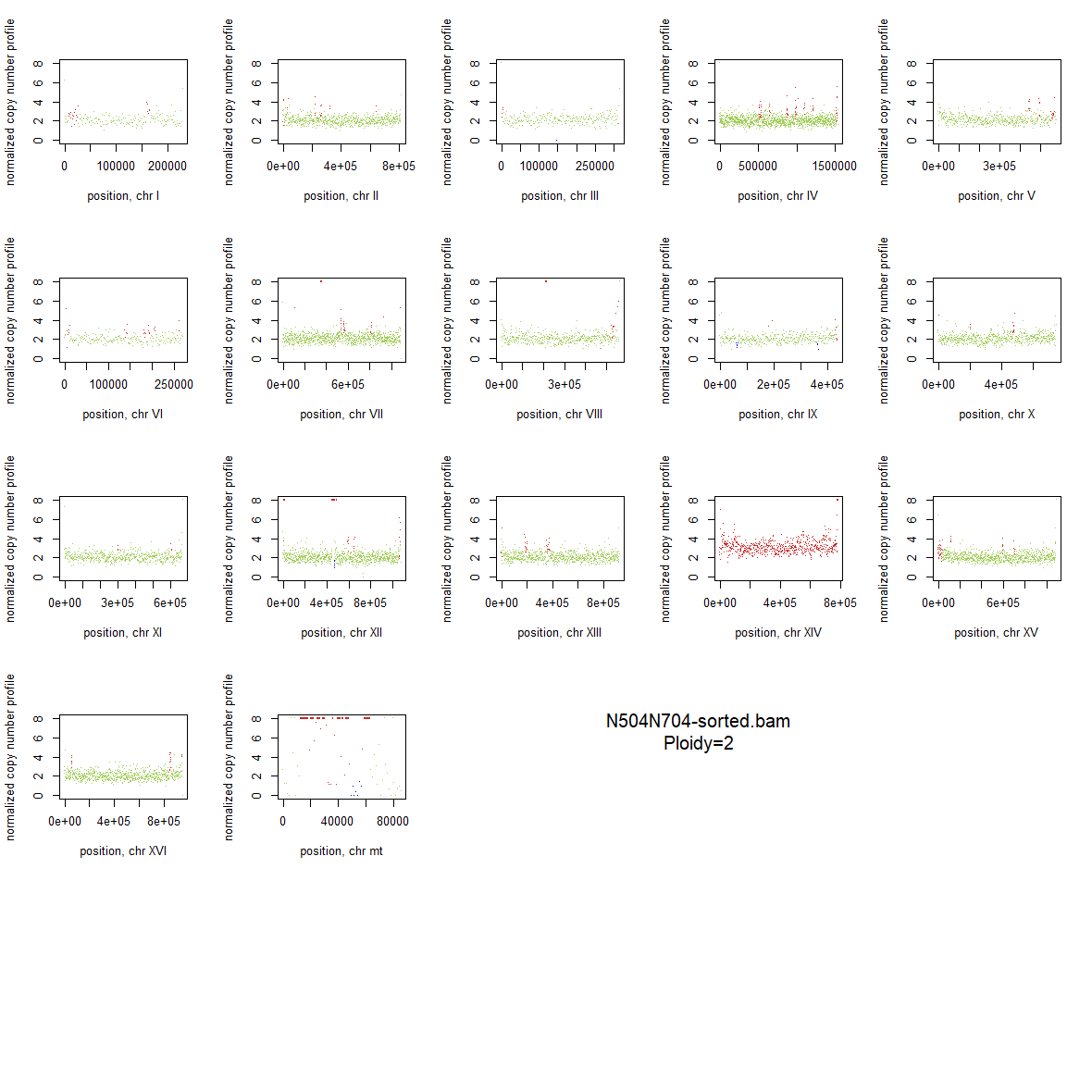

### 2x_V238M_27.png

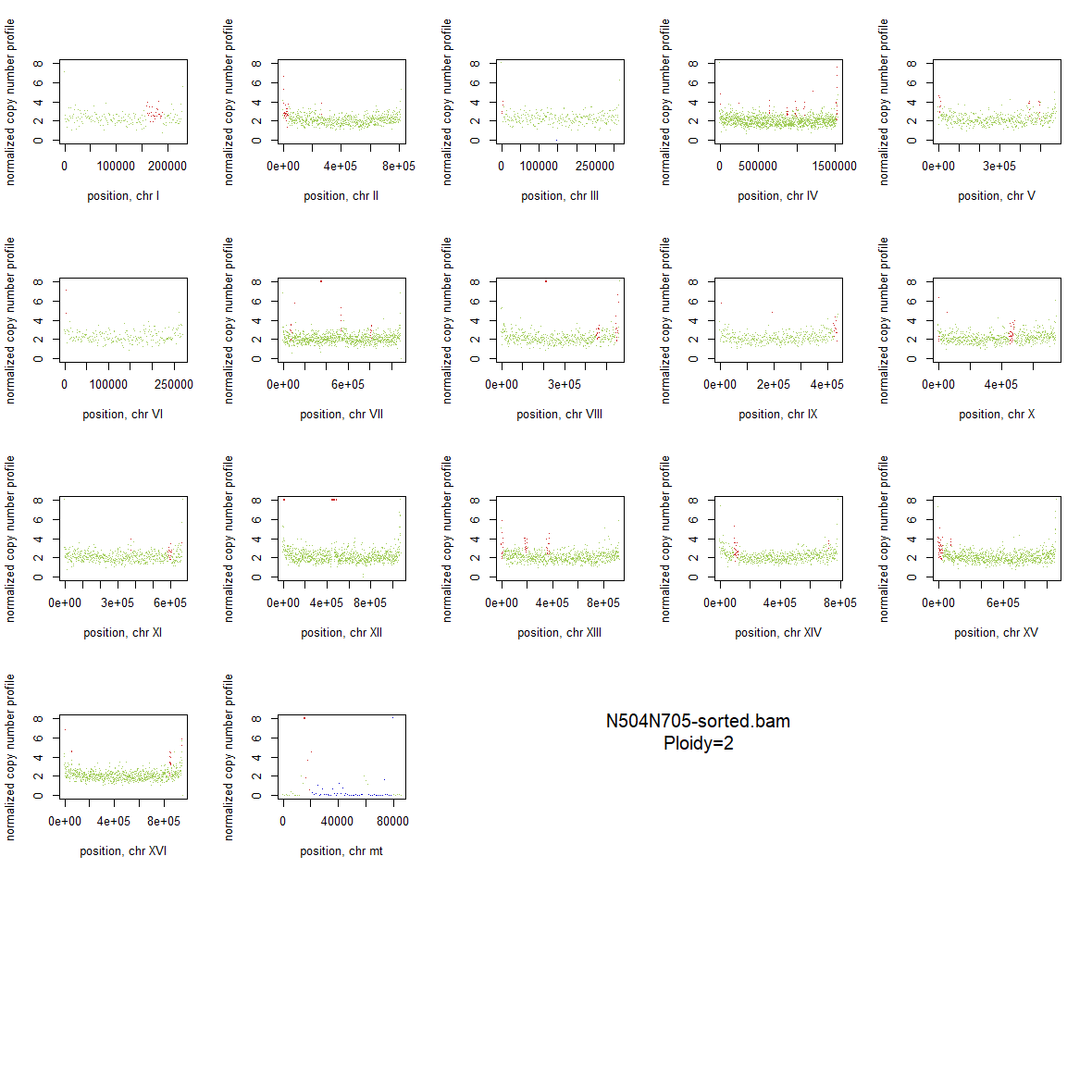

### 2x_V238M_28.png

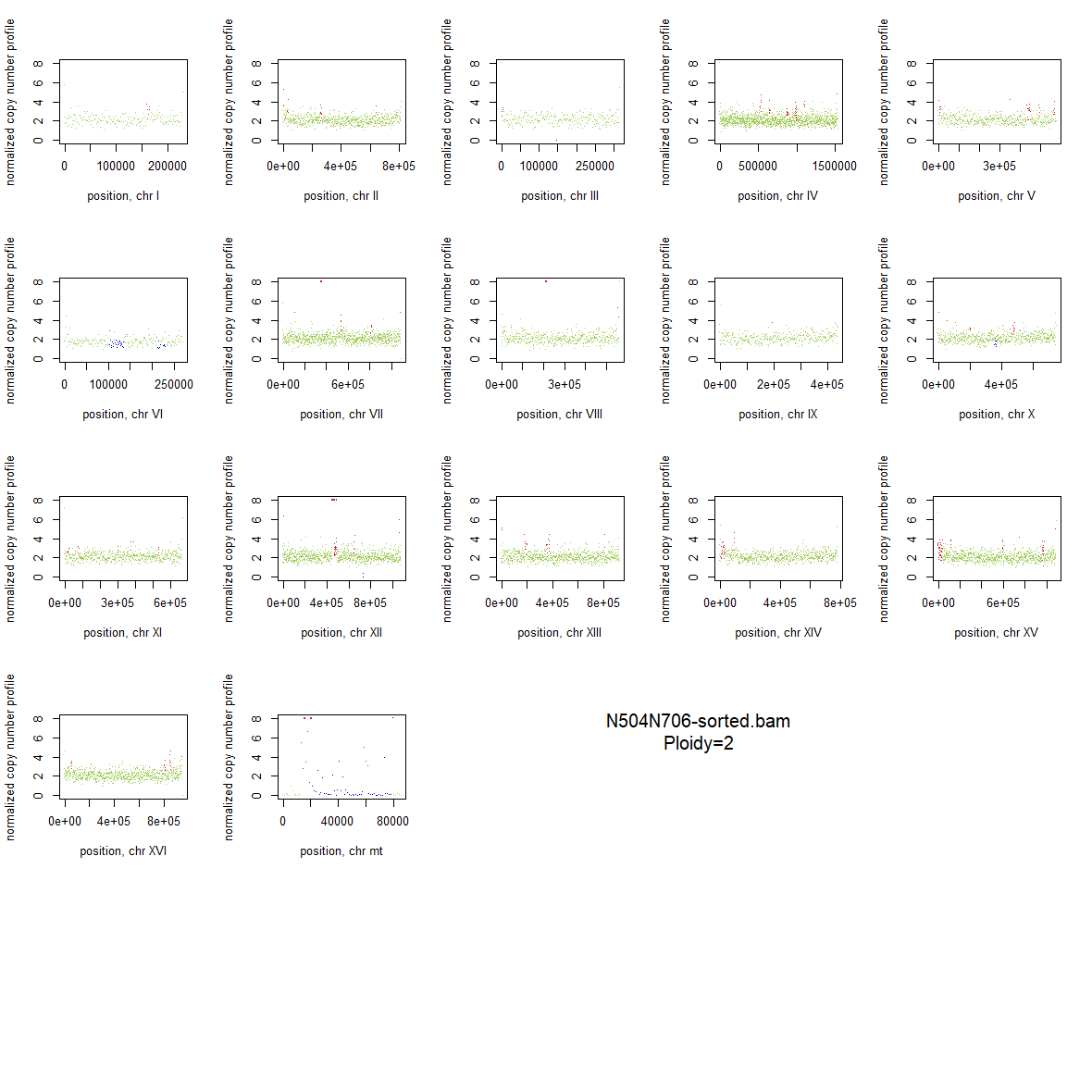

### 2x_V238M_29.png

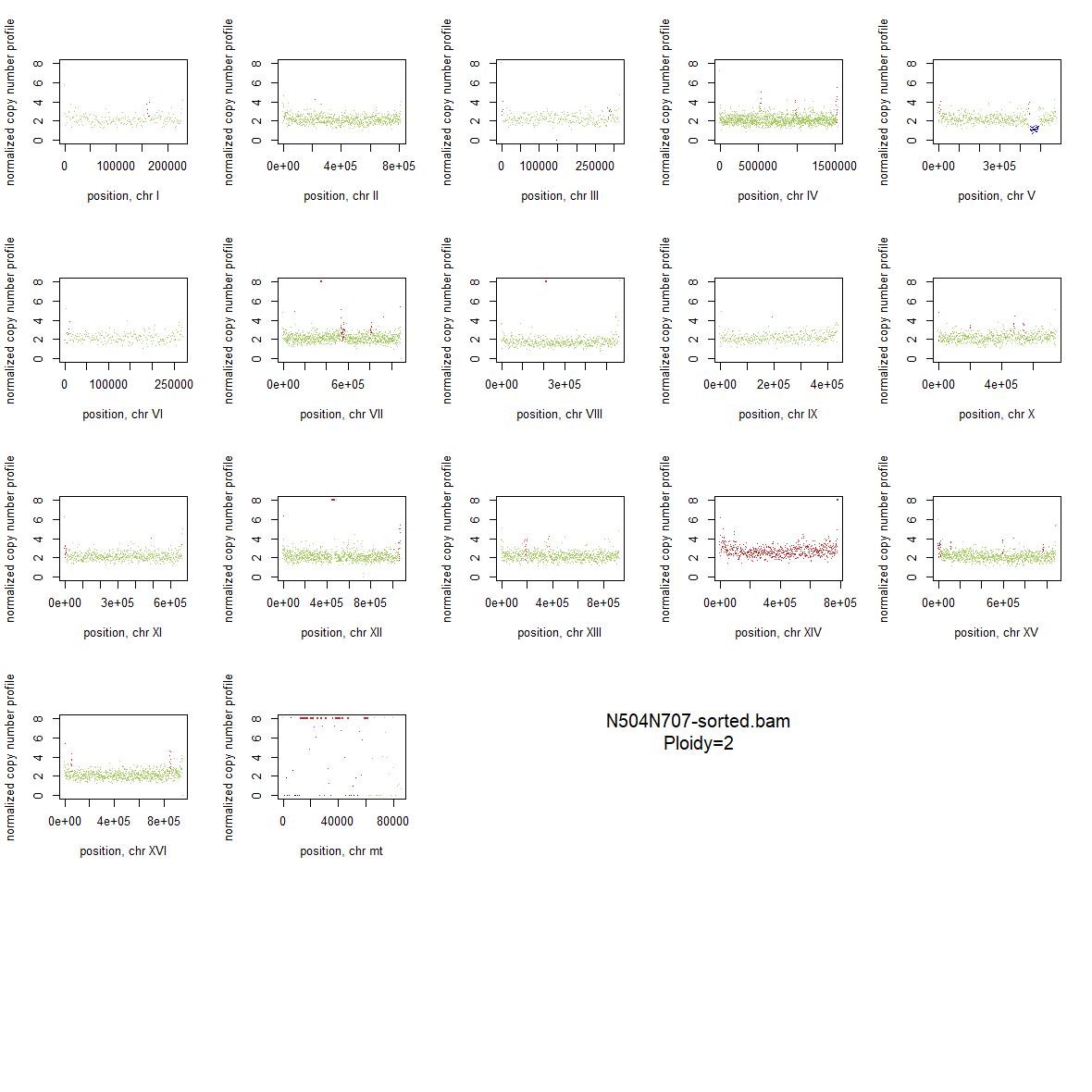

### 2x_V238M_30.png

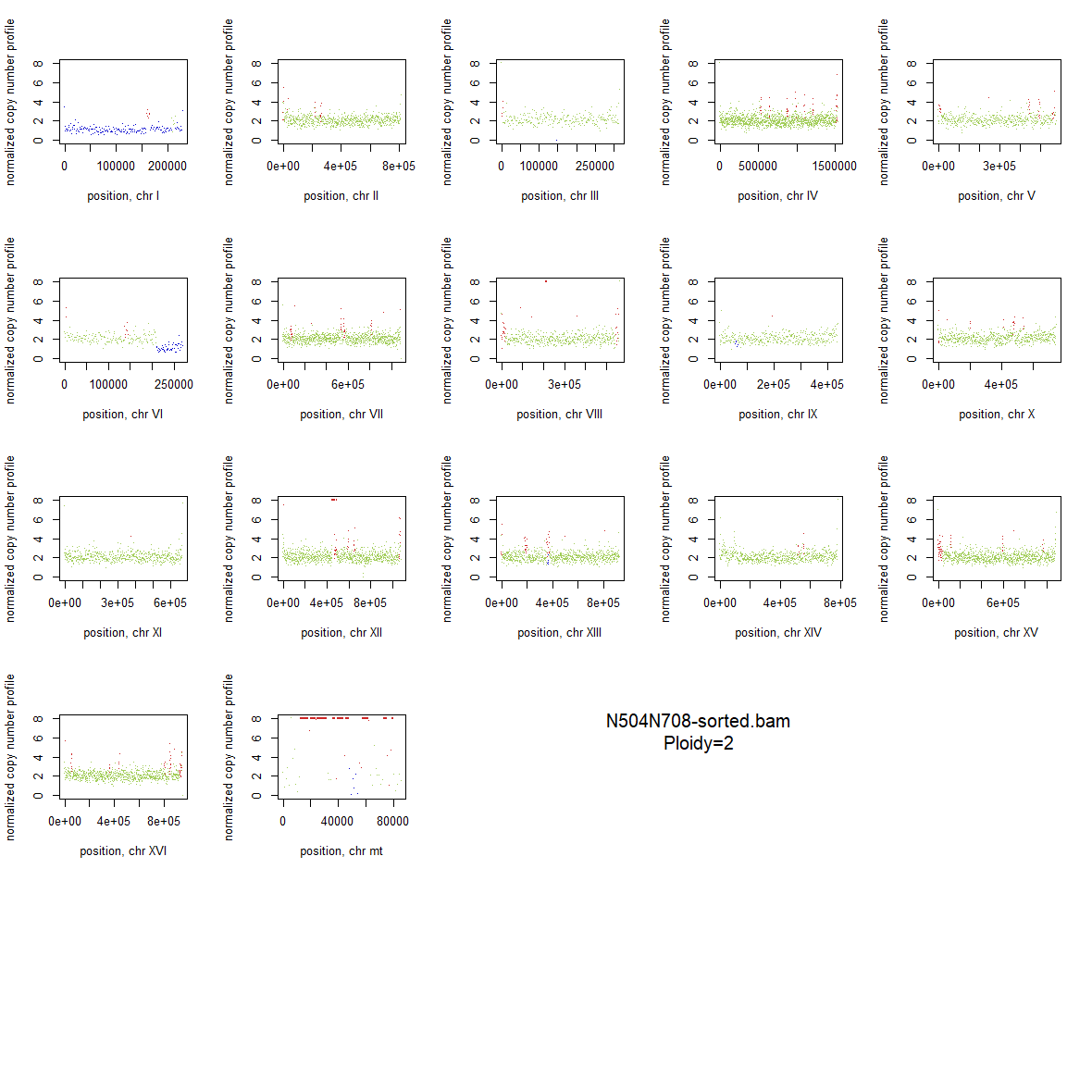

### 2x_V238M_31.png

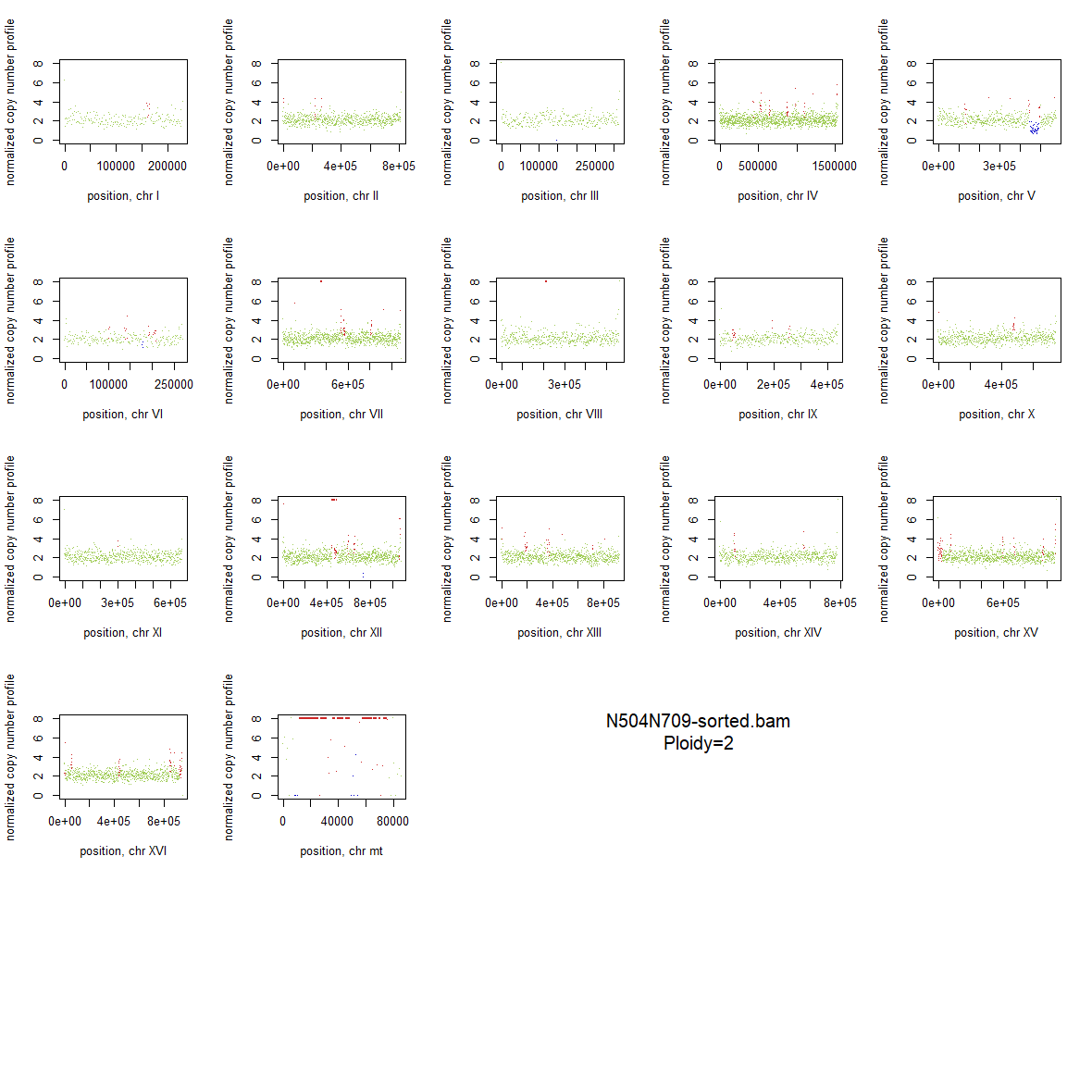

### 2x_V238M_32.png

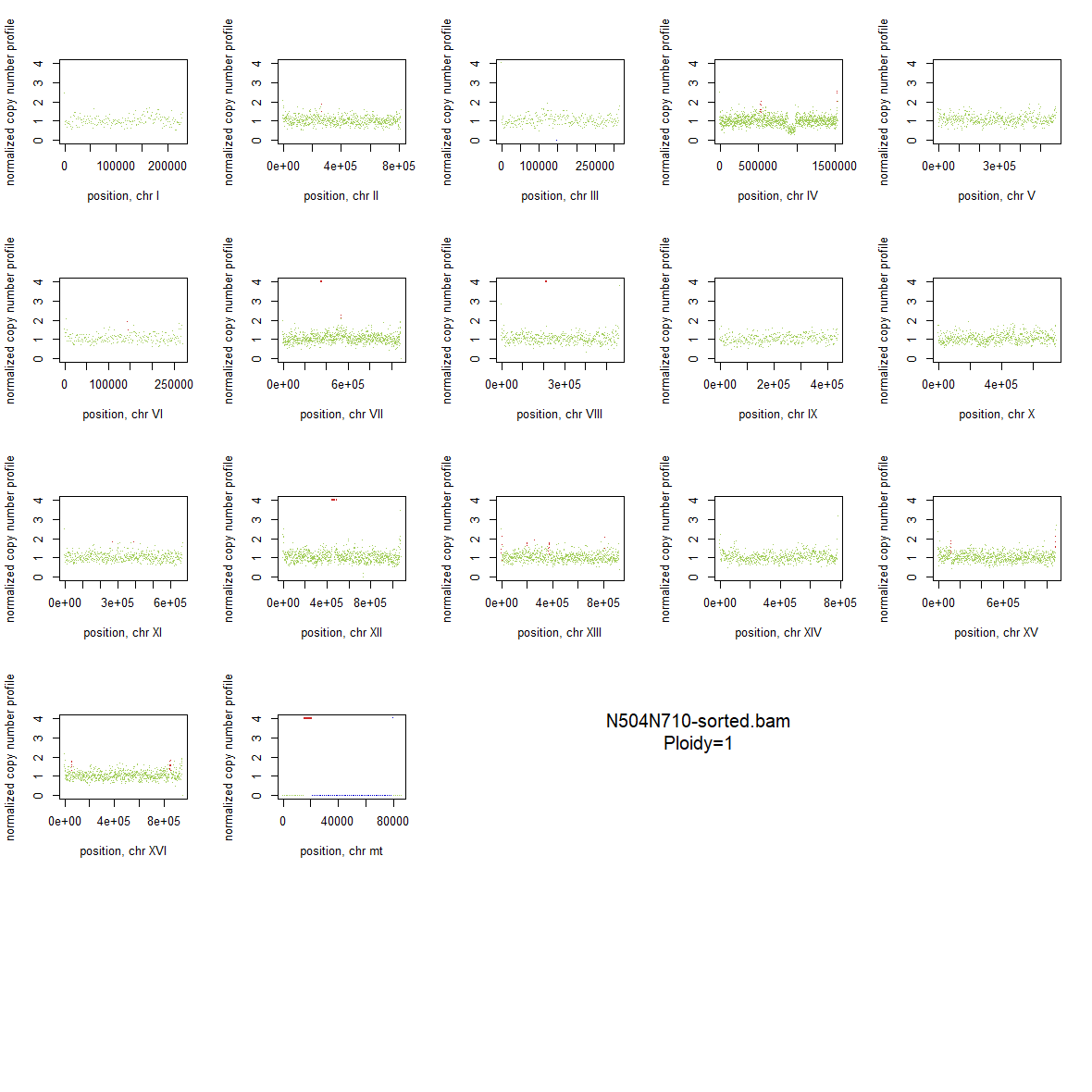

### 2x_V238M_33.png

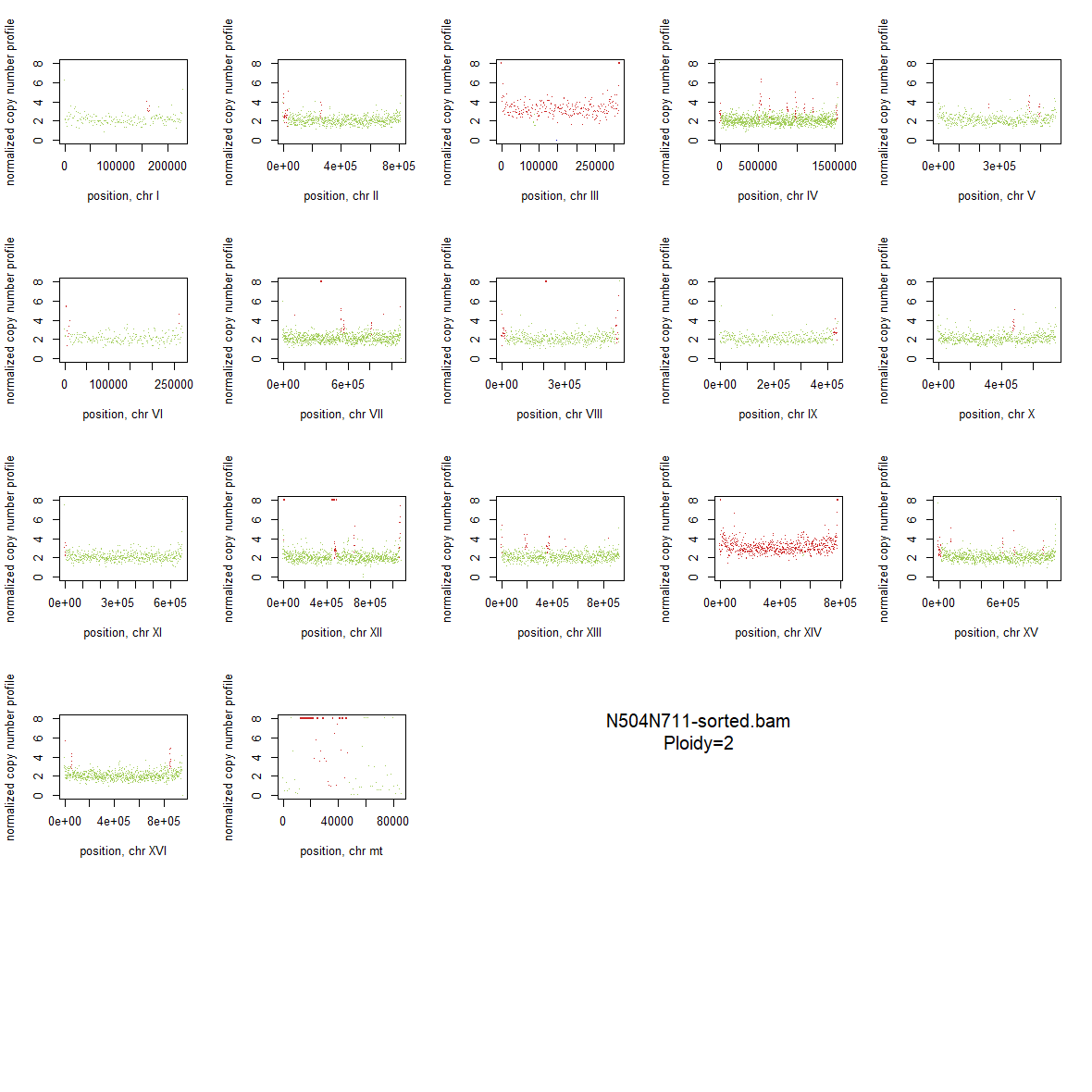

### 2x_V238M_34.png

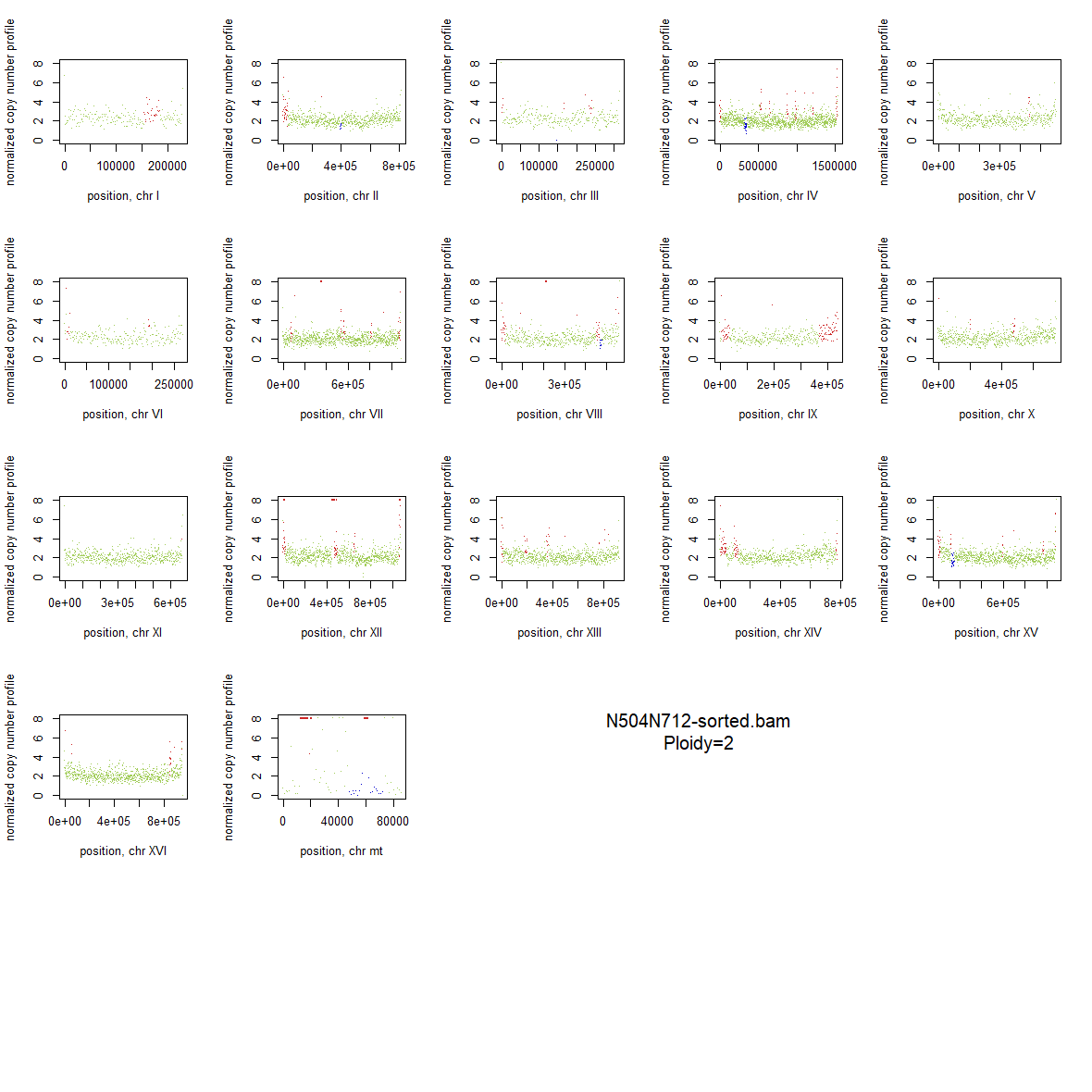

### 2x_V238M_35.png

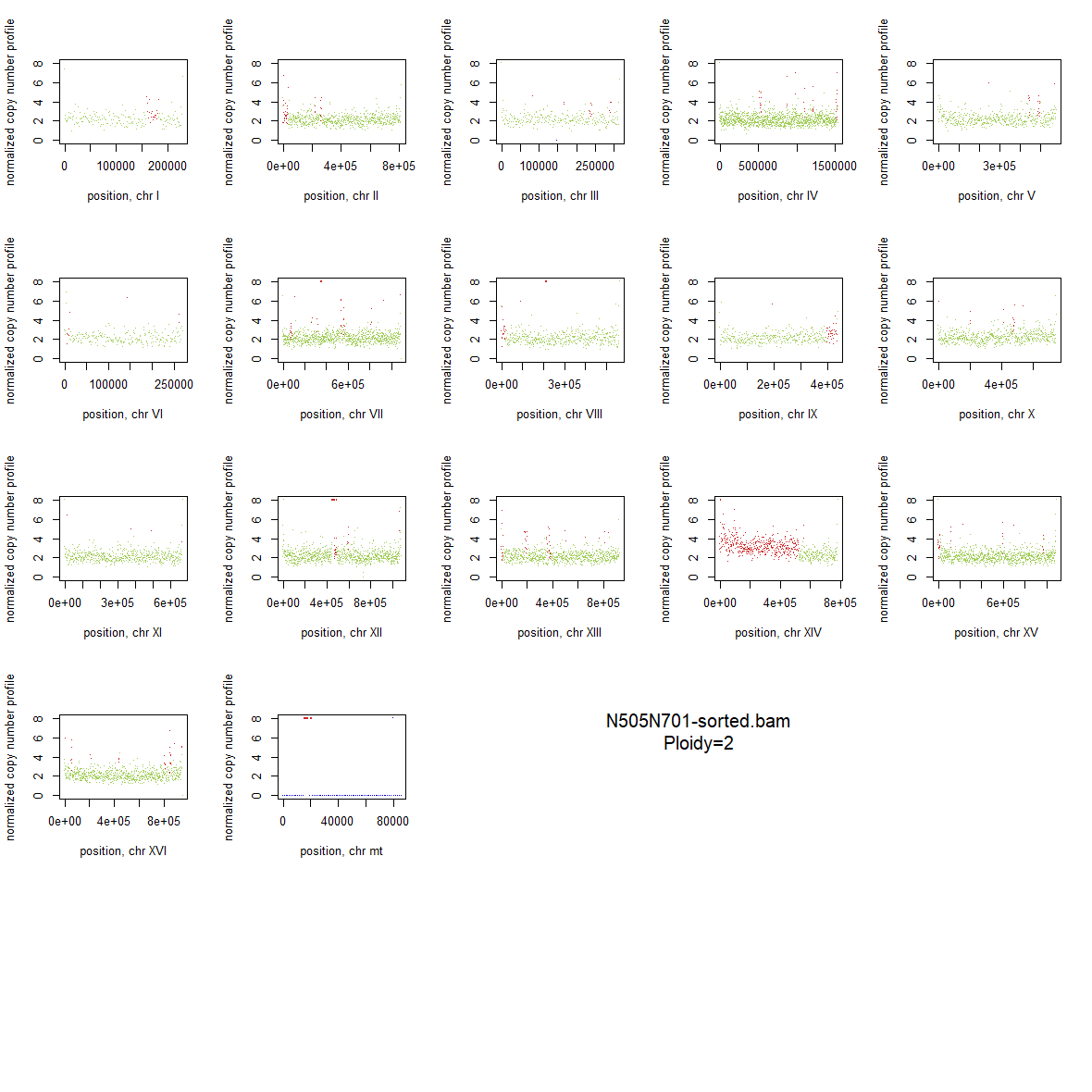

### 2x_V238M_36.png

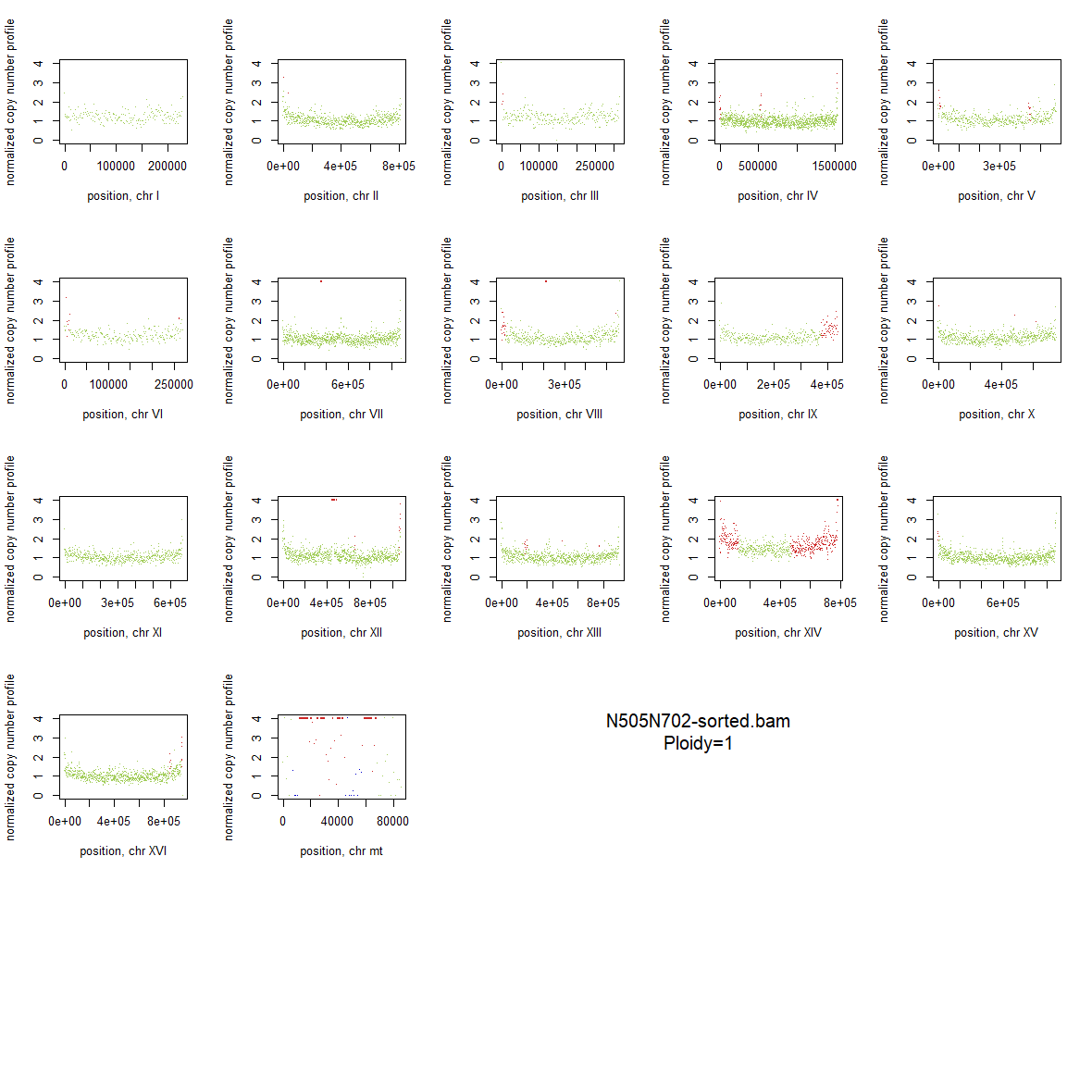

### 2x_V238M_37.png

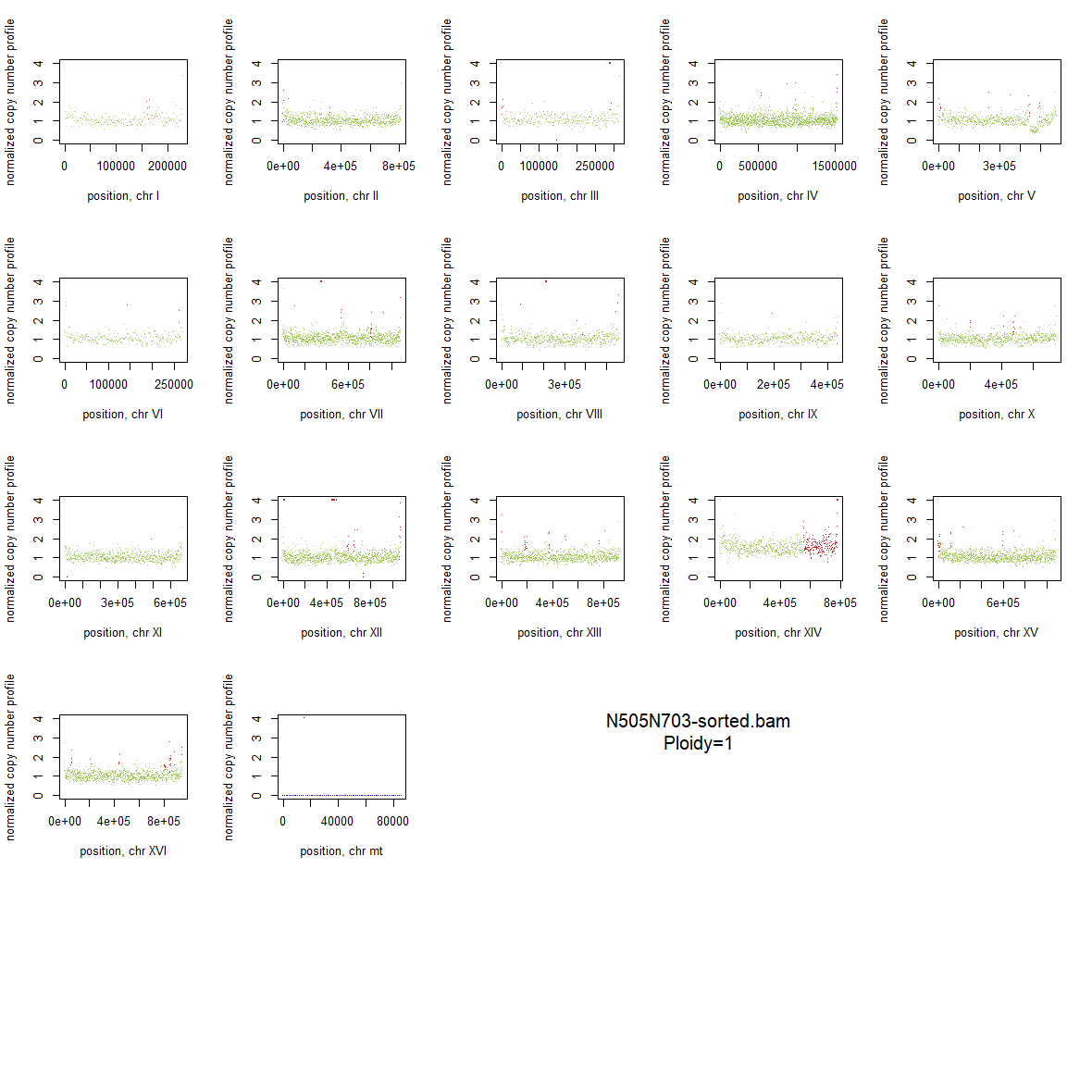

### 2x_V238M_38.png

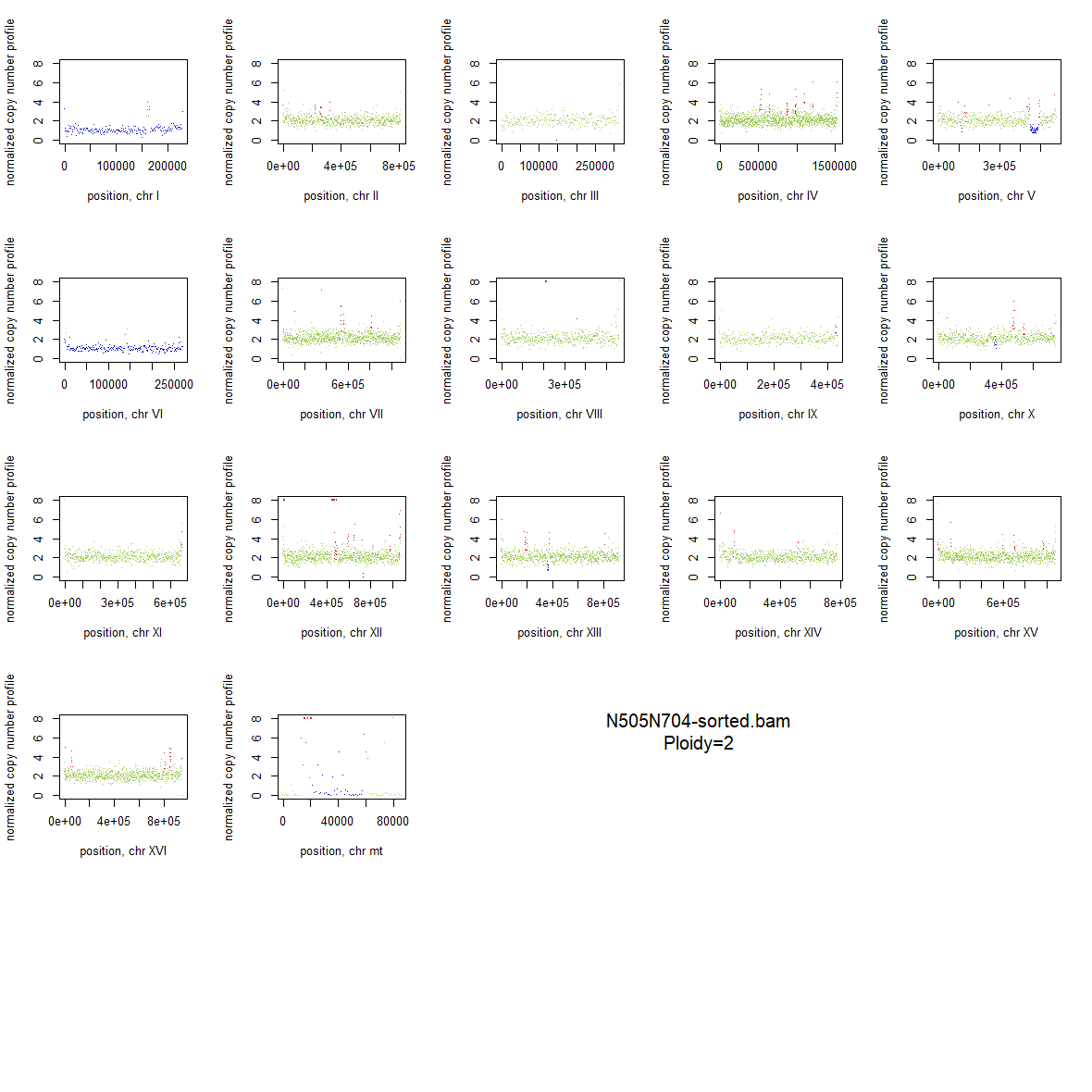

### 2x_V238M_39.png

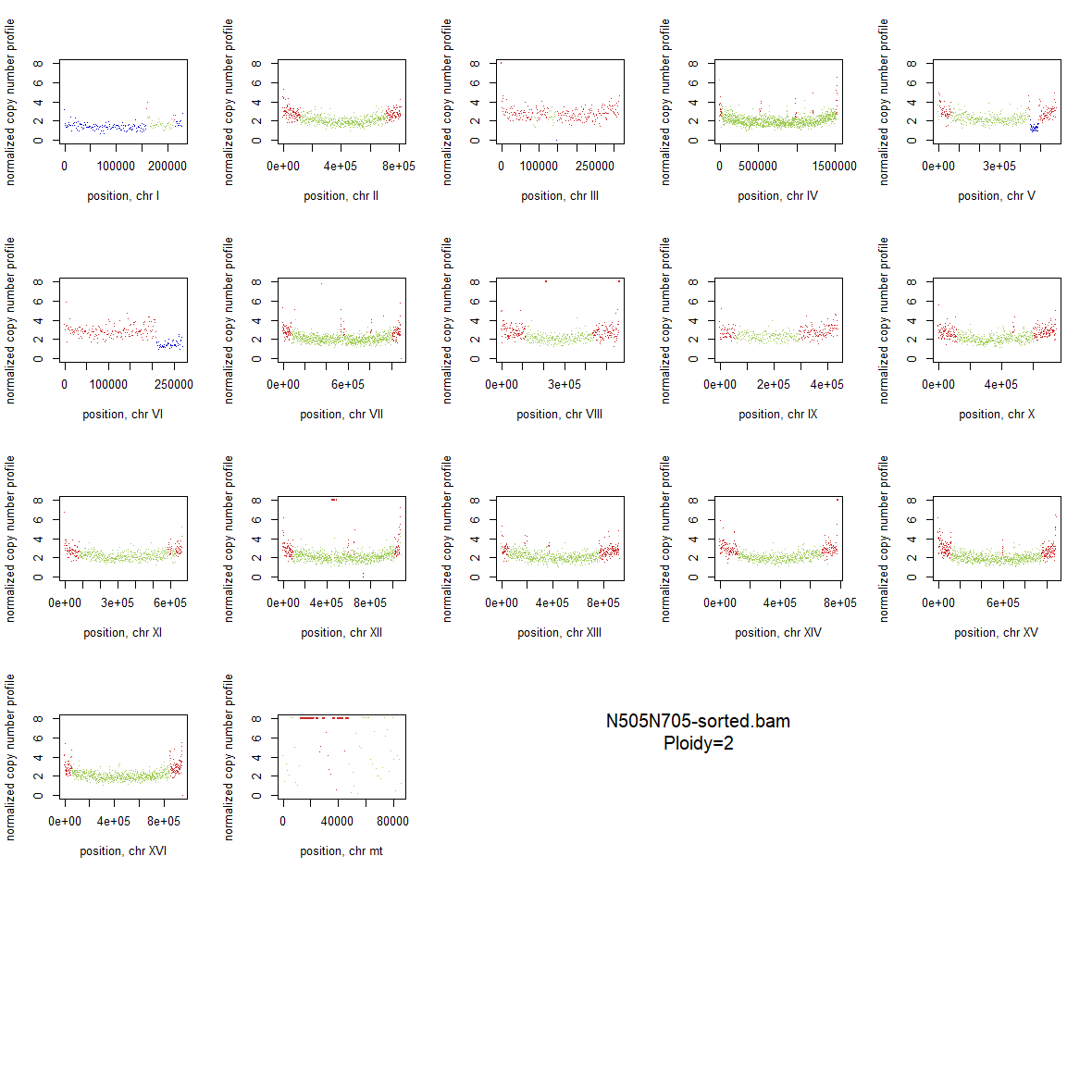

### 2x_V238M_40.png

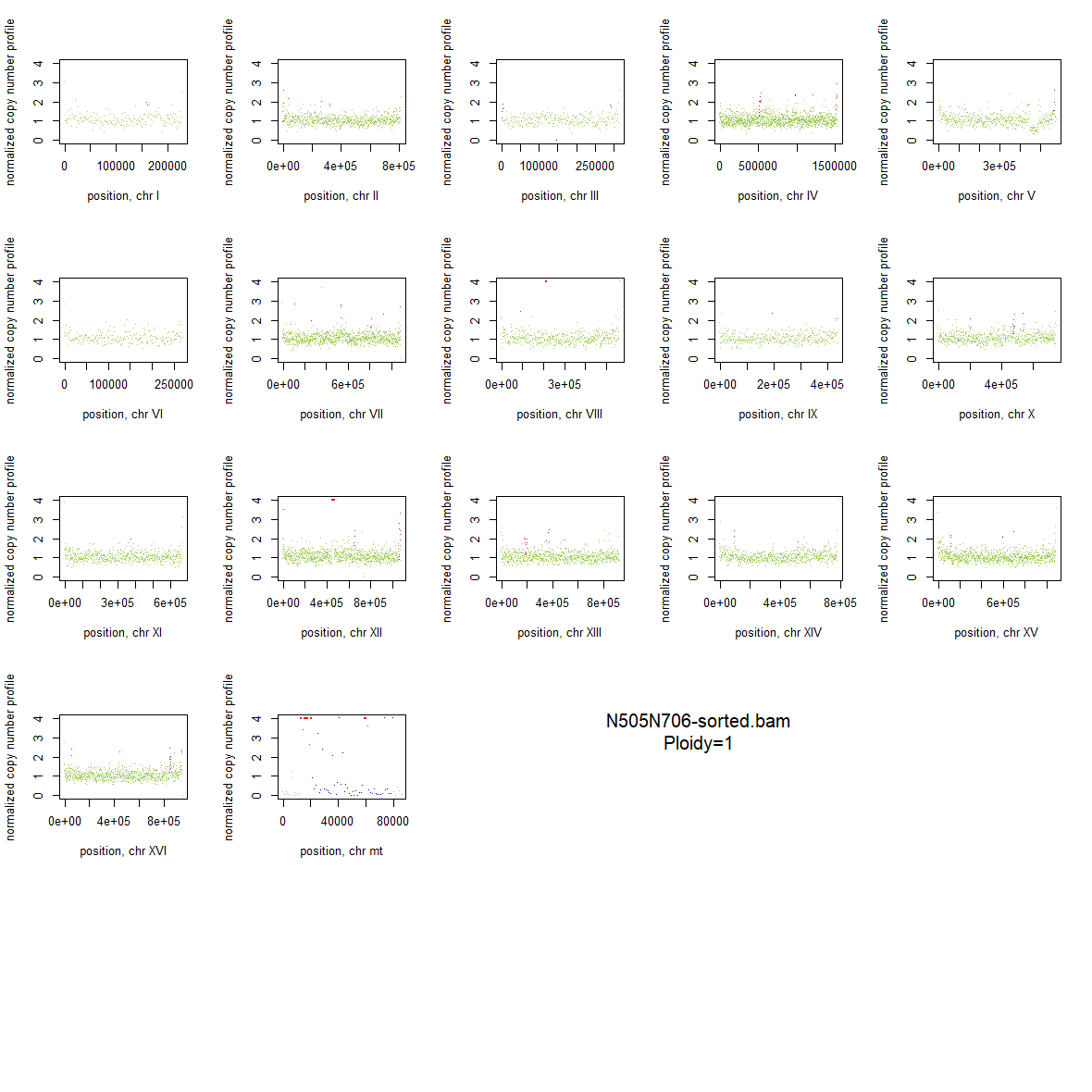

### 2x_V238M_41.png

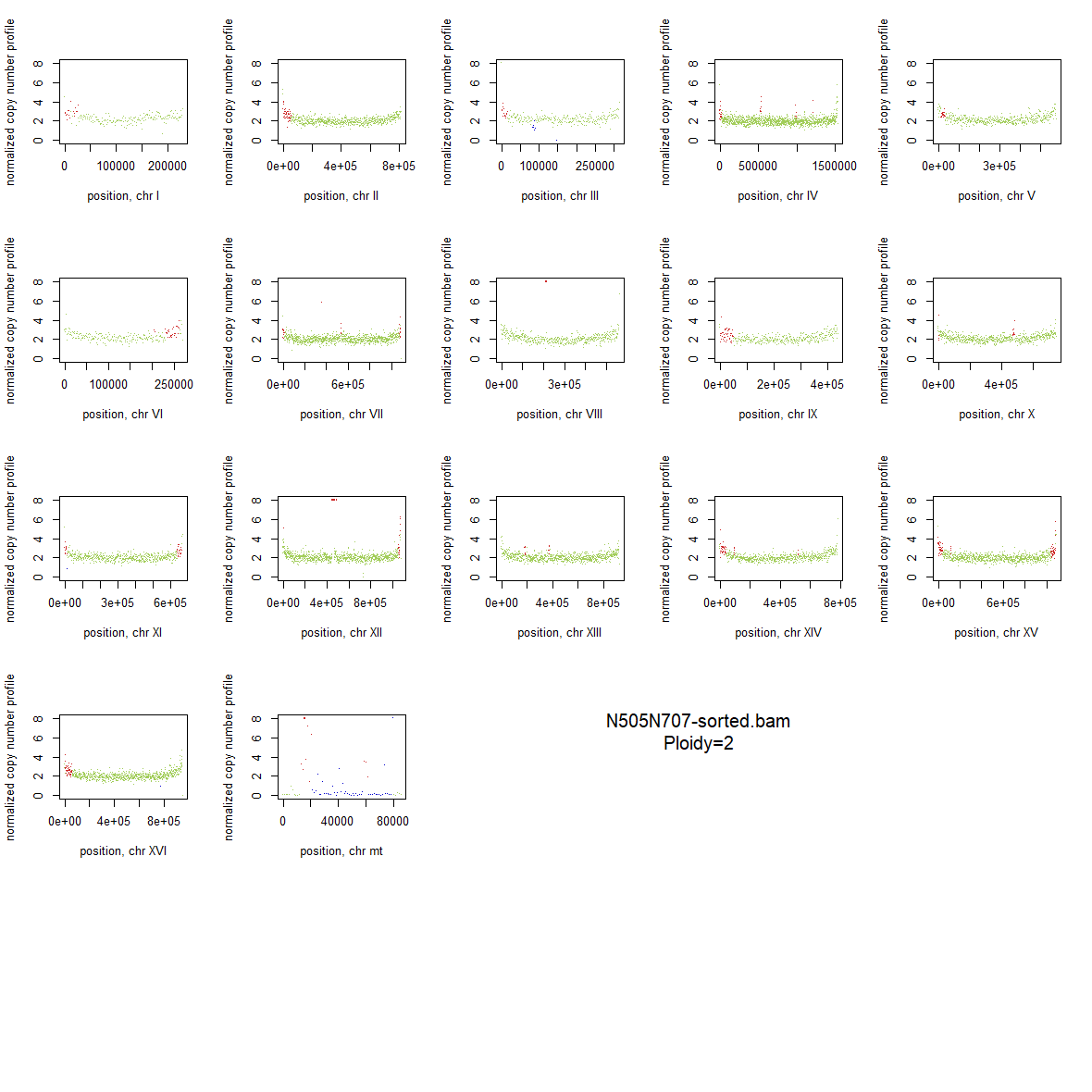

### 2x_V238M_42.png

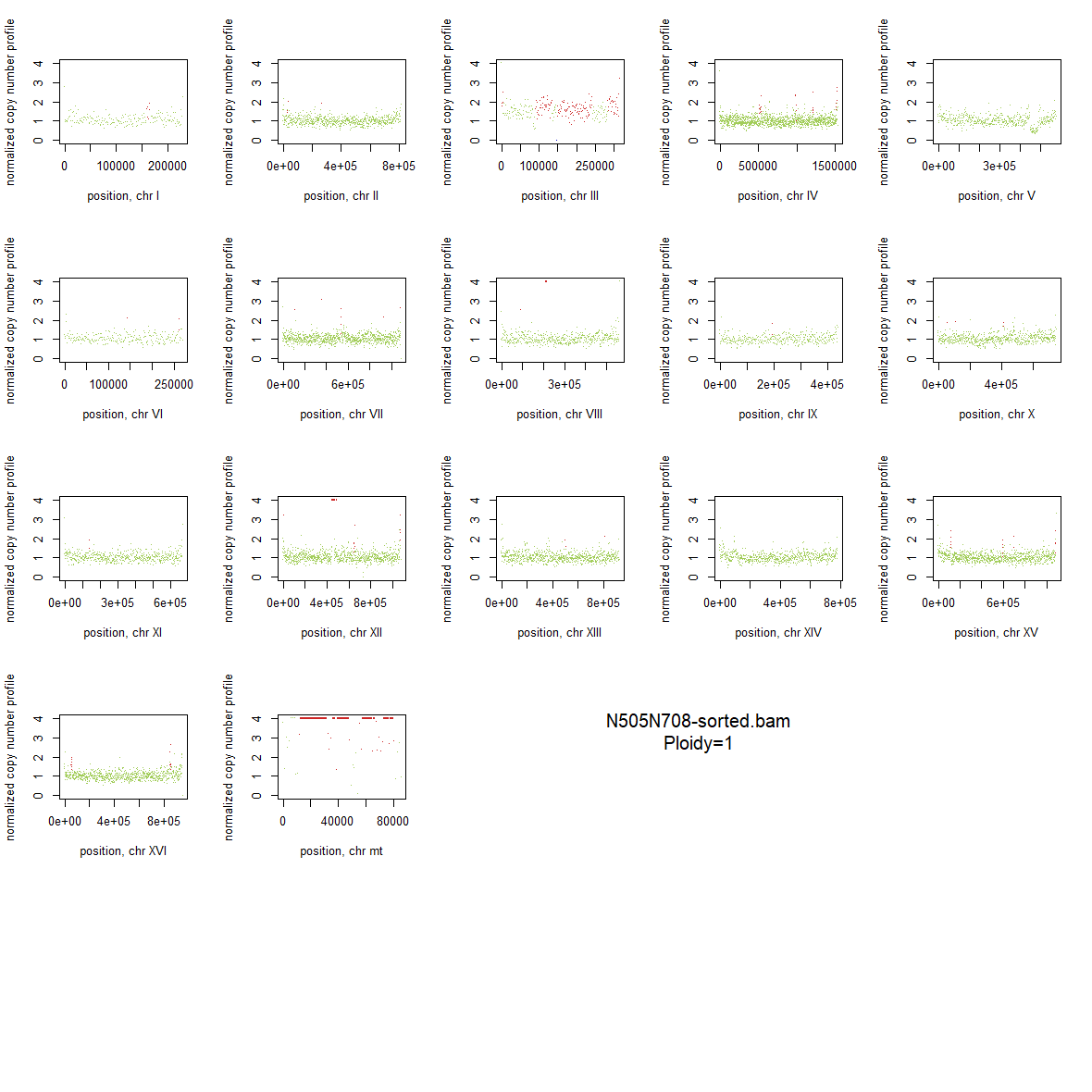

### 2x_V238M_43.png

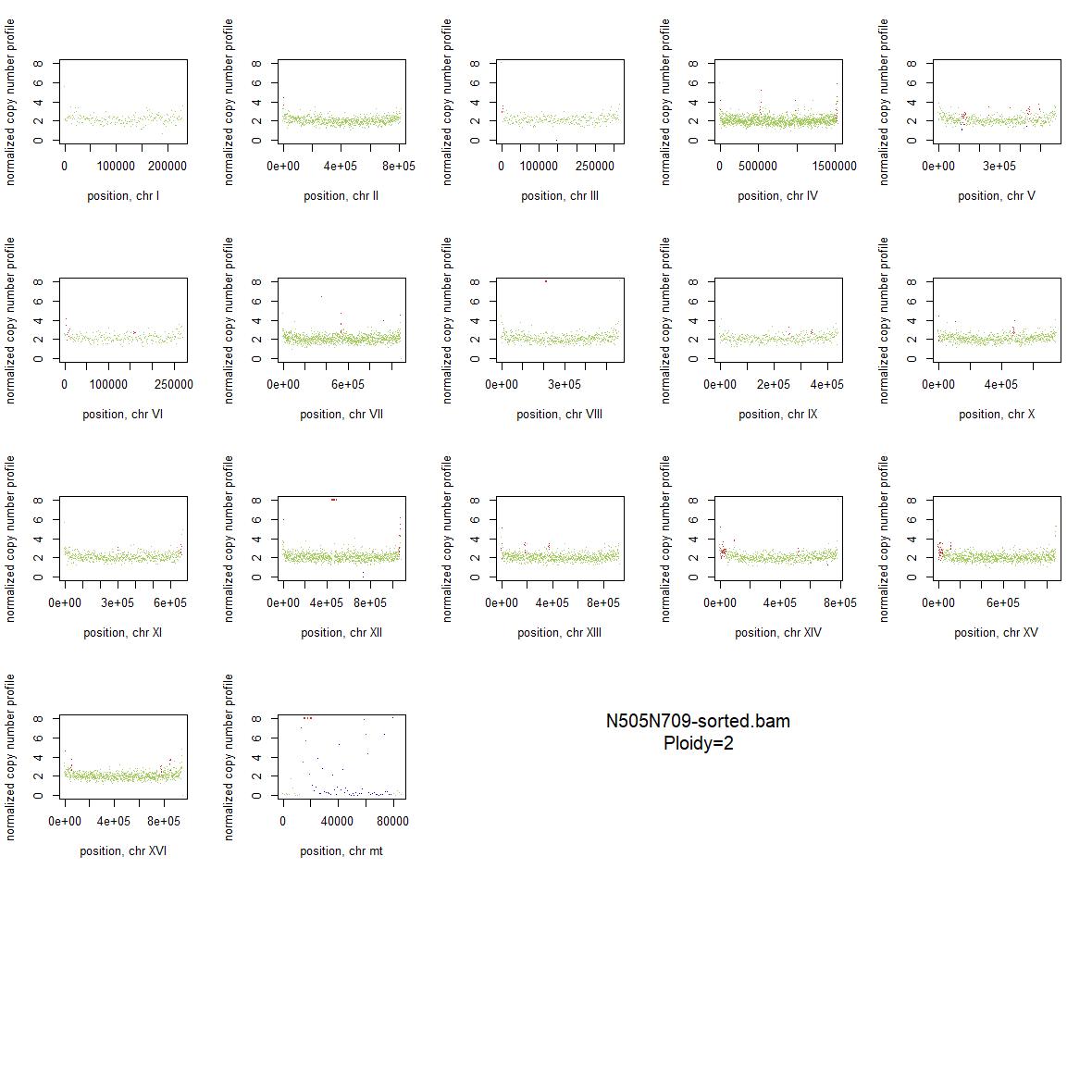

### 2x_V238M_44.png

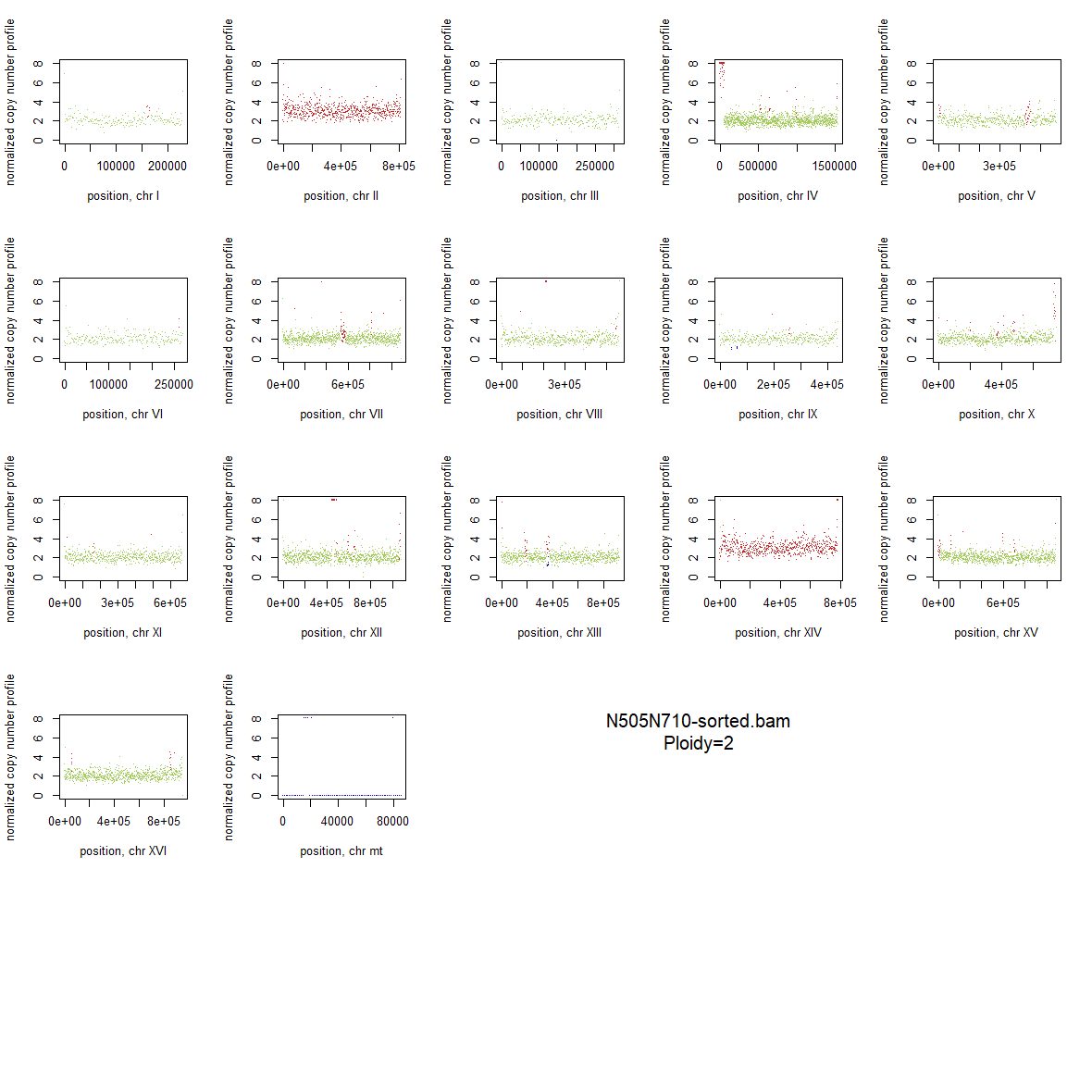

### 2x_V238M_45.png

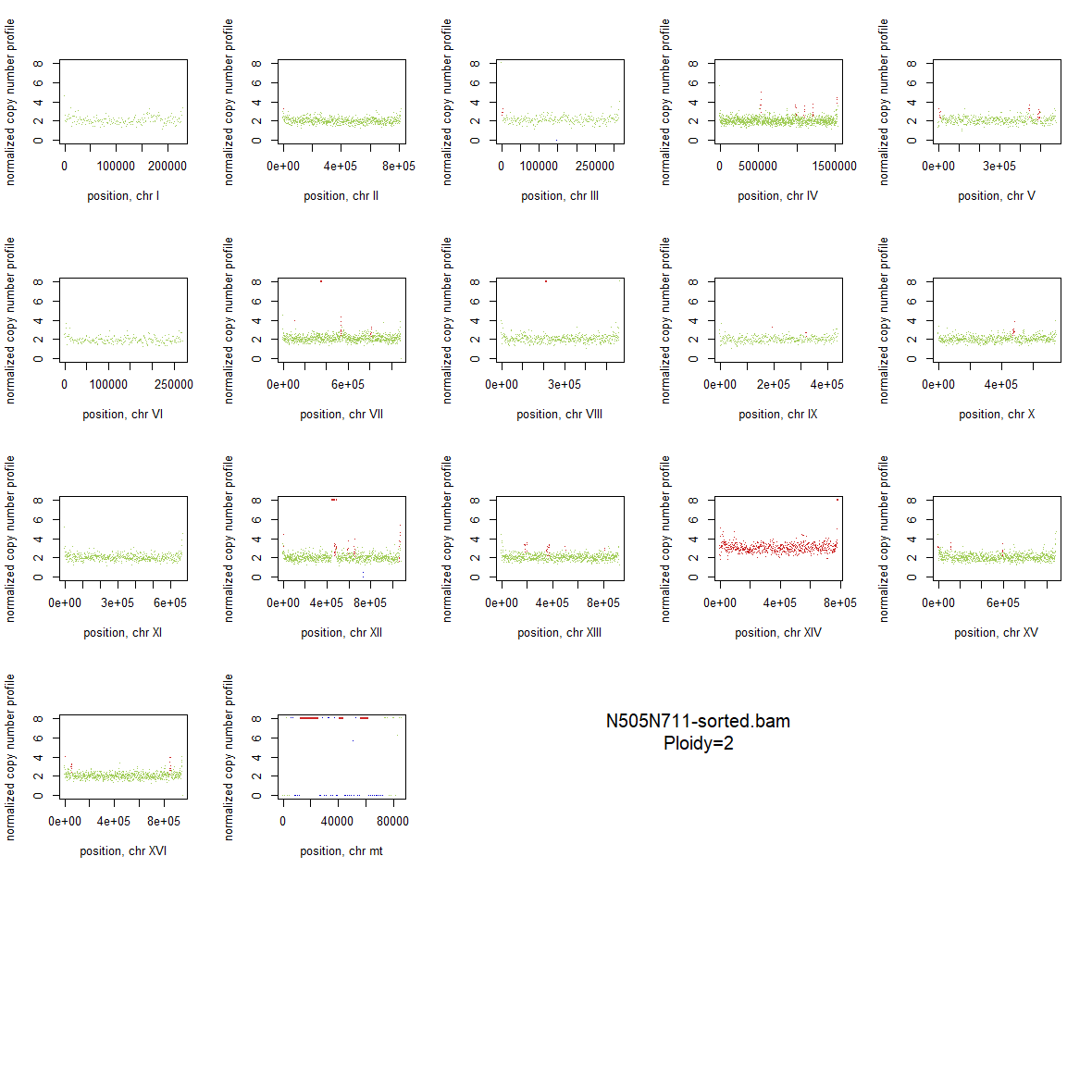

### 2x_V238M_46.png

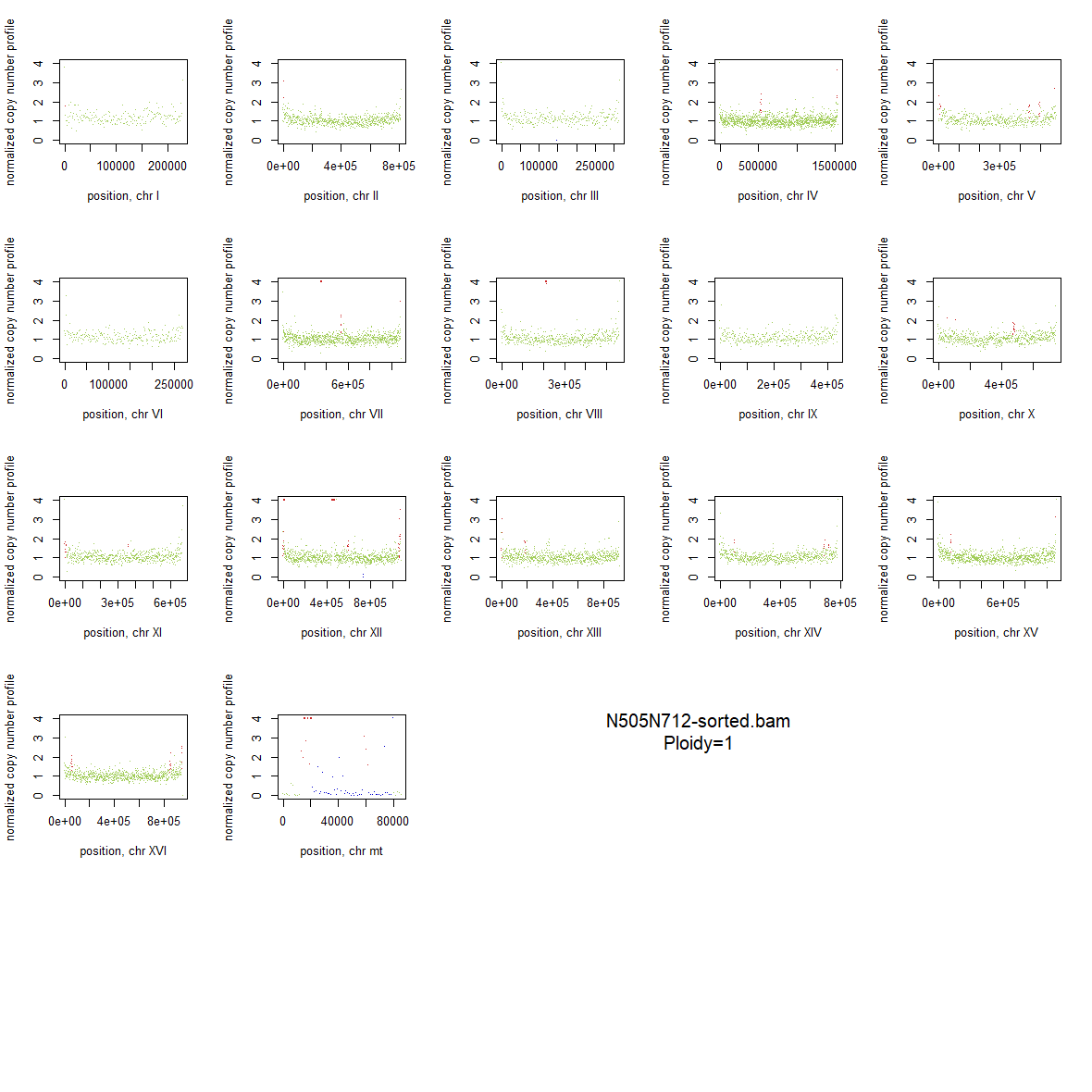

### 2x_V238M_47.png

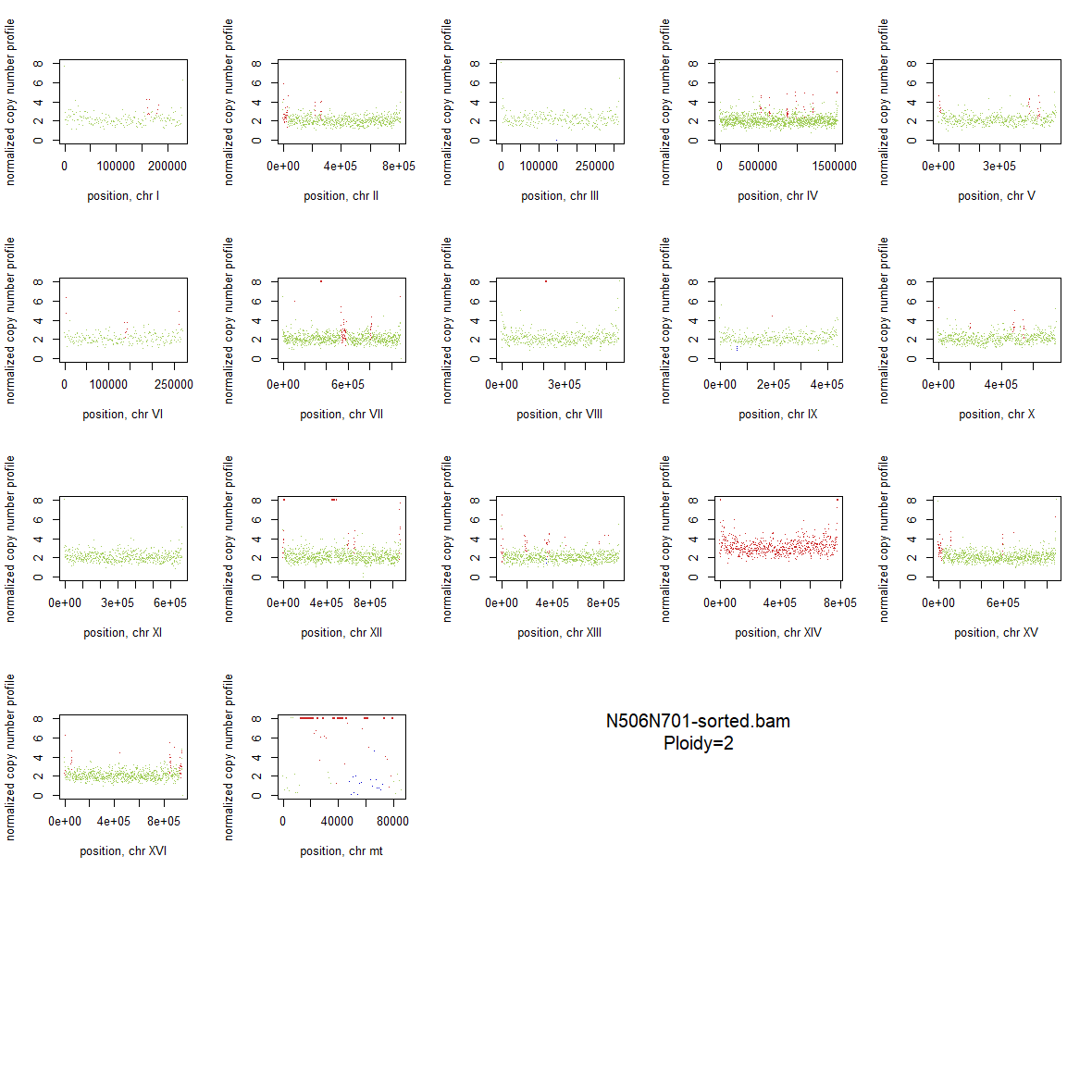

### 2x_V238M_48.png

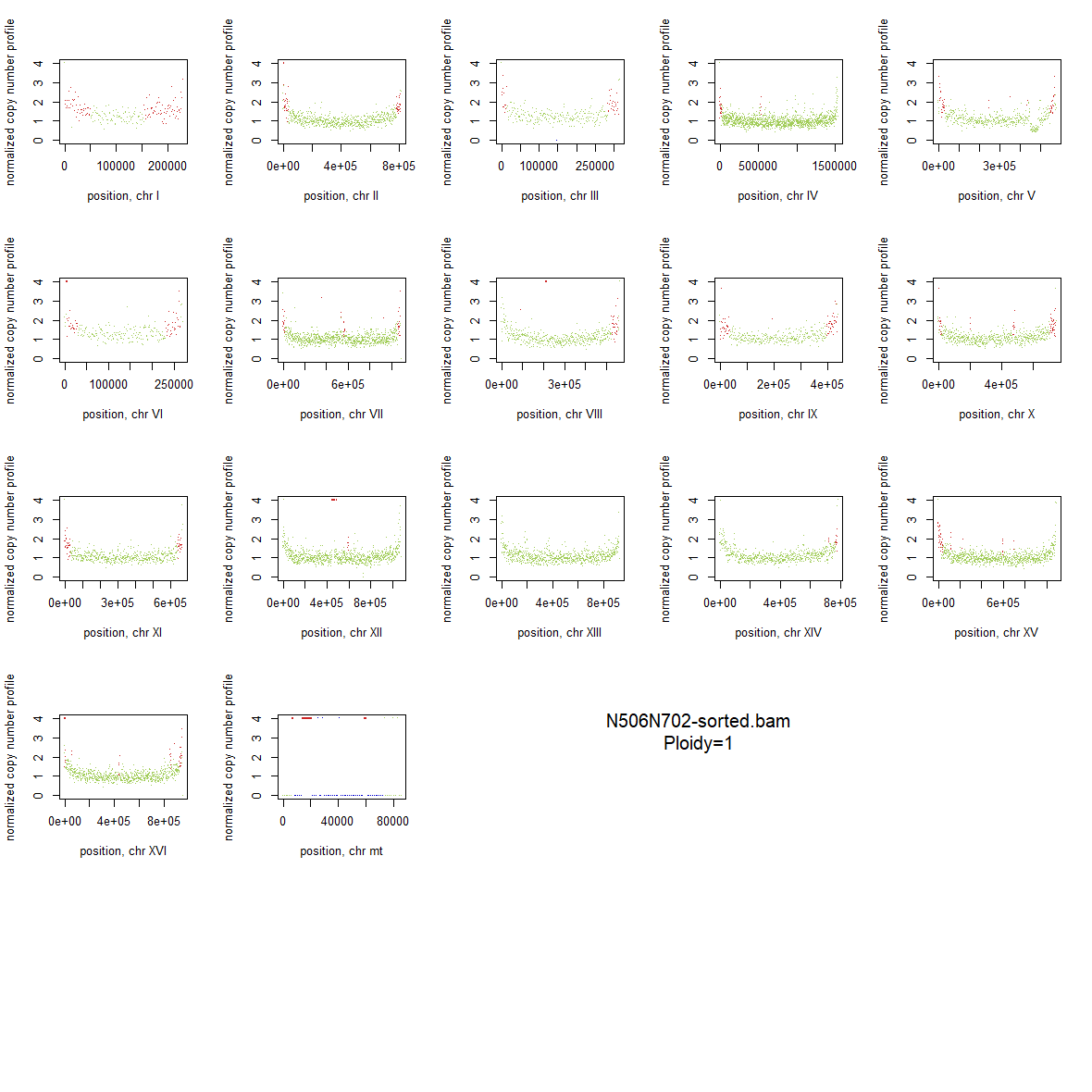

### 2x_V238M_49.png

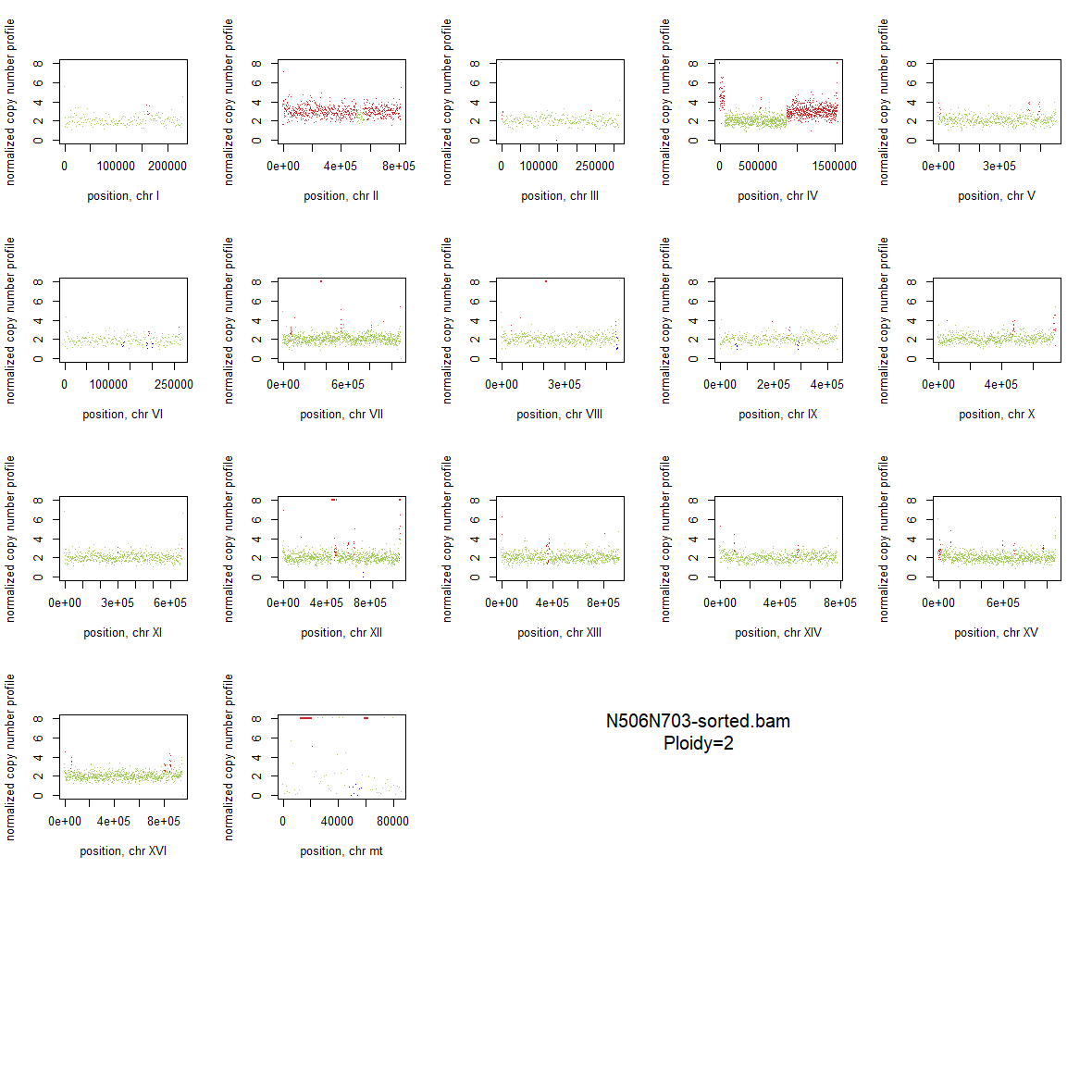

### 2x_V238M_50.png

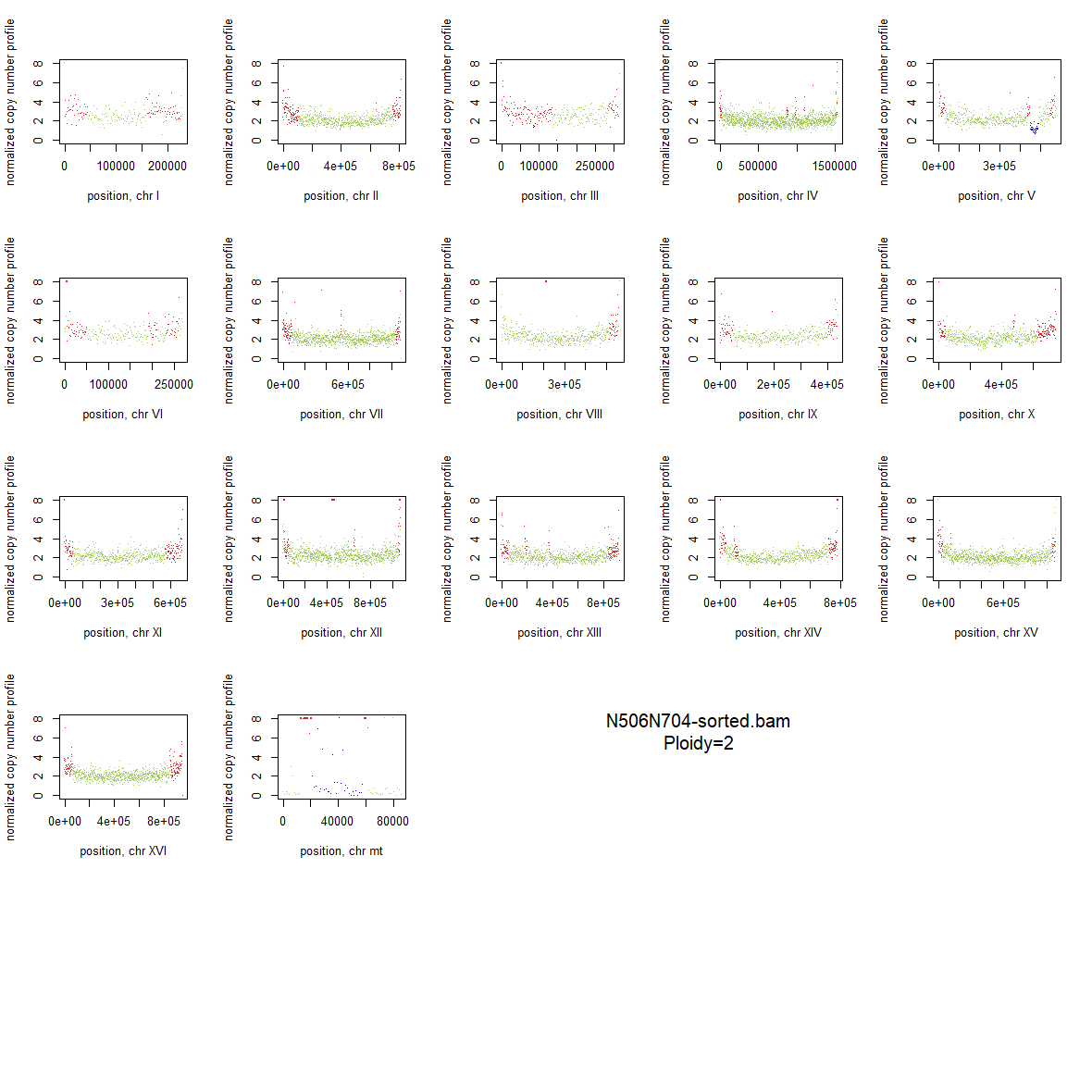

### 2x_V238M_51.png

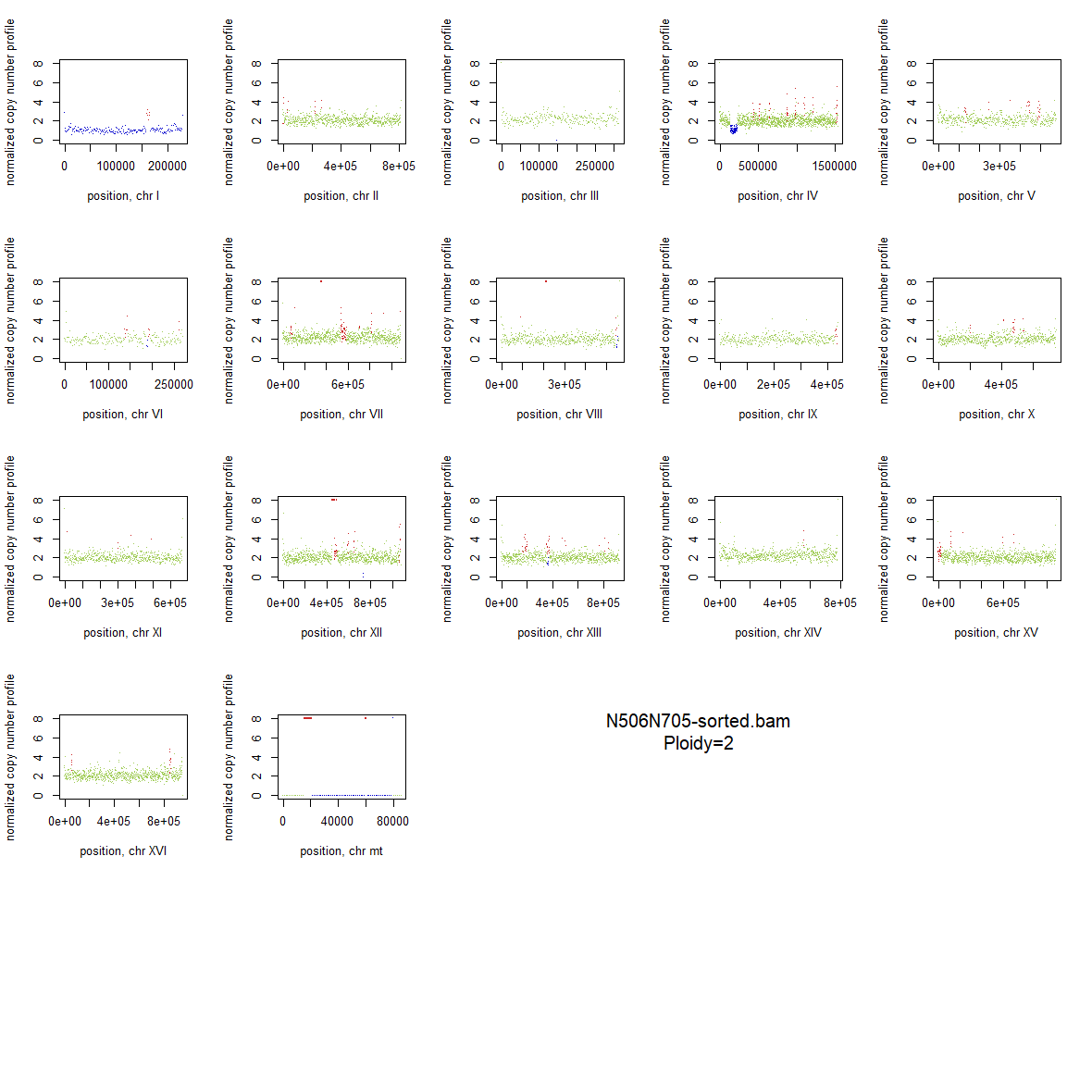

### 2x_V238M_52.png

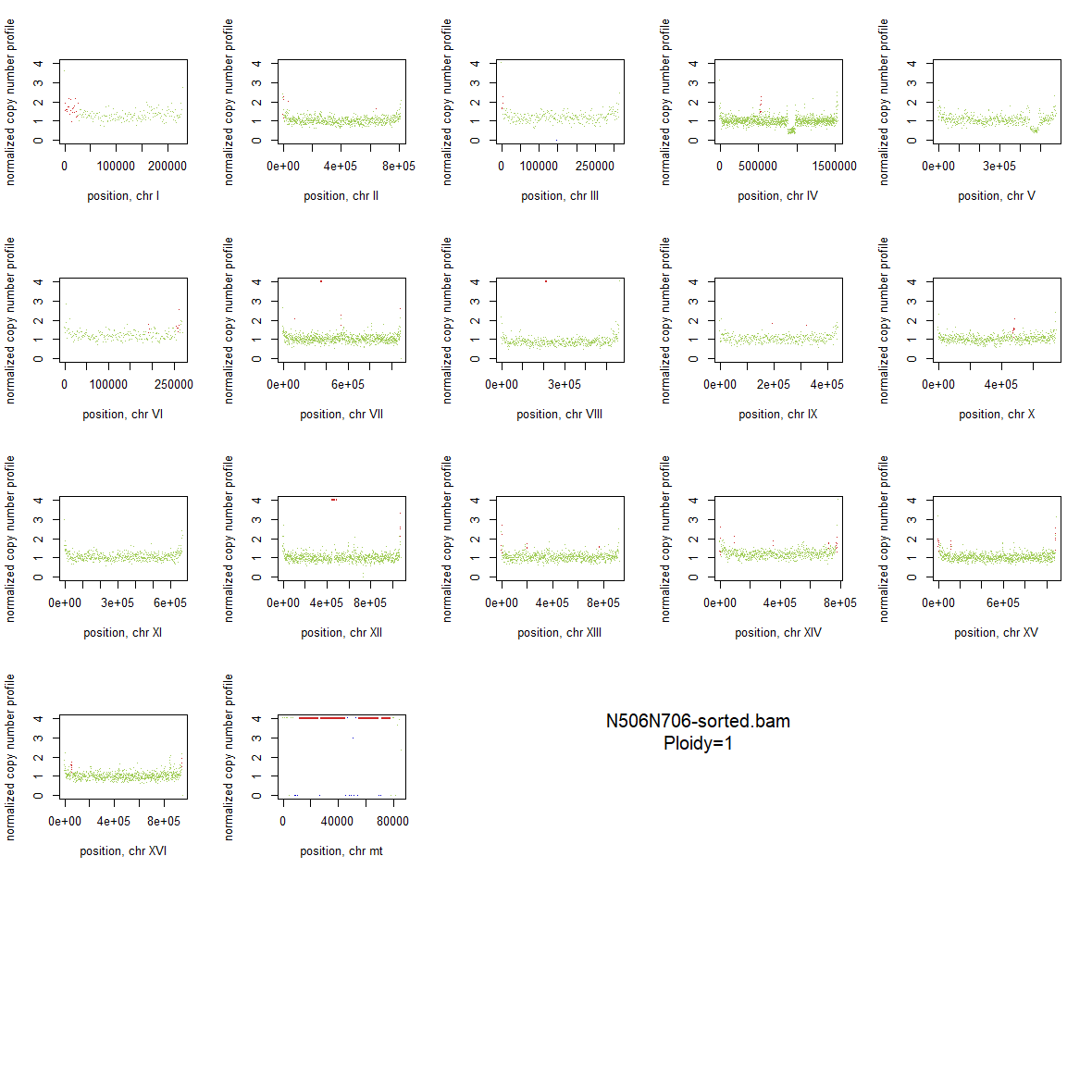

### 2x_V238M_53.png

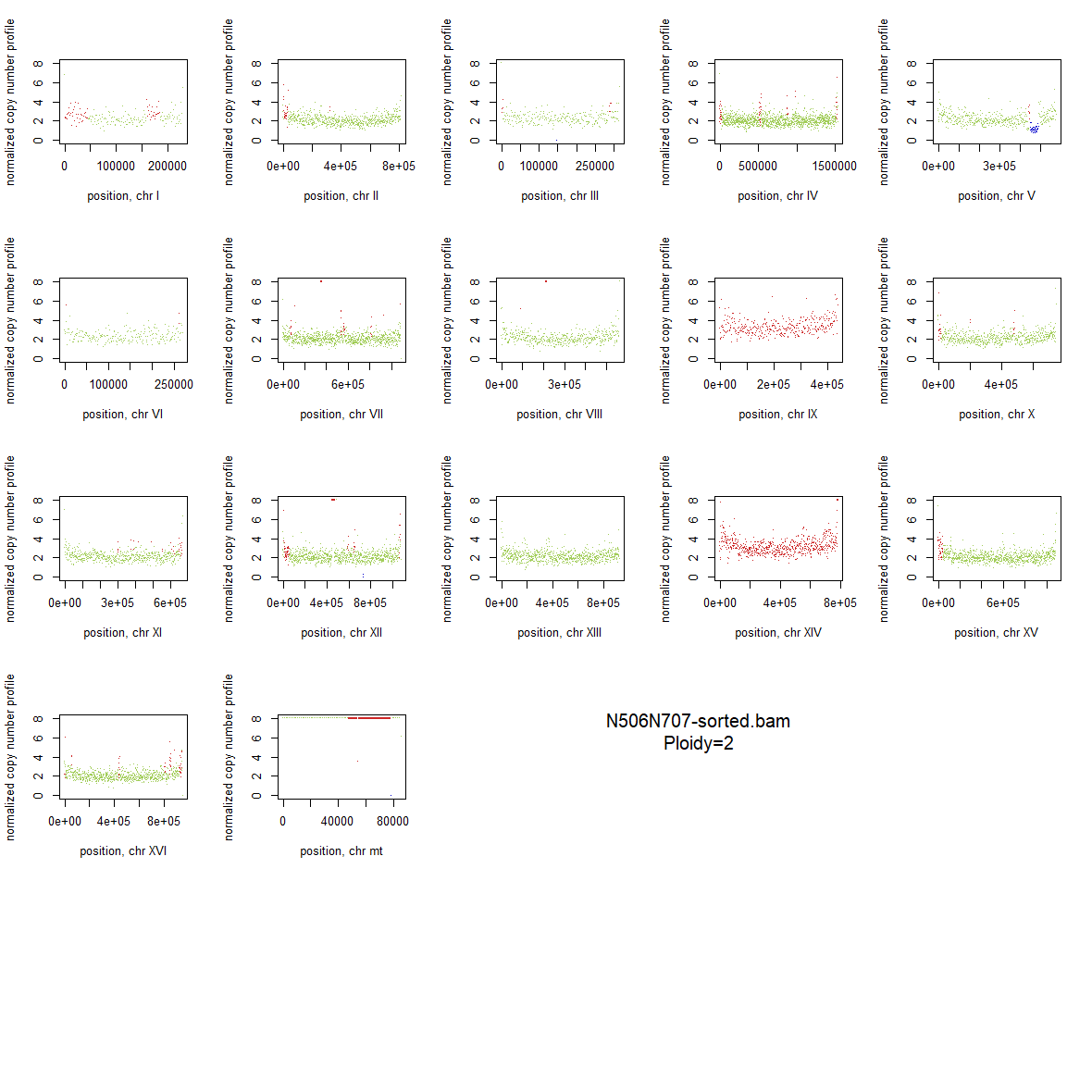

### 2x_V238M_54.png

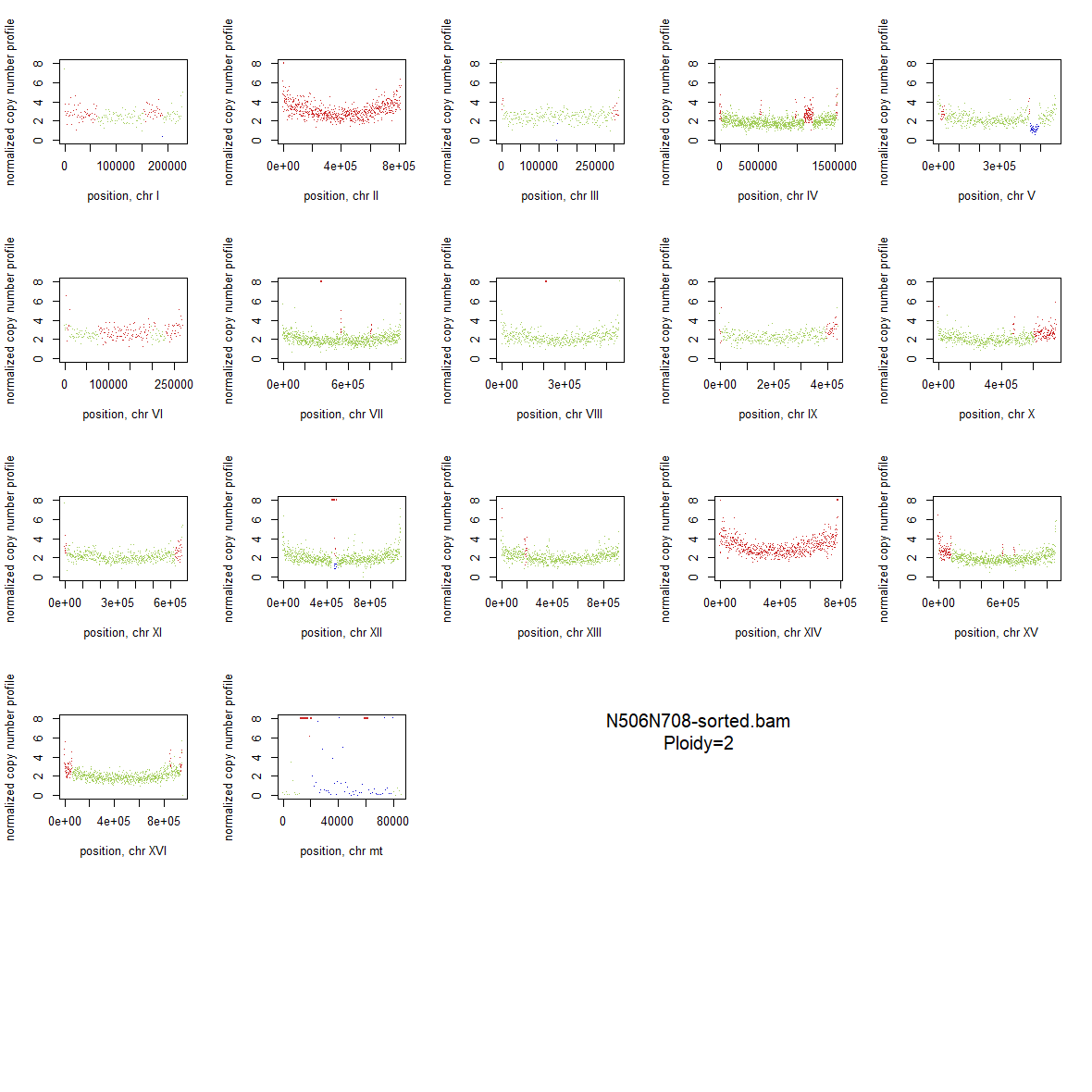

### 2x_V238M_55.png

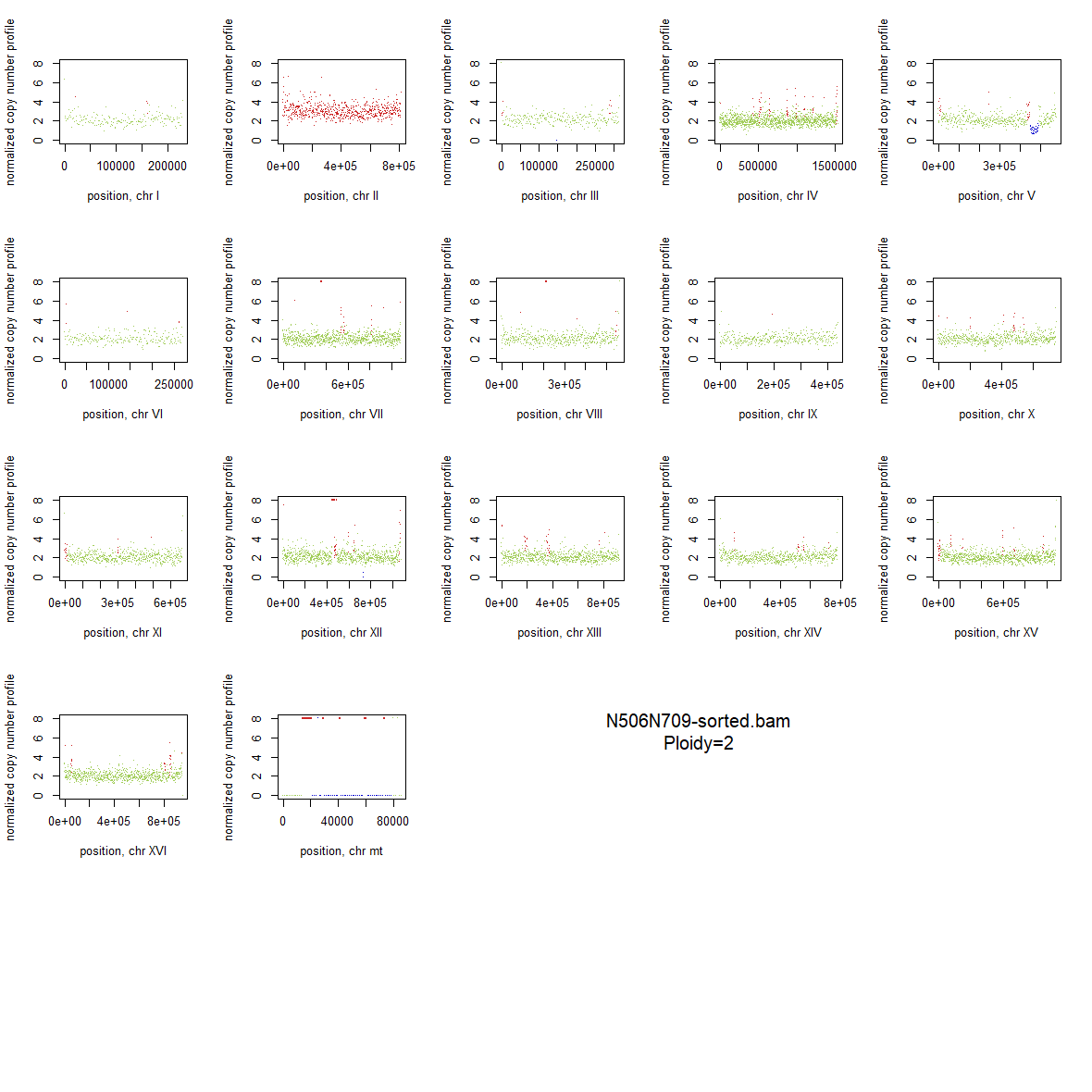

### 2x_V238M_56.png

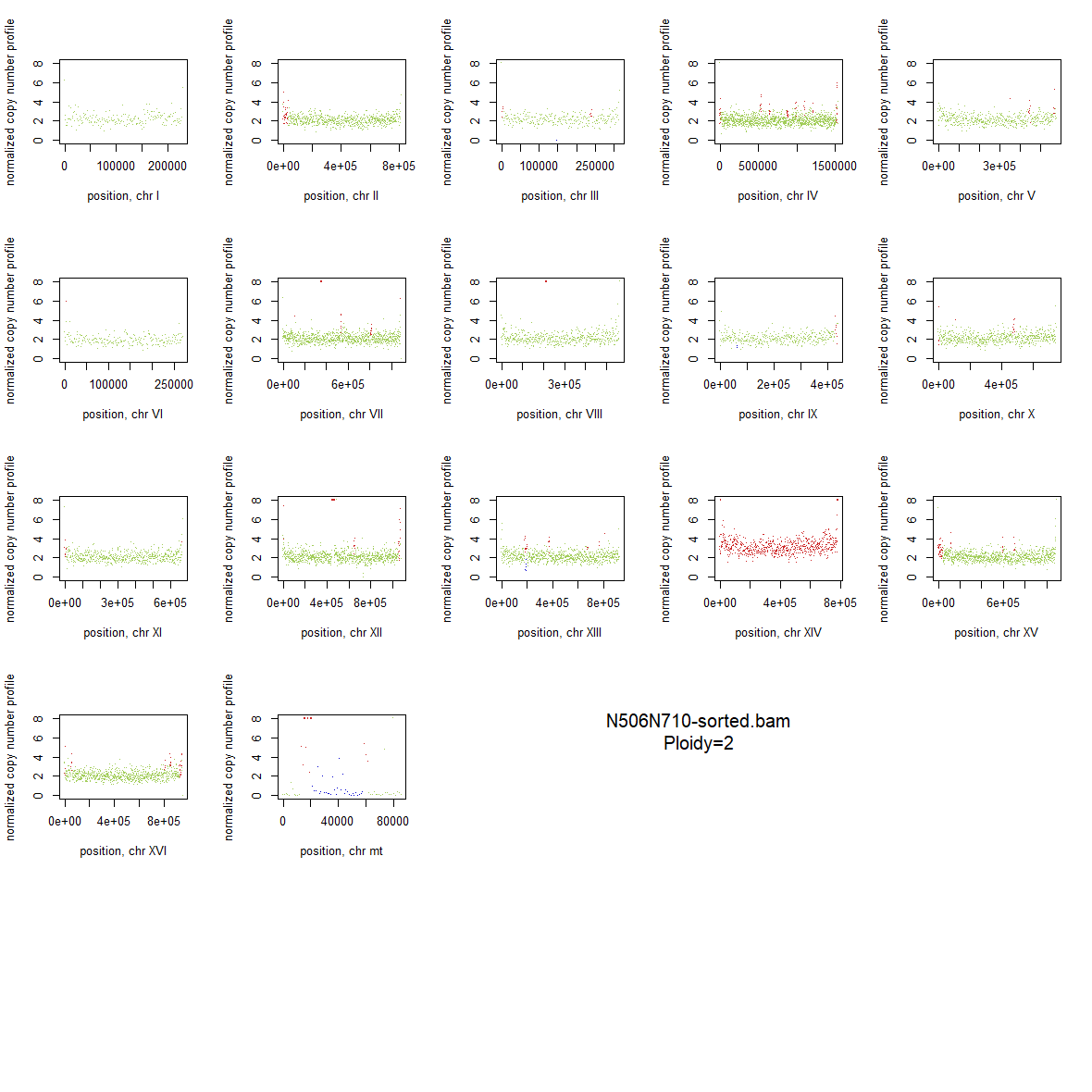

### 2x_V238M_57.png

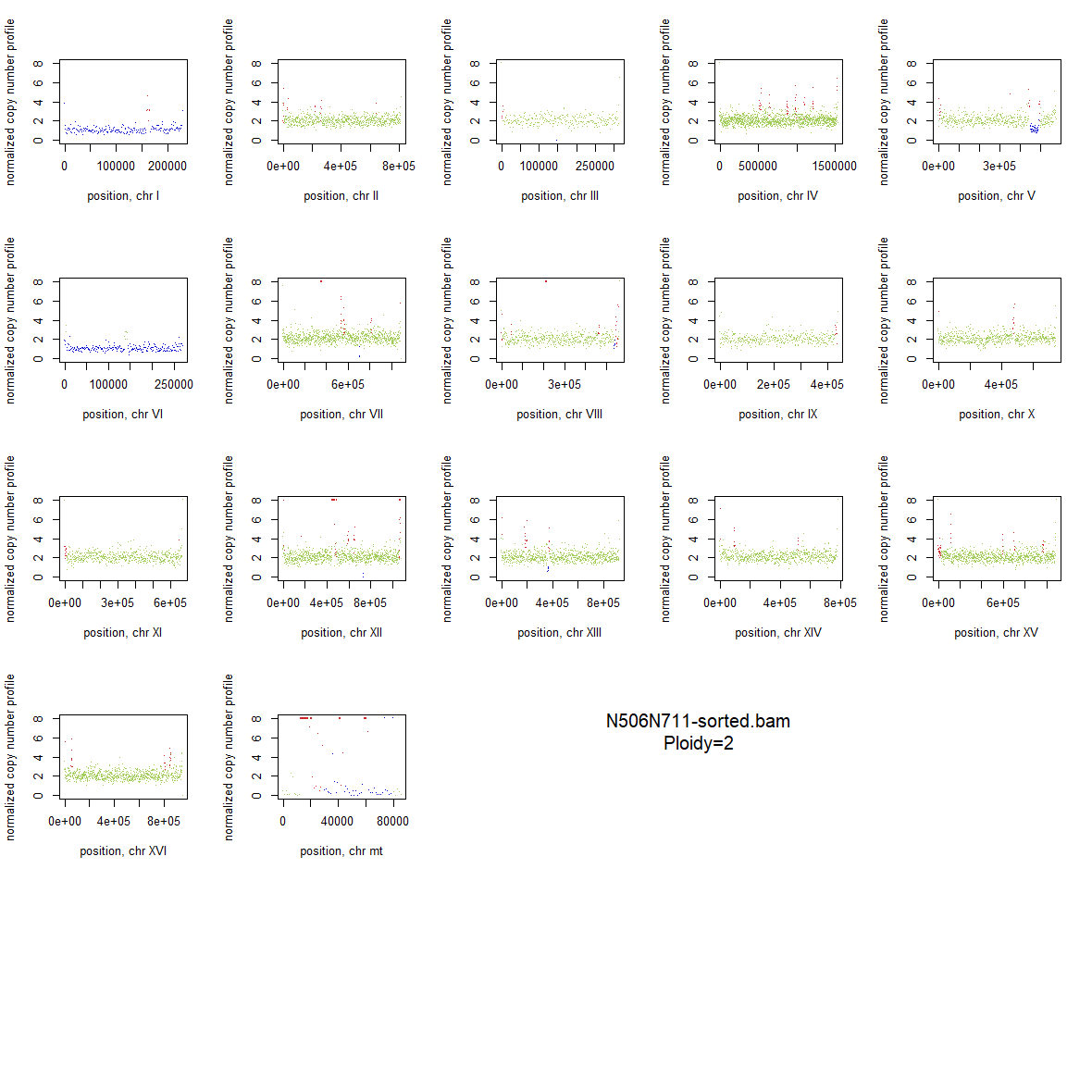

### 2x_V238M_58.png

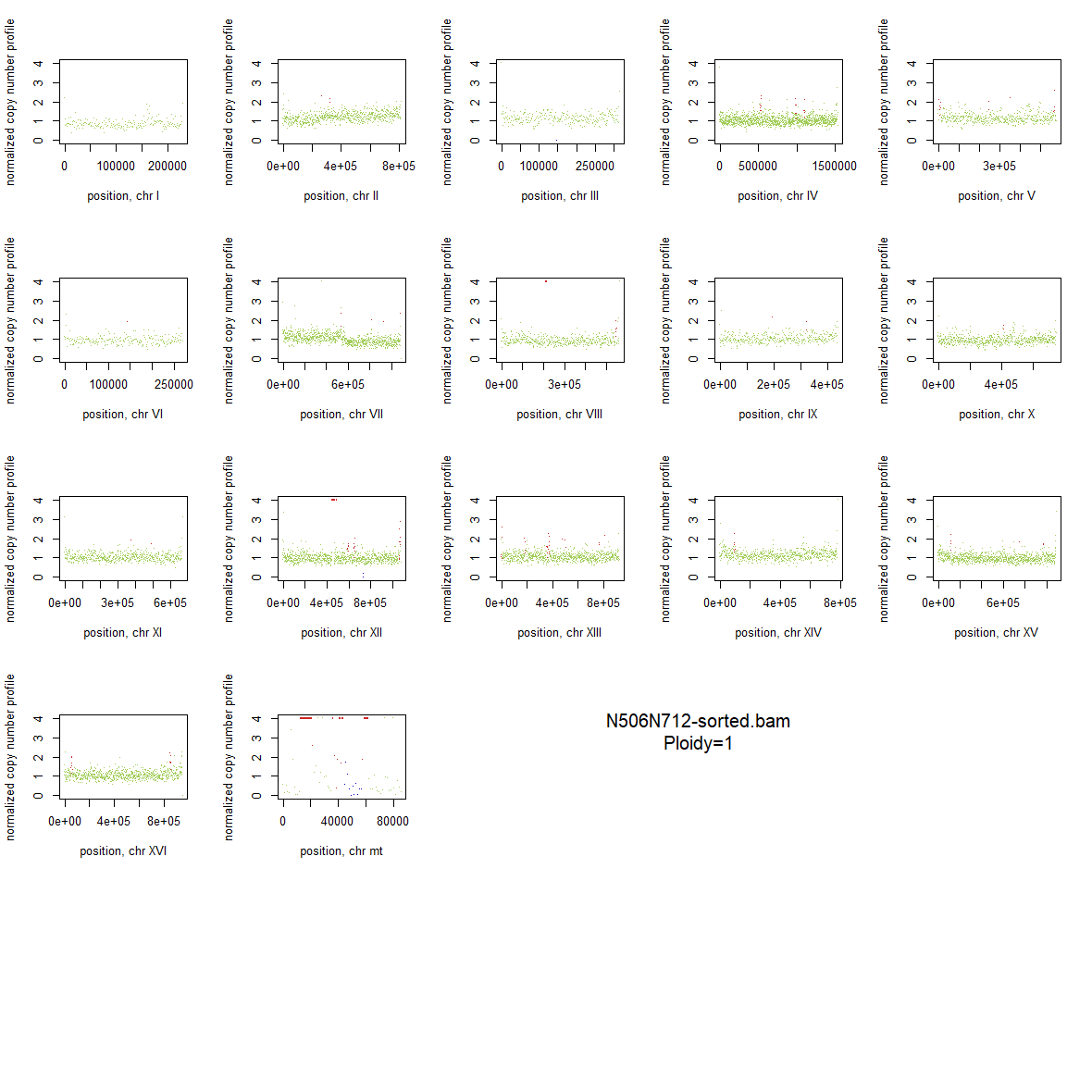

### 2x_V238M_59.png

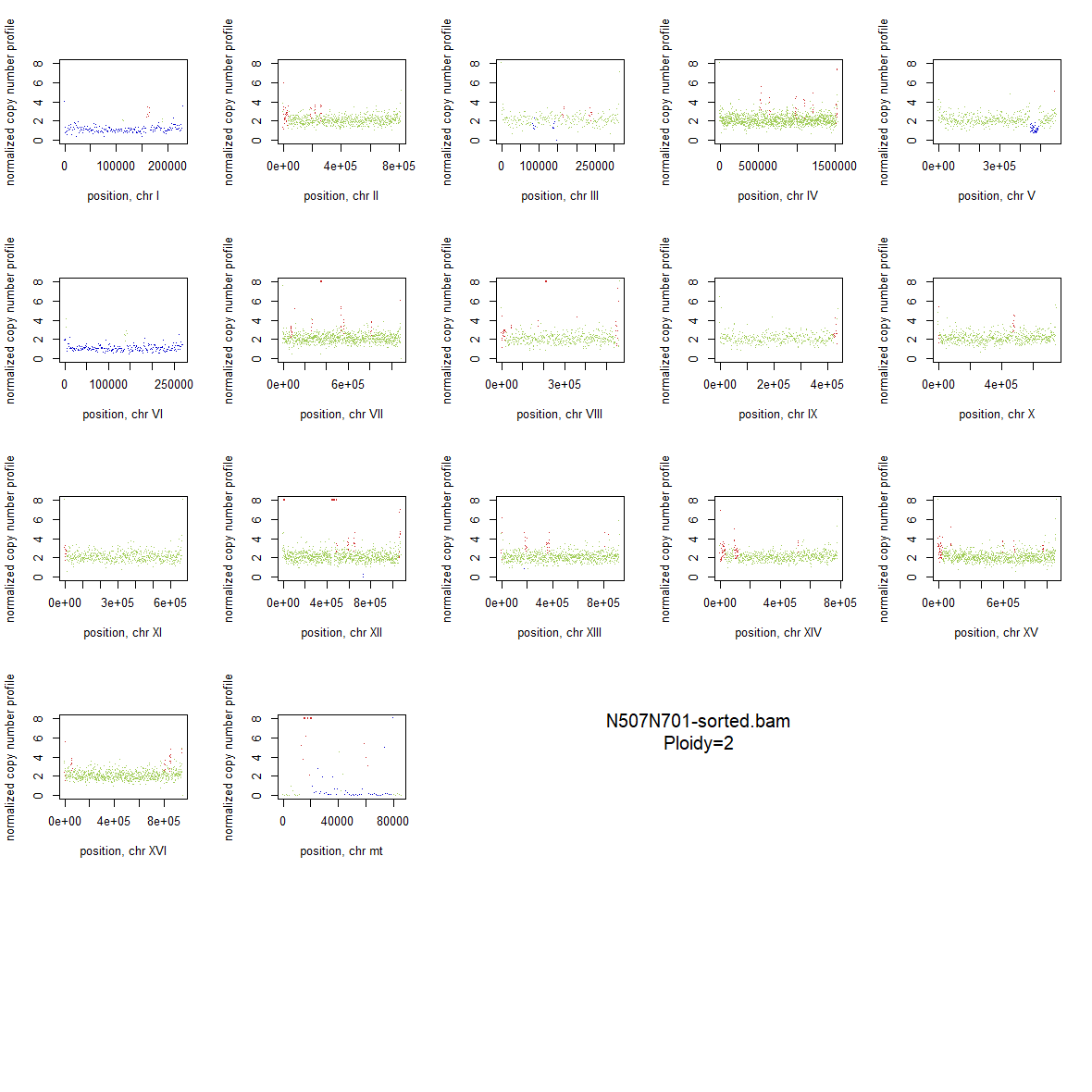

### 2x_V238M_60.png

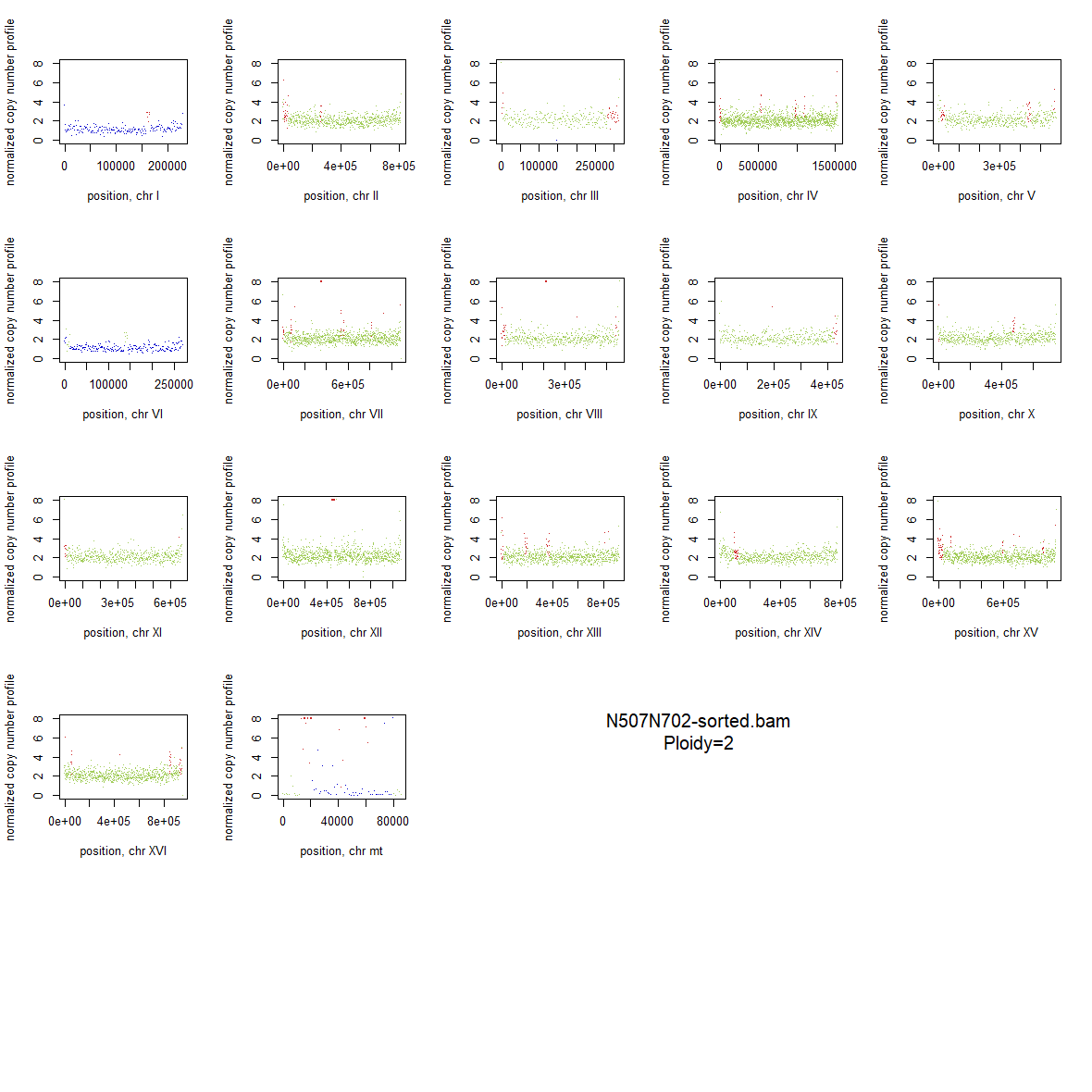

### 2x_V238M_61.png

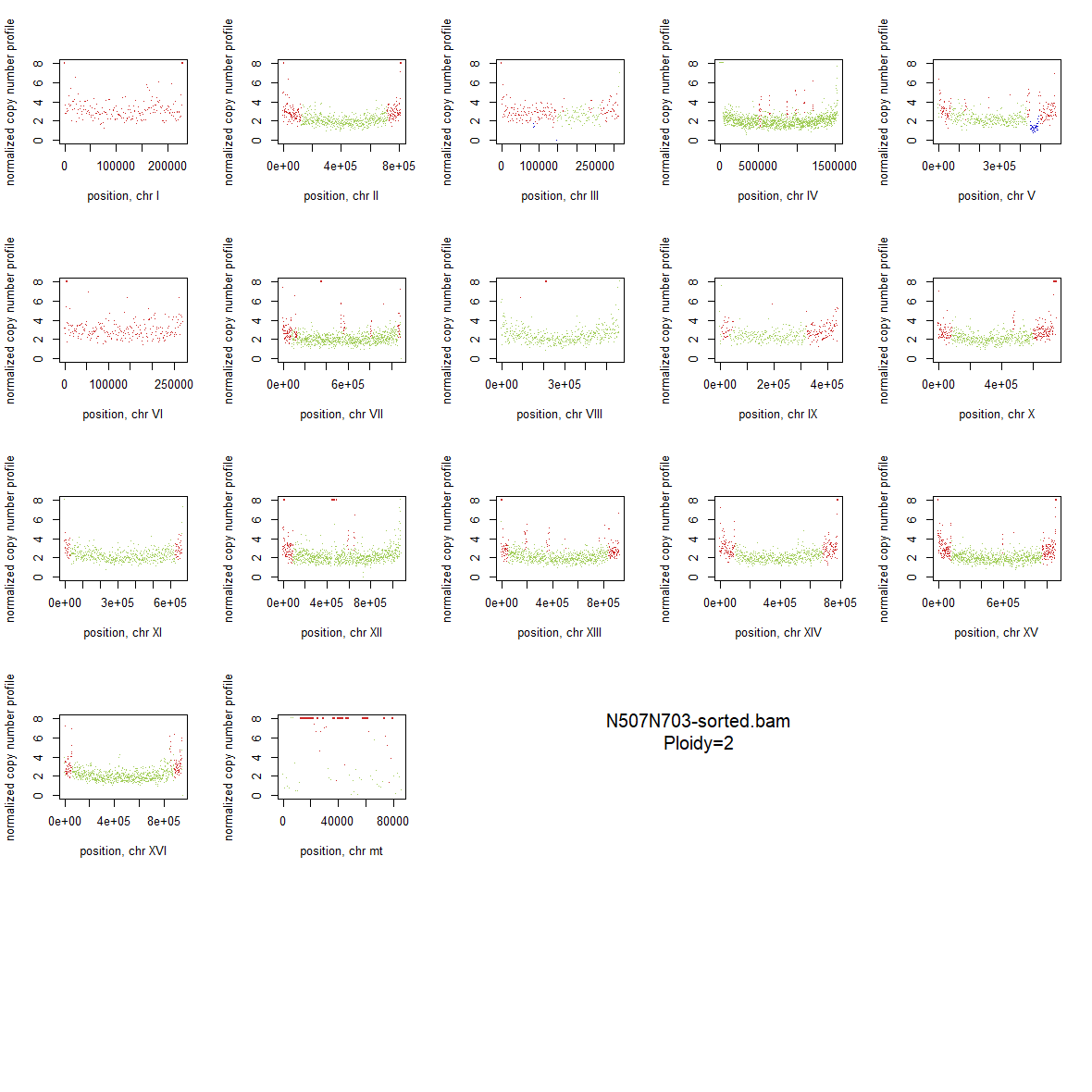

### 2x_V238M_62.png

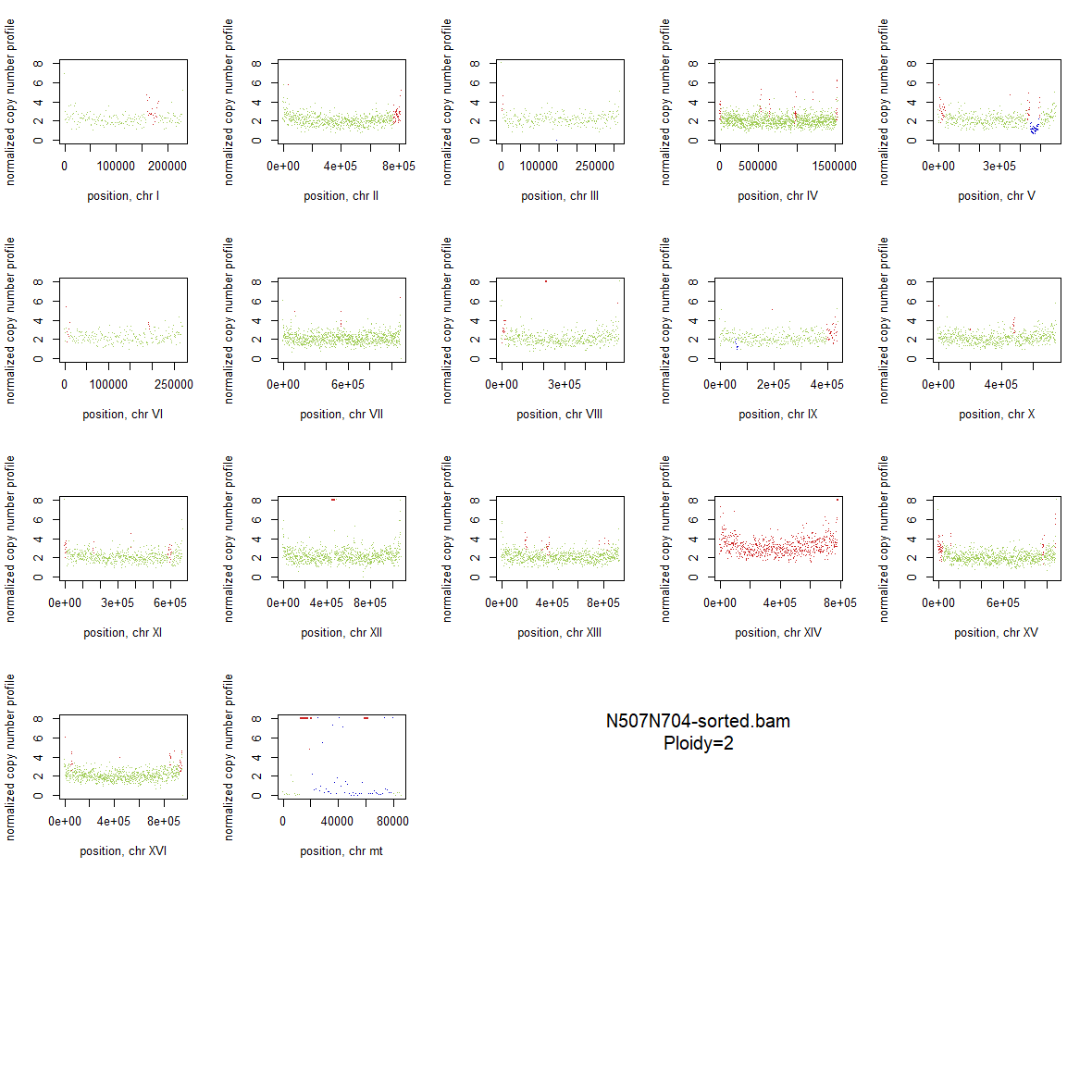

### 2x_V238M_63.png

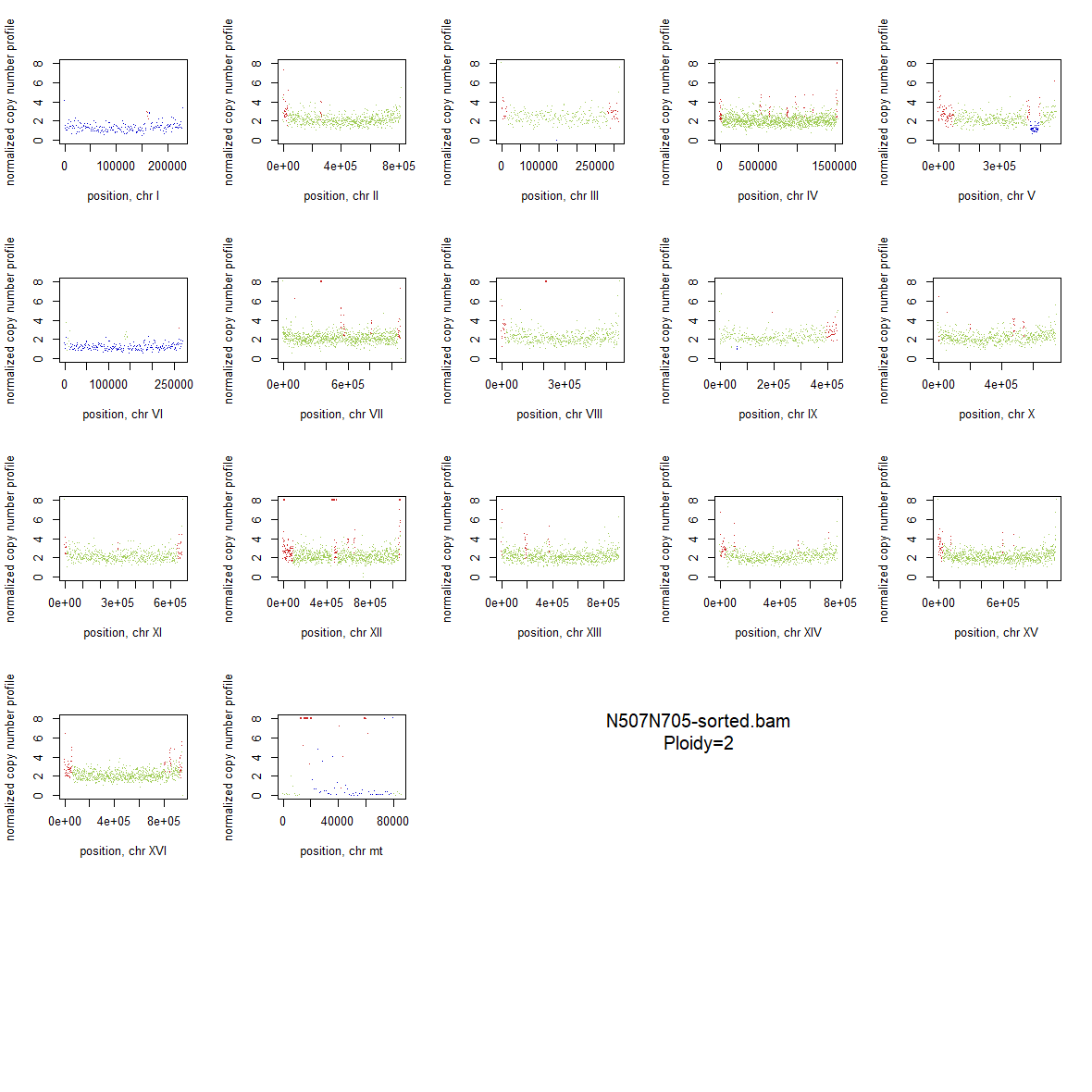

### 2x_V238M_64.png

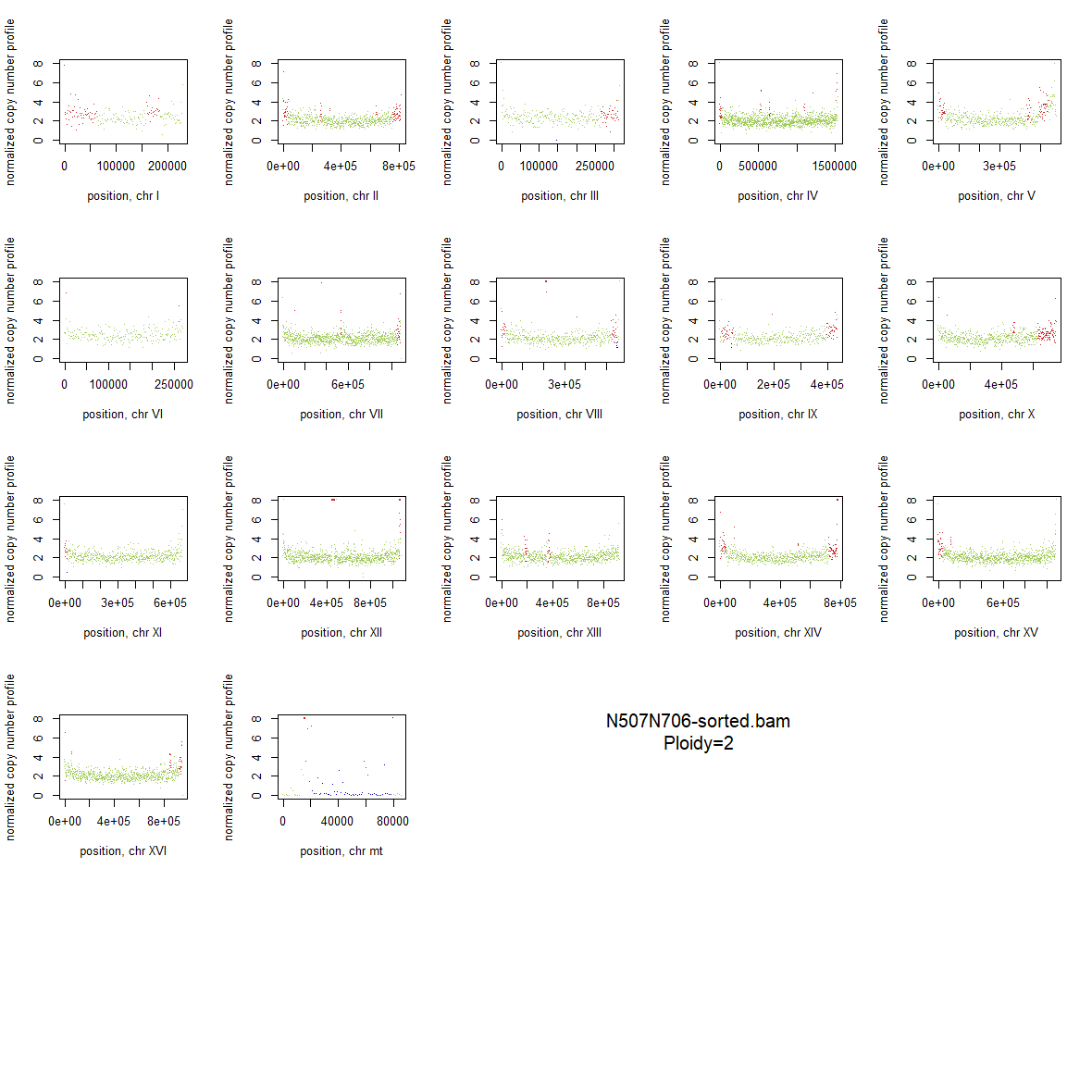

### 2x_V238M_65.png

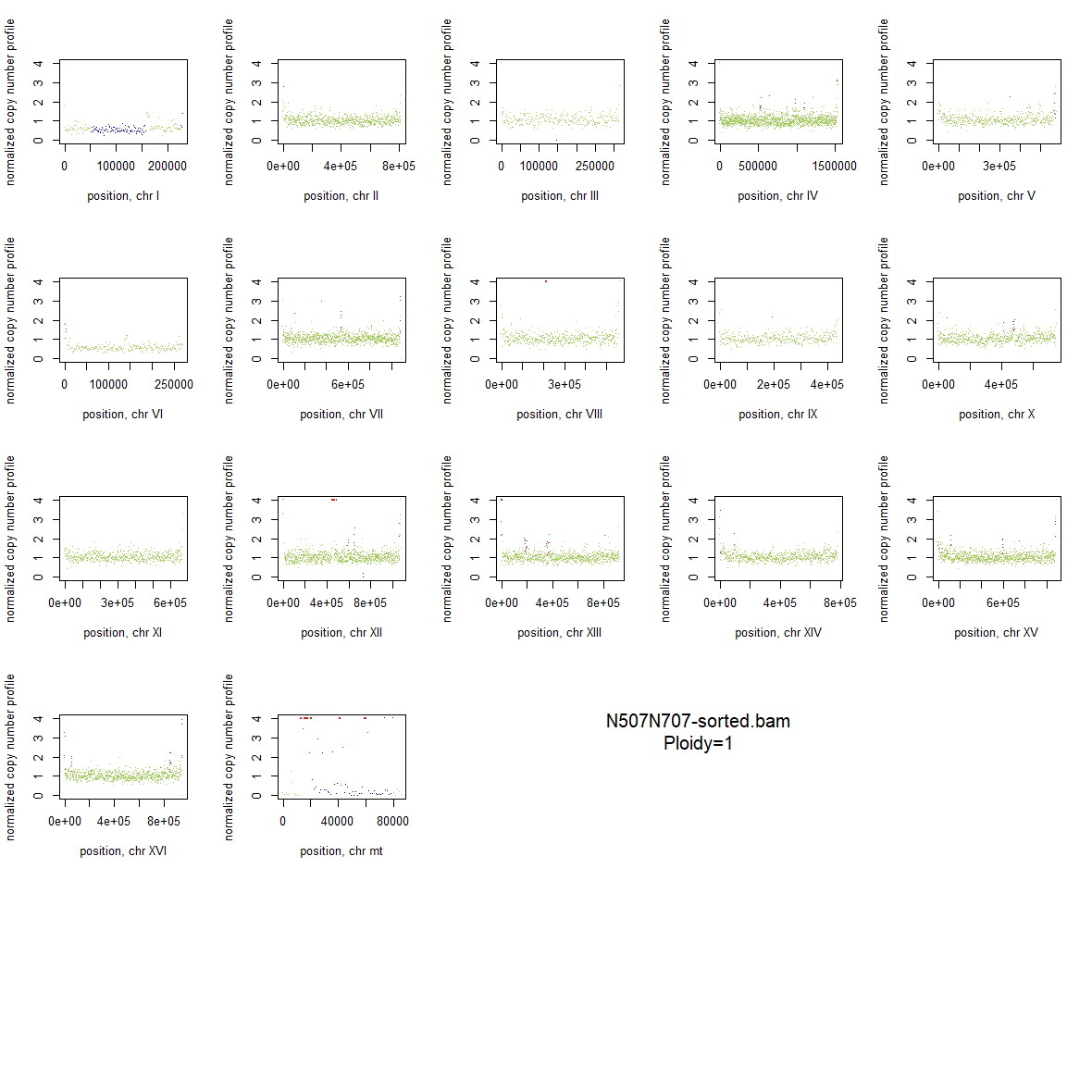

### 2x_V238M_66.png

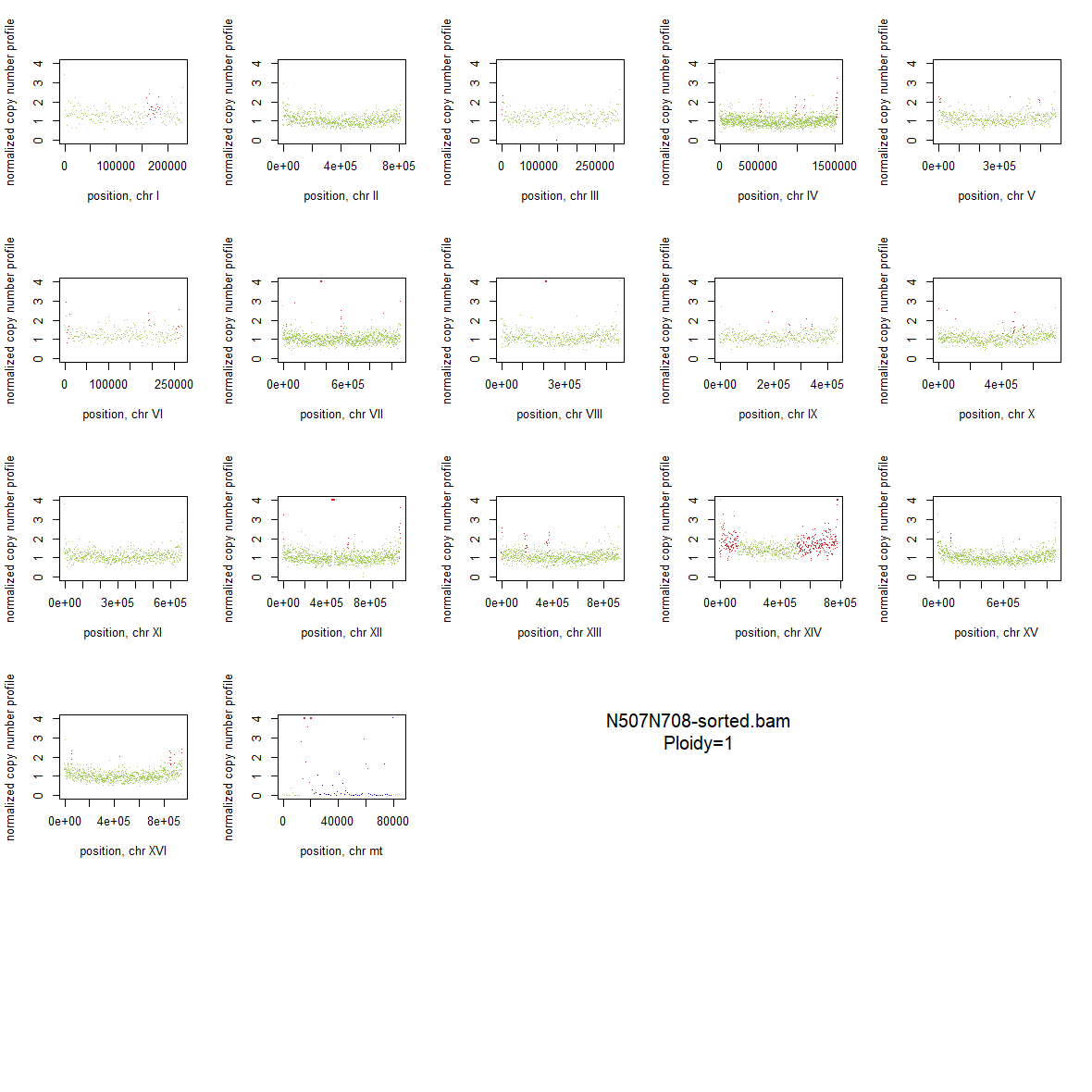

### 2x_V238M_67.png

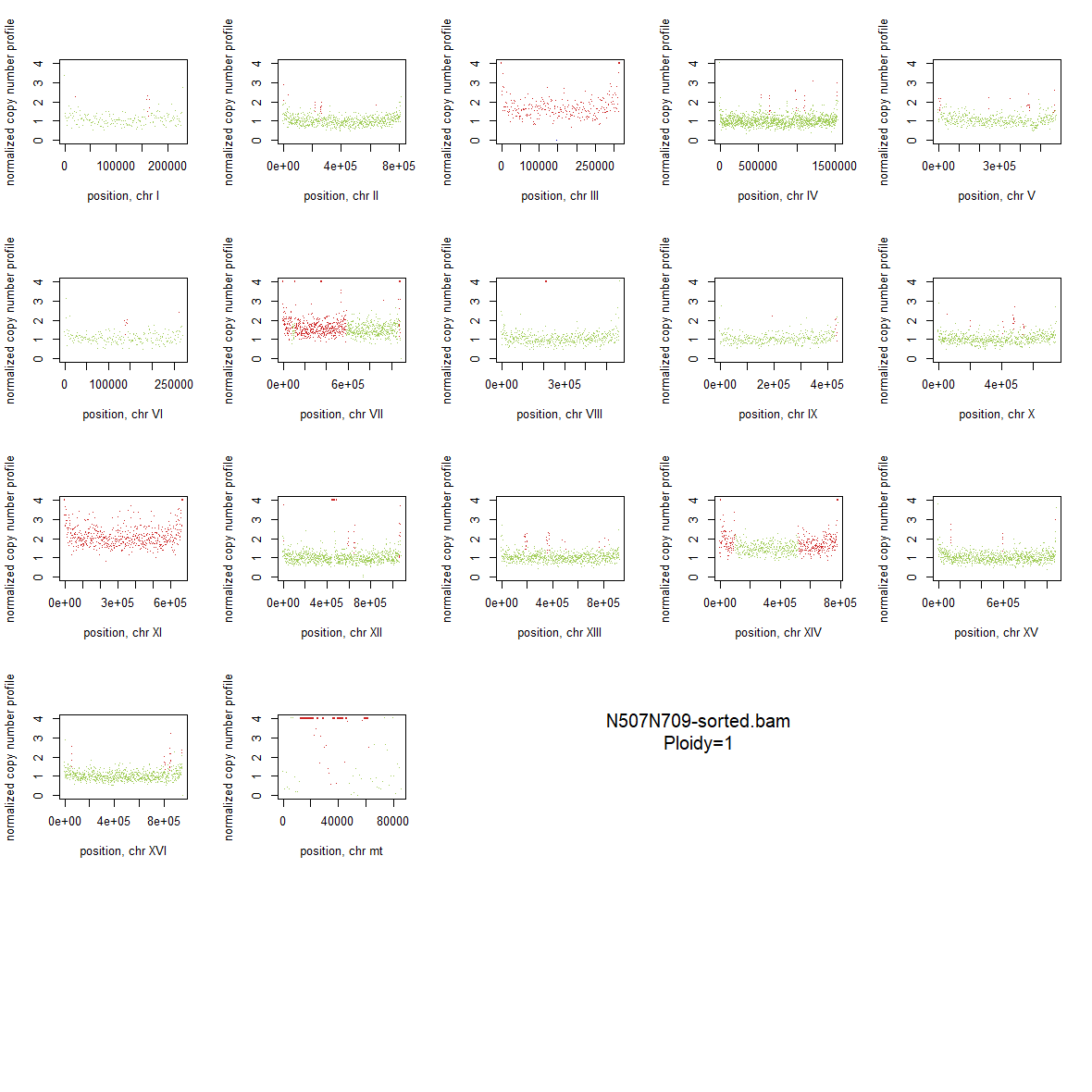

### 2x_V238M_68.png

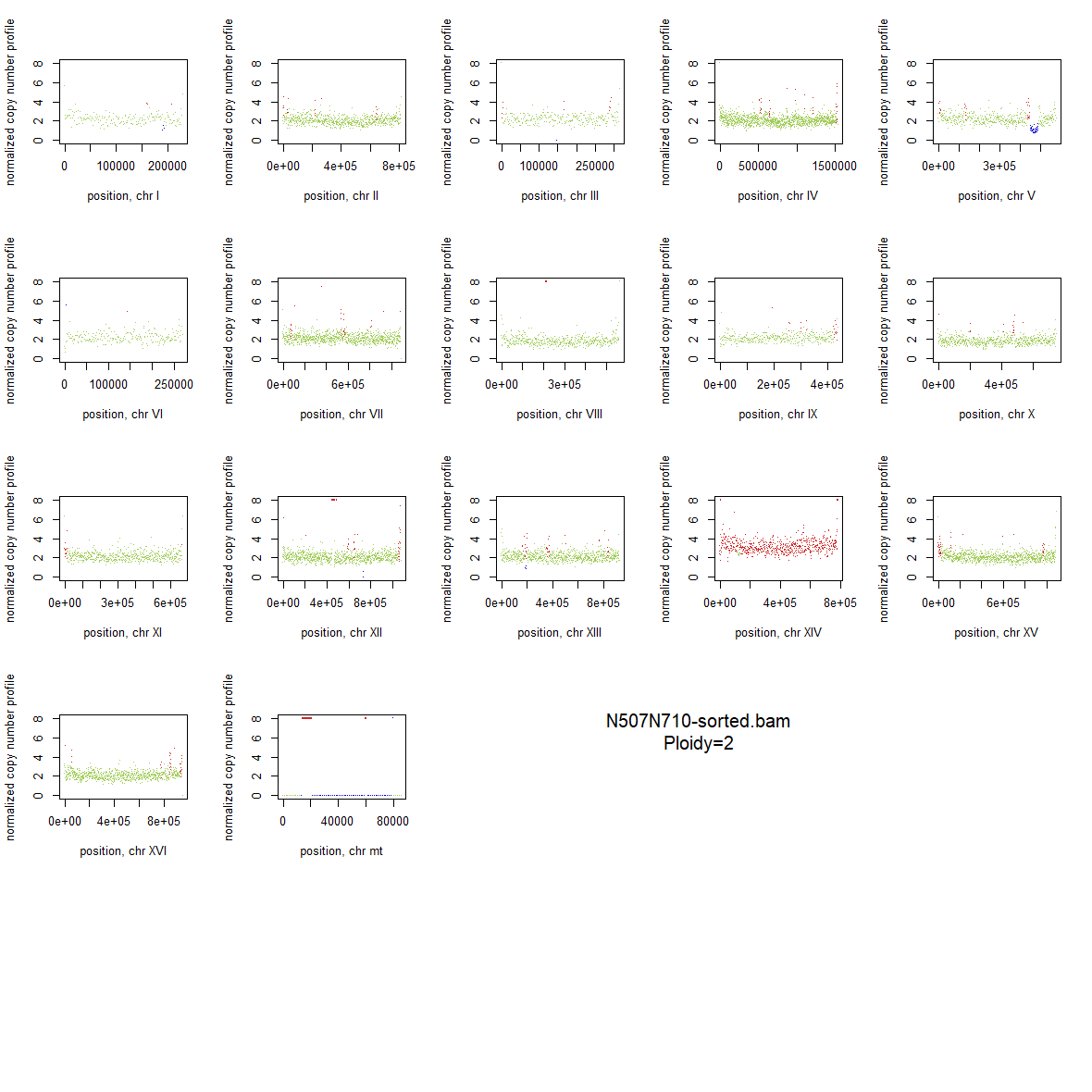

### 2x_V238M_69.png

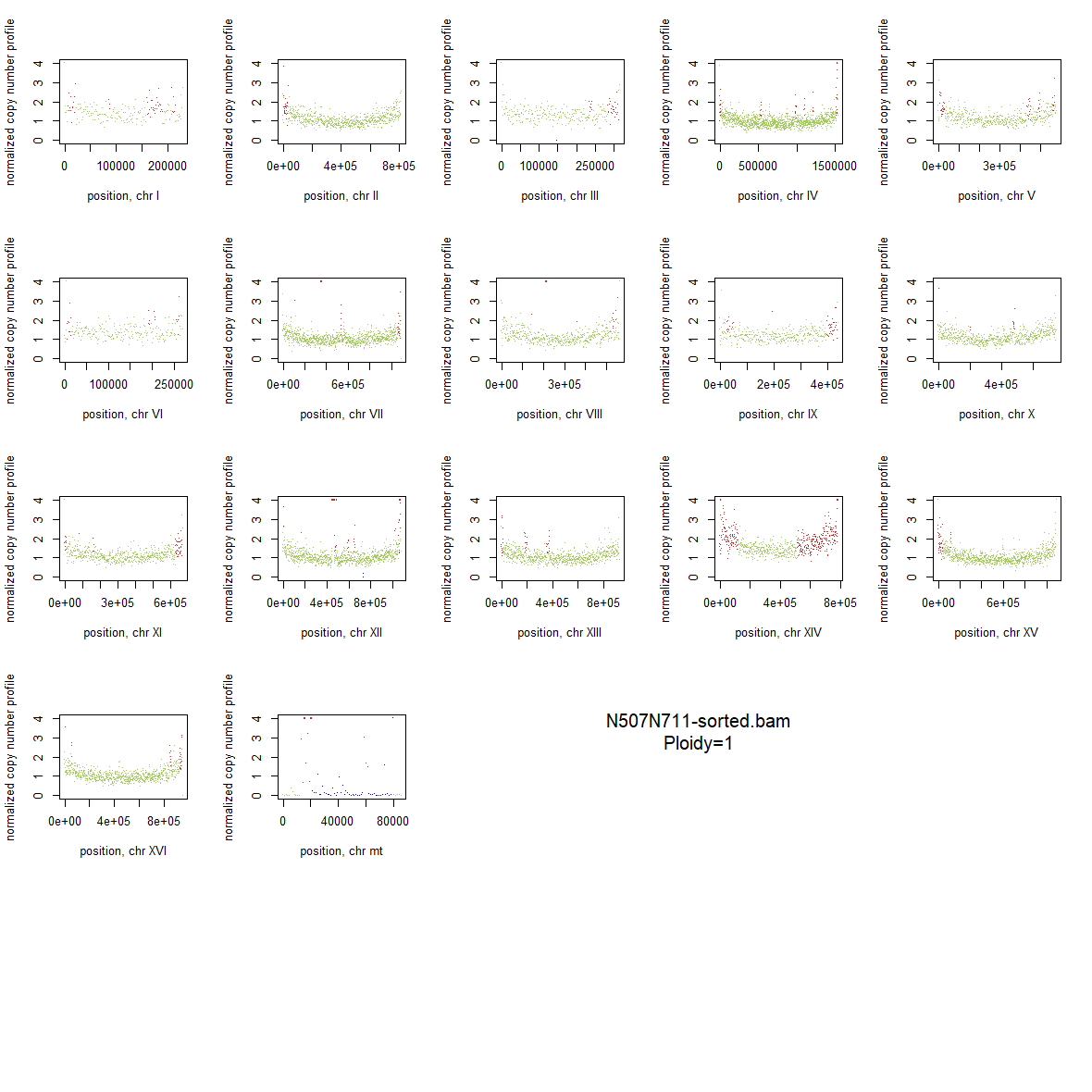

### 2x_V238M_70.png

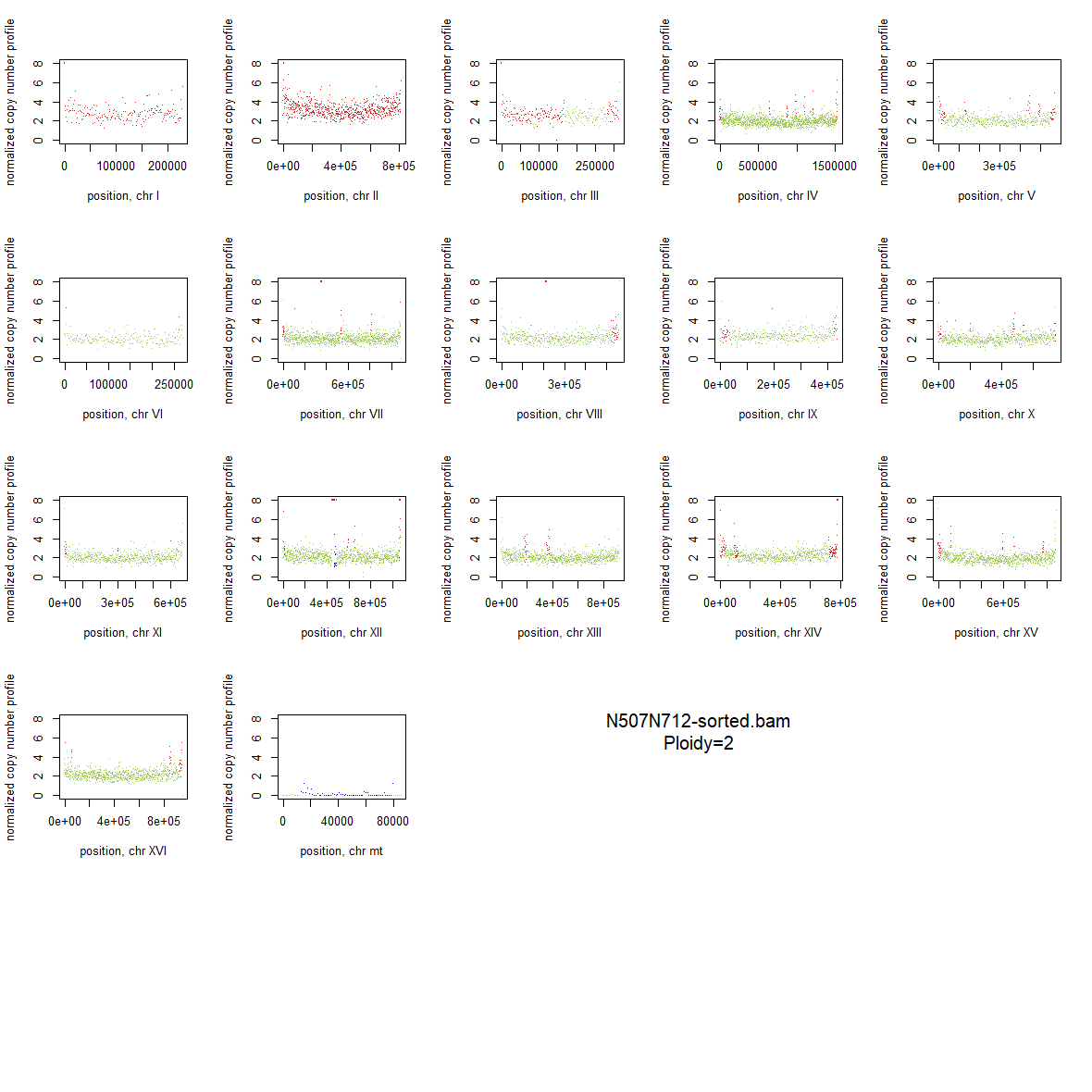

### 2x_V238M_71.png

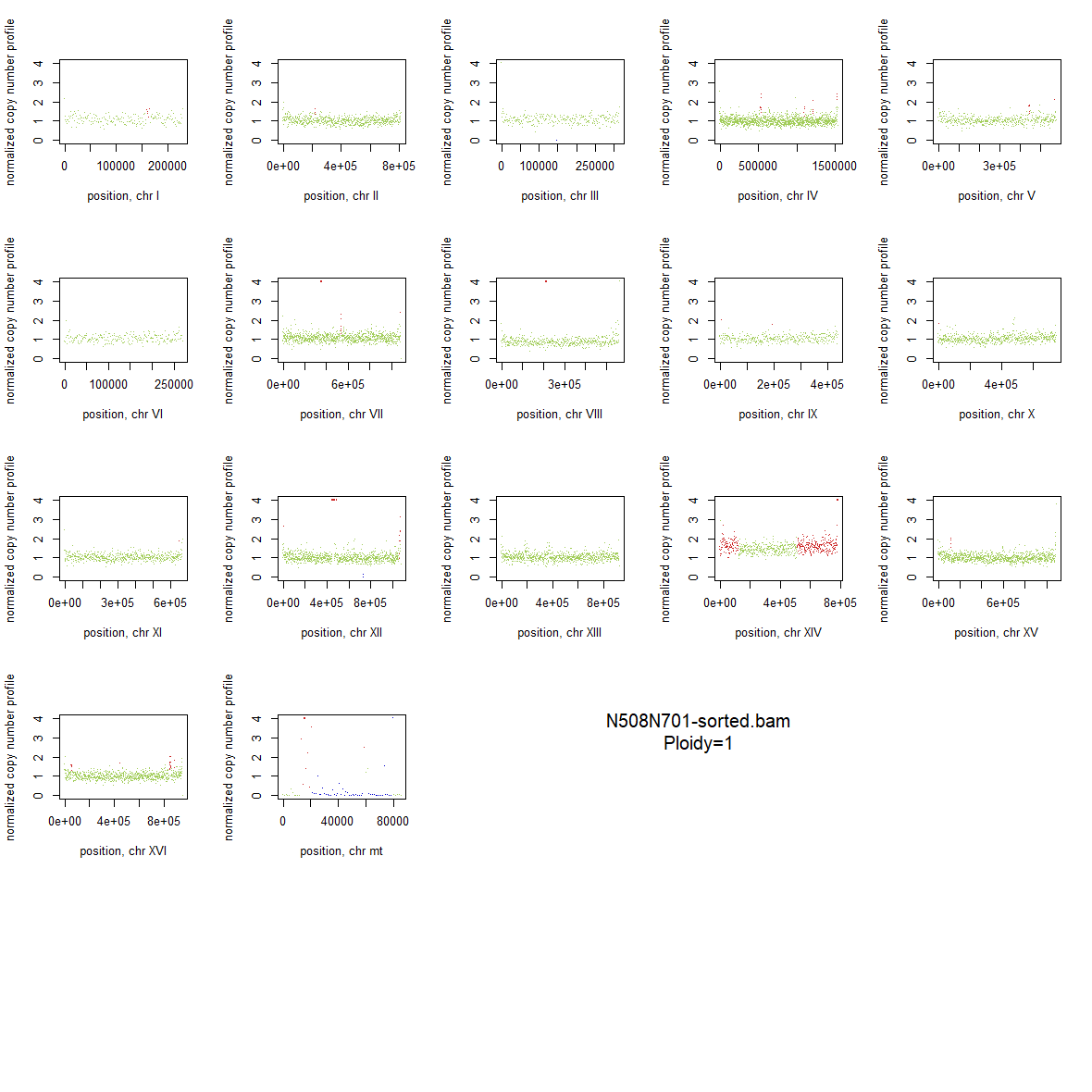

### 2x_V238M_72.png

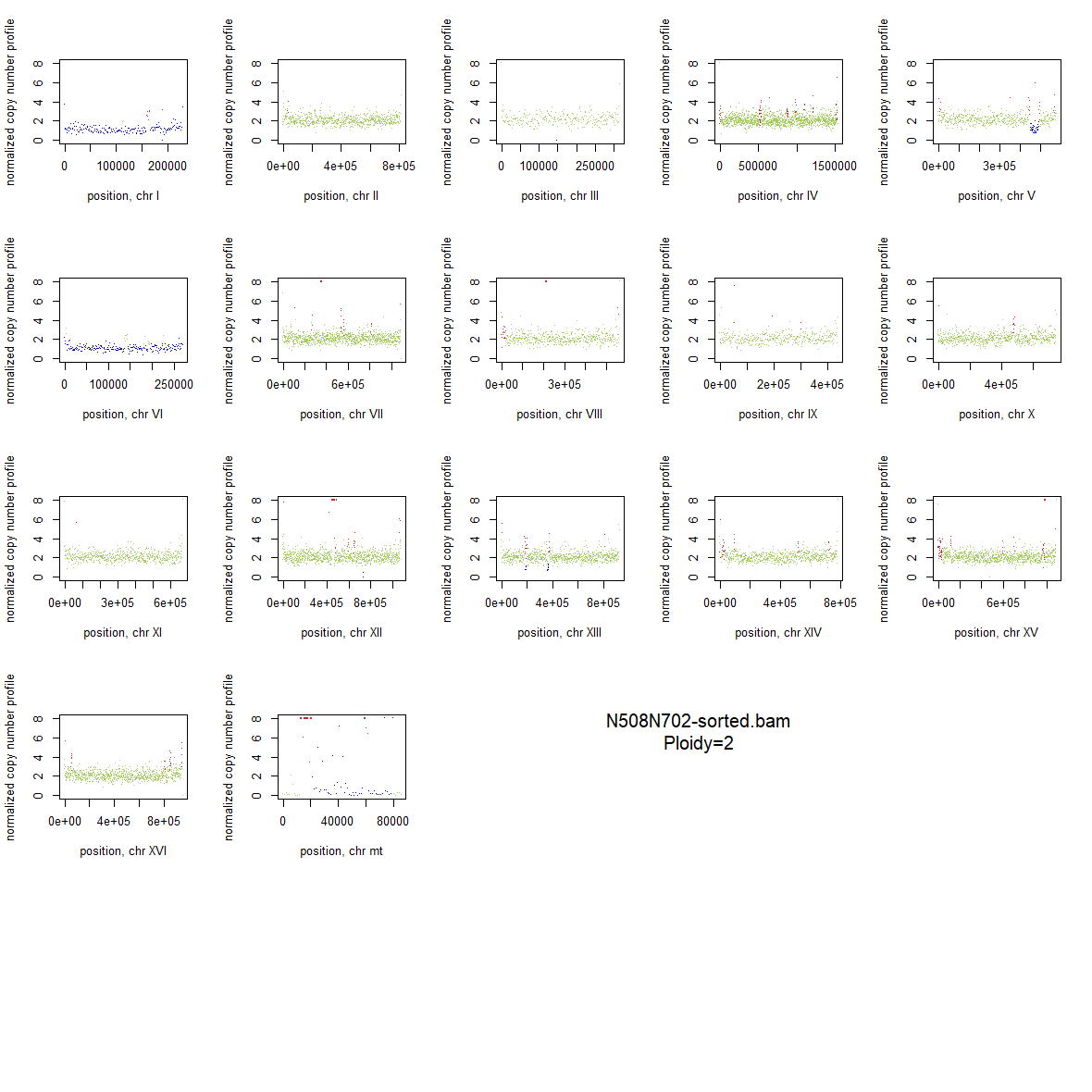

### 2x_V238M_73.png

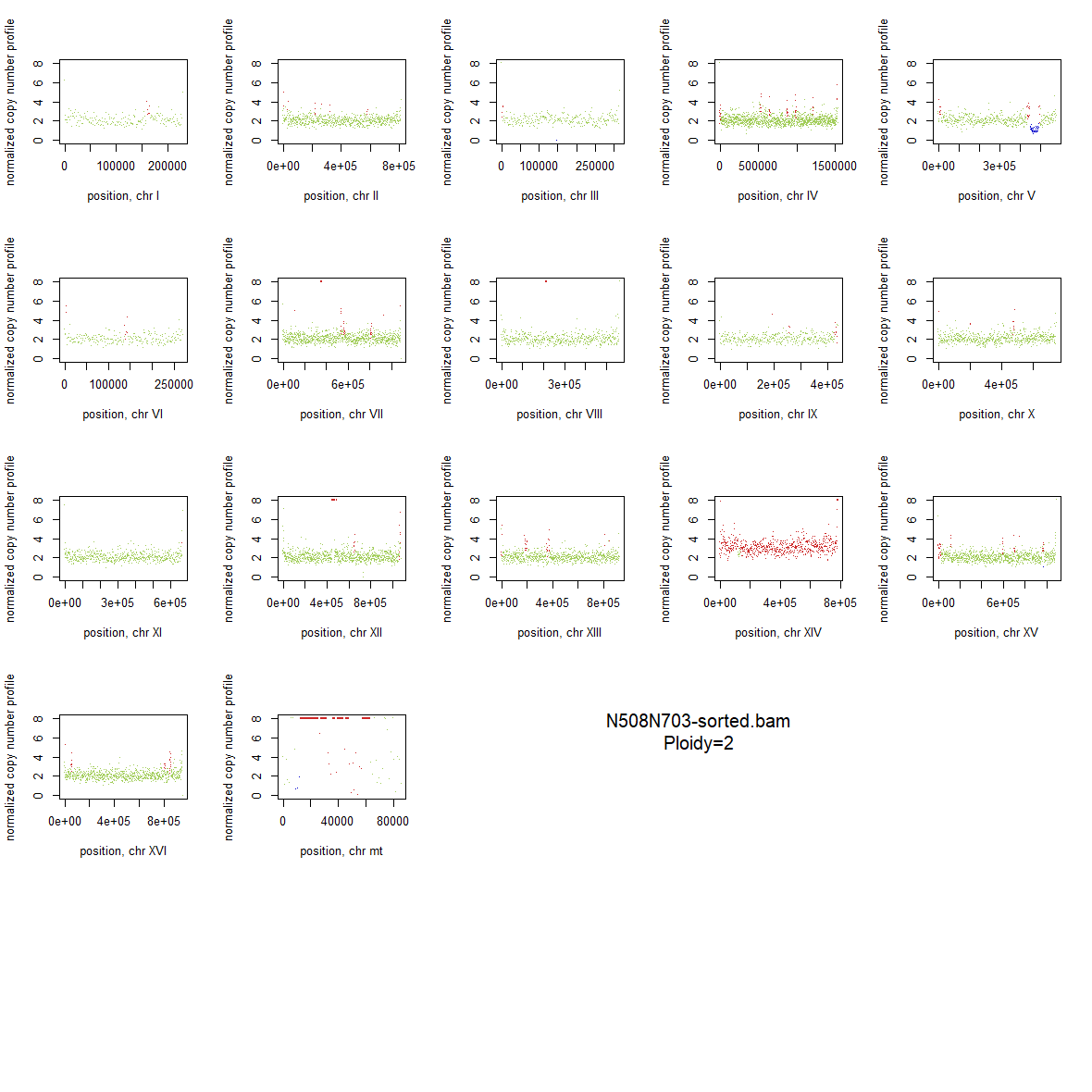

### 2x_V238M_74.png

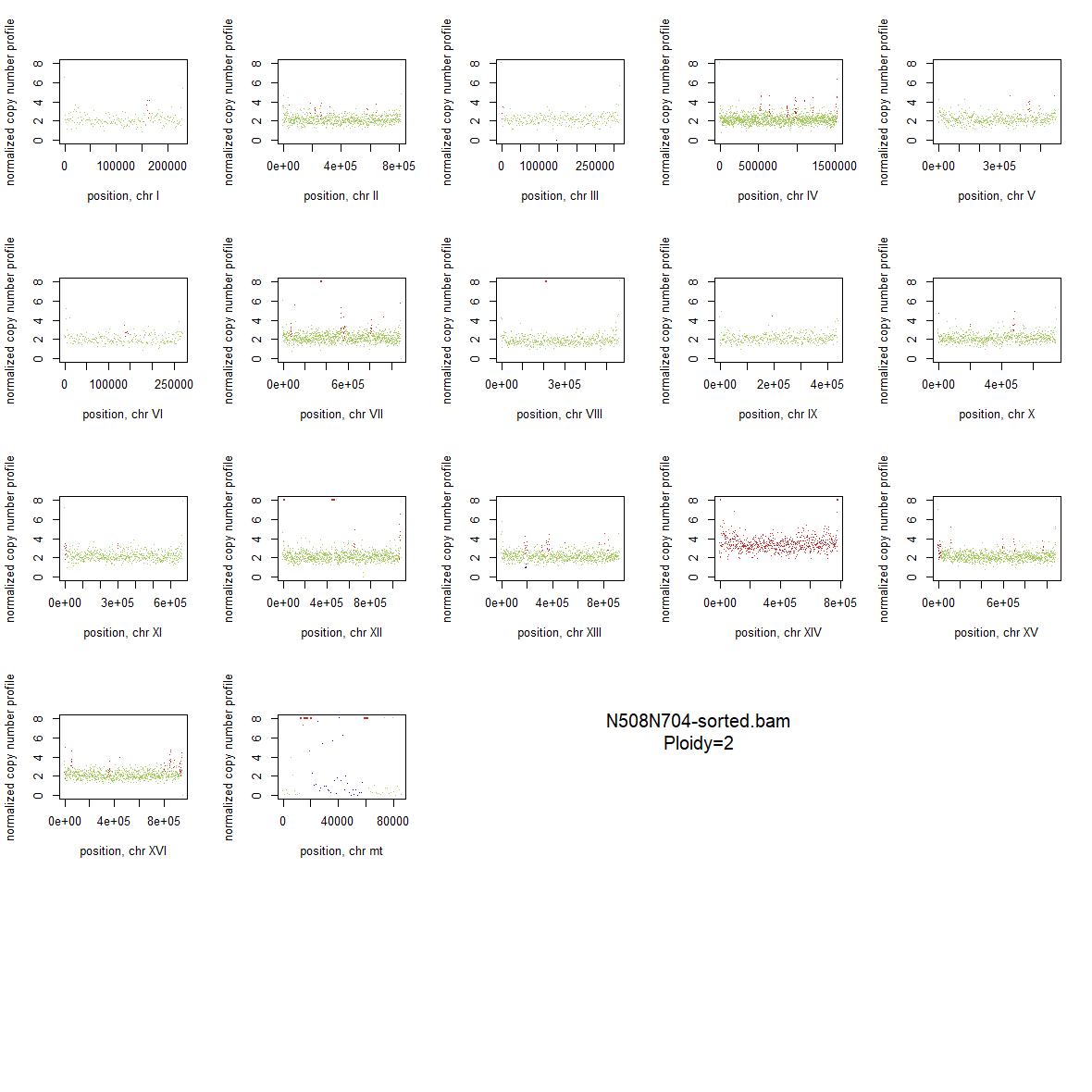

### 2x_V238M_75.png

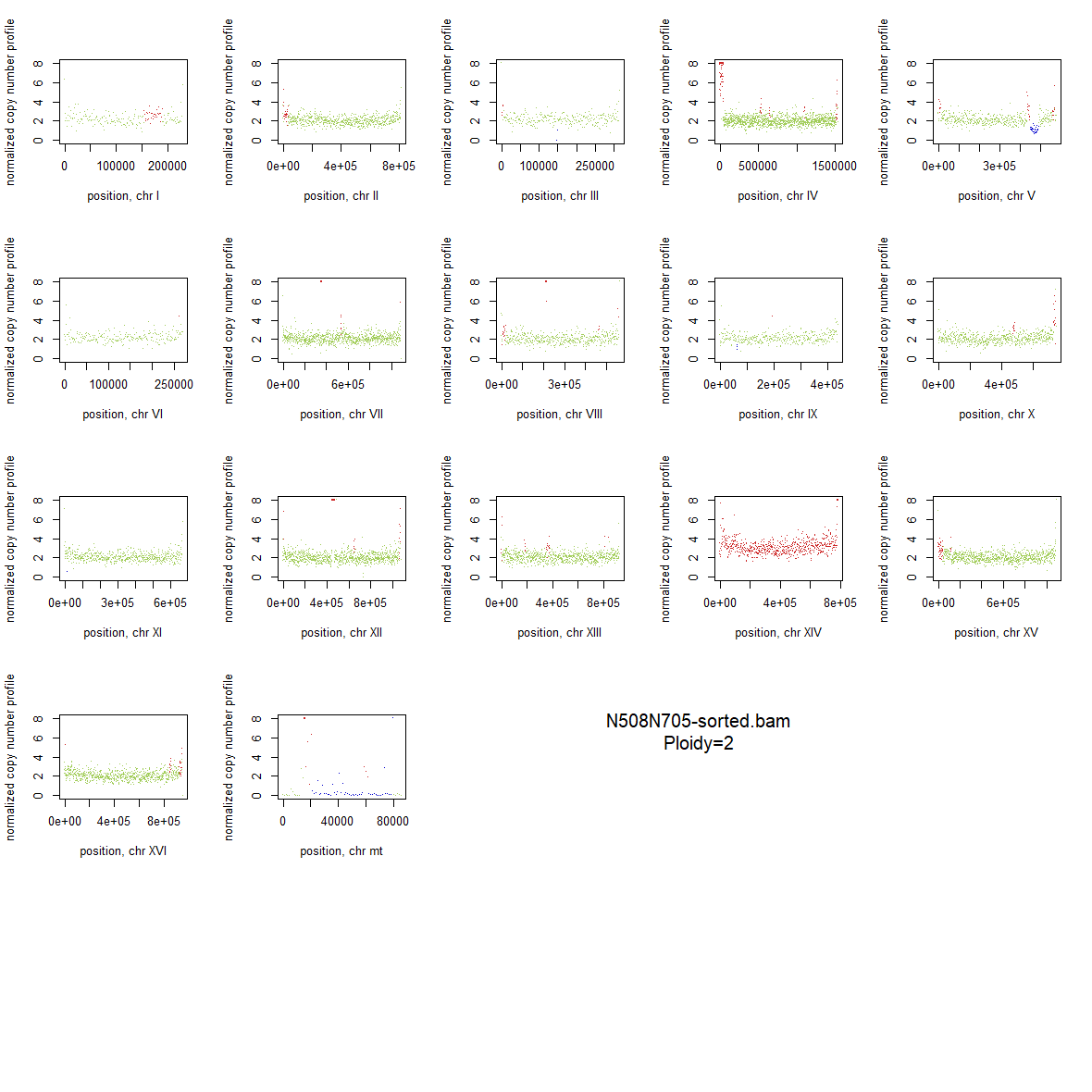

### 2x_V238M_76.png

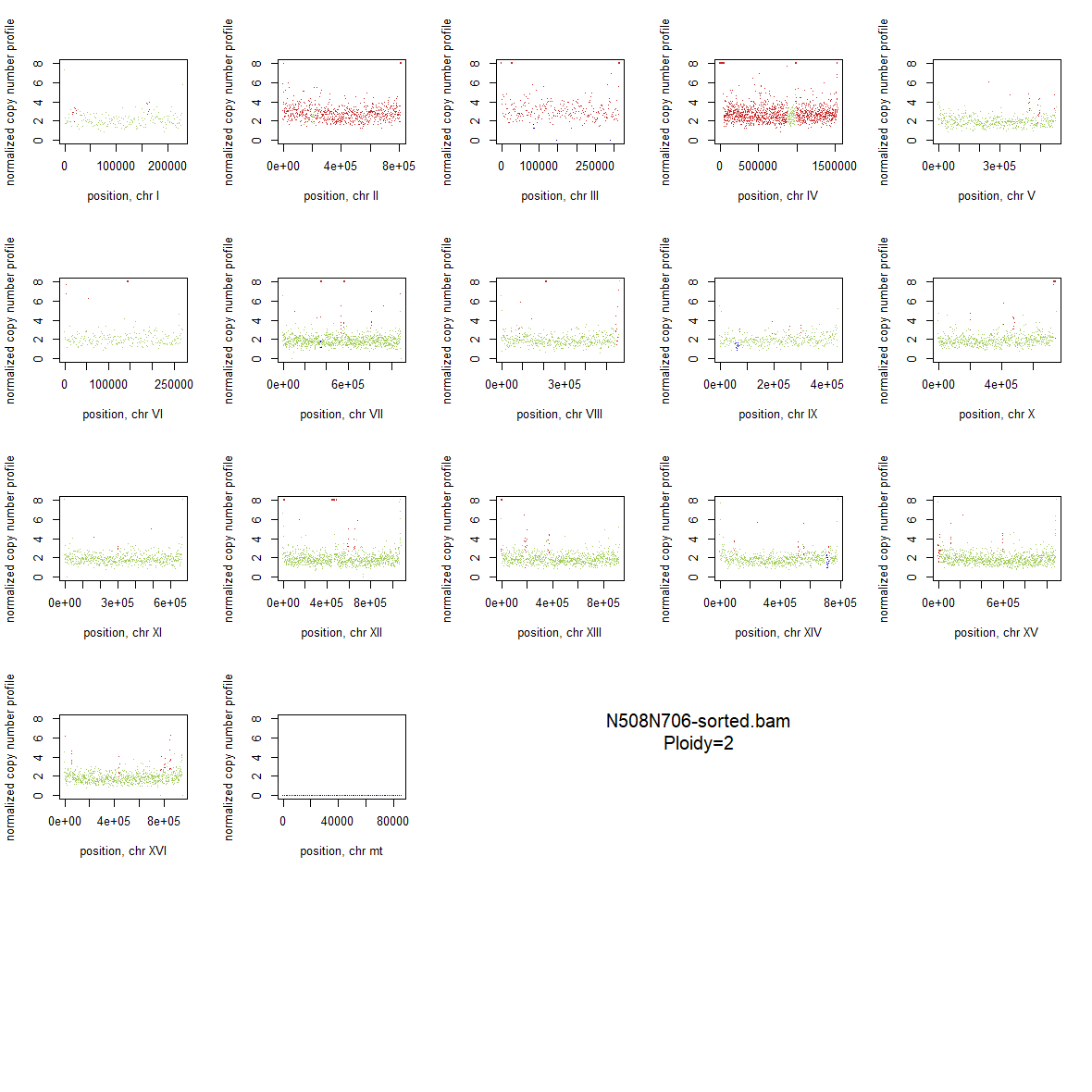

### 2x_V238M_77.png

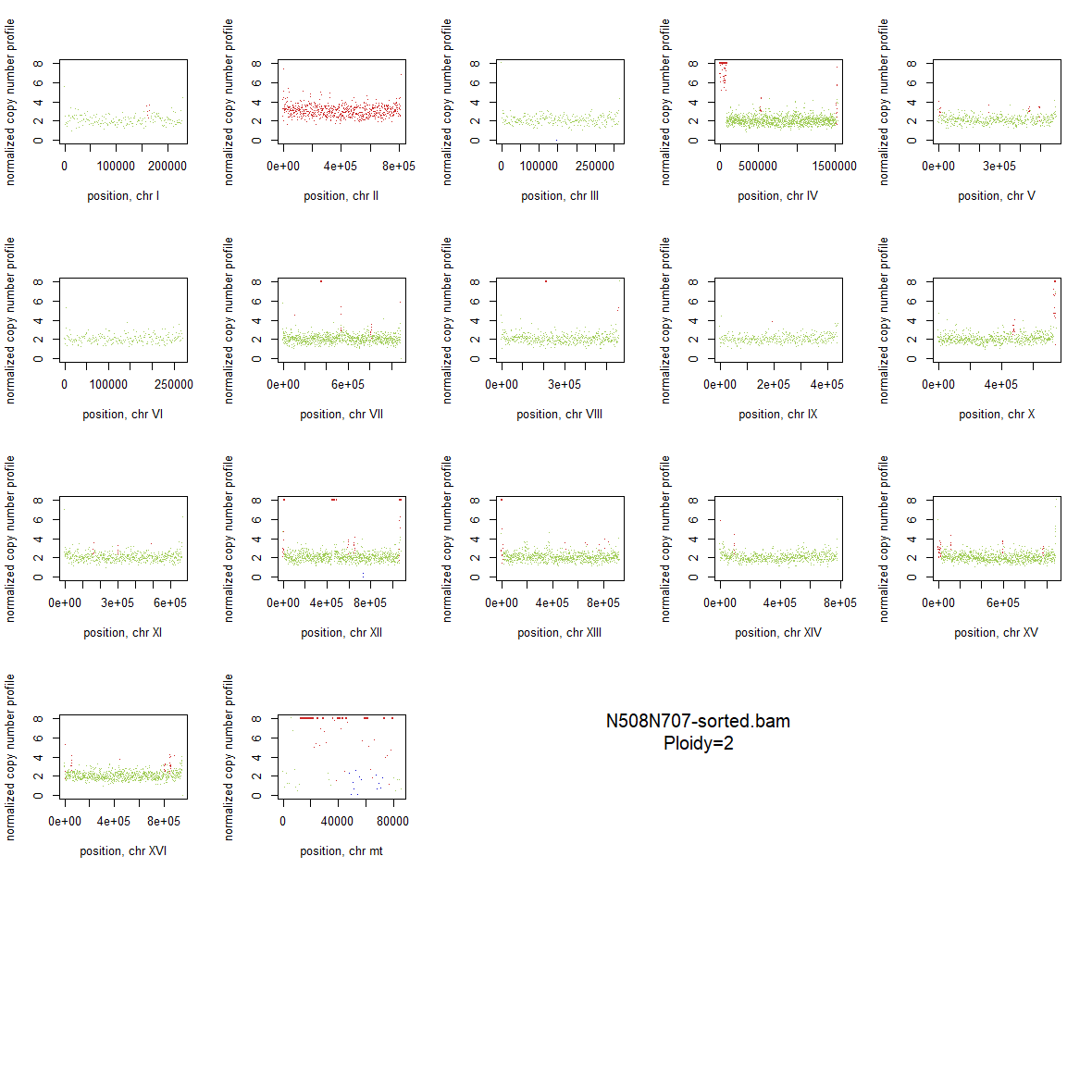

### 2x_V238M_78.png

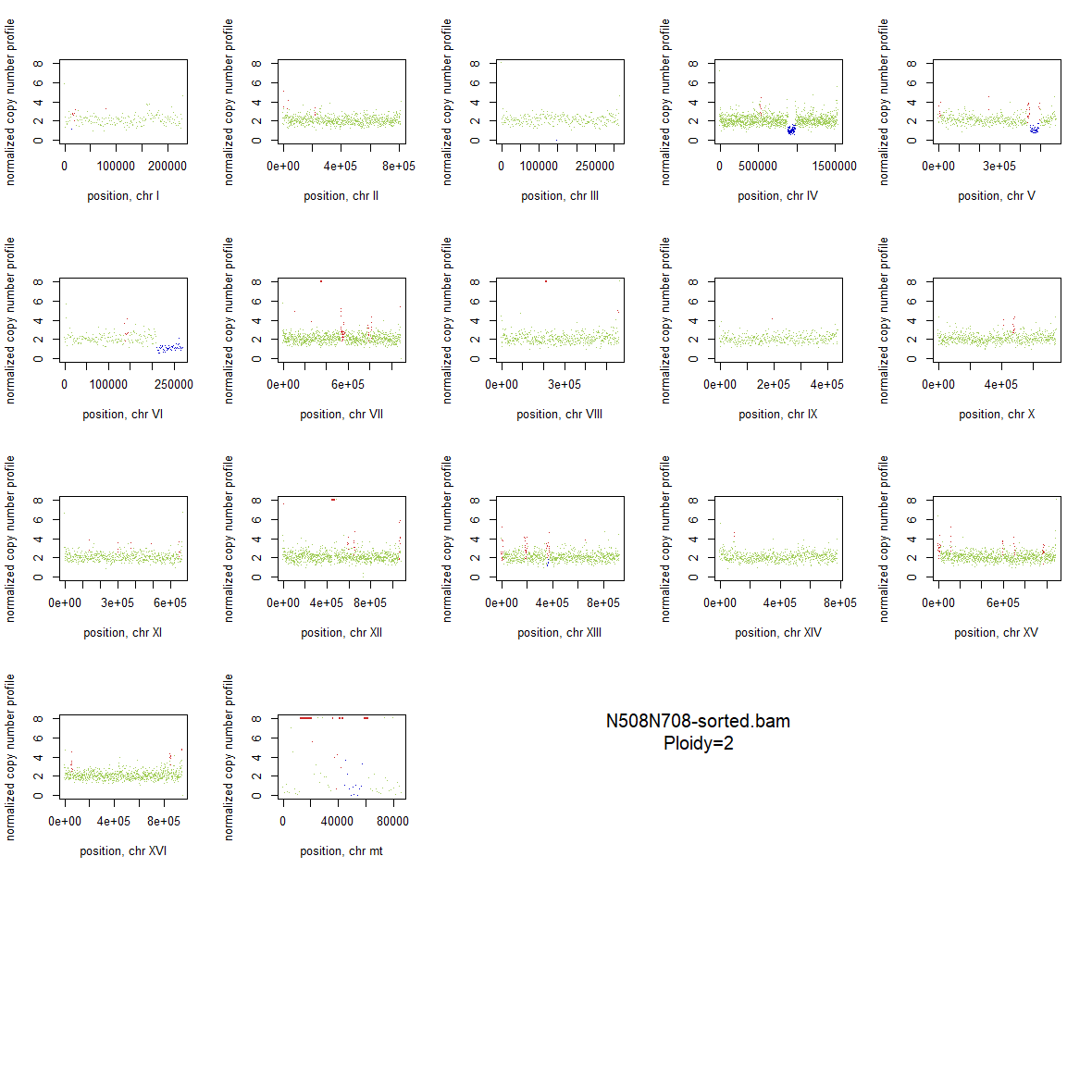

### 2x_V238M_79.png

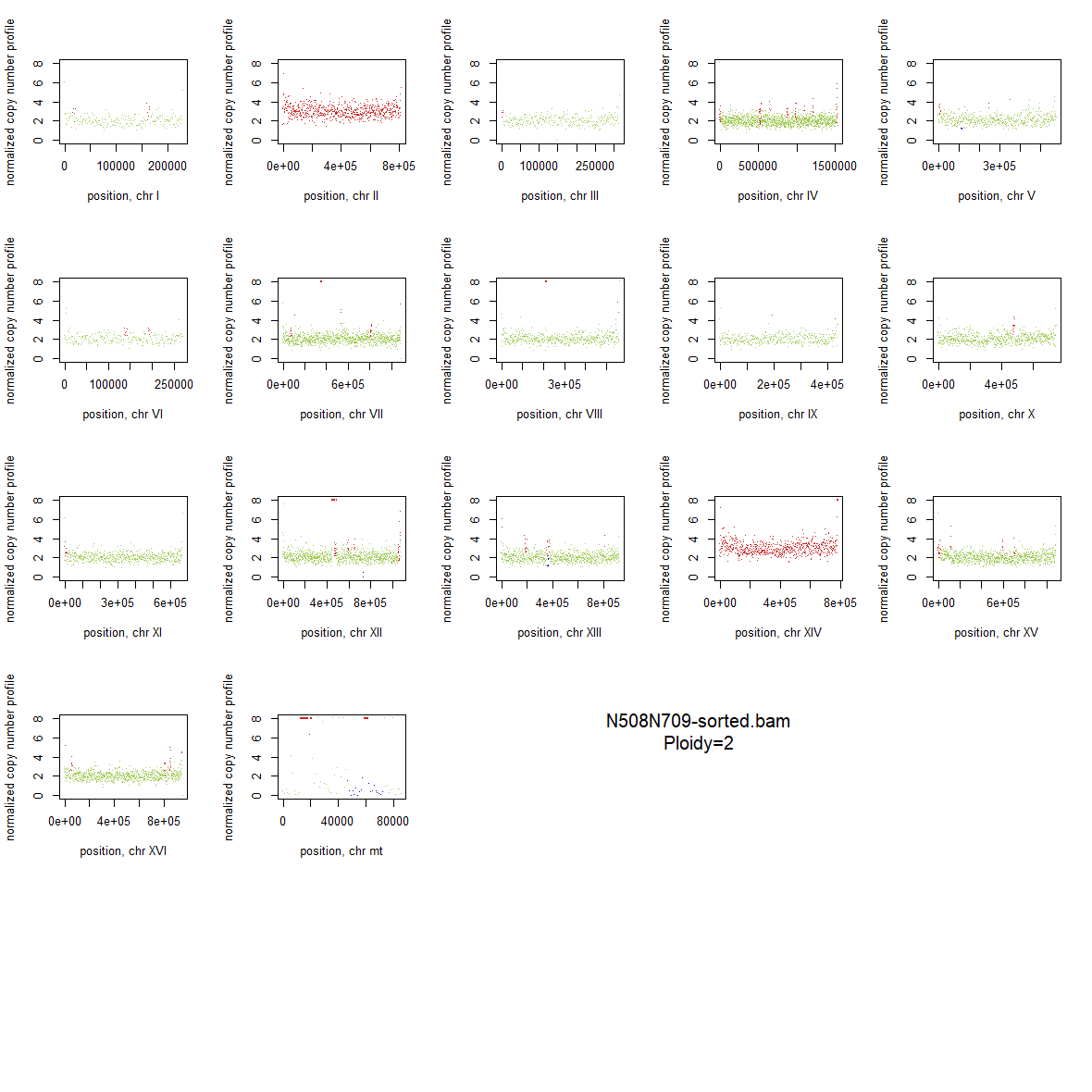

### 2x_V238M_80.png

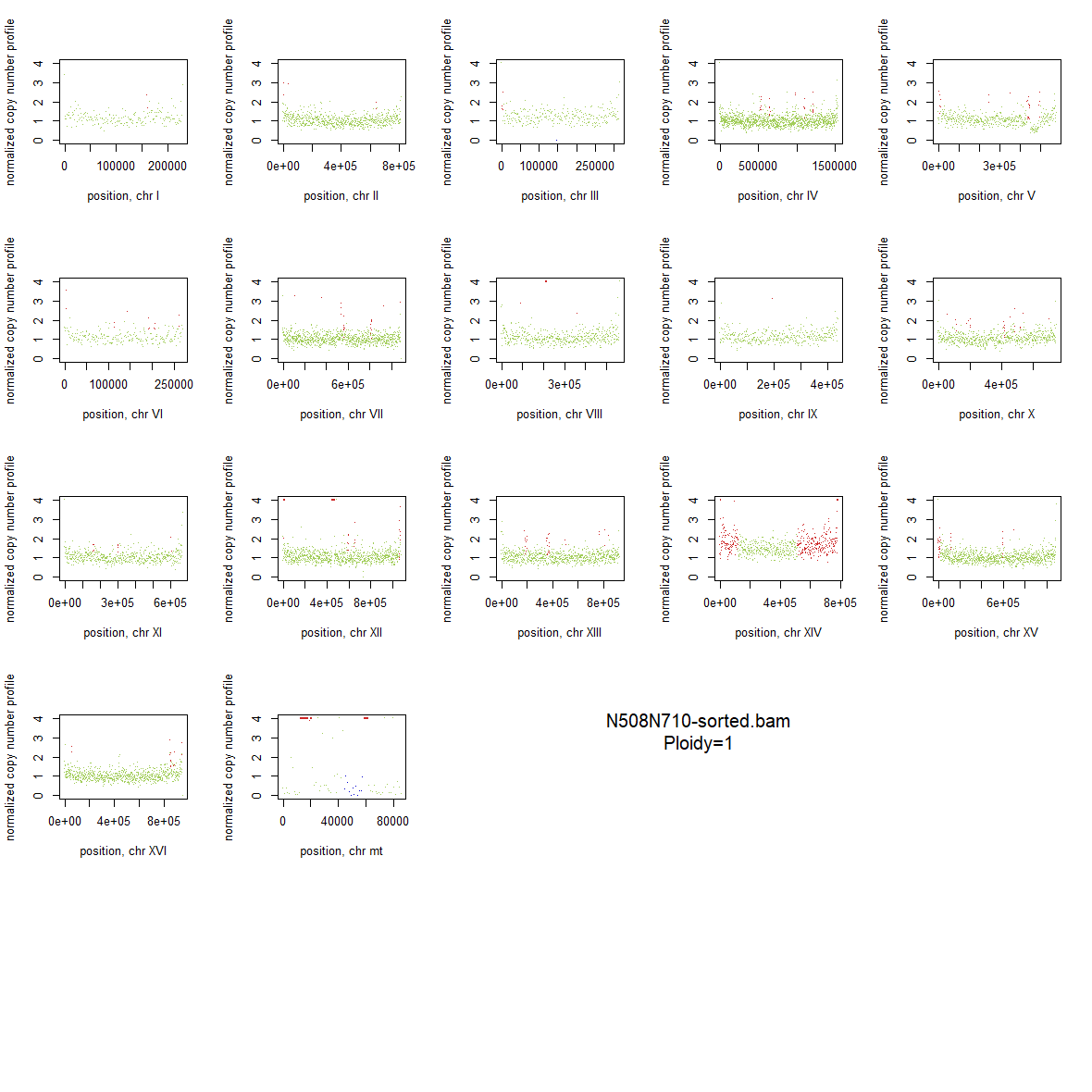

### 2x_V238M_81.png

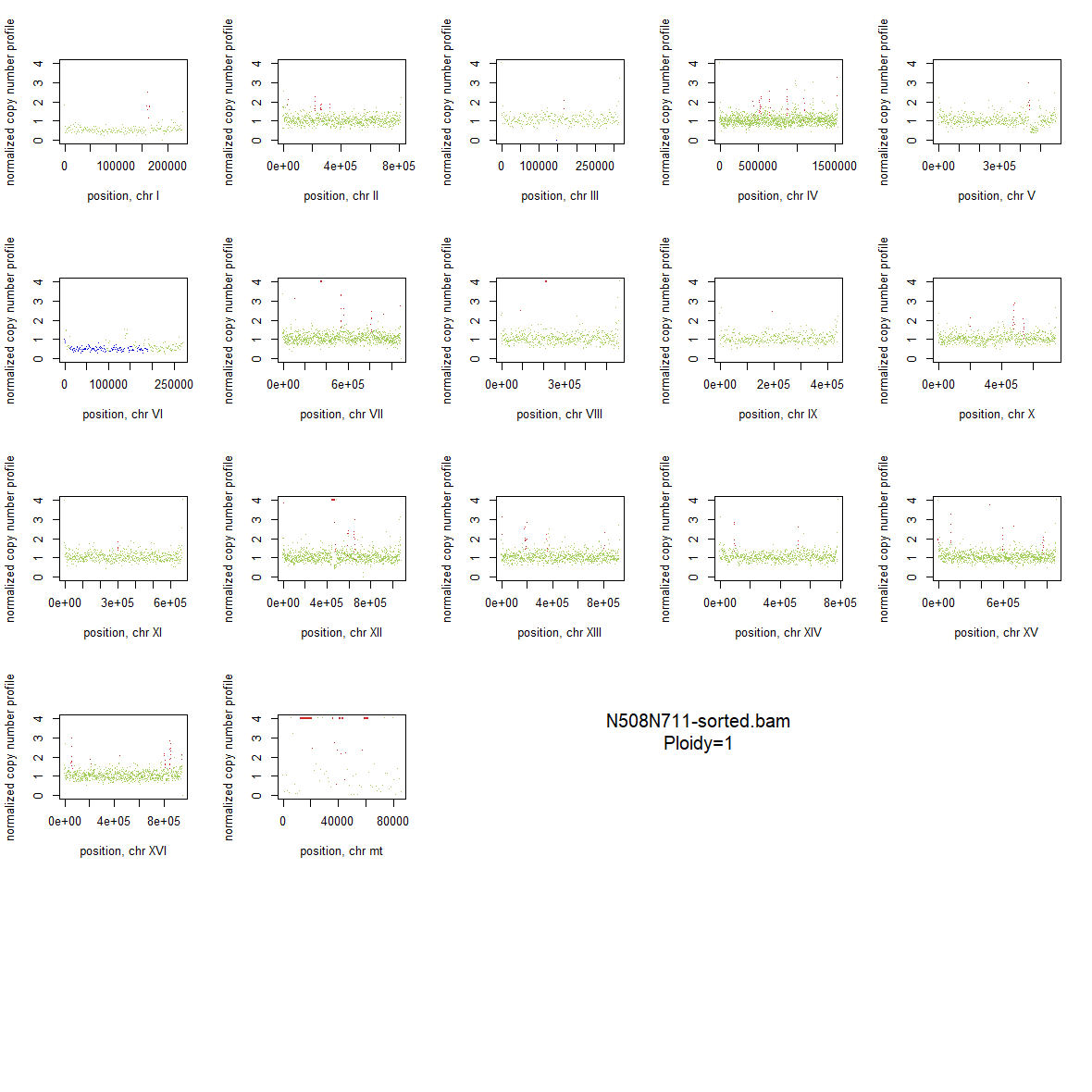

### 2x_V238M_82.png

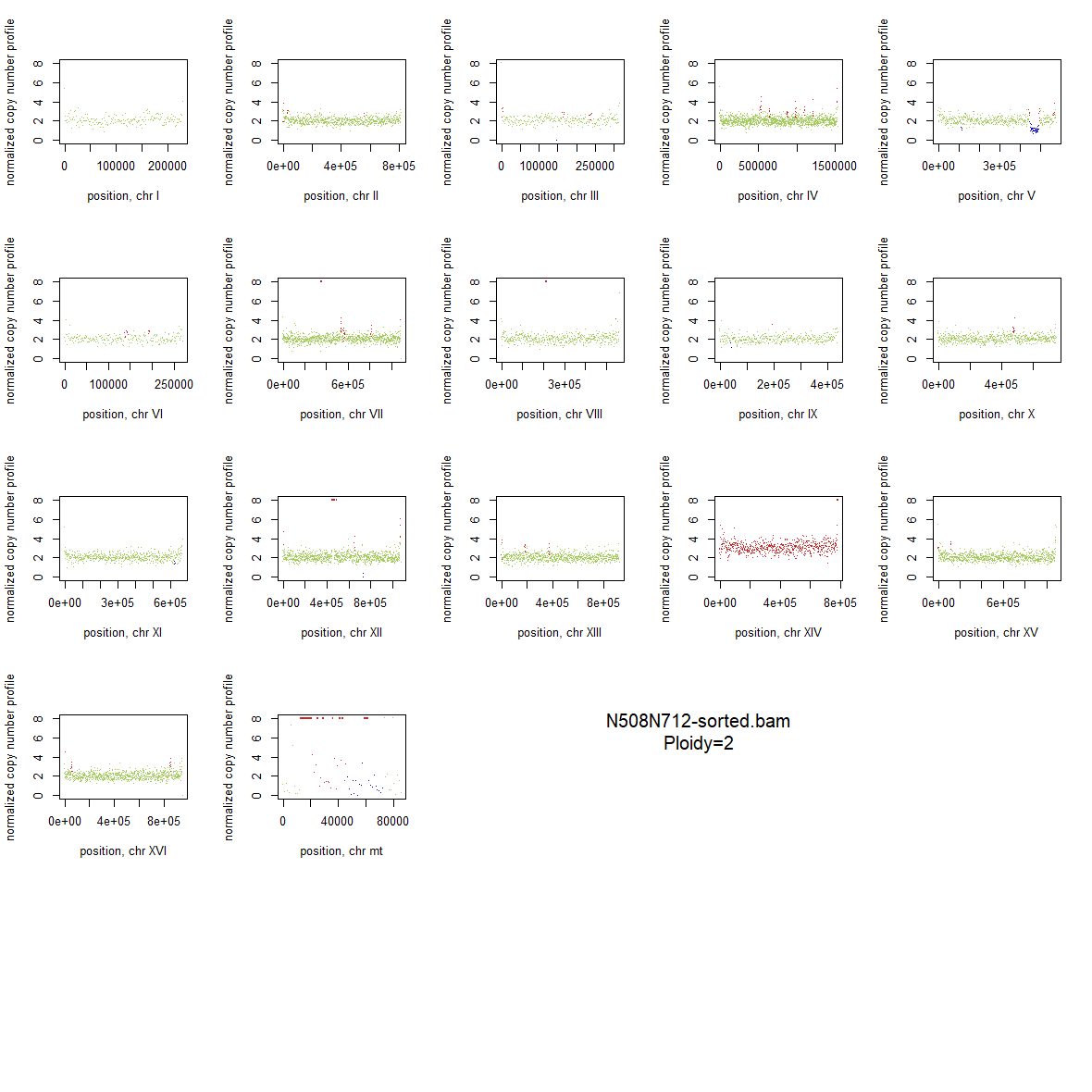

### 2x_V238M_83.png

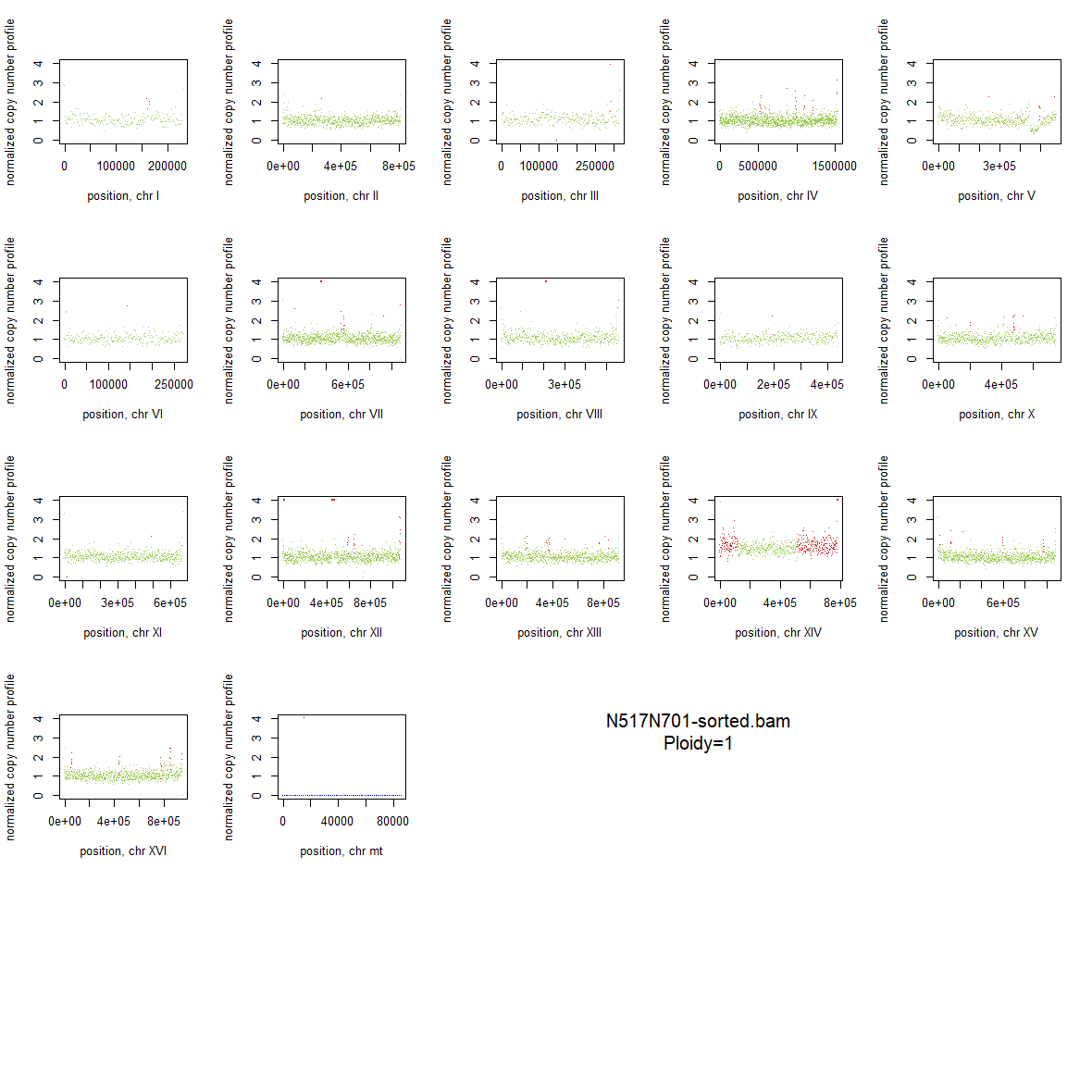

### 2x_V238M_84.png

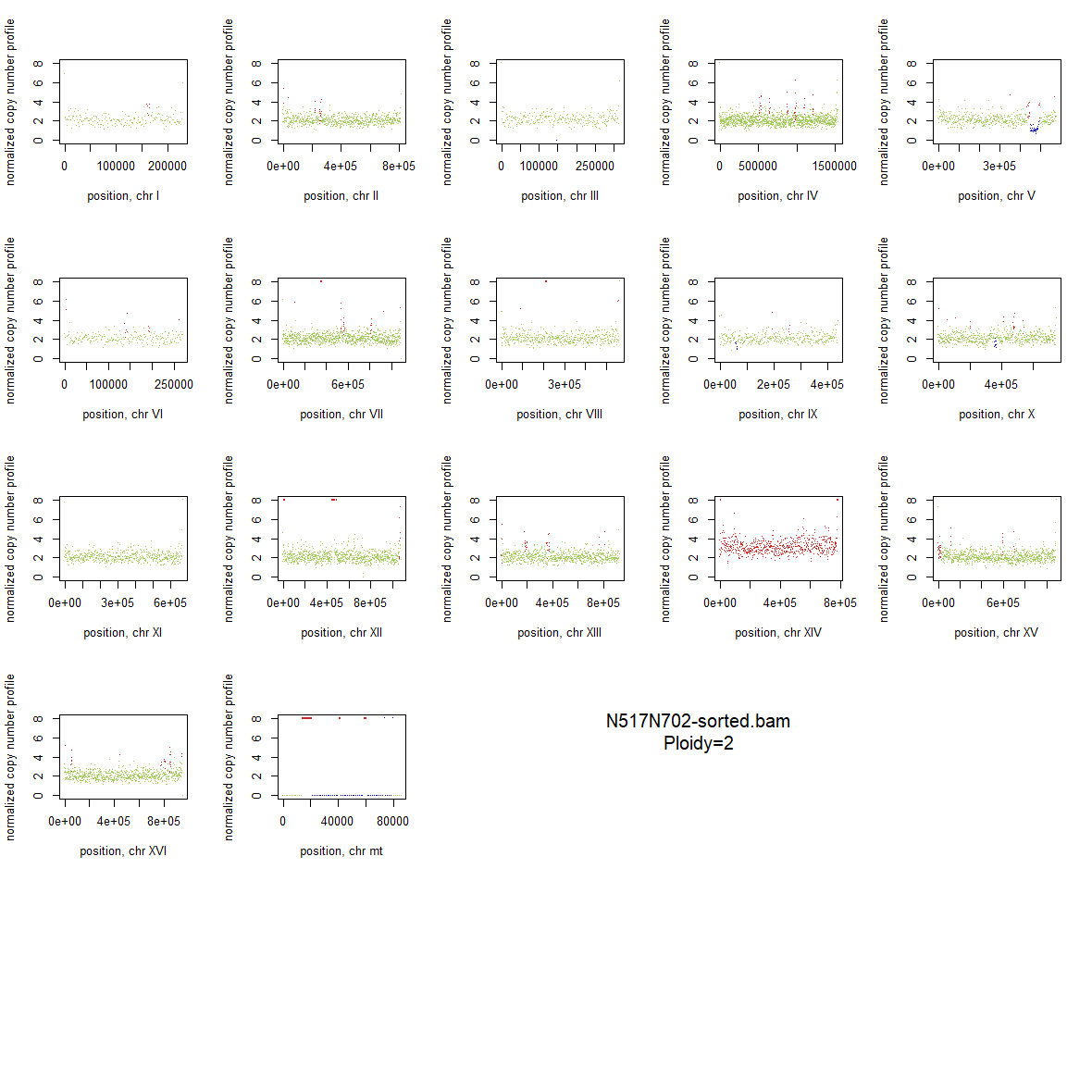

### 2x_V238M_85.png

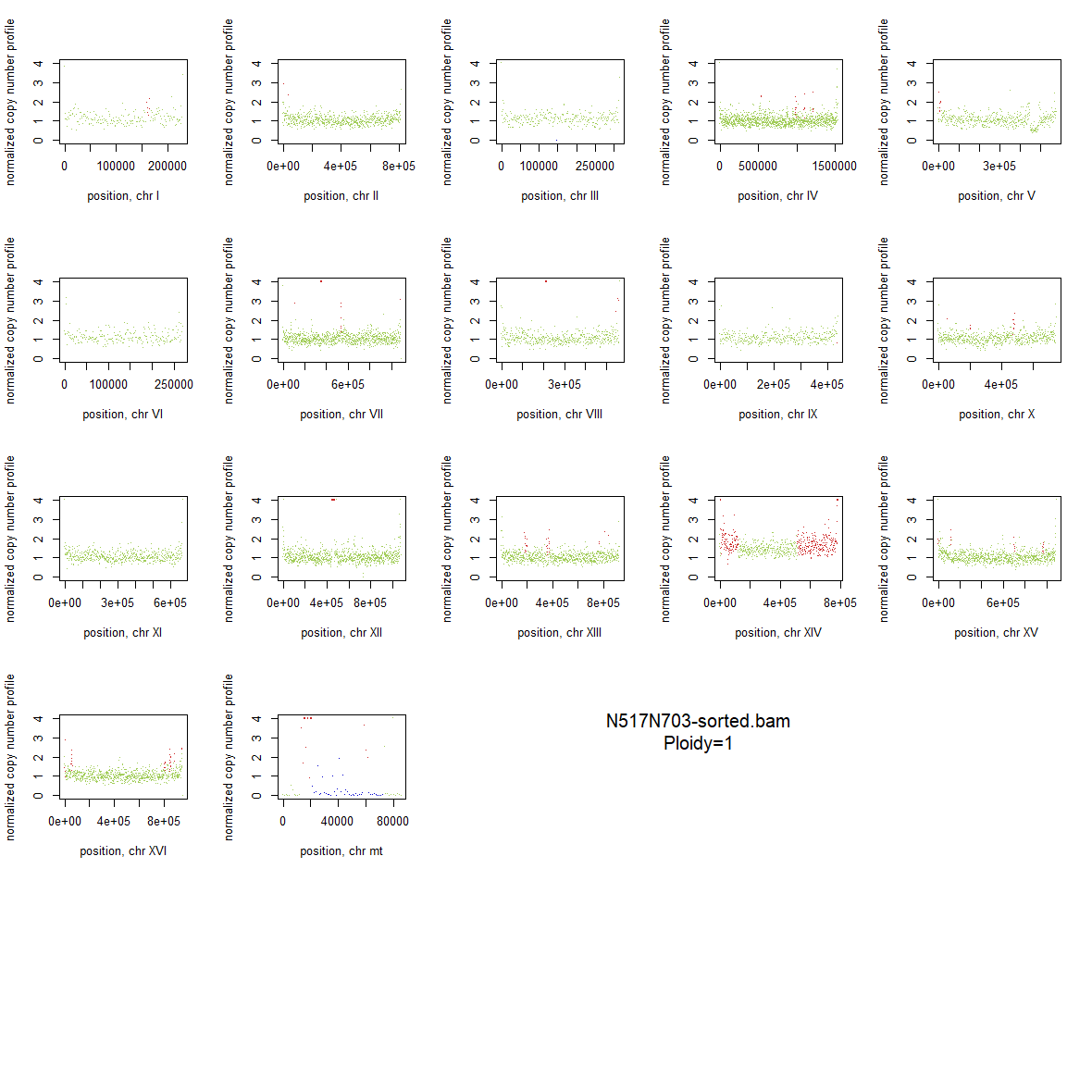

### 2x_V238M_86.png

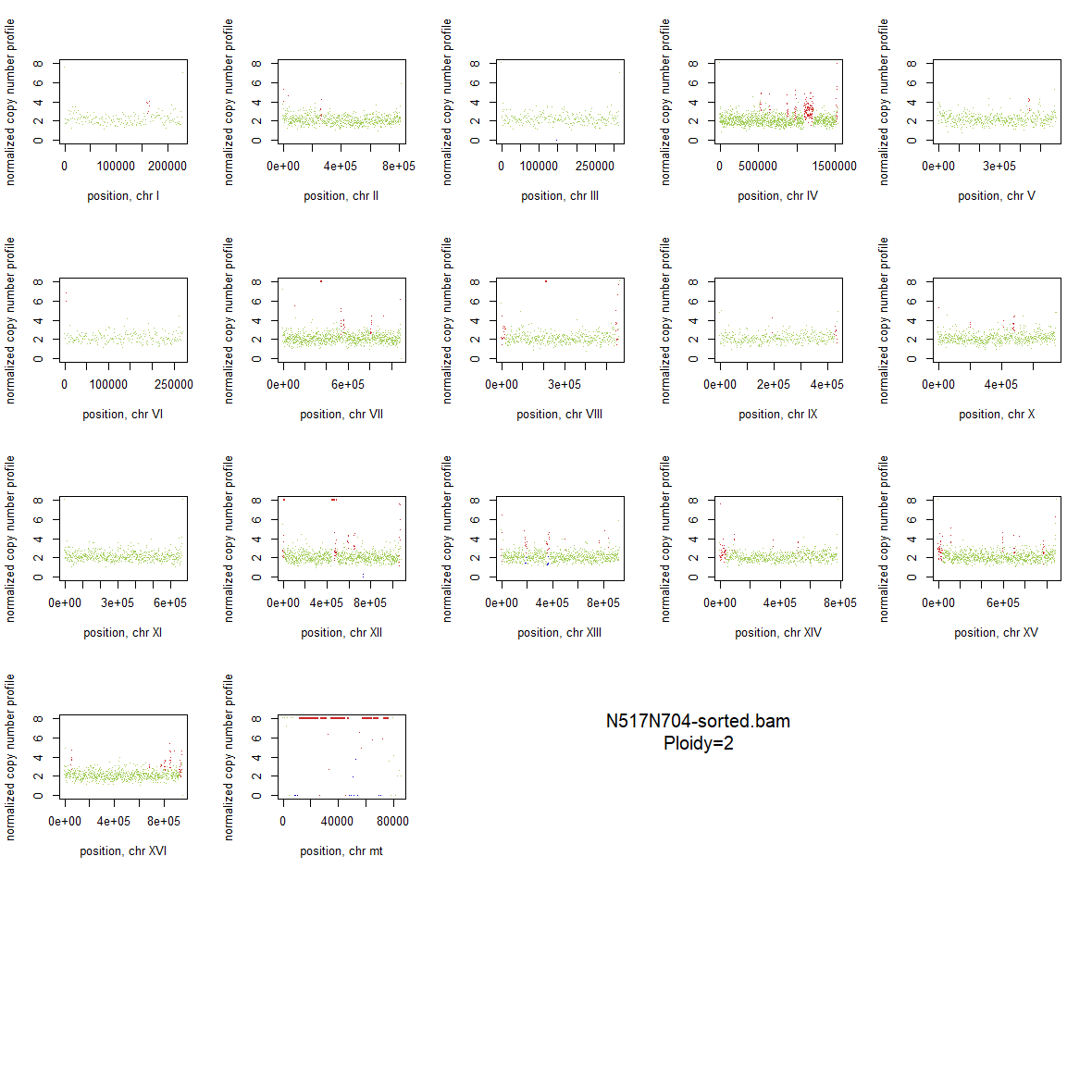

### 2x_V238M_87.png

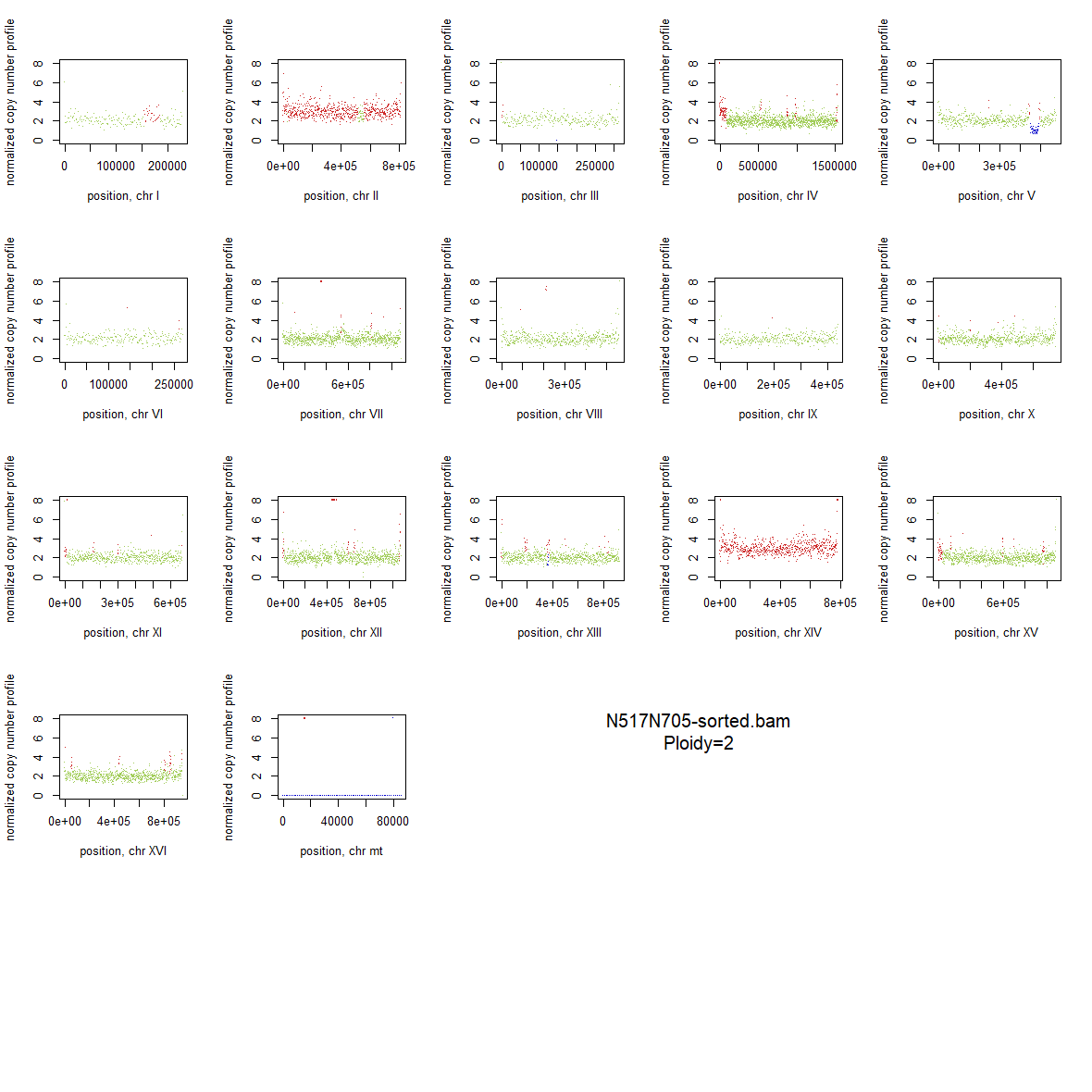

### 2x_V238M_88.png

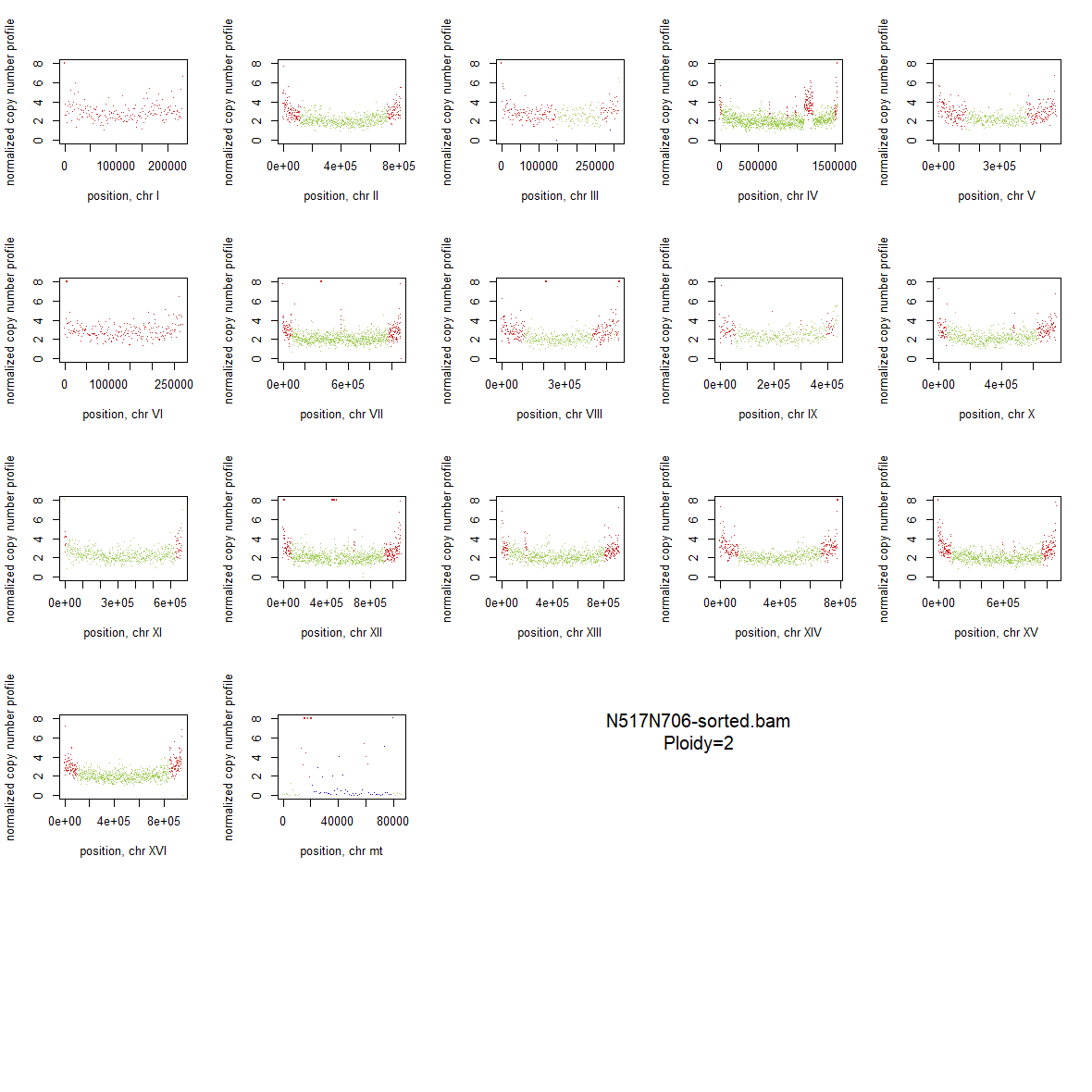

### 2x_V238M_89.png

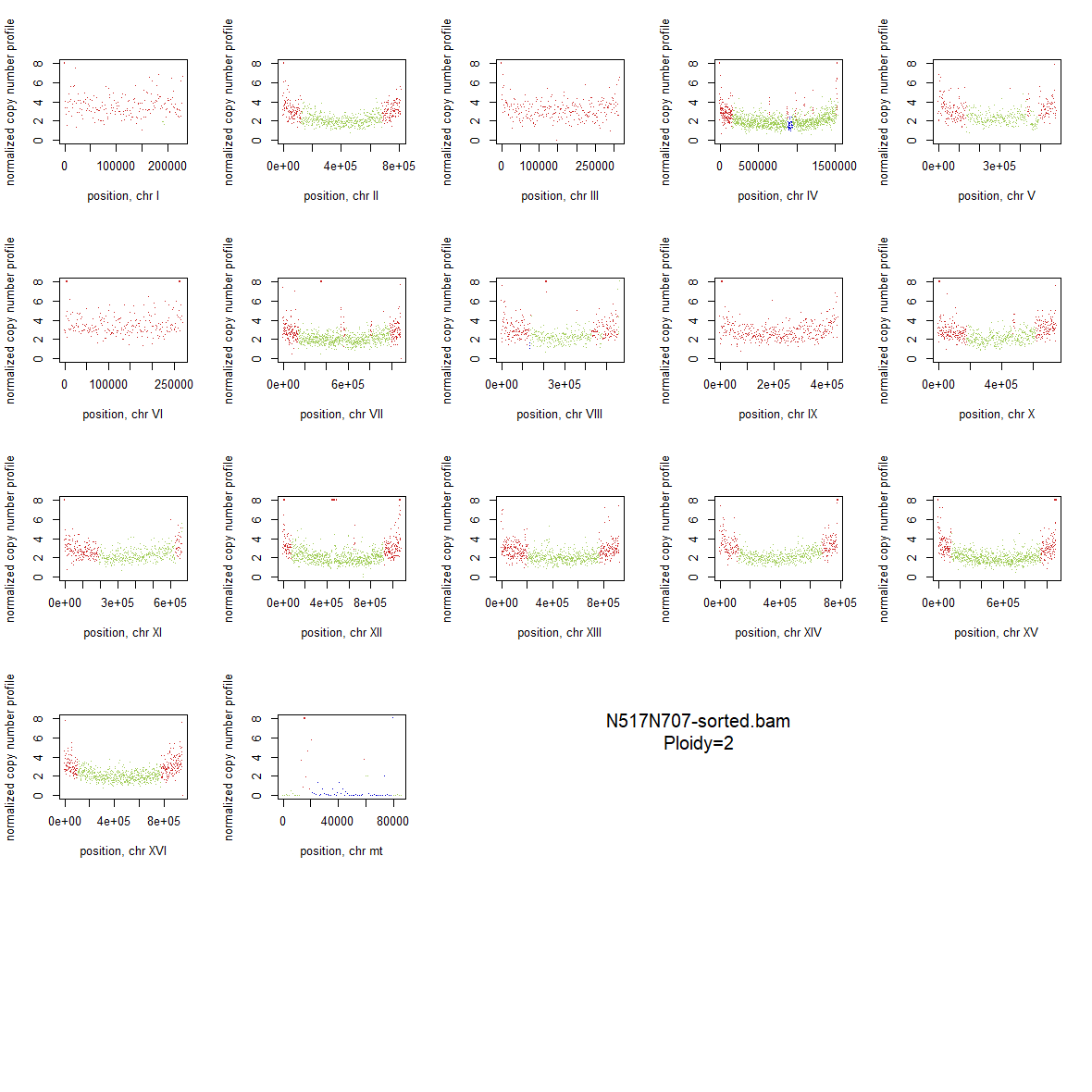

### 2x_V238M_90.png

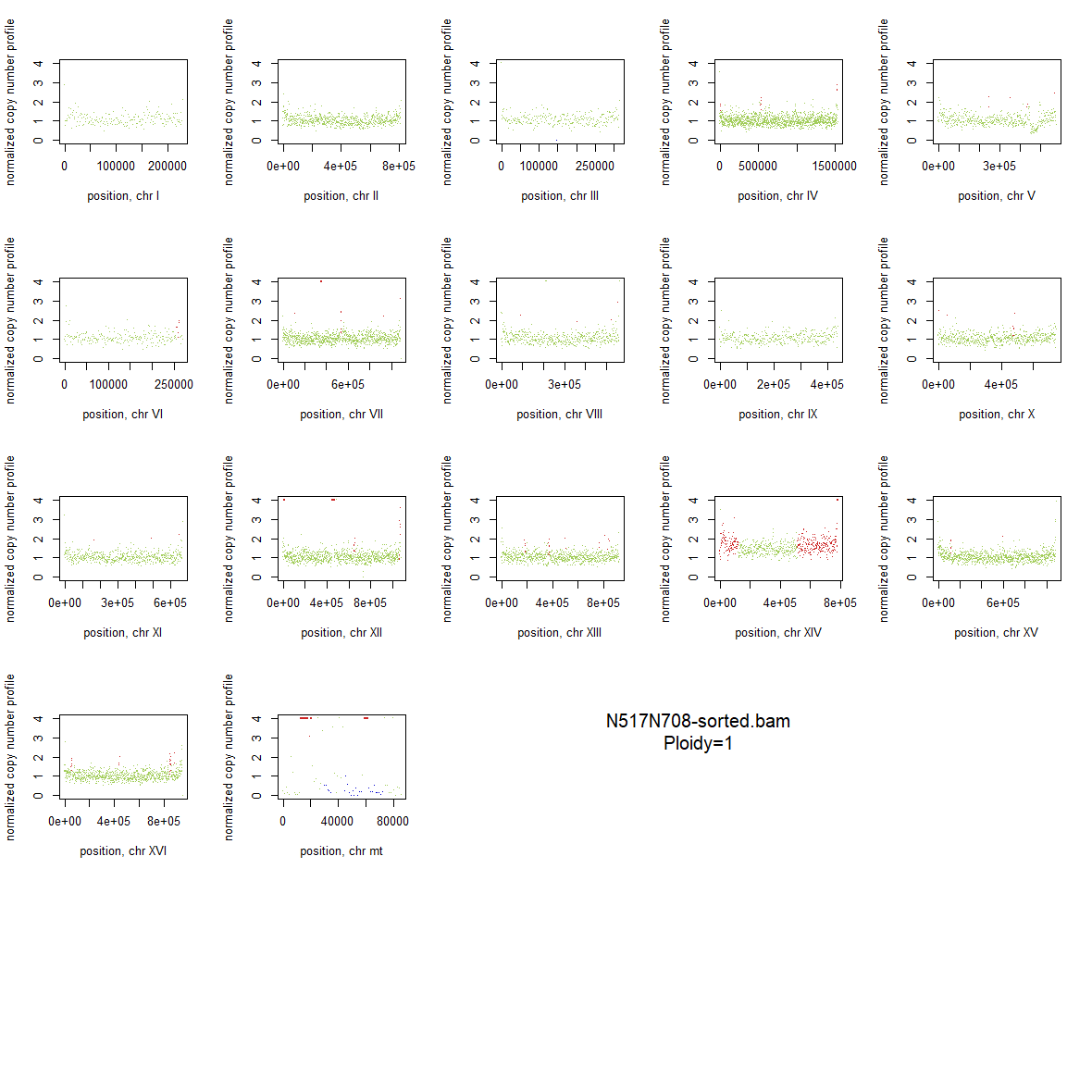

### 2x_V238M_91.png

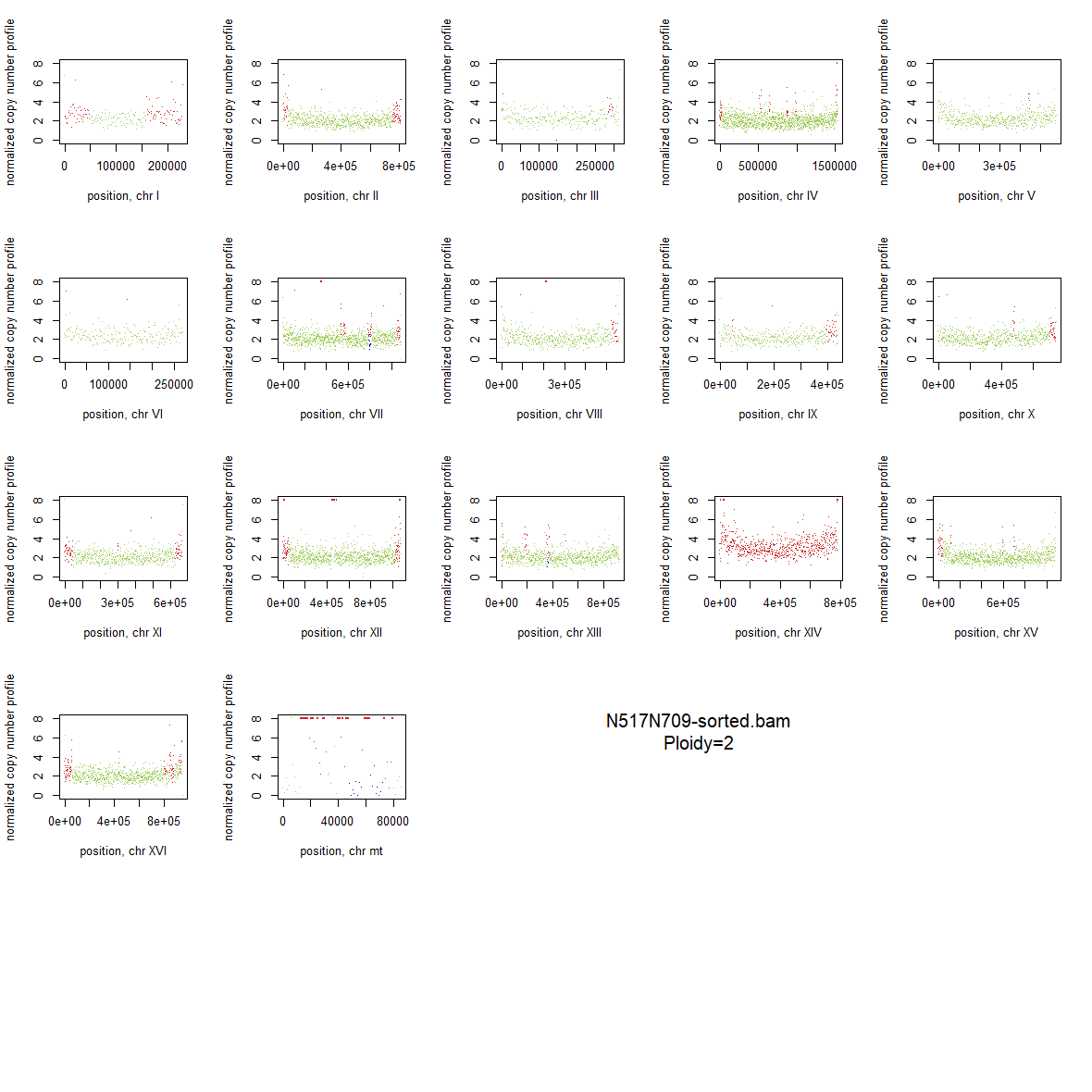

### 2x_Wildtype_01.png

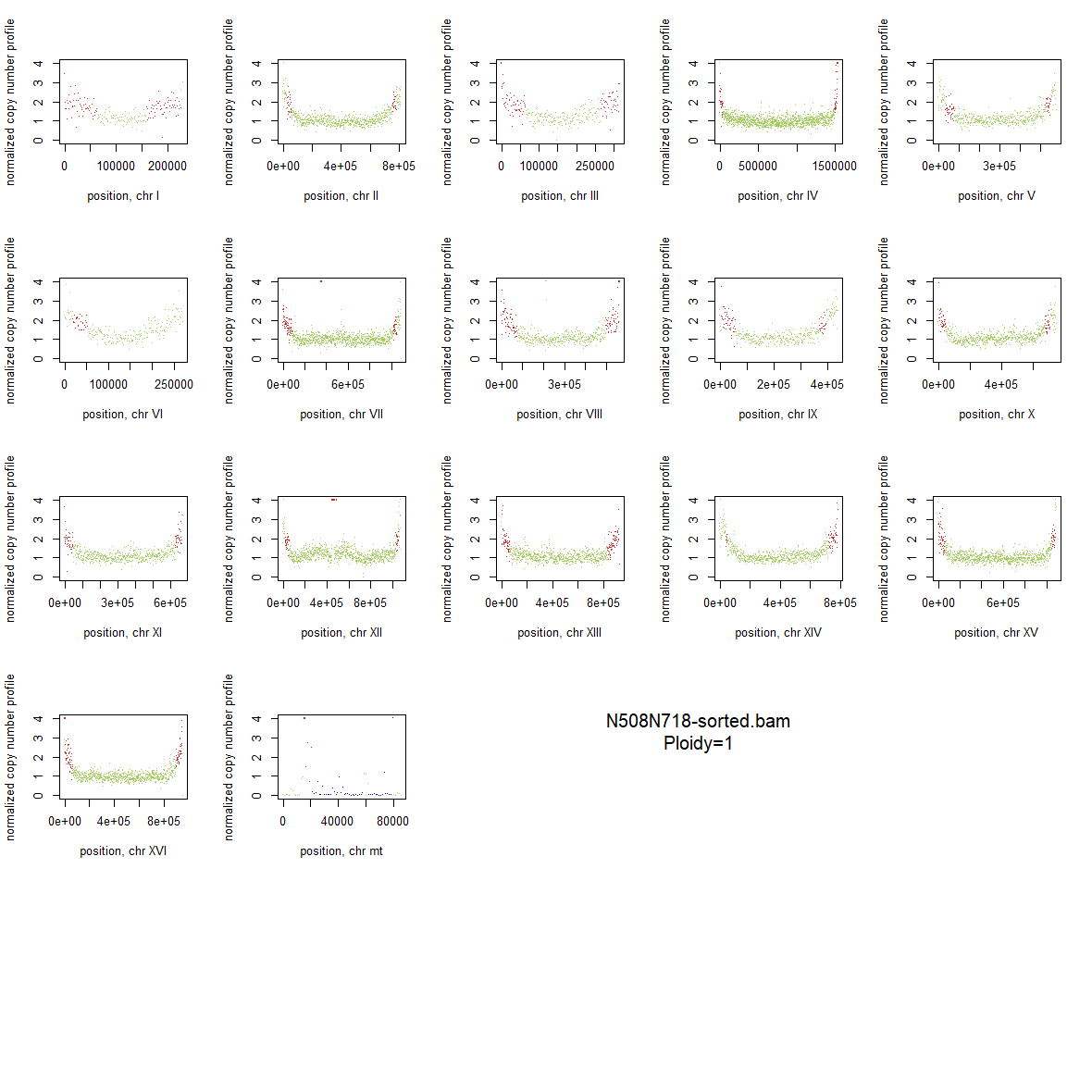

### 2x_Wildtype_02.png

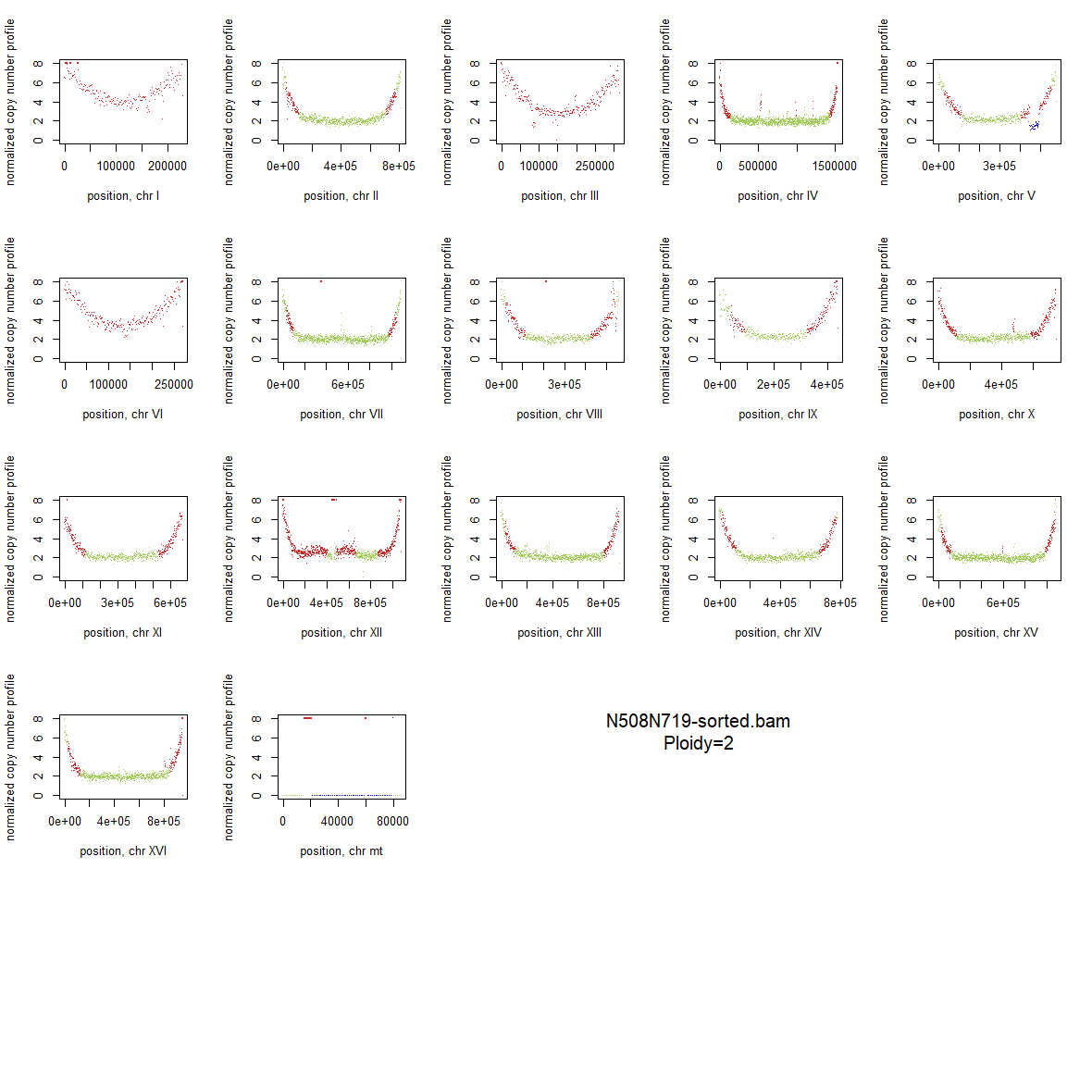

### 2x_Wildtype_03.png

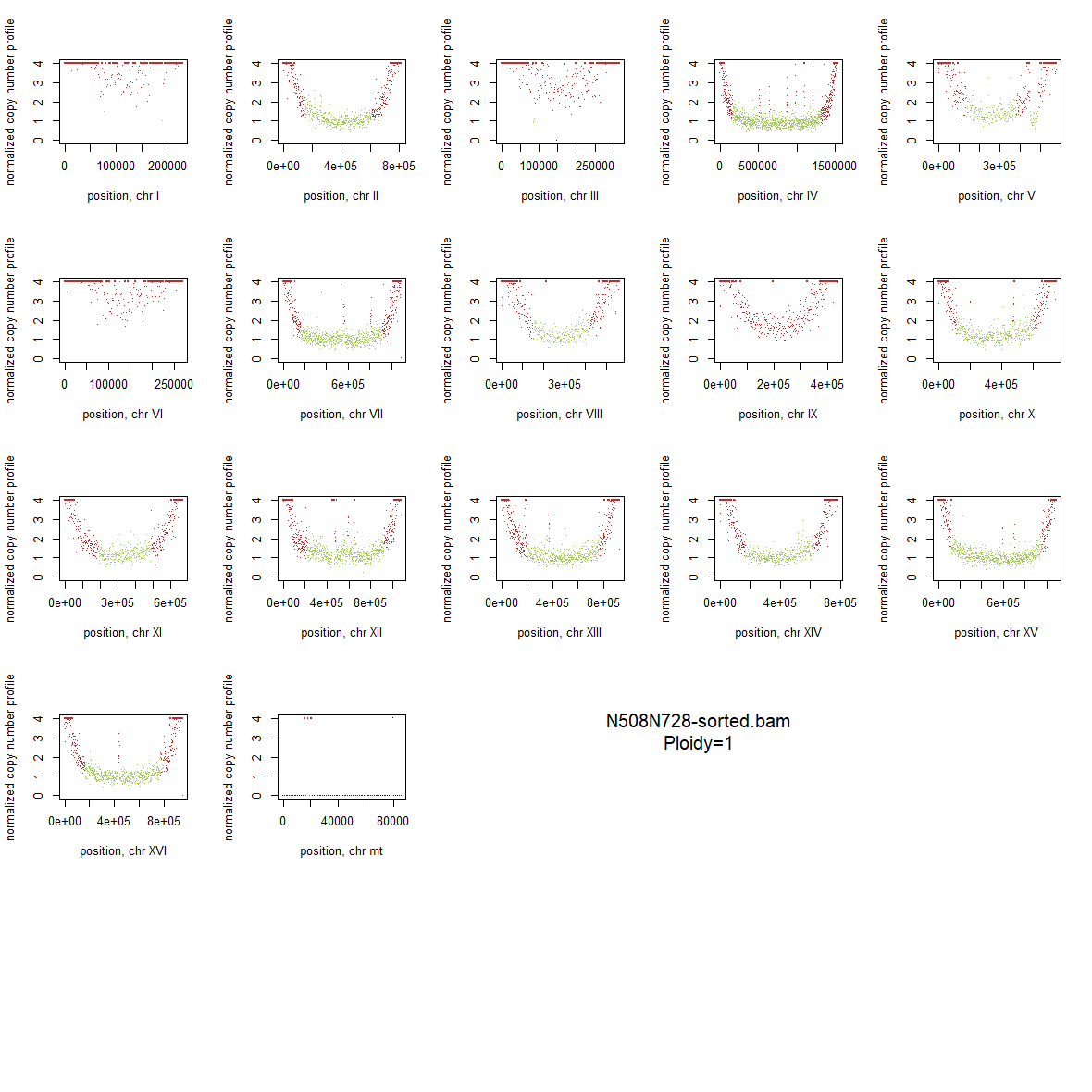

### 2x_Wildtype_04.png

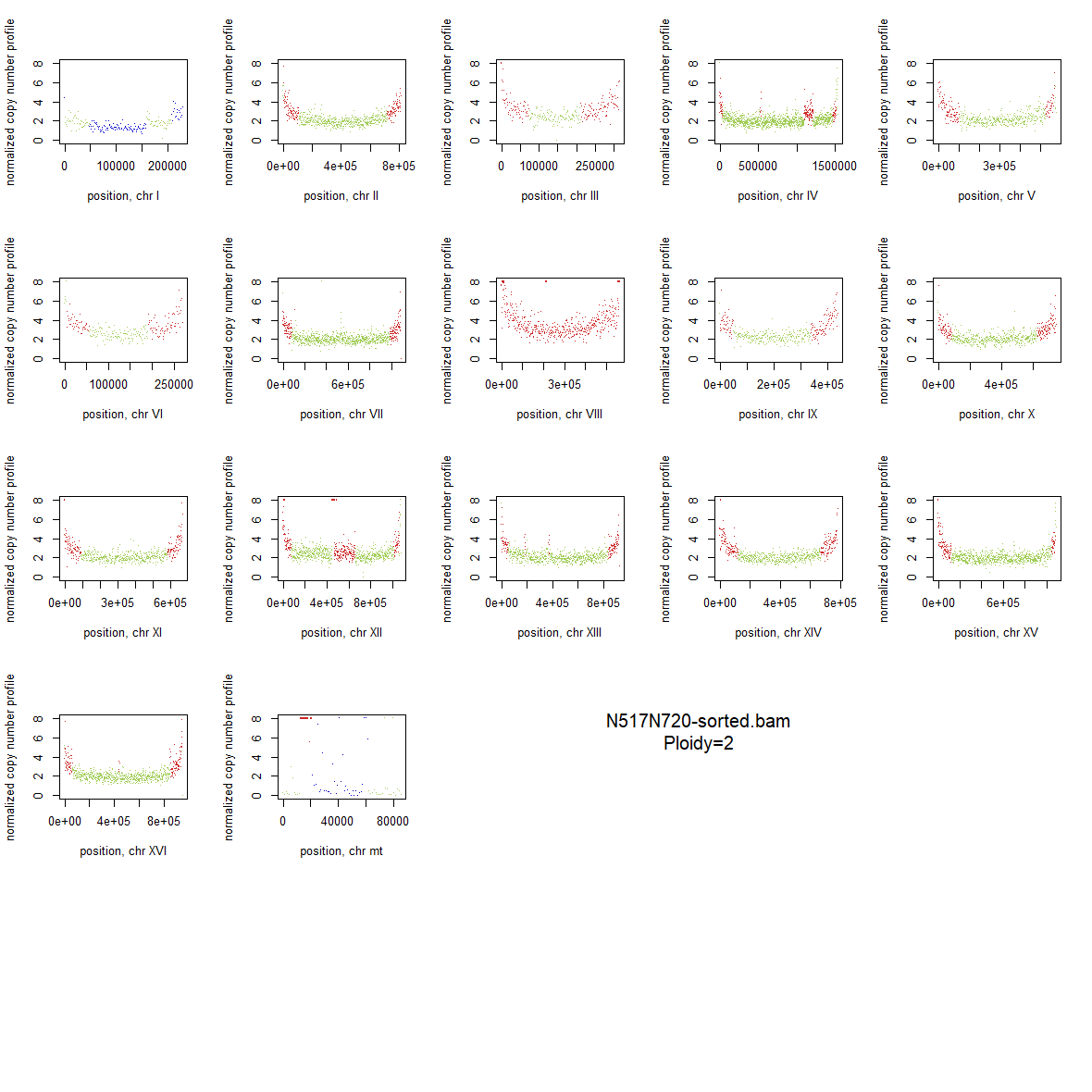

### 2x_Wildtype_05.png

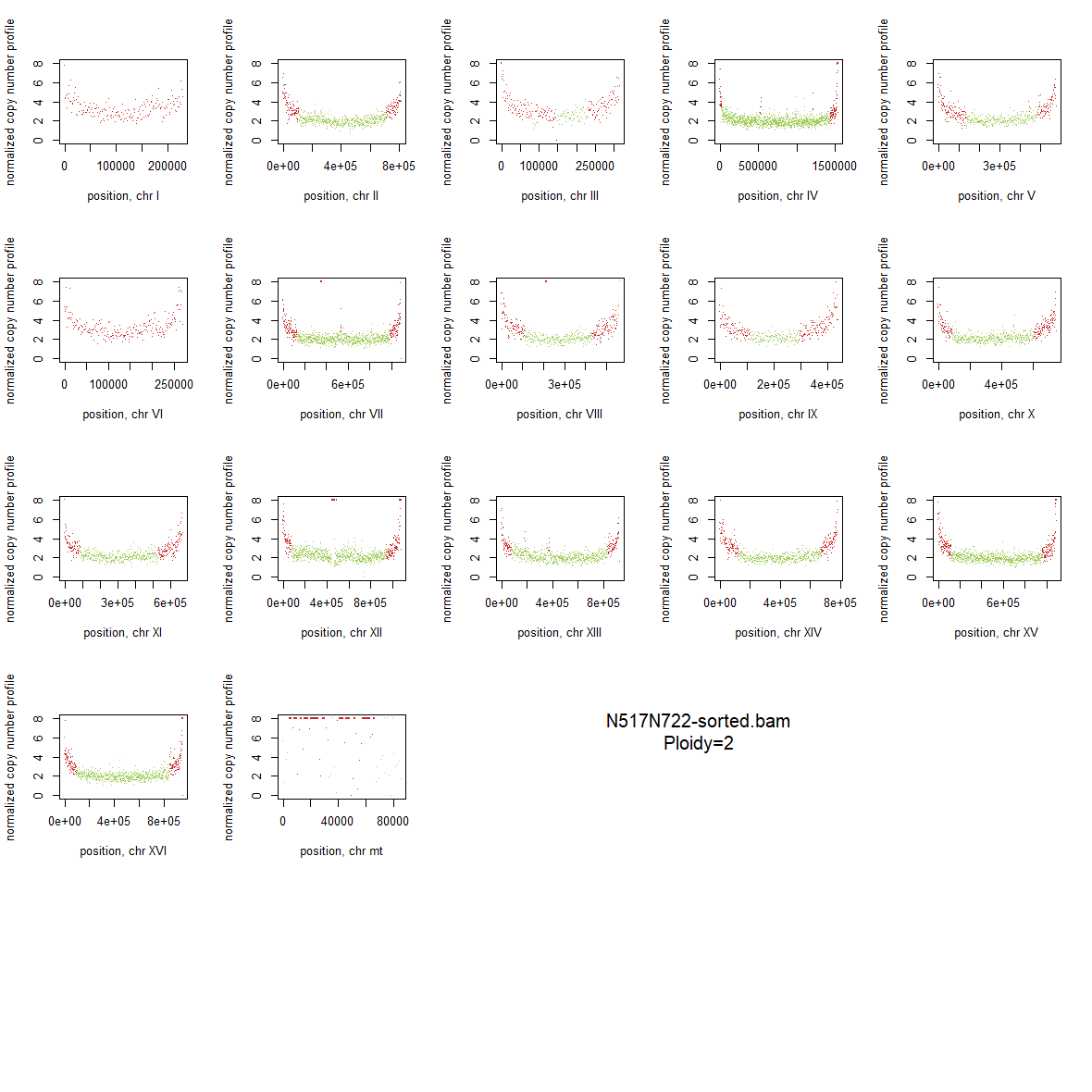

### 2x_Wildtype_06.png

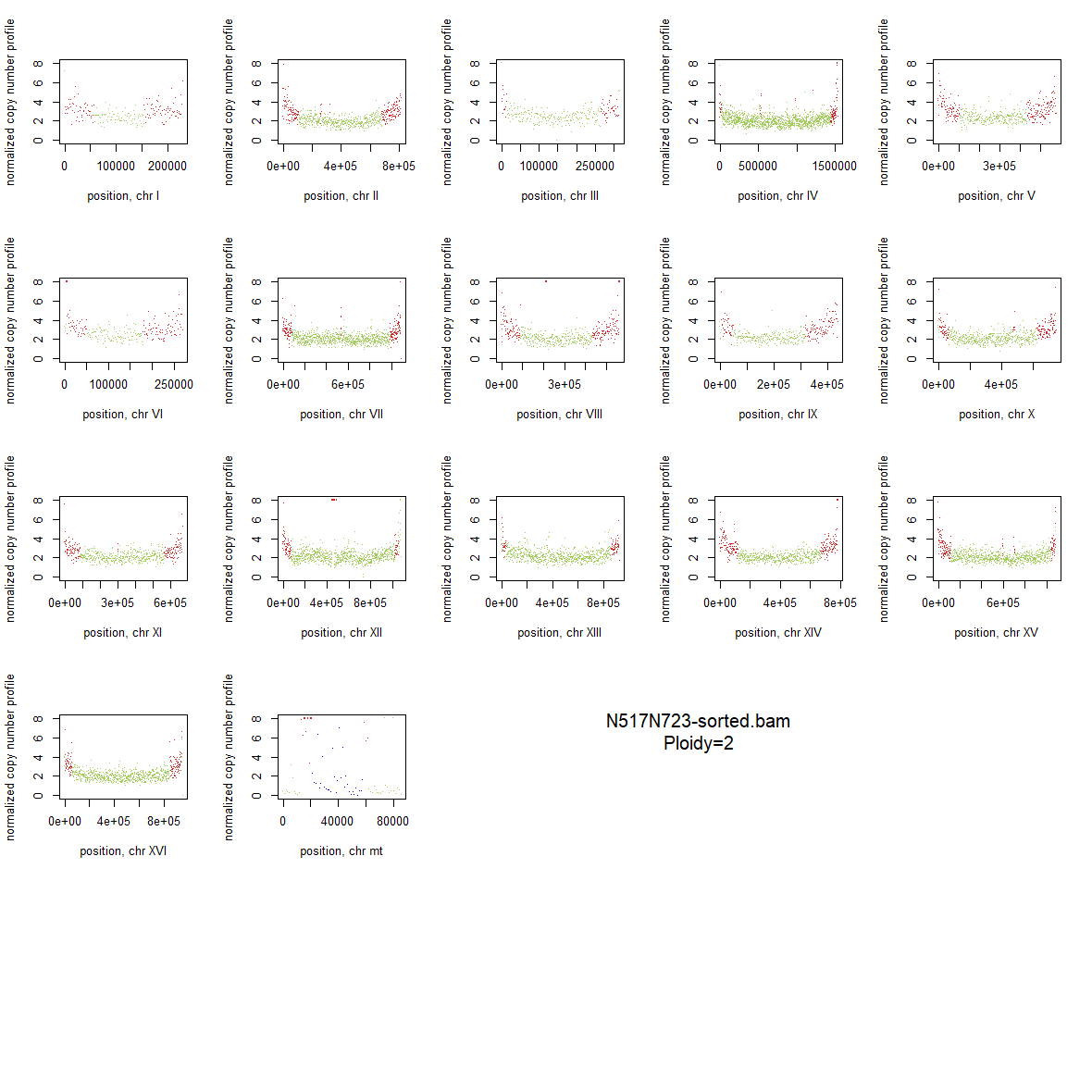

### 2x_Wildtype_07.png

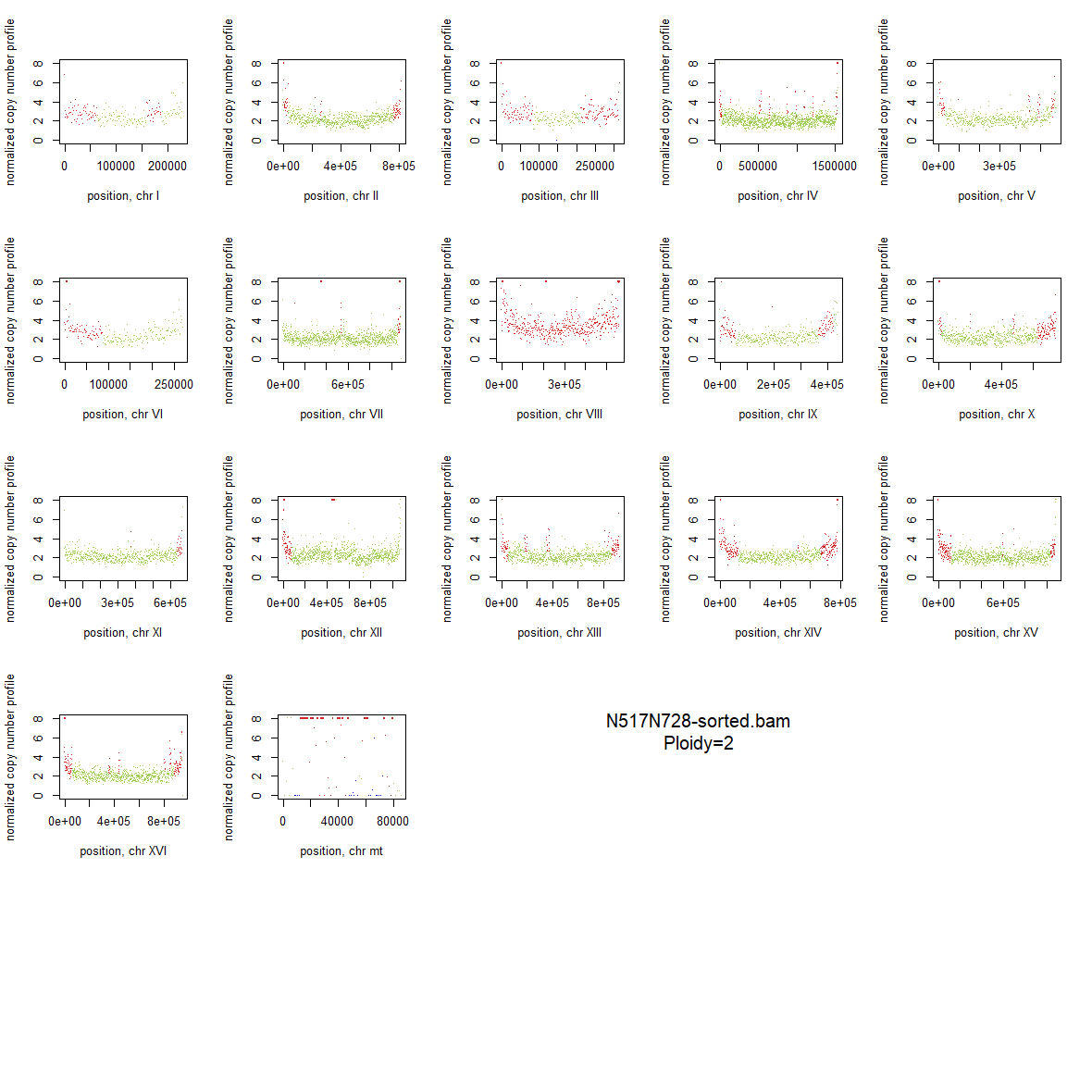

### 2x_Wildtype_08.png

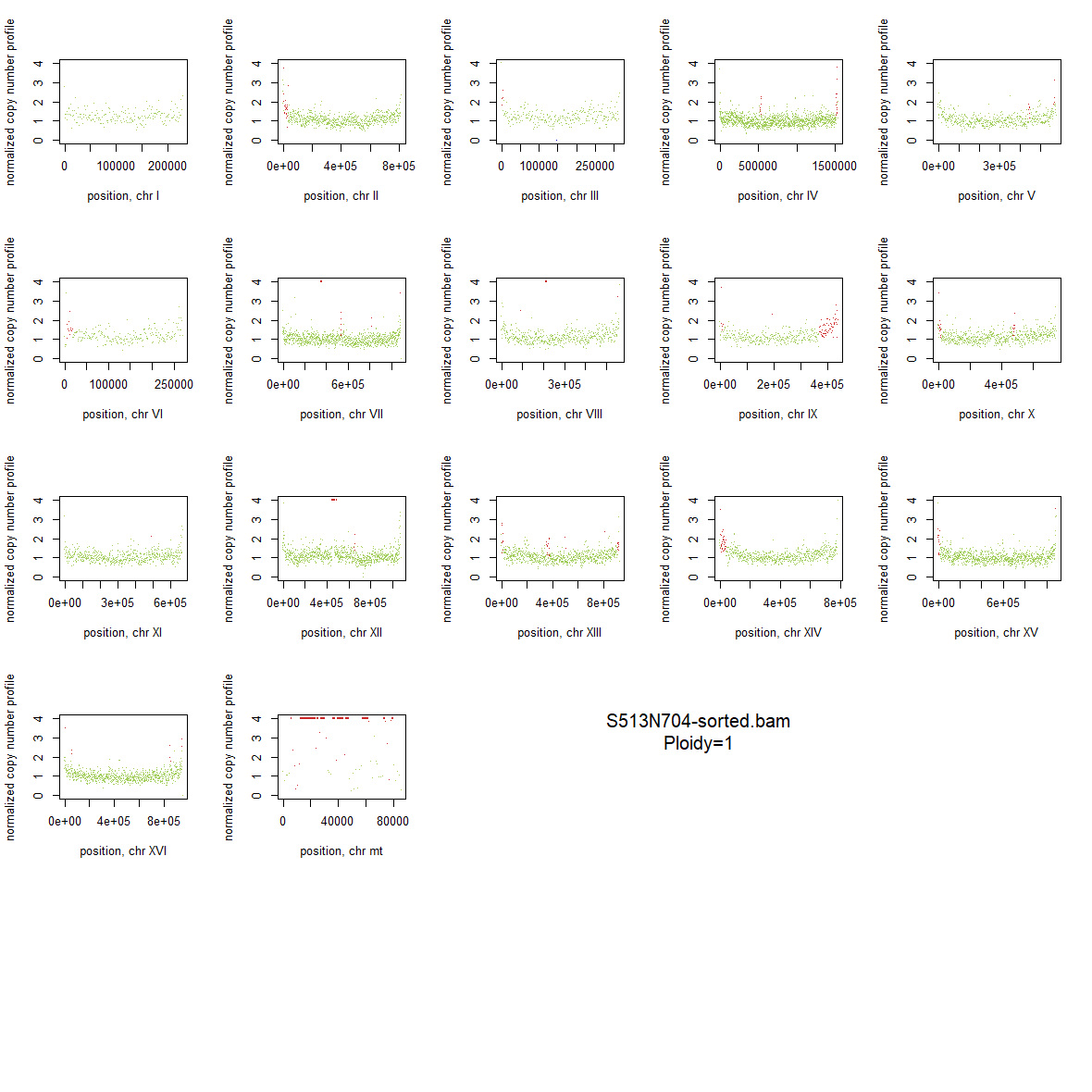

### 2x_Wildtype_09.png

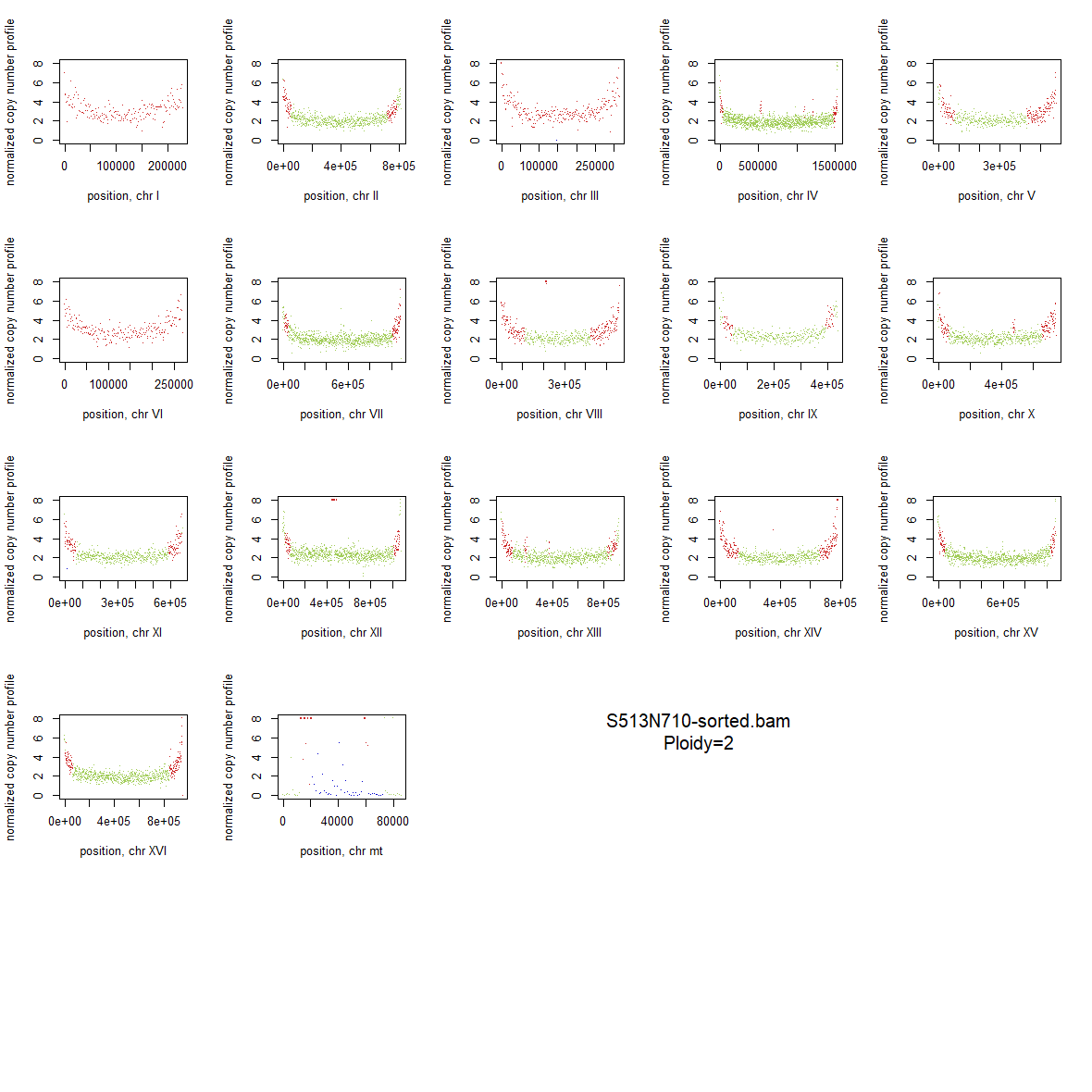
